# Transcriptome analysis reveals a novel DNA element that may interact with chromatin-associated proteins in *Plasmodium berghei* during erythrocytic development

**DOI:** 10.1101/2024.03.10.584310

**Authors:** Adaobi Okafor, Yagoub Adam, Benedikt Brors, Ezekiel Adebiyi

## Abstract

**Background:** The life cycle of *Plasmodium* parasites is intricate and multistage, alternating between dynamic environments. Temporal regulation of transcription by stage-specific transcription factor binding at particular regulatory regions within gene promoters facilitates its progression. As a result, each new developmental stage is endowed with its unique gene sets, whose just-in-time expression enables the parasite to completely adapt to the necessary circumstances. Our understanding of these transcriptome-level regulatory processes is limited, and more so, a thorough examination of the entire life cycle in the experimentally tractable rodent model organism *P. berghei* is lacking.

**Results:** We performed a genome-wide analysis of RNA-Seq data from different developmental stages of *P. berghei*. Integrated data from the human malaria parasites *P. falciparum* and *P. vivax* demonstrated that *Plasmodium* parasites have a unique transcriptional signature. We identified the sets of genes differentially expressed at each stage, clustered them based on similarities of their expression profiles, and predicted the regulatory motifs governing their expression. We interpreted the motifs using known binding sites for established eukaryotic transcription factors, including those of the ApiAP2s, and identified eight potentially novel motifs. Additionally, we expanded the annotation of another motif—AGGTAA—found in genes exclusive to erythrocytic development and identified members of the *Pf*MORC and GCN5 complexes among its possible interacting proteins.

**Conclusion:** This study provides new insights into gene usage and its regulation during *P. berghei* development.

## Background

Malaria, an acute fever illness, is caused by plasmodial infection [1]. The intricate life cycle of *Plasmodium* parasites is marked by multiple developmental stages and environmental transitions and includes sexual reproduction in female *Anopheles* mosquito vectors and asexual reproduction in vertebrate host tissues, particularly liver cells (hepatocytes) and red blood cells (erythrocytes) [2]. Temporal regulation of transcription by stage-specific transcription factor binding at particular regulatory elements upstream of genes facilitates the progression of the stage transitions [3]. The major family of stage-specific transcription factors, the Apicomplexan AP2s (ApiAP2s), first identified in the human parasite *Plasmodium falciparum*, play an essential role in the *Plasmodium* life cycle progression [4]. Protein binding microarray experiments also revealed the DNA binding preferences for 20 of the 27 *P. falciparum* ApiAP2 factors during the intraerythrocytic development cycle (IDC) [5]. However, the binding patterns of these factors across the entire life cycle stages remain unknown.

An important and often used rodent model organism in biomedical research is *Plasmodium berghei*. *P. falciparum*, has been the subject of the majority of studies on malaria, putting *P. berghei* somewhat behind. The entire genomes of *Plasmodium* parasites are now accessible for functional genomics research due to extensive genome sequencing efforts [6,7]. Furthermore, much effort has been made in the last 20 years to utilize RNA-Seq to analyse the bulk transcriptomes of human malaria parasites, with a primary focus on the blood stages and sporadic exploration of the mosquito stages (Additional file 1). Moral and ethical issues hamper performing in-depth studies of the whole *Plasmodium* life cycle in humans [8]. *P. berghei* is an option that provides experimental robustness and has a well-established and standardised genetic modification methodology for *in vivo* research [8,9]. A few studies have recently investigated the bulk transcriptome of *P. berghei*, focusing especially on individual analyses of the blood and mosquito stages (Additional file 1). Notably, the two most recent transcriptome studies throughout liver stages were made possible by the *P. berghei* model, resulting in data that now cover the whole life cycle [8,10]. Both studies examined *P. berghei* gene transcription using extensive available data, but their primary areas of interest were the liver stages. Moreover, only one [10] examined transcriptional regulation and also restricted the scope of their research to the liver stages. Therefore, a deeper understanding of gene transcription and transcription control across the entire life cycle is lacking, hence, we performed a genome-wide analysis of RNA-Seq data from different developmental stages of *P. berghei* to dissect gene usage and the interplay between regulatory elements and transcription factors controlling the life cycle progression.

## Methods

### Dataset

Transcriptome data comprising most of the life cycle stages of *P. berghei*, *P. falciparum* and *P. vivax*, namely, salivary gland sporozoite; liver stages; ring, trophozoite, schizont (or collectively known as asexual blood stages); gametocyte and ookinete, generated from two independent studies (Additional file 2), were downloaded as Sequence Read Archive (SRA) files from the public archives at the National Center for Biotechnology Information (NCBI) using sratoolkit version 2.11.0 [11]. The SRA files were converted into FASTQ files using ‘fastq-dump’ within the sra-toolkit. Reference genome FASTA files and associated gene annotation GTF files for each genome [accession numbers: ASM276v2 (*P. falciparum*), *PB*ANKA01 (*P. berghei*) and GCA 900093555 (*P. vivax*)] were also downloaded directly from the Ensembl Protists genome browser (release 50) [12].

### Data preprocessing

All the raw read sequences were checked for quality with FASTQC version 0.11.9 [13]. Sequencing adapters, low-quality reads and short reads were trimmed off using Trimmomatic version 0.39 [14] with the following parameters: ILLUMINACLIP: TruSeq2 (SE) or TruSeq3 (PE) or Nextera (PE) LEADING: 3 TRAILING: 3 SLIDINGWINDOW: 4:20 MINLEN: 20. The read sequences with a minimum length of 20 nucleotides and whose bases had a minimum Phred quality score of 20 were retained. Trimmed read sequences were rechecked with FastQC and aligned to their respective reference genomes and gene annotation files (Ensembl Protists release 51) using STAR version 2.7.8a [15] with the following parameters: -sjdbOverhang (“read length” – 1) - limitBAMsortRAM 50000000000. The strand specificities of the read sequences were confirmed prior to read quantification using the tool “how are we stranded here” version 1.0.1 [16]. To quantify the aligned reads, the reads that only uniquely mapped to exonic regions in the alignment files were counted using ‘featureCounts’ from the Subread package version 2.0.1 [17] with the following parameters: -s Stranded (Reverse) or Unstranded -p (for PE reads) -t exon -g gene id.

### Differential gene expression analysis

Differential gene expression analysis was performed with DESeq2 version 1.22.2 [18]. The read counts for *P. berghei*, *P. falciparum* and *P. vivax* were integrated into a single matrix by transforming the *P. falciparum* and *P. vivax* genes by orthology into *P. berghei* gene ids using the BioMart tool in the Ensembl Protists genome browser (release 51), which included only the genes with 1:1:1 orthologues across the three species. A Spearman correlation test with hierarchical clustering was performed on the 500 most variable genes to evaluate the correlation of the *P. berghei* datasets with the *P. falciparum* and *P. vivax* datasets. Batch effects in the datasets were corrected using ‘removeBatchEffect’, implemented in limma [19] prior to dimensionality reduction by principal component analysis (PCA) on the 500 most variable genes. The raw counts for the entire *P. berghei* dataset were analysed for differential expression using DESeq2 version 1.22.2, which included the parasite stages in the design formula as the variable of interest. Each stage was compared with all other stages, and the results were generated, including the log2-fold changes, p values and FDR-adjusted p values that control for multiple testing [20]. The differentially expressed genes (DEGs) were then filtered out (padj < 0.01, abs(log2FC) > 2). Normalized read counts for use in subsequent clustering analysis were also extracted with DESeq2 version 1.22.2. Furthermore, for each parasite stage, the DEGs that were consistent in the results of the analysis of the datasets from the corresponding two independent studies were filtered out for use in downstream analysis and visualized with a Venn diagram [21]. Plots were obtained using the ggplot2 R package version 3.4.2 [22], and all heatmaps relating to the gene expression analysis were obtained using the ComplexHeatmap R package version 2.16.0 [23].

### Functional annotation

Gene Ontology (GO) terms related to biological processes for the upregulated genes (release 43) and genes in the coexpression clusters were retrieved from PlasmoDB (release 51) [24]. To interpret unknown motifs, their enriched GO terms for biological processes, cellular components, and molecular functions, as well as enriched KEGG pathways, were also retrieved from PlasmoDB (release 65). The gene ontology terms were summarised with REVIGO version 1.8.1 at medium (0.7) allowed similarity using GO term sizes from *P. falciparum* and semantic similarity measures from SimRel [25].

### Coexpression clustering analysis

For the identification of groups of genes with similar expression profiles, PAM analysis with hierarchical clustering was performed on scaled, normalized counts using the ComplexHeatmap R package version 2.16.0. The optimal number of clusters of similarly expressed genes in the dataset was determined using the factoextra R package version 1.0.7 [26] employing the elbow and silhouette methods.

### *De novo* motif discovery

To identify putative regulatory sequences underlying the observed coexpression patterns, the promoter regions of the genes in each coexpression cluster were searched for overrepresentation of sequence motifs. First, the 5’UTR sequences 1000 bp upstream of the start codons of the genes were extracted from PlasmoDB (release 51). Motif analysis was performed with the ‘oligo analysis’ tool implemented in RSAT version 2020-02-03 [27] with default parameters, background model (expected frequency of each oligonucleotide) estimated from *Plasmodium berghei*.*PB*ANKA01.34 genome, number of pattern assemblies set to 50 and matrix clustering turned off, and with XSTREME from the MEME Suite version 5.5.4 [28] with default universal and STREME parameters and MEME expected motif site distribution set to ‘any number of repetitions’. For each coexpression cluster, the resulting motif collections from both approaches were combined into a single collection. To ensure uniformity, the MEME motif collections in .meme format were converted to .transfac formats compatible with RSAT using the ‘convert matrix’ tool from RSAT with default parameters, selecting the *Plasmodium berghei*.*PB*ANKA01.34 genome from the list of provided organisms. Each transfac collection was clustered using the ‘matrix-clustering’ tool from RSAT with default parameters to group similar motifs together and determine a root motif representing the motifs in each collection, thus reducing the redundancy of the motifs and simplifying the interpretation of motif discovery results.

### Biological interpretation of the discovered motifs

The similarity of the discovered motifs to known motifs was determined by comparison to motifs in the JASPAR database [29] and transcription factor-binding sites discovered in *P. falciparum* through protein binding microarrays (*PB*M) [5] and assay for transposase-accessible chromatin using sequencing (ATAC-Seq) [30]. The TOMTOM tool from the MEME Suite was used with the following parameters: Euclidean distance as the comparison function, e-value less than 1 as the significance threshold and complete scoring turned off to ensure that only the aligned motif columns were considered in computing the comparison function. For compatibility, the .transfac motif collections from RSAT were converted to .meme format using the command ‘transfac2meme’ from the MEME Suite with default parameters.

### Promoter architecture

To visualize the promoter architectures of *P. berghei* genes with suspected related functions in *P. falciparum*, the 5’UTR sequences 1000 bp upstream of the start codon of their orthologues were queried for the individual genomic locations of the regulatory motifs of interest within them using ‘matrix-scan’ in RSAT with default parameters. The scan result or “feature list” was visualized using the ‘feature map’ tool within RSAT with default parameters.

### Gene regulatory network construction

To identify functional relationships that may exist between motifs and their target genes, the target genes harbouring the motifs were identified using ‘matrix-scan’. The gene regulatory network of the interactions of the motifs with their target genes was visualized using Cytoscape version 3.10.0 [31], and the numbers of unique and shared target genes among the motifs were visualized with a Venn diagram.

### Motif activity inference

Motif activity scores were calculated using an adaptation of the motif activity response analysis (MARA) model [32]. Read counts were used as a proxy for promoter expression levels [33], and the number of sites (genomic locations) of the motifs was predicted using the ‘matrix-scan’ tool. The temporal motif activity profiles were scaled and visualized as a line plot.

### Cross-correlation analysis

Cross-correlation analysis was performed for the blood stages (ring, trophozoite and schizont) on ring stage-delayed transcription factor expression data [7] and leading motif activity profile data after scaling to identify transcription factors expressed in the ring stage that could subsequently bind to motifs in the trophozoite and schizont stages. Pearson correlation with hierarchical clustering (Euclidean distance, “complete” method) was applied to the time series data and visualized with a heatmap using the Heatmaply package version 1.4.2 [34].

## Results

### *Plasmodium* parasites have a unique transcriptional signature

The correlation of the highest variable genes in *P. berghei*, *P. falciparum* and *P. vivax* revealed two large clusters in the datasets. The expression profiles of the asexual blood stages and liver stages clustered together nicely in the first cluster, whereas gametocytes, ookinetes, and sporozoites made up the majority of the second cluster (Fig. 1A). Regardless of the species or study from which the data were derived, a general grouping of all biological replicates by parasite stages was found using PC analysis of the most variable genes. Within the first two PCs, 82.8% of the total variance explained by the parasite stages was observed. The parasite stages were divided into two sizable groups, much like in the correlation analysis, to show two distinct underlying patterns. The sporozoites and liver stages composed the first cluster along PC1, and the asexual blood stages, gametocytes, and ookinetes composed the second cluster. Nonetheless, the clustering pattern along PC2 was similar to what was observed in the correlation analysis (Fig. 1B).

**Figure 1:**
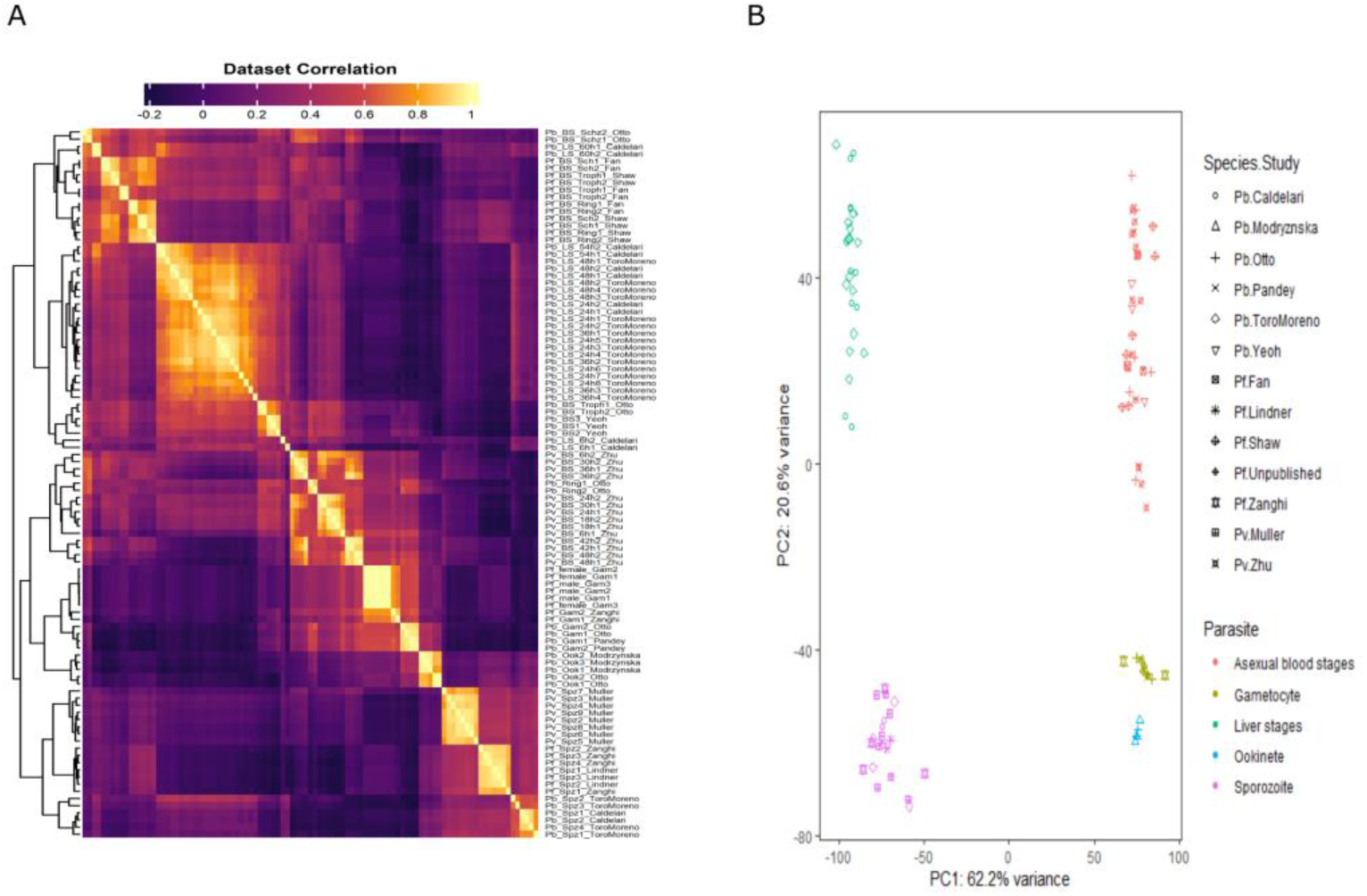
Overview of the *Plasmodium* transcriptome in different species. A. Heatmap of sample-to-sample Spearman correlation of RNA-Seq expression values and B. PCA plot showing the clustering of samples based on their similarity. The rows (A) and data points (B) depict all samples and their biological replicates from different *Plasmodium* spp. generated in different studies.

### RNA-Seq data analysis of *P. berghei*

Differential gene expression was investigated throughout the life cycle stages of *P. berghei* to better understand gene transcription during its development. A total of 3355 DEGs were detected at an adjusted p value (FDR) of less than 0.01 and a minimum log2FC of 2. With the exception of the liver 6 hours postinfection (hpi) stage (L6h) [8] which contained 701 DEGs (Fig. 2A), over 1000 statistically significant DEGs (padj < 0.01, abs(log2FC) > 2) were found in each stage. With 1231 genes identified, the liver stage at 48 hpi (L48h) [8] had the second-lowest number of DEGs discovered. The highest number of DEGs found was 2321 genes during the sporozoite stage [10], while the median number of DEGs found (apart from L6h) was 1706 genes during the ookinete stage [35]. As anticipated, there were variations in the number of DEGs produced for identical samples from two separate studies, underscoring the technical distinctions in RNA extraction, library preparation techniques, RNA sequencing depth, and sequencing platforms (Fig 2A). Filtering the genes found for each stage in the results of their corresponding two independent studies (Fig. 2B) revealed 1832 genes for use in downstream analysis.

**Figure 2:**
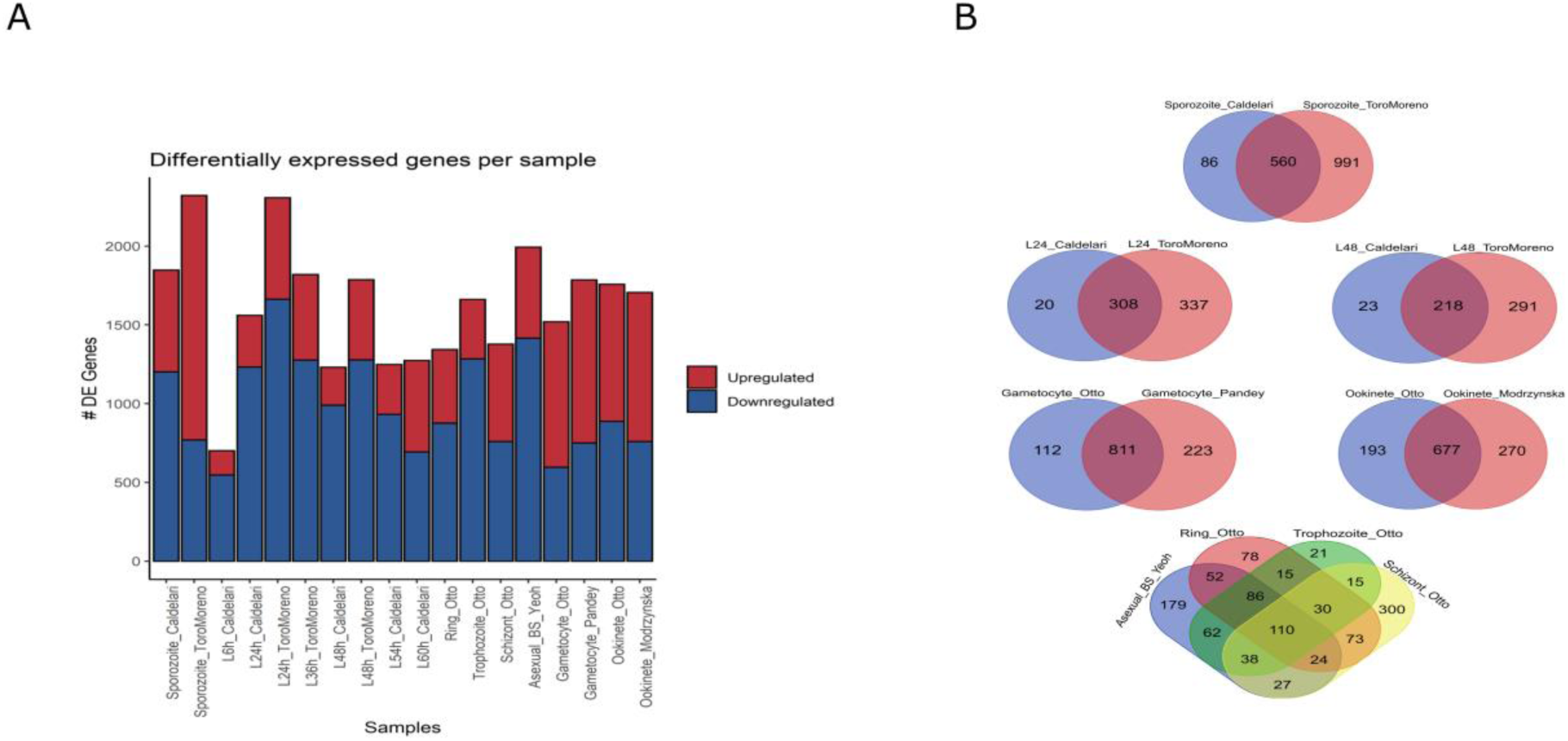
Identification of DEGs in the *P. berghei* samples. A. Total DEGs from the comparison of each stage vs all other stages summarized for each *P. berghei*. sample from each study (padj < 0.01, abs(log2FC) > 2). B. Intersection of upregulated genes identified in each *P. berghei* sample from two independent studies.

Gene ontology terms that describe the related biological processes of the DEGs at each developmental stage were identified. Several GO terms were found to be shared across comparable phases, suggesting similar functional themes. Among the biological processes enriched in the liver stages were metabolic processes (such as fatty acid metabolism), “translation”, “gene expression”, “oxidation‒reduction process”, and “interaction with host” (Fig. 3A). Genes encoding *PK2, pdhA, and ACC*, among other enzymes involved in the type II fatty acid synthesis (FASII) pathway, and genes encoding Fab proteins (*FabB/FabF, FabZ, FabG*, and *FabI*) were notably upregulated (Additional file 3). Additionally, there was an upregulation of genes encoding enzymes involved in lipid precursor synthesis (*G3PDH, G3PAT*), lipoic acid synthesis (*lipA, libB*) and acyl-CoA synthetase (*ACS*), which are involved in fatty acid transport. Furthermore, the liver-specific genes *LISP1* and *LISP2,* which are known for their functions in the maturation of late liver stages and export to host liver cells, are upregulated (Toro-Moreno et al. 2020).

**Figure 3:**
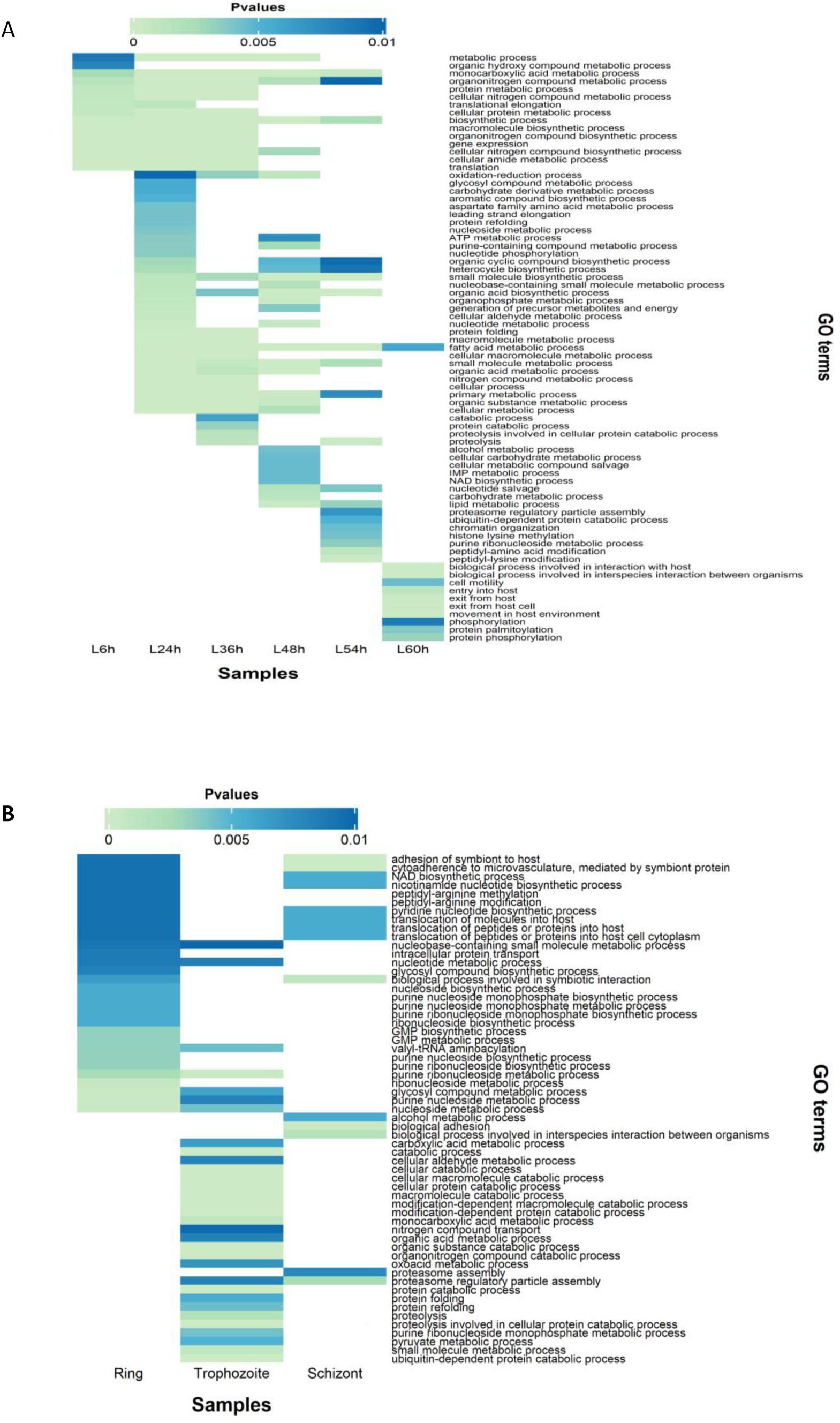
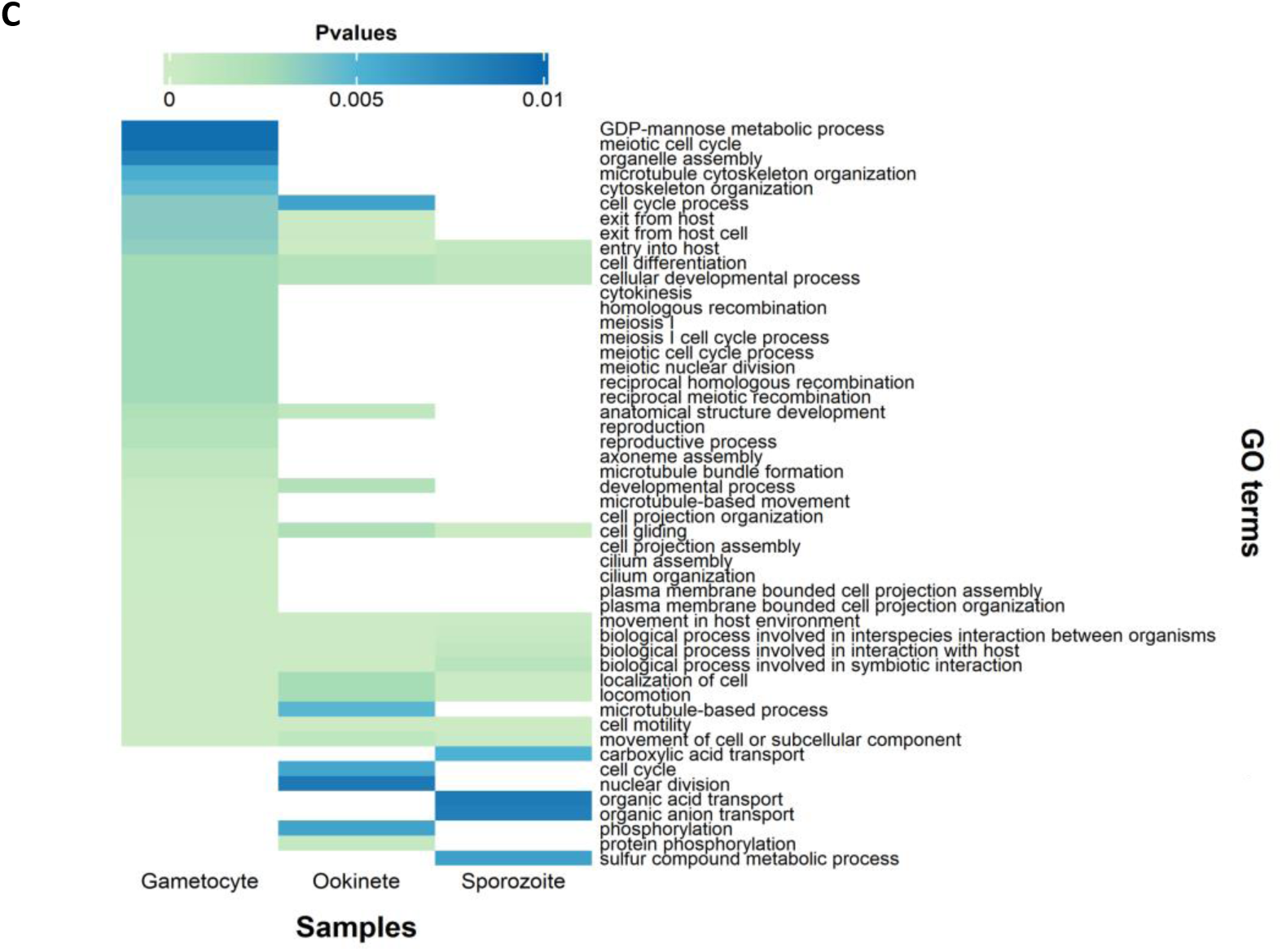
Heatmap of enriched Gene Ontology terms for biological processes associated with *P. berghei*. DEGs in A. liver stages, B. asexual blood stages, and C. sexual and transmission stages (p value<0.01)

The ring and trophozoite blood stages, which were identical to the early and mid-liver stages, respectively, were similarly associated with GO terms related to asexual growth. In addition to metabolic processes, some notable terms enriched in the blood stages were “translocation of peptides or proteins into the host cell cytoplasm” and “cytoadherence to microvasculature” (Fig. 3B). The genes that were found to be upregulated included those that encode proteins involved in invasion, such as SERA proteins (*SERA3, SERA4*), Rhoptry genes (*RhopH2, RhopH3*), and merozoite surface proteins (MSPs), such as *MSP2* and *MSP7*. Throughout the stages, there was a constant upregulation of genes from the lysine-rich small, exported protein/early transcribed membrane protein families (*SEP/ETRAMP*) and multigene families of exported proteins, including PIR, fam-a, and fam-b proteins (Additional file 3). Furthermore, the biological processes related to symbiotic or interspecies interactions with the host were enriched in the schizonts (Fig. 3B).

Genes involved in sexual reproduction and the egress of parasites from host cells were upregulated during the gametocyte stage. These genes were implicated in various activities, including “meiotic cell cycle”, “homologous recombination”, “cytokinesis”, “axoneme assembly”, “cytoskeleton organization”, “locomotion”, and “exit from host cell” (Fig. 3C). The upregulated genes included those encoding nuclear fusion proteins (*NEK2* and *NEK4*); surface protein-coding genes such as *P230, P48/45, CCp, P25*, and *P28*; the female-specific gene *G377*; the shared gene *MDV1*; and male-specific genes such as guanyl cyclase (*GC*), phospholipase C, *CDPK4*, *MAPK2*, and *HAP2* (Additional file 3).

The upregulated genes in the mosquito stages were enriched in biological processes, such as “cell differentiation”, “cellular developmental process”, “entry into the host”, “cell gliding”, “movement in the host environment”, and biological processes involving symbiotic or interspecies interactions (Fig. 3C). In particular, there was an upregulation of genes encoding invasion proteins, including TRAP proteins, TRAP-related proteins, *CSP*, and genes encoding cell traversal proteins, including *PL*, *SPECT1*, and *CelTOS* (Additional file 3).

### Coexpression clustering analysis reveals functionally enriched clusters

Notably, clustering analysis also split the parasite stages into two clusters, one containing gametocytes, ookinetes, and sporozoites and the other containing liver stages and asexual blood stages, similarly to the correlation analysis. A variety of clustering techniques and algorithms were used to find the ideal number of clusters within the DEGs. Three clusters received the greatest “vote” according to the majority rule (Additional file 4). For this reason, clustering was performed with k = 3. The GO enrichment tool from PlasmoDB (version 51) was used to extract the biological processes connected with each cluster to better understand them, and then these terms were summarized using REVIGO.

Cluster 1 included 629 genes (Additional file 5) that were typically downregulated in the liver and asexual blood stages and mostly upregulated in the sporozoite stage and partially upregulated in the gametocyte and ookinete stages (Fig. 4). Biological processes associated with migration in the host environment and interspecies interactions, which are compatible with being ready to infect a new host, were strongly enriched in this cluster (Additional file 6). A total of 536 genes (Additional file 5) from Cluster 2 were downregulated in the sporozoite and ookinete stages, upregulated in the liver and asexual blood stages and partially upregulated in the gametocytes (Fig. 4). The metabolism of various biochemical components, gene expression, proteolysis, and biological adhesion to the host were among the biological activities that were substantially enriched (Additional file 6). A total of 667 genes (Additional file 5) whose expression was downregulated in the sporozoite, liver, and asexual blood stages and upregulated in the gametocyte and ookinete stages were found in Cluster 3. (Fig. 4). Many biological processes related to cell motility, microtubule-based activities, and host interactions were enriched similarly to Cluster 1, as the parasite transitions from a nonmotile to a motile state and is ready to move from the vertebrate host to a mosquito vector (Additional file 6).

**Figure 4:**
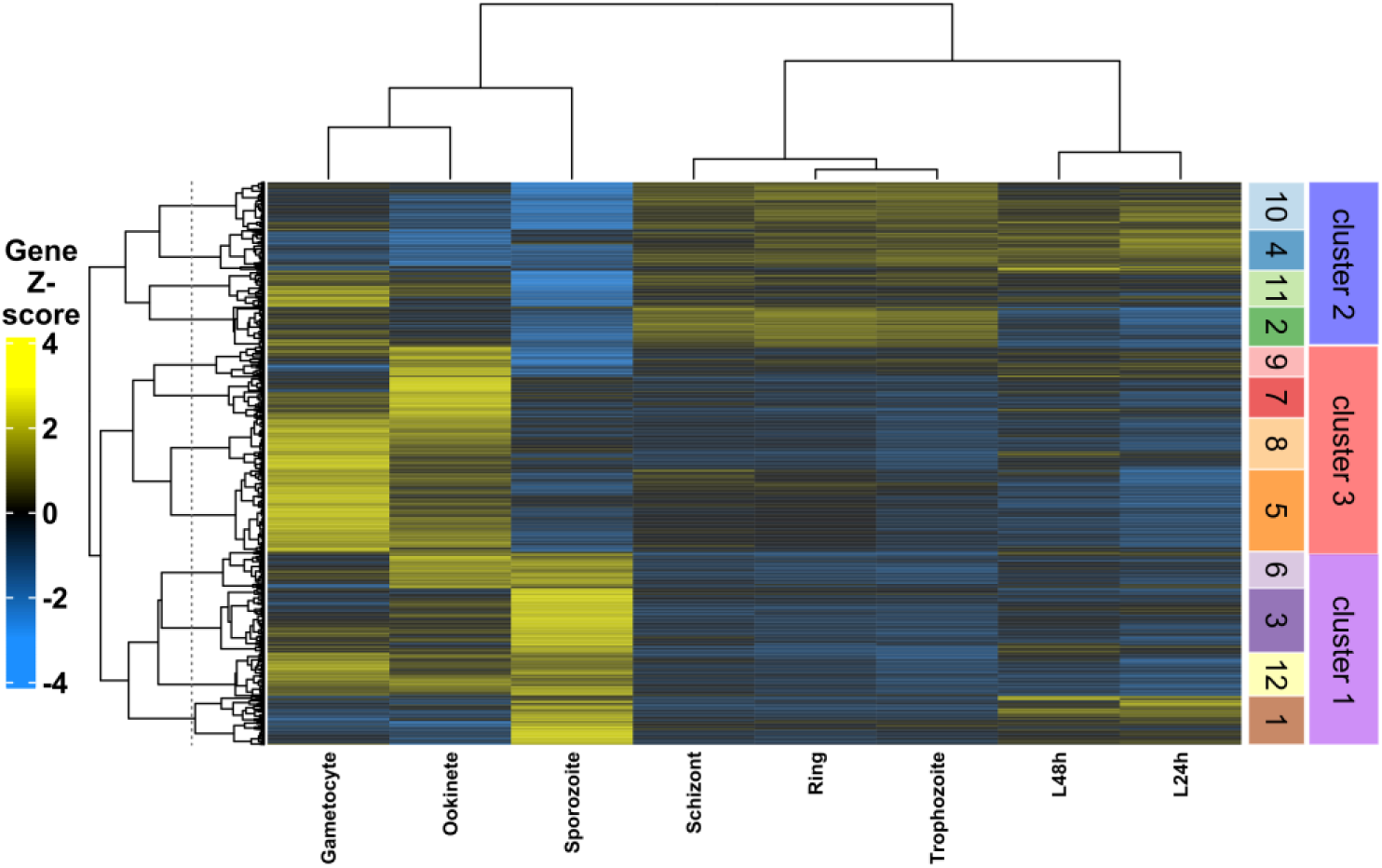
Heatmap of RNA-Seq-based average expression values across replicates, scaled by z-transformation. A total of 1832 genes (rows) are shown across the life cycle stages (columns). Clusters from PAM with k=3 and k=12 are shown as colour bars on the right-hand side. The dendrograms on the left-hand side show the within-cluster and cluster-to-cluster relationships of the PAM clusters, while the dendrograms on the top of the heatmap show the relationships among the stages.

Furthermore, mixed clusters were still discernible even after 3 was determined to be the ideal number for k by the majority rule. Thus, we choose k = 12 to produce smaller (sub)clusters and enhance the clusters’ homogeneity. Nevertheless, selecting 12 was arbitrary. The PAM clusters could be reorganised into larger k = 3 clusters by applying hierarchical clustering to the individual subclusters (Fig. 4). Subclusters 1, 3, 6, and 12 comprised Cluster 1; subclusters 2, 4, 10 and 11 comprised Cluster 2; and subclusters 5, 7, 8, and 9 comprised Cluster 3 (Fig. 4).

A total of 159 genes were found in subcluster 1, including the sporozoite-specific UIS protein-coding genes (*UIS2, UIS3, UIS4, UIS8, UIS12*), *TREP*; the sporozoite and liver-specific SLARP proteins; genes encoding proteins involved in fatty acid and lipid metabolism (Fab proteins (*FabB/FabF, FabG*), *pdhA, ACS*); chaperone and trafficking protein genes (*DnaJ, ClpM, MFR1, NPT1, FNT*); mediators of RNA Pol II transcription; liver-specific *LISP1,* and the ApiAP2s *AP2-L* and *PBANKA 0932300* (Additional file 7). This cluster showed downregulation in gametocytes and ookinetes, strong upregulation in sporozoites and very little upregulation until the blood-stage trophozoites (Fig. 4). The metabolic processes of fatty acids and lipids, “organic acid biosynthetic process”, “pyruvate transport” and “DNA-templated transcription” were the most enriched biological processes in this cluster (Additional file 8).

The 130 genes in subcluster 2 included the following: genes encoding parasitophorous vacuole protein 5 (*PV5*), merozoite surface proteins (*MSP1, MSP8*), export and membrane-associated protein-coding genes (*PIR, fam-a, fam-b, ETRAMP, EXP, SEP, MAHRP* families, *SMAC*, and *IBIS1*), rhoptry protein-coding genes (*RON11, RhopH1A, RhopH2, RhopH3*), the SERA protein-coding gene *SERA4*, and the ApiAP2 *AP2-G* (Additional file 7) expressed in the ring, trophozoite, schizont, and gametocyte stages, with strong downregulation in the sporozoite (Fig. 4). “Biological process involved in interspecies interaction between organisms”, “adhesion of symbiont to microvasculature”, “adhesion of symbiont to host”, “translocation of peptides or proteins into host”, “translocation of molecules into host”, and “protein unfolding” were among those enriched in this cluster (Additional file 8).

Subcluster 3 comprised 211 genes, including the gene encoding the most abundant surface protein on sporozoites (*CSP*); other genes encoding surface proteins (*TRAP, TLP, TRSP, SPELD, P113, SSP3*); the gene encoding the cell traversal protein (*SPECT1*); the gene encoding microneme protein (*CLAMP*); the genes encoding apical membrane proteins (*AMA1, TRAMP*); the gene encoding *SERA5*; iron-sulfur cluster assembly protein-coding genes (*SufE, NFU1, ISCU, NFS1*); and ApiAP2s (*AP2-SP, AP2-HS, PBANKA 0521700*) (Additional file 7) expressed in the gametocyte and ookinete, peaking in the sporozoite and downregulated in the liver and asexual blood stages (Fig. 4). “Biological processes involving sulfur compound metabolic process”, “entry into host”, “movement in host”, “iron-sulfur cluster assembly” and “metallo-sulfur cluster assembly” were the main biological processes enriched in this cluster (Additional file 8).

A total of 133 genes were found in subcluster 4, which were mainly expressed in the L24 stage and through the asexual blood stages, with downregulation in gametocytes, ookinetes, and sporozoites (Fig. 4). These genes included translation initiation factor genes (*IF2a, EIF5A*), liver-specific *LISP2*, chromatin remodelling genes (*CARM1, PRMT5*), and genes encoding proteins involved in carbohydrate metabolism (*6PGD, TPT, INO1*) and fatty acid synthesis (*ACP, LipA*) (Additional file 7). “Translation”, “peptidyl-amino acid modification”, “carbohydrate metabolic process”, “peptidyl-arginine modification”, and “peptidyl-arginine methylation” were the biological processes with the highest levels of enrichment (Additional file 8).

The 267 genes in subcluster 5 included *PIR* genes; genes encoding gametocyte and ookinete surface proteins (*P28, P48/45, P47, MiGS*); genes specific to males (*CDPK4, HAP2, MD2*); genes specific to females (*G377, FD2, FD4*); the shared gene *MDV1*; the gene encoding nuclear fusion protein *NEK4*; genes encoding secreted ookinetes (*PSOP13, PSOP20*); inner membrane complex protein-coding genes (*IMC1b, IMC1c, IMC1e, IMC1h, IMC1k, IMC32, ALV7, ISC3*); glideosome-associated protein-coding genes (*GAP40, GAP50, GAPM1 - 3*); and the ApiAP2 *AP2-Z* (Additional file 7). The genes in this cluster were strongly expressed in the gametocyte, with reduced expression in the ookinete and downregulated expression in the sporozoite and throughout the liver and asexual stages (Fig. 4). “Microtubule-based processes”, “microtubule-based movement”, “cilium assembly”, “biological_process”, and “organelle assembly” were the biological processes that were most enriched in this cluster (Additional file 8).

Subcluster 6 comprised 119 genes, including those encoding proteins for the inner membrane complex (*IMC1g, IMC1l, and IMC1m*); the ookinete maturation gene 1 (*OMG1*); genes encoding proteins for invasion (*CelTOS, SIAP1*); RNA-associated protein-coding genes (*HAS1, UTP4*); phospholipase *PLA1*; and the ApiAP2 *AP2-O4* (Additional file 7). These genes were expressed in the ookinete, with greater expression in the sporozoite stage and were downregulated throughout the liver stages until the gametocyte stage(Fig. 4). “Protein phosphorylation”, “RNA secondary structure unwinding”, “carbon fixation”, “regulation of the lipoprotein metabolic process”, and “regulation of protein lipidation” were among the most enriched biological activities (Additional file 8).

A total of 134 genes were found in subcluster 7, including genes encoding secreted ookinete proteins (*PSOP1, PSOP2, PSOP7, SOAP, CTRP, TRP1, PIMMS1, PIMMS2, PIMMS43, PIMMS57*); genes associated with motility (*MyoB, MyoE, MLC-B, ARP4*); and ApiAP2 *AP2-LT* (Additional file 7), which were expressed in gametocytes and strongly expressed in the ookinete and downregulated for the remainder of the life cycle (Fig. 4). This cluster was primarily enriched for “entry into host”, “movement in host”, “biological process involved in interaction with host”, “biological process involved in interspecies interaction between organisms” and “protein phosphorylation” (Additional file 8).

The 168 genes in subcluster 8 included genes encoding enzymes involved in the metabolism of mannose and nucleotide sugars (*GMD, HAD5, UDG*); motility-associated protein-coding genes (*ARP1*, kinesin-20, dynein light chains); nuclear fusion protein-coding genes (*GEX1, NEK2*); genes specific to sexual development (*G37, GEP1*); the ookinete surface protein-coding gene *P25*; ookinete secreted protein-coding genes (*PSOP6, PSOP12, PSOP17*); and the ApiAP2 *AP2-O5* (Additional file 7). These genes exhibit strong expression in gametocytes and reduced expression in ookinetes, with downregulation occurring throughout the remainder of the life cycle (Fig. 4). The metabolic processes of GDP-mannose metabolic process and nucleotide-sugar, “cell motility”, “karyogamy” and “viral process” were the most enriched biological processes (Additional file 8).

Subcluster 9 comprised 98 genes, including methyltransferases (*RCM, NEP1*), autophagy-related protein-coding genes (*ATG5, ATG7, VPS15*), and enzymes involved in tRNA modification and RNA editing (*GATA, KAE1, GluRS, NCS6*, cytidine deaminase, and dihydrouridine synthase) (Additional file 7) which showed very little expression from the liver stages to the asexual blood stages and gametocyte stage, the strongest expression in the ookinete stage and strong downregulation in the sporozoite stage (Fig. 4). The biological processes associated with tRNA metabolism, “RNA modification”, “tRNA modification”, “methylation”, and “autophagy” were the most enriched in this cluster (Additional file 8).

There were 155 genes in subcluster 10, including those related to protein folding (*HSP60, CCT2*, peptidyl-prolyl cis-trans isomerase, *DnaJ*); genes encoding translation initiation factors (*EIF-3H, eIF2gamma*); proteasome-related protein-coding genes (*RPT4, RPN5, RPN6*); and the ApiAP2 *AP2-G2* (Additional file 7), which were expressed in the L24 stage and through the asexual blood and gametocyte, with downregulation in the ookinete and strong downregulation in the sporozoite (Fig. 4). This cluster was primarily enriched for biological processes related to “translation”, “ubiquitin-dependent protein catabolism”, “nucleotide metabolic process”, “nucleobase-containing small molecule metabolic process”, and “protein folding” (Additional file 8).

Subcluster 11 comprised 118 genes, including sex-determining and reproduction-associated protein-coding genes (*FD1, MD3, MD5*) and the ApiAP2 *PBANKA 1403700* (Additional file 7), which was expressed from the L24 stage to the ookinete stage and was significantly downregulated in the sporozoite (Fig. 4). Nonetheless, the highest level of expression was detected in the gametocyte, revealing the most enriched biological process for “sex determination”, “alcohol metabolic process”, “reproduction”, “reproductive process”, and “organic hydroxy compound metabolic process” (Additional file 8).

A total of 140 genes were found in subcluster 12, including those necessary for mobility and structure (*PAT, IMC1a, LIMP, SIMP, and GEST*) and the ApiAP2 *AP2-O* (Additional file 7), which were downregulated for the remainder of the life cycle after being expressed from the gametocyte to the sporozoite (Fig. 4). “Cell gliding”, “movement in host”, “biological_process”, “biological process involved in interaction with host” and “regulation of cell shape” were among the most enriched biological processes (Additional file 8).

### *De novo* motif discovery

To identify the resident putative regulatory sequences upstream of the genes showing the observed co-expression patterns, we searched their promoter regions for over-representation of short sequences using two *de novo* motif discovery tools from RSAT and MEME. Each tool identified a collection of motifs in the coexpression clusters, which were then clustered to remove the redundancy from both tools and identify the representative root motif for each motif cluster. The clustering of the motif collections for each coexpression cluster and subcluster are shown in Additional files S9 and S10. Similarities between the discovered root motifs and those in JASPAR, a sizable collection of recognised, nonredundant motifs from plants, animals, nematodes, fungi, and insects, were detected. JASPAR contains binding motifs for a number of transcription factor (TF) types, including nuclear receptors, zinc fingers, bZips, helix turn helices, forkheads, POU/Homeodomains, and Myb proteins. Notably, the majority of the root motifs in all of the clusters and subclusters matched the known DNA binding patterns of the ApiAP2 TFs, which are important regulators of *Plasmodium* gene transcription and were initially found in *P. falciparum* via PBM experiments [5]. Additionally, the majority of the motifs (Fig. 5) exhibited similarities to *de novo* motifs predicted by an ATAC-Seq analysis [30].

**Figure 5:**
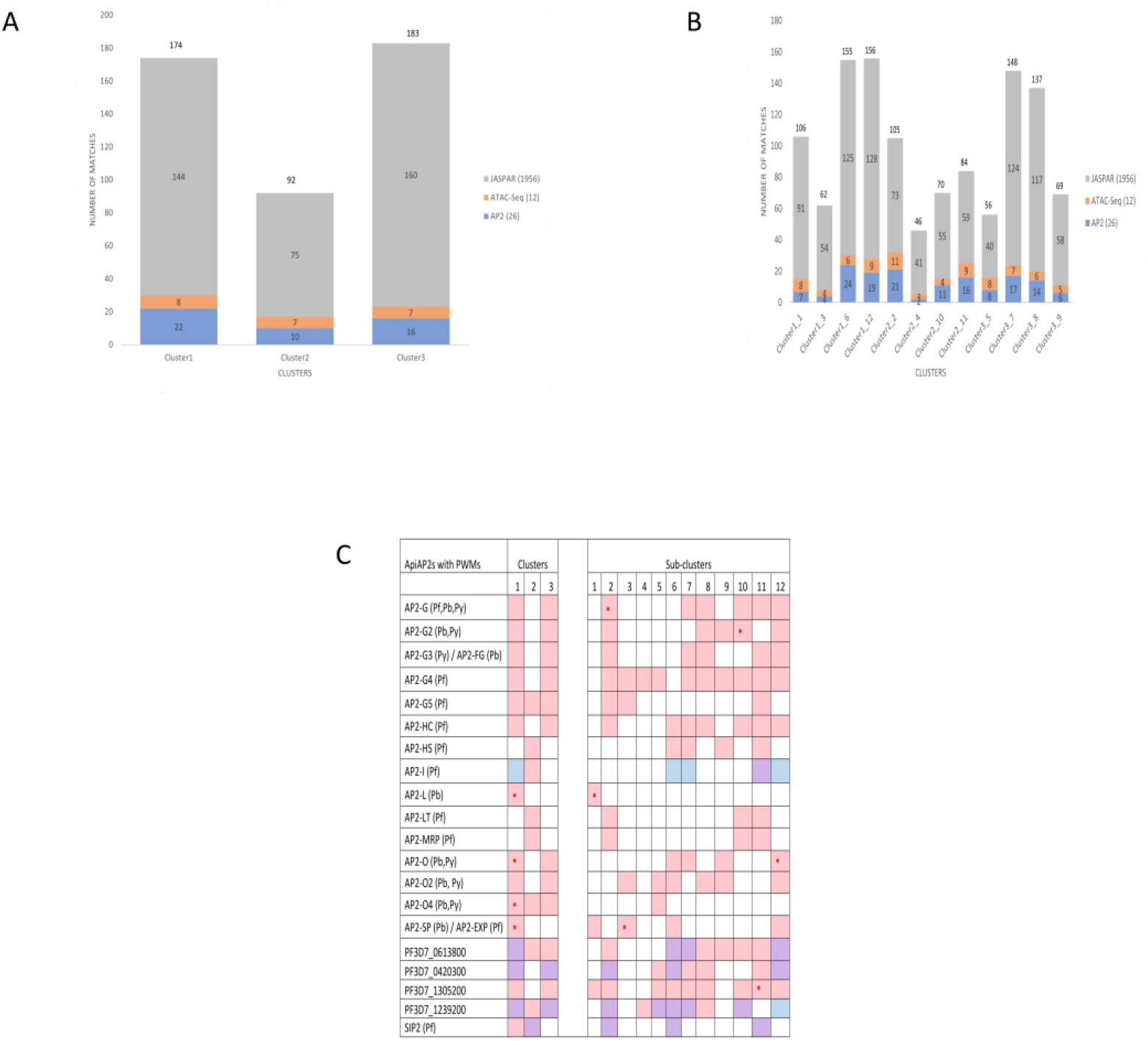
Similarities of the discovered root motifs to known motifs. A-B. Comparison of the discovered motifs from each PAM cluster (A, k= 3; B, k=12) to known motifs. Similarities were computed based on Euclidean distances. The total number of motifs in each database of known motifs is shown in brackets. Each stack of the bars corresponds to the number of known motifs in the respective collection that had matches to the discovered motifs in the cluster. Each subcluster is named according to its main cluster, followed by an underscore and the actual subcluster. The total number of matching motifs found for each cluster is represented by the number on top of the stacked bar. C. Comparison of the discovered motifs from each PAM cluster to known motifs of AP2 domains with position weight matrices. Clusters = PAM clusters with k=3; subclusters = PAM clusters with k=12; Colours = similarity of the discovered motif to known AP2 binding sites detected: pink = motif similar to the binding site of the first AP2 domain, blue = motif similar to the binding site of the second AP2 domain, purple = motif similar to the binding site of both AP2 domains; * motif similarity detected in the same stage AP2 TF is expressed

The majority of the root motifs in each main cluster and subcluster showed redundancy in their resemblance to the motifs predicted by ATAC-Seq, the binding sites of the AP2 TFs, and the known motifs in JASPAR. Similar motifs for the well-known transcriptional activator of sporozoites, AP2-SP, as well as other gametocyte- and ookinete-relevant AP2s, such as AP2-G, AP2-G2, AP2-FG, AP2-G4, AP2-G5, AP2-O, AP2-O2, and AP2-O4, were identified in Cluster 1, which contains genes highly expressed in sporozoites and with lower expression in gametocytes and ookinetes (Additional file S11). Similarly, motifs pertaining to the pertinent AP2s were also noted in the related subclusters: AP2-SP and AP2-L (subcluster 1); AP2-SP, AP2-O2, *Pf*AP2-G4, and *Pf*AP2-G5 (subcluster 3); AP2-SP, AP2-O, and AP2-O2 (subcluster 6); and AP2-G, AP2-G2, AP2-FG, *Pf*AP2-G4, AP2-O, AP2-O2, and AP2-SP (subcluster 12) (Additional file S12).

A motif resembling the binding site of *Pf*AP2-I, the TF that orchestrates red blood cell invasion, was found among the identified motifs in Cluster 2, which is primarily composed of genes related to the asexual stage with a small number of genes related to gametocytes and ookinetes (Additional file S11). Other motifs included AP2-O4 and *Pf*AP2-G5. Nonetheless, motifs for other AP2s important for the asexual stage, such as *Pf*AP2-LT and *Pf*AP2-MRP, were present in the related subclusters. *Pf*AP2-I, *Pf*AP2-MRP, *Pf*AP2-LT, AP2-G, AP2-G2, AP2-FG, *Pf*AP2-G4, *Pf*AP2-G5 (subcluster 2), *Pf*AP2-G4 (subcluster 4), *Pf*AP2-MRP, *Pf*AP2-LT, AP2-G, AP2-G2, *Pf*AP2-G4 (subcluster 10) and *Pf*AP2-I, *Pf*AP2-MRP, *Pf*AP2-LT, AP2-G, AP2-FG, *Pf*AP2-G4, and *Pf*AP2-G5 (subcluster 11) are specifically the observed AP2s (Additional file S12).

Similar patterns for important AP2 TFs, such as AP2-G, AP2-G2, AP2-FG, AP2-G4, *Pf*AP2-G5, AP2-O, AP2-O2, and AP2-O4, were found in Cluster 3, which primarily contains genes related to gametocytes and ookinetes (Additional file S11). The AP2 TFs *Pf*AP2-G4, AP2-O2, AP2-O4 (subcluster 5), AP2-G, AP2-FG, *Pf*AP2-G4, AP2-O (subcluster 7), AP2-G, AP2-G2, AP2-FG, *Pf*AP2-G4, AP2-O2 (subcluster 8), and AP2-O, AP2-O2, AP2-G2, *Pf*AP2-G4 (subcluster 9) are among the specific motifs that were found (Additional file S12). *Pf*SIP2 (Clusters 1 and 2, subclusters 2, 6 and 11), *Pf*AP2-HS (Cluster 2, subclusters 6, 7, 9 and 11), *Pf*AP2-HC (Clusters 1 and 3, subclusters 2, 6-8 and 10-12) (Additional files S11 and S12), and uncharacterized AP2s (Fig. 5C) are additional motifs for AP2 factors that demonstrated similarities to the identified motifs.

Eight of the *de novo* discovered motifs did not share commonalities with the extensive database of known motifs in JASPAR, the predicted motifs from the ATAC-Seq analysis, or the known ApiAP2 motifs, despite the majority of the identified motifs demonstrating similarities. GO terms for biological processes, cellular components, and molecular functions as well as KEGG pathway enrichments from PlasmoDB (version 65) were used to annotate these motifs (Fig. 6), and revealed the following:

**Figure 6:**
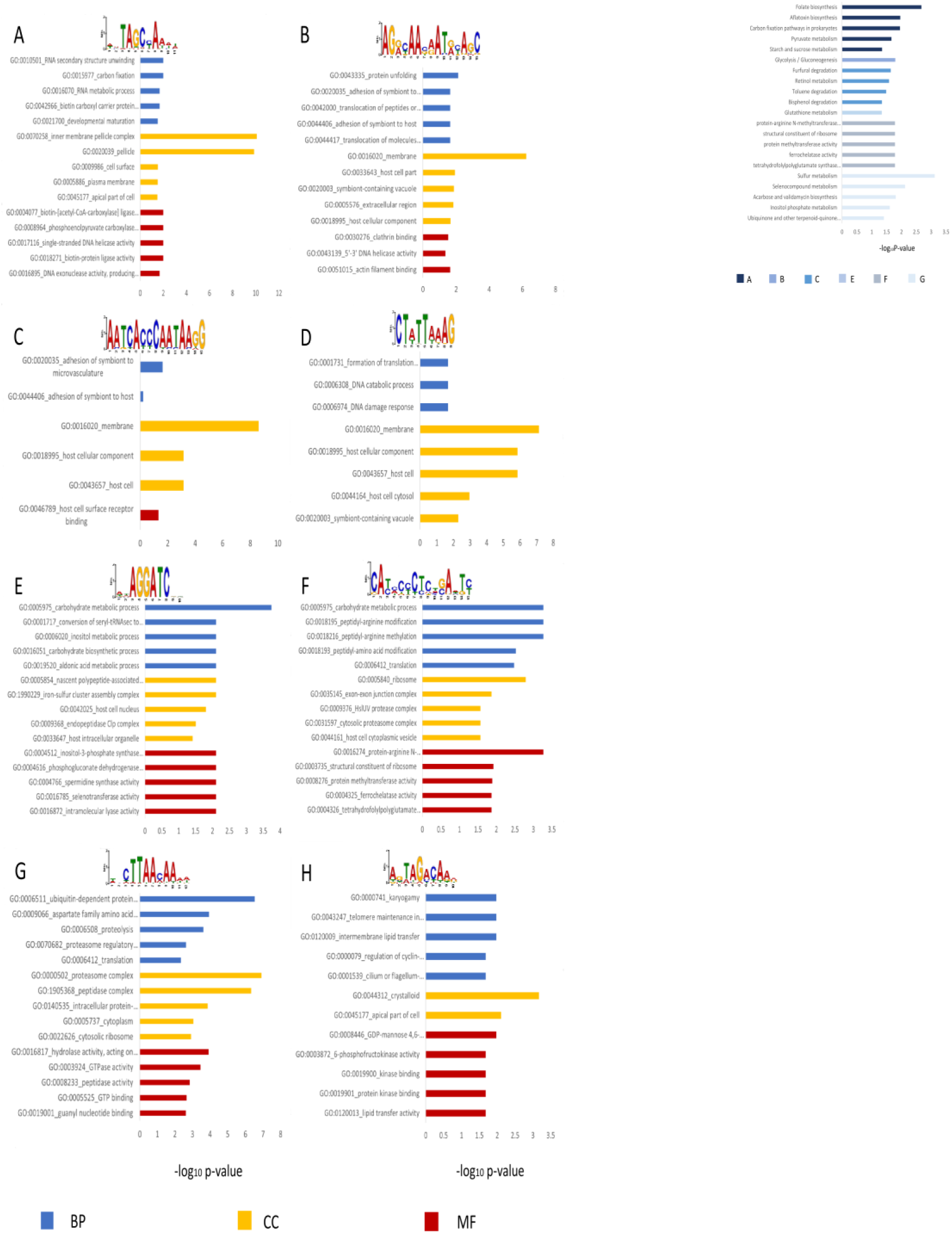
Enrichment of GO terms (left) and KEGG metabolic pathways (right) related to the unknown motifs. The labels A-H represent each motif that did not resemble known motifs. BP: biological process, CC: cellular component, MF: molecular function.

The fifth motif in Cluster 1_6 (Fig. 6A) was primarily enriched in the promoter regions of genes related to cellular components such as “inner membrane pellicle complex’, “pellicle”, “cell surface”, “plasma membrane”, and “apical part of cell”, as well as biological processes such as “RNA secondary structure unwinding”, “carbon fixation”, “RNA metabolic process”, “biotin carboxyl carrier protein biosynthetic process”, and “developmental maturation”. The primary enriched molecular functions were “DNA exonuclease activity producing 5’-phosphomonoesters”, “phosphoenolpyruvate carboxylase activity”, “single-stranded DNA helicase activity”, “biotin-protein ligase activity”, and “biotin-[acetyl-CoA-carboxylase] ligase activity” (Fig. 6A and Additional file 13). Pathway analysis revealed an enrichment of “folate biosynthesis”, “aflatoxin biosynthesis”, “carbon fixation pathways in prokaryotes”, “pyruvate metabolism”, and “starch and sucrose metabolism” (Fig. 6 and Additional file 14).

The biological processes of “protein unfolding”, “adhesion of symbiont to microvasculature”, “translocation of peptides or proteins into host”, and “translocation of molecules into host” were among those enriched in the genes harbouring the 11th motif in Cluster 2_2 (Fig. 6B). “Membrane”, “host cell part”, “symbiont-containing vacuole”, “extracellular region”, and “host cellular component” were the most enriched cellular components; “clathrin binding”, “5’-3’ DNA helicase activity”, and “actin filament binding” were the most enriched molecular functions (Fig. 6B and Additional file 13). Pathway analysis revealed participation in “glycolysis/gluconeogenesis” (Fig. 6 and Additional file 14).

The 13th motif in Cluster 2_2 (Fig. 6C) was enriched in the promoters of genes involved in biological processes related to “adhesion of symbiont to microvasculature” and “adhesion of symbiont to host”; and cellular components of “membrane”, “host cellular component”, and “host cell”. Its molecular function is “host cell surface receptor binding” (Fig. 6C and Additional file 13). “Furfural degradation”, “retinol metabolism”, “toluene degradation”, and “bisphenol degradation” were the enriched pathways (Fig. 6 and Additional file 14).

The genes exhibiting the 14th motif in Cluster 2_2 (Fig. 6D) were associated with enriched biological processes, including “formation of translation preinitiation complex”,” DNA catabolic process”, and “DNA damage response”. “Membrane”,” host cellular component”, “host cell”, “host cell cytosol”, and “symbiont-containing vacuole” were among the enriched cellular components (Fig. 6D and Additional file 13). There was no enrichment for molecular pathways or activities in this motif.

The metabolic processes of carbohydrates, aldonic acid, inositol, “conversion of seryl-tRNAsec to selenocystRNAsec”, and “carbohydrate biosynthetic process” were among the biological activities enriched in the genes harbouring the 9th motif in Cluster 2_4 (Fig. 6E).“Host cell nucleus”, “endopeptidase Clp complex”, “iron-sulfur cluster assembly complex”, “nascent polypeptide-associated complex”, and “host intracellular organelle” were the enriched biological components. The activities of phosphogluconate dehydrogenase (decarboxylating), spermidine synthase, selenotransferase, intramolecular lyase, and inositol-3-phosphate synthase were among the enriched molecular functions (Fig. 6E and Additional file 13). Participation in “glutathione metabolism” was shown by pathway analysis (Fig. 6 and Additional file 14).

The promoters of genes harbouring the 11th motif in Cluster 2_4 (Fig. 6F) were enriched for “carbohydrate metabolic process”, “peptidyl-arginine modification”, “peptidyl-arginine methylation”, “peptidyl-amino acid modification”, and “translation”, as well as cellular components such as “ribosome”, “exon‒exon junction complex”, “HslUV protease complex”, “cytosolic proteasome complex”, and “host cell cytoplasmic vesicle”. “Protein-arginine N-methyltransferase activity”, “ferrochelatase activity”, “protein methyltransferase activity”, and “tetrahydrofolylpolyglutamate synthase activity” were the primary enriched molecular functions (Fig. 6F and Additional file 13). The top pathways identified by pathway analysis were “tetrahydrofolylpolyglutamate synthase activity”, “ferrochelatase activity”, “protein methyltransferase activity”, “structural component of the ribosome”, and “protein-arginine N-methyltransferase activity” (Fig. 6 and Additional file 14).

The promoters of genes harbouring the fourth motif in Cluster 2_10 (Fig. 6G) were enriched in biological processes related to “ubiquitin-dependent protein catabolism”, “aspartate family amino acid metabolism”, “proteolysis”,” proteasome regulatory particle assembly”, and “translation”. The top enriched molecular functions included “hydrolase activity, acting on acid anhydrides”, “GTPase activity”, “peptidase activity”, “GTP binding”, and “guanyl nucleotide binding”; the top enriched cellular components included “proteasome complex”, “peptidase complex”, “intracellular protein-containing complex”, “cytoplasm”, and “cytosolic ribosome” (Fig. 6G and Additional file 13). Pathway analysis revealed that the metabolism of sulphur, selenocompounds, inositol phosphate, “acarbose and validamycin biosynthesis”, and “ubiquinone and other terpenoid-quinone biosynthesis” were the most important pathways (Fig. 6 and Additional file 14).

Finally, “karyogamy”, “telomere maintenance in response to DNA damage”, “intermembrane lipid transfer”, “regulation of cyclin-dependent protein serine/threonine kinase activity”, and “cilium or flagellum-dependent cell motility” were the top enriched biological processes for the genes harbouring the 5th motif in Cluster 3_8 (Fig. 6H), whereas the enriched cellular components were “crystalloid” and “apical part of cell”. The enriched molecular functions included “GDP-mannose 4,6-dehydratase activity”, “6-phosphofructokinase activity”, “kinase binding”, “protein kinase binding”, and “lipid transfer activity” (Fig. 6H and Additional file 13). There was no enrichment for KEGG pathways for this motif.

### Further analysis of selected motifs

The fifth and seventh root motifs within Cluster 2, which included motifs with putative regulatory roles during asexual development, did not resemble any of the known ApiAP2 motifs or the *de novo* motifs predicted by ATAC-Seq in a different study to be enriched in accessible chromatin regions of the *P. falciparum* genome during the IDC (Additional files S9 and S11). This finding was made through motif comparison analysis using the TOMTOM tool. Nevertheless, additional investigations revealed that the fifth motif, [A/T][A/T]AGGTAA[A/T][A/T], or simply AGGTAA, might be related to *de novo* motif 031, one of the ATAC-Seq-predicted motifs. TTATTACAC is the binding sequence for *de novo* motif 031. According to DNA pull-down experiments [30], T**<u>T</u>**A**<u>TTAC</u>**AC interacts with PF3D7 0420300/PBANKA 0521700, the second AP2 domain of the ApiAP2 TF, which has an unclear function and a binding sequence of G**<u>T</u>**G**<u>TTAC</u>**A. This finding implies that this ApiAP2 TF may also bind to the reverse complement of the AGGTAA motif (**<u>T</u>**T**<u>TTAC</u>**CTTT), suggesting a direct interaction.

The ATAC peaks localised to a variety of locations in the *P. falciparum* genome, including promoters, exons, and intergenic regions; hence, only the ATAC-Seq peak regions that mapped exactly to 5’ UTRs were desired. The regions where the 5’ UTRs and the ATAC peaks overlapped were subsequently extracted. Using the RSAT “matrix-scan” tool, the individual genomic coordinates for each occurrence of the AGGTAA motif and the *de novo* motif 031 within the accessible chromatin 5’UTRs were extracted, revealing that both motifs shared some genomic locations (Fig. 7A). The predicted motifs in the same cluster as the *de novo* motif 031 (TTATTACAC) [30] included the motif for *Pf*AP2-I (PF3D7 1007700/PBANKA 1205900), a well-characterized ApiAP2 TF whose third AP2 domain contains the binding sequence GTGCACTA. During asexual development, *Pf*AP2-I orchestrates the invasion of red blood cells [36]. Similarly, in our analysis, the motif for *Pf*AP2-I was validated by the motif comparison tool TOMTOM to be comparable to the [A/T][A/T]GCACTA[A/T][A/T] motif, or GCACTA for short. This motif appears in the same cluster as the [A/T][A/T]AGGTAA[A/T][A/T] motif, which has a reverse complement TTTTACCTTT. This finding implies that the TFs that bind both motifs probably coregulate the target genes that harbour them. Overall, the connections between *Pf*AP2-I, AGGTAA, and *de novo* motif 031 were investigated through the construction of a gene regulatory network. Using the “matrix-scan” tool, 287 target genes from Cluster 2 that harboured the three motifs were extracted. Fifty-eight genes (≈20%) had binding sites for two of the motifs, whereas 218 genes (≈76%) had binding sites for only one of the motifs. Among all the genes, only 11 (approximately 4%) had binding sites for all three motifs (Fig. 7B).

**Figure 7:**
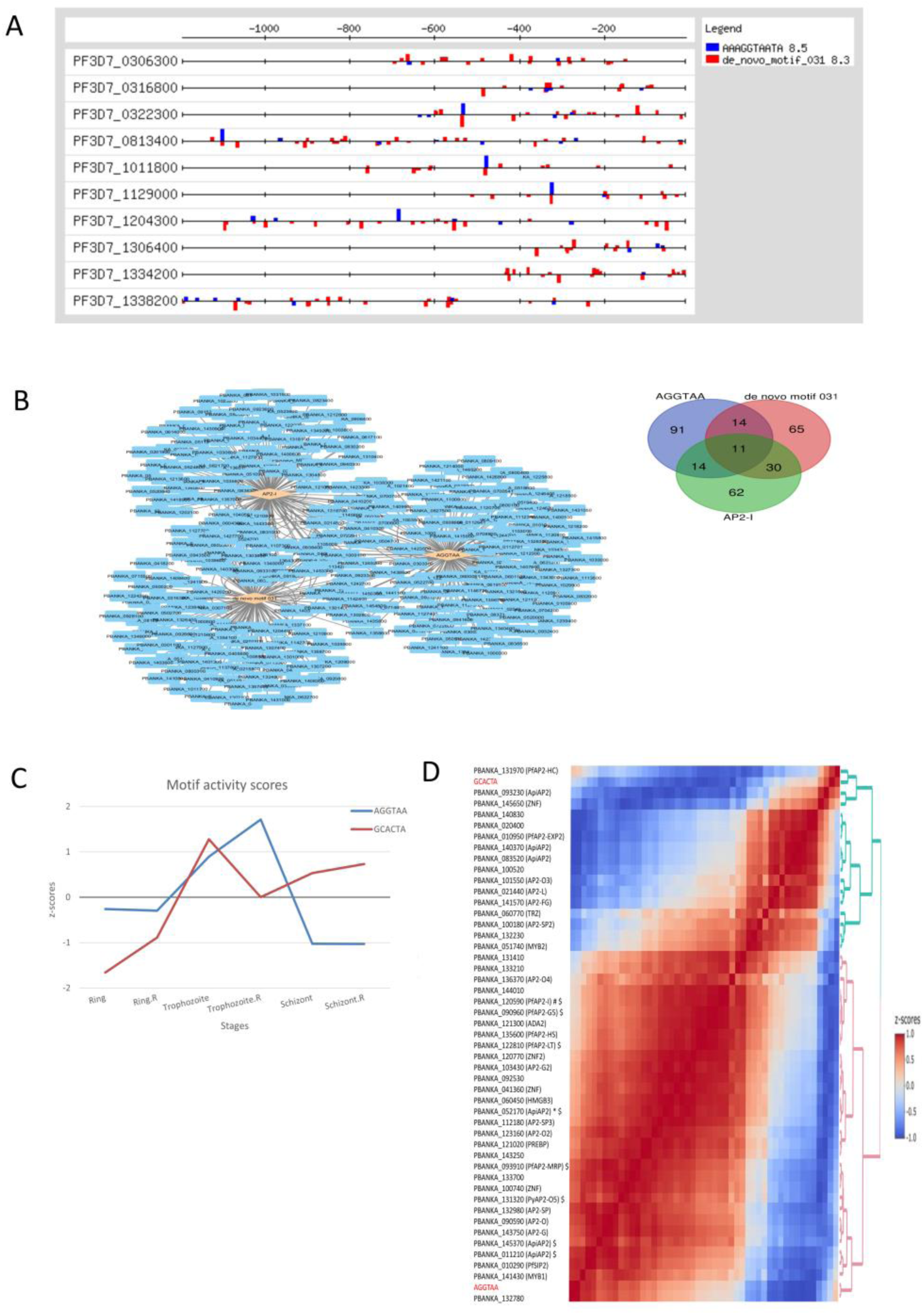
A novel motif likely coregulated with the *Pf*AP2-I-related motif and prediction of its interactors. A. Promoter architecture of some of the common target genes of the AGGTAA motif and the *de novo* motif 031 in *P. falciparum*. The promoter region of each gene represents its entire 5’UTR upstream of the start codon, and the bars represent the genomic locations of each instance of a motif. The AGGTAA motif is represented in blue, while the *de novo* motif 031 is represented in red. Overlapping instances of both motifs in opposite directions can be seen as expected for complementary sequences. B. Relationships among the AGGTAA, *de novo* motif 031 and *Pf*AP2-I motifs in *P. berghei*. Left: Gene regulatory network among the motifs (orange diamonds) and the target genes harbouring them (blue rectangles). Nodes within the network represent the genes and their regulators. The edges (grey lines) denote the physical and/or regulatory connections between nodes. Right: Intersection of target genes harbouring the motifs. C. Activities of the AGGTAA and GCACTA motifs in *P. berghei* as inferred by MARA [32]. The motif activities are scaled by z-transformation. The activity of the AGGTAA motif is represented by the blue line, while that of the GCACTA motif is represented by the red line. The motif activities are shown during the ring, trophozoite and schizont stages of asexual blood stage development. R represents biological replicates. D. Heatmap of the cross-correlation of inferred motif activity scores for the AGGTAA and GCACTA motifs and the expression values of known transcription factors and coactivators in *P. berghei*, scaled by z-transformation. The rows represent the motifs and regulatory proteins. The AGGTAA and GCACTA motifs are highlighted in red. ∗ and ♯ denote the potential direct and indirect interactors of the AGGTAA motif, respectively. $ denotes orthologues of members of the *Pf*MORC and GCN5 protein complexes, chromatin-associated proteins.

According to the motif activity inference analysis, the GCACTA motif is active throughout the schizont and trophozoite stages and inactive during the ring stage, whereas the AGGTAA motif is active during the trophozoite stage (Fig. 7C). Thirty of the forty-seven TFs that were examined during the cross-correlation analysis were found to correlate well with the AGGTAA motif, and the rest were found to correlate with the GCACTA motif. Remarkably, the well-characterized *Pf*ApiAP2-I (PF3D7 1007700/PBANKA 1205900) and the unknown ApiAP2 PBANKA 0521700/PF3D7 0420300, which were previously demonstrated to be involved in a direct interaction (**<u>T</u>**T**<u>TTAC</u>**CTTT vs. G**<u>T</u>**G**<u>TTAC</u>**A, respectively), were among the potential interactors of the AGGTAA motif revealed by cross-correlation (Fig. 7D).

## Discussion

### Transcriptomic model of the *Plasmodium* life cycle

Two main clusters of parasite stages were found in the datasets using Spearman correlation analysis with clustering and PCA of the most variable genes. The expression profiles of the asexual blood and liver stages clustered together nicely in the first cluster, whereas gametocytes, ookinetes, and sporozoites made up the majority of the second cluster (Fig. 1A). This clustering pattern was observed specifically along PC2 in the PC analysis (Fig. 1B). Previous research has also shown similar clustering patterns [8,10]. In general, the two clusters of stages can be described as primarily consisting of (i) intracellular, metabolically active stages and (ii) extracellular, sexual, and motile stages (Fig. 1; [8,10]). This is in contrast with the canonical description of the *Plasmodium* life cycle, which describes the life cycle in terms of the host environment. With the exception of gametocyte formation from asexual progenitors in host red blood cells, the life cycle is essentially the same in this “host model.” Transcriptomics, however, has made it possible to add more details to the life cycle description. The gametocytes in this “transcriptomic model” resemble the motile, extracellular stages of the mosquito vector more closely than the metabolically active, internal asexual stages found in the vertebrate host.

### Stage-specific transcriptome of *P. berghei*

Differential gene expression analysis was employed to identify the genes that were differentially expressed at each stage (Fig. 2). This consequently revealed the biological processes related to the asexual and sexual development of parasites as well as host-parasite interactions (Fig. 3). The L6h data had the fewest reads mapping to genes and thus the fewest DEGs discovered when compared to the other stages with at least 1,000 genes (Fig. 2A). This is most likely because at the L6h stage, there are fewer transcripts from the parasite than from the host cell [8]. These DEGs are functionally relevant (Additional file 3). From L24h to L60h, there was a steady enrichment of fatty acid and lipid metabolism-related processes in the liver stages (Fig. 3). This is because lipids and fatty acids are involved in a wide range of biological processes, but they are especially crucial for the asexual reproduction of *Plasmodium* parasites in the liver. Each parasite produces thousands of merozoites, and membrane biogenesis is the primary reason for this high requirement for fatty acids [37]. The type 2 fatty acid (FASII) pathway is a *de novo* process used by *Plasmodium* parasites to synthesise fatty acids in the apicoplast. A significant byproduct of the FASII pathway is octanoyl-ACP, which acts as a precursor to lipoic acid, a cofactor for the FASII-essential apicoplast pyruvate dehydrogenase complex. After being synthesised, fatty acids are either exported to different parts of the cell or integrated into lipid precursors inside the apicoplast to be used in the synthesis of other lipids [38]. Some proteins are exported to the parasitophorous vacuole membrane (PVM) by the *Plasmodium* parasite during its intracellular stages. Among these proteins secreted during the liver stages are LISP1 and LISP2. During the asexual replication phase, LISP1 aids in the breakdown of the PVM, releasing the merozoites. However, LISP2 is localised in the PV as well as the PVM. Additionally, it is exported to the plasma membrane and cytoplasm of liver cells [39].

Merozoites enter the bloodstream after exiting liver cells and invade erythrocytes to begin many rounds of asexual replication. Well-known protein families were among the upregulated genes in the blood stages (Additional file 3). SERA proteins are essential for the rupture of mature schizonts and remain attached to the surface of merozoites until erythrocyte adhesion and invasion [3,40]. Once they enter erythrocytes, parasites alter infected cells by exporting proteins to different parts of the host cell. Because it includes the trafficking of virulence components to the host cell surface, this remodelling of erythrocytes is crucial for pathogenicity and antigenic variety, allowing the parasite to evade the host’s immune response [41,42]. The most prevalent multigene family in *Plasmodium* is the PIR family, which encodes exported proteins [7,43]. The fam-a gene family encodes proteins exported into the cytoplasm of the host erythrocyte and capable of being transported to the erythrocyte surface membrane. Additionally, Fam-b proteins are exported into the cytoplasm of host erythrocytes [7,44]. The degree of infection is also associated with the capacity of parasite-infected red blood cells to sequester in the microvasculature via interactions with the host receptor CD36, which offers an additional means of evading the host immune system [44].

By the time that sexually committed schizonts eventually mature into gametocytes, their sex has already been determined. Male gametocytes exflagellate into motile, flagellated microgametes as a result of gametocyte activation in the midgut of the mosquito [45]. By identifying environmental cues, including temperature and pH changes, as well as the presence of xanthurenic acid, a molecule unique to insects, gametocytes are able to distinguish between warm-blooded and cold-blooded hosts [46,47]. Increased intracellular calcium and cyclic guanosine monophosphate (cGMP) levels (Additional file 3) are associated with the exflagellation process [48]. Guanyl cyclase activity is stimulated by xanthurenic acid, which leads to cGMP signalling and gametogenesis [49]. Despite being expressed in gametocytes, GCbeta is necessary for ookinete motility, as evidenced by the fact that disrupting GCbeta in *P. berghei* inhibits ookinete motility but does not influence exflagellation [50]. Also among the upregulated genes were sex-specific proteins and enzymes such as kinases and lipases (Additional file 3). Phospholipase C [51] mediates calcium signalling, which in turn activates CDPK4 [52], a calcium-dependent protein kinase. Cell cycle progression and the polymerisation of axonemes, the motile backbone of microgametes, are initiated by CDPK4. The subsequent generation of male gametes during cytokinesis is mediated by the mitogen-activated protein kinase MAPK2 [53]. Male fertility and the fusion of a male gamete with a female gamete to generate a zygote depend on the male-specific protein HAP2 [54]. Meiosis is preceded by nuclear fusion and cell fusion. The kinases NEK2 and NEK4 initiate genome replication during nuclear fusion, resulting in the production of a zygote with a higher concentration of nuclear DNA [55,56]. A hydrophilic protein, G377, is located within gametocyte-specific secretory organelles called osmiophilic bodies, which mediate the egress of activated female gametocytes from host erythrocytes [57]. MDV1 is present in both sexes, but it is more prevalent in females, where it is linked to osmiophilic bodies. It has also been implicated in the egress of gametes [58]. Moreover, surface proteins were upregulated (Additional file 3). Surface proteins are expressed in concert during both gametocytogenesis and gametogenesis and are targets for transmission-blocking agents. The upregulated surface proteins included the EGF domain-containing proteins P25 and P28, the multiadhesion domain CCp proteins, and the cysteine motif-rich proteins P230 and P48/45. While P25 and P28 are translationally repressed by DOZI until gametocyte fertilisation in the mosquito midgut, P230, P48/45, and CCp proteins are expressed in developing gametocytes [59,60].

The resulting diploid zygote then undergoes nuclear division to form a motile, tetraploid ookinete [61]. Ookinete gliding motility, which is promoted by phosphorylation via calcium-dependent protein kinase 3 (CDPK3), is essential for traversing the midgut epithelial wall during parasite transmission to the mosquito vector [62] (Additional file 3). Certain enzymes, including lipases, proteases, and pore-forming proteins, are also involved in penetrating the cell membrane (Additional file 3). Among the upregulated genes (Additional file 3), four different proteins have been identified as necessary for cell traversal: phospholipase (PL), SPECT, SPECT2, and cell traversal protein for ookinete and sporozoite (CelTOS) [63]. Among these proteins, pore formation has been linked to SPECT2 [64] and PL [65], whereas CelTOS [66] is thought to be required for movement in the host cell cytoplasm. Before reaching the basal lamina, the ookinete passes through one or more epithelial cells. There, it forms an oocyst, which later matures into hundreds of motile sporozoites. These invasive sporozoites infiltrate the salivary glands of mosquitoes and, eventually, the liver cells of their vertebrate host. Prior to invasion, liver cell surface proteoglycans are recognised by the sporozoite surface proteins CSP and TRAP (Additional file 3). Similarly, sporozoites must penetrate the liver sinusoidal layer and pass through numerous hepatocytes before reaching the final hepatocyte where infection is established to infect liver cells [63].

Specific regulatory sequence motifs are associated with the dynamics of transcription Redundant motifs are frequently found when high-throughput datasets, such as those from ChIP-Seq or RNA-Seq, are analysed (Fig. 5A-B). It is also typical to employ several *de novo* motif discovery approaches concurrently, enabling one to take advantage of their complementary qualities. Furthermore, the majority of motifs will be independently found by multiple tools, although some motifs may be identified solely by one tool. This means that redundant motifs with slight variations in length and/or nucleotide frequencies at specific positions may arise, and this redundancy can be reduced by employing motif clustering (Additional files S9 – S10). Unrelated TFs also exhibit similar patterns [67], which explains why a particular motif could resemble one or more binding sites for TFs in the ApiAP2, ATAC-Seq, and JASPAR motif collections (Fig. 5A-B, Additional files S11 and S12).

Many studies have been conducted on the transcriptional regulation of gene expression, especially in *P. berghei* and *P. falciparum,* during the IDC and mosquito phases. While the majority of the AP2s have well-established biological roles in regulating the transitions between distinct developmental stages, the functions of other AP2s remain unknown. The majority of AP2 factors are essential for pathogenesis, optimal morphology and maturation, host cell invasion, sexual commitment, and sex differentiation processes [68]. The *de novo* discovered motifs were compared with the 26 experimentally obtained PWMs of the AP2s, which demonstrated varying degrees of similarity between them (Fig. 5C).

AP2s are often named based on the direct transcriptional activator roles they play at the relevant developmental stage. Consequently, parallels to well-known AP2s with the underlying expression profiles of the genes in the corresponding coexpression clusters were observed (Fig. 5C). The expression of AP2-SP in Cluster 1 and all of its subclusters typically occurs during the late oocyst and sporozoite stages [35,69]. For proper ookinete development and oocyst production, AP2-O (clusters 1 and 3, subclusters 6, 7, 9 and 12) is required [35,70–72]. Proper ookinete morphogenesis and subsequent differentiation depend on AP2-O2 (clusters 1 and 3, subclusters 3, 5, 6, 8, 9 and 12), which is normally expressed during sexual differentiation up to the oocyst stage [35,72]. A component of oocyst formation and subsequent differentiation, AP2-O4 (Cluster 1), is only expressed during the sexual and mosquito phases [35,72].

Prior to sexual differentiation, there is sexual commitment by AP2-G (Cluster 1 and subclusters 2, 10, 11 and 12) [73–75]. AP2-G is an essential regulator of gametocytogenesis since it initiates the differentiation programme [76]. AP2-G2 (cluster 1 and subclusters 2, 10 and 12) functions downstream of AP2-G in morphological differentiation and maturation, exflagellation, and gametocyte transfer to mosquitoes [74,77]. Furthermore, in gametocytes, AP2-G2 functions as a translational repressor of genes involved in the asexual stage [35]. AP2-FG, found in Cluster 1 and its subclusters 2, 11, and 12, is responsible for regulating genes particular to female development. In *P. berghei*, it is only expressed in female gametocytes in the first four to six hours before sexual dimorphism in developing gametocytes appears. Its disruption also results in the restriction of female maturation but has no effect on male development [78]. In trophozoites, AP2-G4 (which functions downstream of AP2-G and is present in Clusters 1 and 3, subclusters 2-5, and 7–12) and AP2-G5, which are found in Clusters 1–3, subclusters 2, 3 and 11, are typically transcribed during sexual commitment. AP2-G5 is a repressor of commitment and early gametocyte development [79].

AP2-L (subcluster 1), which is typically expressed in the sporozoites and trophozoites of *P. falciparum* and *P. berghei*, is the only known stage-specific TF involved in parasite liver infection [80,81]. *P. vivax* has also been shown to exhibit high transcript levels in sporozoites [82]. When sporozoites enter the liver, AP2-L aids in their proper growth into the hepatic stage. Red blood cell invasion is a crucial and well-regulated process in the *Plasmodium* life cycle. *Pf*AP2-I (cluster 2, subclusters 2 and 11) is essential for controlling invasion genes during blood stage development [36]. In a similar vein, invasion gene regulation is facilitated by *Pf*AP2-LT (subclusters 2, 10 and 11), which is often expressed in the latter stages of IDC [83]. The development of the trophozoite stage and the regulation of var genes during antigenic variation, merozoite development, parasite egress, and pathogenicity during the schizont stage are dependent on a third component, *Pf*AP2-MRP (subclusters 2, 10 and 11), it is hence referred to as the master regulator of pathogenesis [84]. During blood-stage infection, periodic fever is one of the most prominent clinical signs of human malaria. The protective heat-shock response controls parasite survival and is regulated by *Pf*AP2-HS (cluster 2, subclusters 6, 7, 9, and 11). Evidence indicates that *Pf*AP2-HS may serve as a general maintainer of proteostasis in other stages in addition to its role as a protective response regulator (Cluster 2) [85].

In contrast to most ApiAP2 TFs, several have been linked to broader roles in transcriptional regulation. By binding subtelomeric and transcriptionally silent areas on chromosomes, *Pf*SIP2 (Clusters 1 and 2, subclusters 2, 6 and 11) contributes to the development of heterochromatin and the preservation of genomic integrity [86]. In contrast to *Pf*SIP2, *Pf*AP2-HC (clusters 1 and 3, subclusters 2, 6-8 and 10-12) is not involved in the development or maintenance of heterochromatin, despite being a core component of heterochromatin [87].

The discovery of *de novo* motifs in coexpression clusters that resemble uncharacterized AP2s suggested that these clusters may be under their regulation (Fig. 5C). These uncharacterized AP2s are resistant to disruption in the blood stages [35], which makes it more difficult for functional genomics research to identify their associated phenotypes. To distinguish between biologically significant and off-target binding similarities, the similarities between the motifs in the clusters and the uncharacterized AP2s were evaluated in light of their documented interactions with other proteins in the literature. There are currently few studies indicating potential functions for these AP2s. Notably, a study that examined the trophozoite and schizont stages employing blue-native PAGE combined with quantitative mass spectrometry and machine learning revealed that in *P. falciparum*, the *Pf*MORC complex is associated with ApiAP2 proteins, comprising three uncharacterized ApiAP2s (PF3D7 0420300/PBANKA 0521700, PF3D7 1239200/PBANKA 1453700, and PF3D7 0613800/PBANKA 0112100); three characterised ApiAP2s (*Pf*AP2-I/PF3D7 1007700/PBANKA 1205900, *Pf*AP2-MRP/PF3D7 1107800/PBANKA 0939100, AP2-O5/PF3D7 1449500/PBANKA 1313200); and machinery for chromatin remodelling [88]. Using targeted immunoprecipitation with LC‒MS/MS proteomic quantification during asexual blood stage development in *P. falciparum*, the associations of the ApiAP2s with the *Pf* MORC complex were also validated in another experiment [89]. The study identified six ApiAP2s previously mentioned, as well as *Pf*AP2-G5/PF3D7 1139300/PBANKA 0909600. Additionally, using a yeast two-hybrid assay of *P. falciparum*-infected erythrocytes, PF3D7 0420300 and PF3D7 1007700 were also previously linked to histone acetylation since they form a complex with the histone acetyltransferase GCN5, as well as the characterised *Pf*AP2-LT/PF3D7 0802100/PBANKA 1228100 [90]. Because there are insufficient ChIP-Seq investigations that explicitly target ApiAP2s, it is currently unknown which of the multiple-domain AP2s are involved in these interactions. For instance, each of the three AP2 domains of PF3D7 1007700 binds a different sequence, making it challenging to consistently correlate the other two AP2 domains with these protein complexes, as only the third AP2 domain of PF3D7 1007700 has been characterised. Together, these findings support the notion that the motifs resembling the binding sites of the unidentified ApiAP2s in Cluster 2 and its related subclusters may have functional roles.

A *de novo* motif that may influence chromatin structure and accessibility A similar coclustering pattern between the AGGTAA motif and a motif resembling the *Pf*AP2-I binding site, GCACTA in Cluster 2 (Additional File S9), was observed between *de novo* motif 031 and the *Pf*AP2-I binding site (TGCACTA) in another ATAC-Seq study [30]. Motif activity inference revealed that the AGGTAA and GCACTA motifs are most likely active in the trophozoite and schizont stages (Fig. 7C). Putative TFs that may control their activity were identified via cross-correlation analysis of ring stage-delayed TF expression data and leading motif activity data (Fig. 7D). Remarkably, the activity of the AGGTAA motif in the trophozoite stage corresponded better with the expression of the uncharacterized *P. berghei* orthologue for *Pf*AP2-I, PBANKA 1205900, in the ring stage than with the activity of its related binding sequence, GCACTA, in the trophozoite and schizont stages. This could be explained by the possibility that *Pf*AP2-I binds to its target gene promoters, including its promoter at the trophozoite stage, and remains associated in schizonts. It has been demonstrated by live microscopy and nuclear fractionation assays that *Pf*AP2-I-GFP localises exclusively to the nucleus of the trophozoite and schizont stages [36]. On the other hand, RNA-Seq revealed that PBANKA 1205900 is expressed as early as the ring stage and peaks in the schizont stage [7]. This shows that PBANKA 1205900 may interact with other proteins, such as the TF for the AGGTAA motif, after it is expressed in the early ring stage, before binding to its targets in the later stages. The association between trophozoite-stage AGGTAA motif activity and early expression of the uncharacterized PBANKA 0521700 was another significant but expected finding. It has been demonstrated that PBANKA 0521700 is expressed preferentially in the ring stage [7,35]. Interestingly, cross-correlation analysis also revealed a strong association between AGGTAA motif activity and the expression of members of the *Pf*MORC and GCN5 complexes, suggesting a potential interaction between chromatin-associated proteins and the TF that recognises the AGGTAA motif (Fig. 7D). Notably, DNA pull-down experiments in *P. falciparum* have demonstrated that the related motif, *de novo* motif 031, interacts with the second AP2 domain of the uncharacterized ApiAP2, PF3D7 0420300/PBANKA 0521700 [30], which is also a member of both protein complexes.

Despite having similar life cycles and biological characteristics, *Plasmodium* species differ greatly in their genetic makeup. These genetic and regulatory differences can be linked to evolutionary adaptations within the species. The variations in the genomes and regulatory components of *P. berghei* and *P. falciparum* are caused by various factors, including host specificity, transmission dynamics, and distinctive features of each species’ life cycle. These variations could manifest as different patterns of gene expression, species-specific TF involvement, or motifs. As such, interpretations of species-specific mechanisms should be made cautiously. In comparison to other eukaryotes, *Plasmodium* parasites lack a lot of TFs. To compensate for this, *Plasmodium* parasites have also developed some characteristics [91]. Research points to a potentially limited role for the ApiAP2 TFs in controlling erythrocytic transcription, especially during the asexual phases. This suggests that while the interaction of AP2s with specific promoters can lead to either repression or activation of gene expression, this interaction is dependent on the presence of many chromatin-associated proteins (apart from the few AP2s with additional chromatin-related functions) and a favourable epigenetic environment, both of which aid in the transcription process [92]. Consequently, studies involving a more thorough examination of the interactions between all the mechanisms would contribute to our knowledge of holistic gene regulation in *Plasmodium*.

## Conclusion

In addition to providing insightful information about transcriptional regulation throughout the life cycle of *P. berghei*, this study revealed patterns of gene transcription at different stages of development. Based on the different expression profiles of the DEGs across the *P. berghei* life cycle, unique clusters were identified. Additionally, a thorough examination of the promoter regions of the genes in each cluster with similar expression profiles was performed to identify shared DNA binding motifs that would aid in our continued comprehension of transcriptional regulation in *P. berghei*. In particular, the smaller clusters allowed the identification of novel DNA motifs that could be candidates for further experimental validation. Not all binding specificities for all ApiAP2 TFs have been resolved. Therefore, it is unclear whether some of the motifs discovered in this study that were not similar to the known ApiAP2 binding sites in *P. falciparum* are, in fact, genuine binding sites that have not yet been identified. Hence, to demonstrate the functional significance of these hypothesised patterns for gene regulation throughout the parasite’s life cycle, experimental validation is valuable. GFP and luciferase reporter assays have previously been used to successfully demonstrate the stage-specific activity of a promoter harbouring ApiAP2 TFs, allowing both exact quantitative analysis and qualitative analysis [93]. For example, site-directed mutagenesis was used to truncate one of the four predicted binding sites for *Pf*AP2-G5 by introducing a mutation inside its sequence in the *PbLISP2* promoter. The mutation resulted in a significant increase in luciferase activity, suggesting that the orthologue of *Pf*AP2-G5 acts as a repressor and showing that one copy of a binding site was sufficient to impact transcription. It is interesting to note that the mutation did not elevate luciferase levels in other parasite stages, maintaining the *PbLISP2* transcription profile specifically to the liver stage. Comparably, one can investigate multistage expression by combining numerous promoter components. A customised synthetic promoter combining the binding sites for *Pb*AP2-SP, *Pb*AP2-O, and *Pb*AP2-G from two genes (*PbCTRP* and *PbCSP*) that are expressed sequentially during the mosquito stages was previously reported [94]. The results of mCherry reporter gene assays showed that this promoter effectively stimulated gene expression in early to late mosquito stages. For such larger mutations, it becomes necessary to design the promoter and synthesise its sequence. It is, therefore, essential to mutate the recipient 5’ UTR sequence by introducing motifs inside it, ensuring that the new motifs are placed at the same distance from the start codon as in the native promoter. Furthermore, it is beneficial to place specific promoter elements at comparable distances from their native TSS to the recipient 5’ UTR when possible. Rapid amplification of cDNA ends (5’ RACE) is a useful tool for identifying the TSS inside targeted 5’ UTRs, as a genome-wide map of the TSS in *P. berghei* is lacking [93,95].

The other motifs were interpreted in light of the purported or reported roles of the well-known ApiAP2s at various life cycle stages. Furthermore, another novel motif was identified in the larger clusters to be similar to a newly identified motif in *P. falciparum* whose function is not well understood. Extending the annotation of this novel motif suggested its possible association with chromatin-associated proteins during erythrocytic development. The transcriptome specific to each stage, obtained from differential gene expression analysis, is a highly significant resource for the application of systems biology techniques to predict metabolic pathways important to different stages of the *P. berghei* life cycle. The data from the coexpression clustering may be helpful in creating multistage vaccination plans that depend on predetermined expression patterns. The underlying regulatory elements governing these expression patterns can be used in the creation of attenuated parasites via targeted mutagenesis employing genetic engineering techniques such as CRISPR-Cas9 or gene knockout, which disrupt or modify them.

## List of abbreviations

abs: absolute value
ApiAP2: Apicomplexan AP2
AP2: apetala2 DNA-binding domain
ATAC-Seq: assay for transposase-accessible chromatin with sequencing bp base pairs
cDNA: complementary DNA
DEGs: differentially expressed genes DNA deoxyribonucleic acid
FC: fold change GC guanyl cyclase GO gene ontology
GTF: Gene transfer format hpi hours post-infection
IDC: intraerythrocytic developmental cycle
KEGG: Kyoto encyclopedia of genes and genomes
LC-MS/MS: liquid chromatograpy with tandem mass spectroscopy log logarithm
MEME: multiple EM (expectation maximization) for motif elicitation padj adjusted p-value
PAM: partitioning around medoids
*Pb*: *Plasmodium berghei*
PBM: protein binding microarray PC principal component
PCA: principal component analysis PE paired end
*Pf*: *Plasmodium falciparum Pv Plasmodium vivax*
*PV*: *parasitophorous vacuole*
PVM: parasitophorous vacuolar membrane REVIGO reduce + visualize gene ontology RNA ribonucleic acid
RNA-Seq: RNA sequencing
RSAT: regulatory sequence analysis tool SE single end
*spp.*: species
STREME: sensitive, thorough, rapid enriched motif elicitation TF transcription factor
TSS: transcription start site
XSTREME: motif discovery and enrichment analysis
5’UTR: five prime untranslated region

## Declarations

Ethics approval and consent to participate

Not applicable

## Consent for publication

Not applicable

Availability of data and materials

Data supporting the results reported in the article are included within the article and its supplementary materials.

## Competing interests

The authors declare no competing interests.

## Funding

AO was supported by the DAAD Doctoral Research Grant 57440921.

Authors’ contributions

AO performed the analyses, interpreted the results, prepared the visualizations, and wrote the manuscript. YA performed the motif activity inference analysis and provided scientific support. BB and EA conceived and supervised the study.

## Acknowledgement

The authors would like to thank Friedrich Frischknecht for his constructive comments and suggestions both during the conception and throughout the study.

## Supplementary Information

Additional file 1: Table S1.pdf

Genome-wide transcriptome data sets retrieved from PlasmoDB release 65

Additional file 2: Table S2.pdf Overview of datasets analysed

Additional file 3: Table S3.csv

List of differentially expressed genes per parasite stage sample

Additional file 4: Figure S1.pdf

Determining the optimal number of coexpression clusters

Additional file 5: Table S4.xlsx

List of genes per coexpression cluster

Additional file 6: Table S5.xlsx

Biological processes per coexpression cluster

Additional file 7: Table S6.csv

List of genes per coexpression subcluster

Additional file 8: Table S7.xlsx

Biological processes per coexpression subcluster

Additional file 9: Supplementary File S1.zip

Motif results per cluster from RSAT and MEME and clustering of the motifs to reduce redundancy

Additional file 10: Supplementary File S2.zip

Motif results per subcluster from RSAT and MEME and clustering of the motifs to reduce redundancy

Additional file 11: Supplementary File S3.zip

Similarities of non-redundant motifs in clusters to known motifs

Additional file 12: Supplementary File S4.zip

Similarities of non-redundant motifs in subclusters to known motifs

Additional file 13: Table S8.xlsx

GO terms associated with unknown novel motifs

Additional file 14: Table S9.xlsx

KEGG pathways associated with unknown novel motifs

## Notes

### Competing Interest Statement

The authors have declared no competing interest.

