## Additional file 1 for "Transcriptome analysis reveals a novel DNA element that may interact with chromatin-associated proteins in *Plasmodium berghei* during erythrocytic development"

Table S1: Genome-wide transcriptome data sets retrieved from PlasmoDB release 65 Sept 2023.

| Data Set | Summary of study |
|---|---|
| ***P. berghei* ANKA** | |
| Exo-erythrocytic stage transcriptomes sporozoite, liver time course and detached cells [1] | Caldelari et al. 2019 did an investigation that encompassed the full transcriptome of the entire life cycle of *P. berghei*, with a meticulous temporal analysis of the mid to late liver stages 24hpi to 60hpi. |
| Female and male gametocyte [2] | Yeoh et al. 2017 propose that male gametocytes utilised existing cellular machinery over their evolutionary history, while female gametocytes primarily evolved through the emergence of new, parasite-specific pathways. |
| 5 asexual and sexual stage transcriptomes [3] | Otto et al. 2014 improved the previously existing genome assemblies. Used RNA-Seq to enhance gene-model prediction and obtain quantitative, comprehensive data on gene expression across the entire genome. |
| *P. berghei* transcriptome during inducible gametocytogenesis [4] | Kent et al. 2018 demonstrated that controlled overexpression of AP2- G may be employed to effectively and simultaneously transform most of the parasite population into fertile gametocytes, enabling the redefinition of the time period of sexual commitment, identifying several potential AP2-G targets, and mapping out the sequence of transcriptional changes throughout gametocyte development. |
| Dietary restriction of *P. berghei* WT and Δkin parasites [5] | Mancio-Silva et al. 2017 demonstrated that blood-stage parasites actively react to changes in the host's calorie intake by reorganising their transcriptome and making significant adjustments to their rate of reproduction. KIN was recognised as a crucial regulator that facilitates the detection of nutrients and governs the transcriptional reaction to the nutritional condition of the host. |
| Transcriptomes of viable ApiAP2 mutants by directional RNA-Seq [6] | Modrzynska et al. 2017 conducted a thorough knockout screen of 25 potential ApiAP2 TFs. The cellular and molecular characterization of the 11 viable mutants discovered uncovered intricate connections between positive and negative regulators that govern the life cycle at various stages. The ApiAP2 TFs form intricate regulatory networks with multiple functions. AP2-G2 was found to be a transcriptional repressor in both asexual and sexual processes. |
| Transcriptome during early and mid-stage *P. berghei* liver infection [7] | Toro-Moreno et al. 2020 also did an investigation that encompassed the full transcriptome of the entire life cycle, with a meticulous temporal analysis of the early to mid liver stages 2hpi to 48hpi. |
| ***P. falciparum*** | |
| Intraerythrocytic development cycle transcriptome [8] | Toenhake et al. 2018 mapped the chromatin structure of IDC parasites throughout the genome using ATAC-Seq. This mapping was then used to analyse the relationship between regulatory DNA elements and TFs that regulate gene expression throughout IDC by integrating RNA-Seq data. They conclude that patterns of chromatin accessibility can be used to predict the dynamics of transcription. Accessible regions are enough to induce gene expression at specified stages and ApiAP2 TFs interact with predicted cis-regulatory regions. |
| Intraerythrocytic cycle transcriptome 3D7 | unpublished |

Table S1 continued

| Data Set | Summary of study |
|---|---|
| Polysomal and steady-state asexual stage transcriptomes [9] | Bunnik et al. 2013 compared the levels of steady-state mRNA with that of polysomal mRNA at various time points throughout the IDC and noticed a delay in the peak levels of transcripts in the polysomal fraction compared to the steady-state fraction for over 30% of its genes, indicating the presence of significant translational regulation. |
| Ribosome and steady-state mRNA-Seq of asexual cell cycle stages [10] | Caro et al. 2014 reported a strong connection between transcription and translation. Specifically, for fewer than 10% of the transcriptome, direct control over translation occurs. Translationally regulated genes are mostly linked to the functions of merozoite egress. Messenger RNA 5′ leader sequences, 3′ UTRs, and antisense transcripts are delineated, while also determining the ribosome occupancy for each. Furthermore, it is confirmed that the aggregation of ribosomes on 5′ leaders is a prevalent characteristic of transcripts. |
| Intraerythrocytic development cycle Transcriptome by DAFT-Seq and UTR- Seq 3D7, HB3, IT [11] | Chappell et al. 2020 developed a novel RNA-Seq protocol which is directional and amplification-free DAFT-Seq to minimise bias against AT-rich cDNA. Although there were strain-specific variations, the overall transcriptional patterns were conserved. Their datasets allowed them to refine the 5′ and 3′ UTRs, which can be varied, lengthy > 1000 nt, and often overlap those of nearby transcripts. |
| Mosquito or cultured sporozoites and blood stage transcriptome NF54 | unpublished |
| Transcriptome of the asexual life stages [12] | Tang et al. 2020 demonstrated the co-localization of histone H3 acetylations H3K18ac and H3K27ac, as well as the variant histone *Pf* H2A.Z, at regulatory regions located upstream of genes. Integrated ChIPseq and RNAseq to identify putative regulatory sequences and showed that in schizonts, H3K18ac, H3K27ac and *Pf* H2A.Z colocalise with *Pf* AP2-I and the bromodomain protein PfBDP1. |
| NSR-Seq Transcript Profiling of malaria-infected pregnant women and children [13] | Vignali et al. 2011 used NSR-Seq to validate previous microarray investigations that showed increased expression of a subset of genes in parasites that infect pregnant women, including the gene encoding var2csa, a widely recognised vaccine candidate against pregnancy-related malaria. They additionally delineated a subset of parasite transcripts that differentiate parasites infecting children from those infecting pregnant women. |
| Asexual blood stages, salivary gland sporozoite and midgut oocyst transcriptomes [14] | Gómez-Díaz et al. 2016 reported that the expression of the PF3D7_1255200 *var* gene in the oocyst is associated with low levels of H3K9me3 at the PF3D7_1466400 AP2 TF binding site. This is also associated with the expression of an antisense lncRNA that was previously demonstrated to enhance *var* gene transcription throughout the IDC *in vitro*. The expression of both the sense protein-coding transcript and the antisense lncRNA significantly increased in sporozoites, indicating that the activation of a specific *var* gene involves an intricate mechanism that includes AP2 TFs and lncRNAs. |
| Trophozoite and Schizont transcriptomes of *Pf* BDP1HA [15] | Josling et al. 2015 identified the bromodomain protein *Pf* BDP1 as a critical activator of essential genes during erythrocyte invasion. *Pf* BDP1 attaches to chromatin located at the promoters of invasion genes and its depletion leads to the dysregulation of invasion gene expression and the dis- ruption of invasion. |

Table S1 continued

| Data Set | Summary of study |
|---|---|
| Transcriptome of sequestration phenotypes [16] | Kamaliddin et al. 2021 reported a strong positive correlation between the mRNA and protein levels in both the ring-stage transcriptomes and trophozoite proteomes. However, twice the level of transcripts were identified at the ring stage than the level of proteins identified in the mature trophozoite stage, indicating that the presence of a significant level of transcript expression does not necessarily ensure the identification of the associated protein. Nevertheless, the level of mRNA does not accurately represent the expression of proteins at the individual level. |
| Gametocyte Transcriptomes [17] | Lasonder et al. 2016 presented the first transcriptome study of male and female *Pf* gametocytes, together with a comprehensive proteome analysis. Male gametocytes were found to have a higher concentration of proteins that play a role in the development of flagellated gametes, as well as proteins involved in chromatin organisation, DNA replication, and axoneme formation. In contrast, female gametocytes possessed a higher concentration of proteins necessary for zygote development and post-fertilization processes, including protein, lipid, and energy metabolism. |
| Dual transcriptomes of malaria-infected Gambian children [18] | Lee et al. 2018 identified differentially expressed human and parasite genes were identified in severe vs uncomplicated malaria, exhibiting distinct patterns specifically linked to coma, hyperlactatemia, and thrombocytopenia. In all severe malaria phenotypes, there was a clear correlation with elevated expression of genes related to neutrophil granules. They also noticed severity-associated variation in the expression of parasite genes responsible for cytoadherence to the vascular endothelium, stiffness of infected erythrocytes, and parasite growth rate. The majority of variable gene expression in severe malaria, accounting for up to 99%, was influenced by variations in parasite load. Conversely, there was minimal correlation between parasite gene expression and parasite load. |
| Oocyst and salivary gland sporozoite transcriptome comparison in *P. falciparum* [19] | Lindner et al. 2019 used dynamic gene expression profiles and protein compositions of oocyst sporozoites and salivary gland sporozoites to identify the mRNAs and proteins that are upregulated in oocyst sporozoites UOS or upregulated in infectious sporozoites UIS. Furthermore, they discovered that malaria parasites employ two overlapping, comprehensive, and separate mechanisms of translational repression during sporozoite development to control the timing of protein expression during sporozoite maturation, migration, and infection, all of which facilitate their effective development and passage from the vector to the host. |
| Intraerythrocytic developmental cycle by RNA-Seq [20] | Otto et al. 2010 used RNA-Seq to acquire novel understanding of the *Pf* transcriptome during the IDC: to identify previously unknown gene transcripts, rectify numerous gene models, propose alternative splicing events, and predict the 5’ and 3’ UTRs. |
| IDC in constant temperature and darkness | Subudhi et al. 2020 found that 6% of *Pf* genes show 24-hour rhythms in expression under free-running conditions. The serpentine receptor 10 gene SR10 has a 24-hour rhythm in transcription. |
| Heat shock response in sensitive mutants LRR5, DHC [21] | Zhang et al. 2021 developed an extensive phenotypic screening system and used it to conduct a comprehensive forward-genetic phenotype screen, uncovering genes that enable parasites to withstand febrile temperatures. Crucial key processes involve essential, conserved pathways related to protein folding, heat shock response, and proteasomal degradation, and surprisingly, isoprenoid biosynthesis. |

Table S1 continued

| Data Set | Summary of study |
|---|---|
| Comparison of *in vitro* versus aseptic mosquito-produced sporozoites by RNA-Seq [22] | Eappen et al. 2022 successfully generated of a vast quantity of *in vitro Pf* SPZ iPfSPZ which were shown to successfully infiltrate human hepatocytes in a laboratory setting and progressed into fully grown liver-stage schizonts that displayed merozoite surface protein 1 *Pf* MSP1 in quantities similar to mosquito *Pf* SPZ m*Pf* SPZ, and also generate both asexual and sexual erythrocytic-stage parasites in laboratory conditions. Additionally, gametocytes developed into *Pf* SPZ when consumed by mosquitoes, thereby completing the entire life cycle. |
| High-resolution intraerythrocytic time course transcriptome by RNA-Seq [23] | Kucharski et al. 2020 developed a detailed approach for efficiently preserving, extracting, amplifying, and sequencing whole transcripts from *Plasmodium*-infected blood. The methodology provided is a comprehensive process for producing high-quality transcriptomic data from a wide range of blood samples, which have different levels of parasitaemia and RNA inputs. The adaptability of this pipeline to robotic handling will encourage future transcriptomic investigations in the field of malaria, both on a small and big scale. |
| *Pf* 3D7 Ralph nanopore RNA-Seq | unpublished |
| Strand-specific transcriptome of the intraerythrocytic developmental cycle [24] | Siegel et al. 2014 performed a strand-specific analysis of the transcriptome which enhanced and reinforced the existing body of proof that antisense transcription is a significant occurrence in *Pf*. A complex relationship exists between the levels of sense and antisense transcripts for a specific group of adjacent genes, but other natural antisense transcripts seem to be regulated independently from mRNA production. |
| Ring, Oocyst and Sporozoite Transcriptomes [25] | Zangh`ı et al. 2018 found that a strain-specific component of *Pf* EMP1 is expressed on the sporozoite surface NF54 SpzPfEMP1. Antibodies targeting this protein hindered the strain-specific entry of sporozoites into hepatocytes. Sporozoites induced the expansion of sub-telomeric heterochromatin in order to suppress the expression of genes specific to the blood stage. The protein elicited an immune response in human volunteers infected with sporozoites. |
| Transcriptome during intraerythrocytic development [26] | Bártfai et al. 2010 presented a new, linear amplification NGS technique that enabled unbiased investigation of the highly AT-rich *Plasmodium* genome. The technique was employed to conduct a detailed examination of a specific histone H2A variation, H2A.Z, as well as two histone H3 modifications across the entire genome during the IDC. In contrast to other organisms, H2A.Z is consistently present in euchromatic intergenic areas during the whole intraerythrocytic cycle. The high degree of colocalization between H2A.Z and H3K9ac and H3K4me3 indicates that these marks are mostly deposited on nucleosomes containing H2A.Z. |
| Transcriptomes of 7 sexual and asexual life stages [27] | López-Barragán et al. 2011 generated data of seven bidirectional libraries from the ring, early and late trophozoite, schizont, gametocyte II and V, and ookinete stages. The cDNA sequences were aligned to the 3D7 reference genome, which resulted in the identification of stage-specific antisense transcripts and previously unknown intron-exon splicing junctions. The existence of antisense transcripts in some gametocyte and ookinete genes implies that these antisense RNA molecules may have a significant impact on the regulation of gene expression and the development of the parasite. |
| Strand-specific transcriptomes of 4 life cycle stages [27] | López-Barragán et al. 2011 also generated four strand-specific libraries from the late trophozoite, schizont and gametocyte II and V stages. Sequencing of strand-specific cDNA libraries indicated a higher expression of genes in one direction in gametocyte compared to schizont. |

Table S1 continued

| Data Set | Summary of study |
|---|---|
| Transcriptome in severe vs uncomplicated malaria [28] | Tonkin-Hill et al. 2018 found that the transcriptomes of some severe malaria parasite isolates revealed a decrease in glycolysis without the ac- tivation of compensatory pathways. Additionally, there were alterations in chromatin structure, likely affecting transcriptional regulation through a de- crease in histone methylation. Furthermore, there was a reduction in the surface expression of *Pf* EMP1 and a down-regulation of multiple chaperone proteins. The analysis also discovered new connections between the severity of the disease and certain *Pf* EMP1 transcripts, domains, and smaller sequence regions. |
| Intraerythrocytic development cycle transcriptome [29] | Wichers et al. 2019 used Deep RNA-Seq to investigate *stevor* gene expression throughout the IDC, including free merozoites. Majority of *stevor* genes exhibited a pattern of expression characterised by two distinct peaks: the first peak occurred during the trophozoite stages, whereas the second peak occurred in late schizonts. |
| ***P. vivax*** | |
| Hypnozoite [30] | Gural et al. 2018 successfully replicated the complete liver stage of *Pv* in a laboratory setting, which involved the creation and reactivation of hypnozoites with the release of merosomes and infection of reticulocytes. Hybrid capture, followed by RNA-Seq, provided an initial glimpse into the transcriptome of hypnozoites. |
| Transcription profile of intraerythrocytic cycle [31] | Zhu et al. 2016 produced a comprehensive transcriptome map providing novel insights into the regulatory processes of individual genes and uncovered their close association with certain biological functions. An area of high transcription activity for vir genes was identified on chromosome 2, indicating the presence of a possible active site that regulates immune evasion in different patients. *Pv* genes exhibited elongated 5'UTR and numerous transcription start sites, distinguishing them from other eukaryotes. Also, in *Pv*, alternative splicing is infrequent. However, its connection to the late schizont stage implies that it plays a role in gene function. |
