## Additional file 2 for "Transcriptome analysis reveals a novel DNA element that may interact with chromatin-associated proteins in *Plasmodium berghei* during erythrocytic development"

Table S2: Overview of datasets analysed

| **Accession number** | **Species/ Strain** | **Library type** | **RNA Selection** | **Instrument** | **Read length** | **Layout** | **Adapter** | **Stages analyzed** | **Study** |
|---|---|---|---|---|---|---|---|---|---|
| ERP105548 | *Pb*ANKA | U | Oligo-dT | Illumina HiSeq 2500 | 100 | PE | Nextera | Sporozoite, liver stages | [1] |
| SRP250329 | *Pb*ANKA | U | cDNA | Illumina HiSeq 4000 | 50 | SE | SMART-Seq | Sporozoite, liver stages | [2] |
| SRP027529/ ERP004740 | *Pb* | U | | Illumina Genome Analyzer IIx | 76 | SE/PE | Truseq | Ring, Trophozoite, Schizont, Gametocyte, Ookinete | [3] |
| SRP099925 | *Pb*ANKA | S (BS2-3) / SR (BS1) | Poly(A) | Illumina HiSeq 2500 | 100 | PE | | Asexual blood stages (mixed) | [4] |
| SRP197607 | *Pb*ANKA | SR | PCR | Illumina NextSeq 500 | 75 | PE | KAPA | Gametocyte | [5] |
| SRP073801 | *Pb*ANKA | SR | cDNA | Illumina HiSeq 2500 | 100 | PE | NEBNext | Ookinete | [6] |
| SRP090611 | *Pf*NF54 and 3D7 | SR | Random | Illumina NextSeq 550 | 150 | SE | Truseq | Gametocyte, Sporozoite | [7] |
| SRP142460 | *Pf*3D7 | SR | cDNA | Illumina HiSeq 2500 | 100 | SE | Truseq | Sporozoite | [8] |
| SRP048710 | *Pf*K1 | SR | cDNA | Illumina MiSeq | | PE | Truseq | Ring, Trophozoite, Schizont | [9] |
| SRP211863 | *Pf*NF54 | SR | cDNA | Illumina HiSeq X Ten | 150 | PE | KAPA AGATCGGAAGAGC AAGATCGGAAGAGC | Ring, Trophozoite, Schizont | [10] |
| SRP069075 | *Pf*3D7 | SR | cDNA | Illumina HiSeq 2500 | | SE | TruSeq | Male and female gametocyte | unpublished |
| SRP100893 | *Pv* | U | Random | Illumina HiSeq 2000 (NextSeq) | | PE | TruSeq | Sporozoite | [11] |
| SRP046739 | *Pv* | U | cDNA | Illumina HiSeq 2000 | 100 | PE | TruSeq | Blood stages | [12] |

S: stranded, SR: stranded reverse, U: unstranded, SE: single end, PE: paired end, oligo-dT: oligo-deoxythymidine
