## Additional file 4 for "Transcriptome analysis reveals a novel DNA element that may interact with chromatin-associated proteins in *Plasmodium berghei* during erythrocytic development"

Supplementary figure

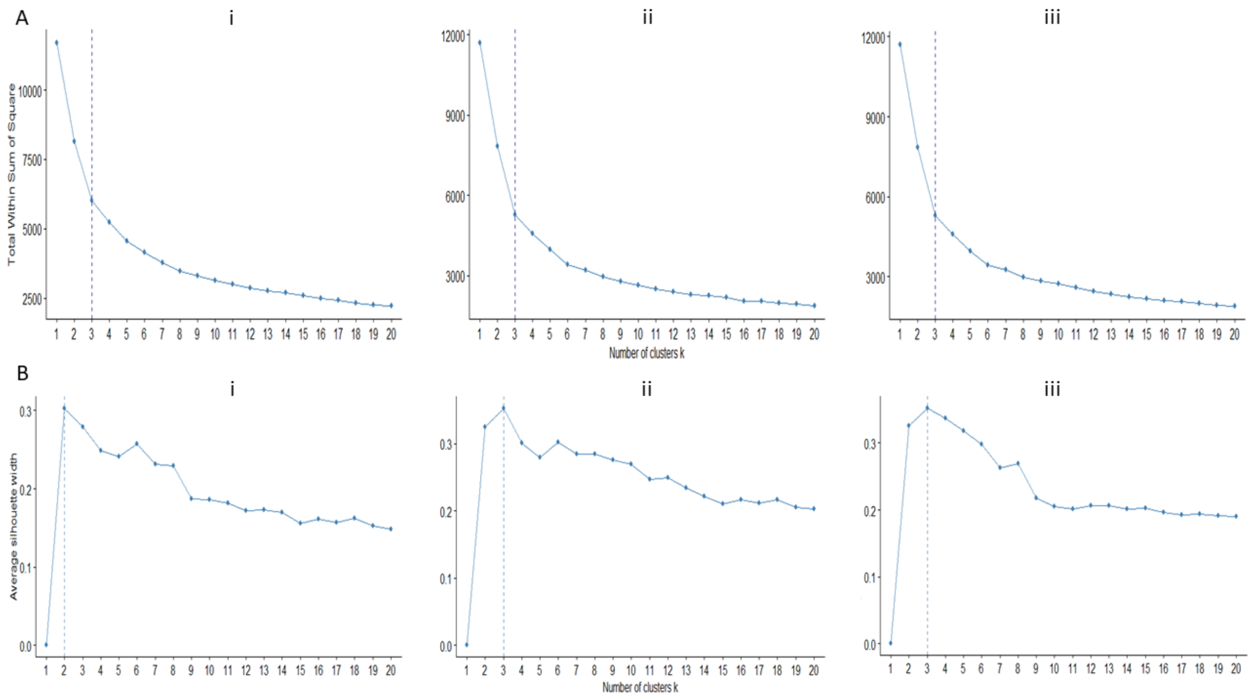


Figure S1: Determining the optimal number of co-expression clusters by A. the elbow method and B. the average silhouette width method. The number of clusters is determined for three algorithms: i. hierarchical ii. K-means and iii. PAM clustering. The vertical dashed line indicates the optimal number of clusters for each algorithm in each method.
