## Additional file 9 for "Transcriptome analysis reveals a novel DNA element that may interact with chromatin-associated proteins in *Plasmodium berghei* during erythrocytic development": matrix-clustering_SUMMARY_cluster1.html

matrix-clustering coexpression\_cluster1

### RSAT - matrix-clustering result

##### Analysis: coexpression\_cluster1 (07/09/2023 11:03)

##### Command

```
matrix-clustering  -v 1 -max_matrices 300 -matrix coexpression_cluster1 $RSAT/public_html/tmp/www-data/2023/09/07/matrix-clustering_2023-09-07.110304_kSNkrp/matrix-clustering_query_matrices.transfac transfac -hclust_method average -calc sum -title coexpression_cluster1 -metric_build_tree Ncor -lth w 5 -lth cor 0.6 -lth Ncor 0.4 -quick -label_in_tree name -return json,heatmap -o $RSAT/public_html/tmp/www-data/2023/09/07/matrix-clustering_2023-09-07.110304_kSNkrp/matrix-clustering
```

#### Results Summary

|  |  |  |  |  |  |
| --- | --- | --- | --- | --- | --- |
| Nb Input motifs | Nb Input collections | Nb Clusters Found | Download   Root motifs | Complete results [zip] | Parameters used Linkage method \| Similarity metric \| Thresholds to partition the tree |
| 44 | 1 | 13 | **Download** | Download | average \| Ncor \| **Ncor** = 0.4 **cor** = 0.6 |

|  |  |  |  |  |  |  |
| --- | --- | --- | --- | --- | --- | --- |
|  | **Logo Forest (dynamic browsing)**  **Logo Forest (rapid overview - low image quality)**  cluster\_1  cluster\_2  cluster\_3  cluster\_4  cluster\_5  cluster\_6  cluster\_7  cluster\_8  cluster\_9  cluster\_10  cluster\_11  cluster\_12  cluster\_13 |  |  |  |  |  |
|  | **Clusters Summary** Clusters Summary Table (Root Motif \| Root Motif (Reverse) \| Cluster ID \| # Motifs \| Motif\_Name::Collection \| Number of Motifs by Collection \| Root Motif (transfac Format)) \| | cluster\_1 | 17 | cluster1\_motif33\_m::coexpression\_cluster1:: cluster1\_motif11\_r::coexpression\_cluster1:: cluster1\_motif2\_r::coexpression\_cluster1:: cluster1\_motif12\_r::coexpression\_cluster1:: cluster1\_motif1\_r::coexpression\_cluster1:: cluster1\_motif15\_r::coexpression\_cluster1:: cluster1\_motif7\_r::coexpression\_cluster1:: cluster1\_motif8\_r::coexpression\_cluster1:: cluster1\_motif3\_r::coexpression\_cluster1:: cluster1\_motif25\_r::coexpression\_cluster1:: cluster1\_motif9\_r::coexpression\_cluster1:: cluster1\_motif21\_r::coexpression\_cluster1:: cluster1\_motif40\_m::coexpression\_cluster1:: cluster1\_motif41\_m::coexpression\_cluster1:: cluster1\_motif28\_m::coexpression\_cluster1:: cluster1\_motif32\_m::coexpression\_cluster1:: cluster1\_motif34\_m::coexpression\_cluster1:: | coexpression\_cluster1::17\_motifs | Show matrix |
| cluster\_2 \| 4 \| cluster1\_motif20\_r::coexpression\_cluster1:: cluster1\_motif24\_r::coexpression\_cluster1:: cluster1\_motif23\_r::coexpression\_cluster1:: cluster1\_motif5\_r::coexpression\_cluster1:: \| coexpression\_cluster1::4\_motifs \| Show matrix \| \| |  |  |  |  |  |  |
| cluster\_3 \| 6 \| cluster1\_motif16\_r::coexpression\_cluster1:: cluster1\_motif17\_r::coexpression\_cluster1:: cluster1\_motif14\_r::coexpression\_cluster1:: cluster1\_motif19\_r::coexpression\_cluster1:: cluster1\_motif22\_r::coexpression\_cluster1:: cluster1\_motif6\_r::coexpression\_cluster1:: \| coexpression\_cluster1::6\_motifs \| Show matrix \| \| |  |  |  |  |  |  |
| cluster\_4 | 7 | cluster1\_motif43\_m::coexpression\_cluster1:: cluster1\_motif26\_m::coexpression\_cluster1:: cluster1\_motif27\_m::coexpression\_cluster1:: cluster1\_motif10\_r::coexpression\_cluster1:: cluster1\_motif31\_m::coexpression\_cluster1:: cluster1\_motif30\_m::coexpression\_cluster1:: cluster1\_motif4\_r::coexpression\_cluster1:: | coexpression\_cluster1::7\_motifs | Show matrix | | |
| cluster\_5 | 2 | cluster1\_motif29\_m::coexpression\_cluster1:: cluster1\_motif38\_m::coexpression\_cluster1:: | coexpression\_cluster1::2\_motifs | Show matrix | | |
| cluster\_6 | 1 | cluster1\_motif18\_r::coexpression\_cluster1 | coexpression\_cluster1 : 1 motif | Show matrix | | |
| cluster\_7 | 1 | cluster1\_motif37\_m::coexpression\_cluster1 | coexpression\_cluster1 : 1 motif | Show matrix | | |
| cluster\_8 | 1 | cluster1\_motif13\_r::coexpression\_cluster1 | coexpression\_cluster1 : 1 motif | Show matrix | | |
| cluster\_9 | 1 | cluster1\_motif36\_m::coexpression\_cluster1 | coexpression\_cluster1 : 1 motif | Show matrix | | |
| cluster\_10 | 1 | cluster1\_motif39\_m::coexpression\_cluster1 | coexpression\_cluster1 : 1 motif | Show matrix | | |
| cluster\_11 | 1 | cluster1\_motif35\_m::coexpression\_cluster1 | coexpression\_cluster1 : 1 motif | Show matrix | | |
| cluster\_12 | 1 | cluster1\_motif44\_m::coexpression\_cluster1 | coexpression\_cluster1 : 1 motif | Show matrix | | |
| cluster\_13 | 1 | cluster1\_motif42\_m::coexpression\_cluster1 | coexpression\_cluster1 : 1 motif | Show matrix | | |

- Note: by default the table displays the first 50 clusters. Click on the *Show X entries* button to display more (or less) clusters.
- Click on the column names to change the order of the data.
- Use the *Search* window to find your motif (s) of interest.

| **Individual Cluster View** Individual Cluster View **Show All** **Hide All** **cluster\_1** **cluster\_2** **cluster\_3** **cluster\_4** **cluster\_5** **cluster\_6** **cluster\_7** **cluster\_8** **cluster\_9** **cluster\_10** **cluster\_11** **cluster\_12** **cluster\_13** **Display Logo Trees** **cluster\_1** **cluster\_2** **cluster\_3** **cluster\_4** **cluster\_5** **cluster\_6** **cluster\_7** **cluster\_8** **cluster\_9** **cluster\_10** **cluster\_11** **cluster\_12** **cluster\_13** **Display Branch-Motifs** **cluster\_1** Branch Motifs \| \| \| \| \| \| \|; Node \| Consensus \| Logo \| Logo (Reverse) \| Collections \| Matrix (Transfac format) \| IC \| Nb sites; Node 1 \| **wwwAAgACAAwwww-** \| \| \| coexpression\_cluster1::2\_motifs \| PSSM \| 8.53 \| 3830; Node 2 \| **wwAAAgACAwww---** \| \| \| coexpression\_cluster1::2\_motifs \| PSSM \| 7.80 \| 5667; Node 3 \| **wwwTAgACAAwwww-** \| \| \| coexpression\_cluster1::2\_motifs \| PSSM \| 8.52 \| 3731; Node 4 \| **wwwAAgACAAwwww-** \| \| \| coexpression\_cluster1::3\_motifs \| PSSM \| 8.55 \| 6444; Node 5 \| **wwAAAgACAwwww--** \| \| \| coexpression\_cluster1::3\_motifs \| PSSM \| 8.09 \| 7402; Node 6 \| **wwaAAgAcAAwwwww** \| \| \| coexpression\_cluster1::4\_motifs \| PSSM \| 8.67 \| 8885; Node 7 \| **rAAAAAAwAAAAAAM** \| \| \| coexpression\_cluster1::2\_motifs \| PSSM \| 13.23 \| 1137; Node 8 \| **wwaWAgACAAwwwww** \| \| \| coexpression\_cluster1::6\_motifs \| PSSM \| 8.20 \| 12616; Node 9 \| **wwAAAgACAwwwaww** \| \| \| coexpression\_cluster1::4\_motifs \| PSSM \| 8.65 \| 8769; Node 10 \| **rAAAAAAarAAAAAM** \| \| \| coexpression\_cluster1::3\_motifs \| PSSM \| 12.32 \| 1412; Node 11 \| **wwATrAAcAAAwwwA** \| \| \| coexpression\_cluster1::2\_motifs \| PSSM \| 10.11 \| 4689; Node 12 \| **wwAAAgACAawwwww** \| \| \| coexpression\_cluster1::10\_motifs \| PSSM \| 7.90 \| 21385; Node 13 \| **rAAAAAAAAAwAAAM** \| \| \| coexpression\_cluster1::4\_motifs \| PSSM \| 12.20 \| 1961; Node 14 \| **waAwaAAmAAAwawA** \| \| \| coexpression\_cluster1::6\_motifs \| PSSM \| 9.10 \| 6648; Node 15 \| **wwAWArAcAAwwwwa** \| \| \| coexpression\_cluster1::16\_motifs \| PSSM \| 7.51 \| 28033; Node 16 \| **wwAWArAcAAwwwwa** \| \| \| coexpression\_cluster1::17\_motifs \| PSSM \| 7.51 \| 28333 **cluster\_2** Branch Motifs \| \| \| \| \| \| \|; Node \| Consensus \| Logo \| Logo (Reverse) \| Collections \| Matrix (Transfac format) \| IC \| Nb sites; Node 1 \| **wwaTGTTGTAww** \| \| \| coexpression\_cluster1::2\_motifs \| PSSM \| 7.92 \| 2634; Node 2 \| **wwwTGTTkTwww** \| \| \| coexpression\_cluster1::3\_motifs \| PSSM \| 7.86 \| 4889; Node 3 \| **wwwTGTTkTwww** \| \| \| coexpression\_cluster1::4\_motifs \| PSSM \| 7.67 \| 5090 **cluster\_3** Branch Motifs \| \| \| \| \| \| \|; Node \| Consensus \| Logo \| Logo (Reverse) \| Collections \| Matrix (Transfac format) \| IC \| Nb sites; Node 1 \| **wwwwaAAmAAAAATwTtTAAAtww** \| \| \| coexpression\_cluster1::2\_motifs \| PSSM \| 12.57 \| 6001; Node 2 \| **--------wwwaATTTTTAAAwww** \| \| \| coexpression\_cluster1::2\_motifs \| PSSM \| 9.22 \| 14538; Node 3 \| **wwAAaaAmAAAAAwwtwtAwAwww** \| \| \| coexpression\_cluster1::3\_motifs \| PSSM \| 11.95 \| 13464; Node 4 \| **wwAwawwaAAAAATwTwtAAAwww** \| \| \| coexpression\_cluster1::4\_motifs \| PSSM \| 11.64 \| 21224; Node 5 \| **wwAwawwaAawaATTTTTAAAwww** \| \| \| coexpression\_cluster1::6\_motifs \| PSSM \| 11.34 \| 35762 **cluster\_4** Branch Motifs \| \| \| \| \| \| \|; Node \| Consensus \| Logo \| Logo (Reverse) \| Collections \| Matrix (Transfac format) \| IC \| Nb sites; Node 1 \| **TATATRTATrTATry** \| \| \| coexpression\_cluster1::2\_motifs \| PSSM \| 13.28 \| 1269; Node 2 \| **---atACACACATww** \| \| \| coexpression\_cluster1::2\_motifs \| PSSM \| 8.69 \| 1868; Node 3 \| **TATATATATryATry** \| \| \| coexpression\_cluster1::3\_motifs \| PSSM \| 13.08 \| 1472; Node 4 \| **--waTrCATGCAyA-** \| \| \| coexpression\_cluster1::2\_motifs \| PSSM \| 7.80 \| 1602; Node 5 \| **--watrCAyrCATaw** \| \| \| coexpression\_cluster1::4\_motifs \| PSSM \| 7.45 \| 3470; Node 6 \| **TATaTryAyrYATay** \| \| \| coexpression\_cluster1::7\_motifs \| PSSM \| 9.83 \| 4942 **cluster\_5** Branch Motifs \| \| \| \| \| \| \|; Node \| Consensus \| Logo \| Logo (Reverse) \| Collections \| Matrix (Transfac format) \| IC \| Nb sites; Node 1 \| **yygtrwyAGCTAGCTrwhtwtr** \| \| \| coexpression\_cluster1::2\_motifs \| PSSM \| 9.46 \| 283 **cluster\_6** Branch Motifs \| \| \| \| \| \| \|; Node \| Consensus \| Logo \| Logo (Reverse) \| Collections \| Matrix (Transfac format) \| IC \| Nb sites; Singleton \| **wwATTACACww** \| \| \| coexpression\_cluster1 : 1 motif \| PSSM \| 7.22 \| 3137 **cluster\_7** Branch Motifs \| \| \| \| \| \| \|; Node \| Consensus \| Logo \| Logo (Reverse) \| Collections \| Matrix (Transfac format) \| IC \| Nb sites; Singleton \| **taatmatartaarr** \| \| \| coexpression\_cluster1 : 1 motif \| PSSM \| 15.20 \| 85 **cluster\_8** Branch Motifs \| \| \| \| \| \| \|; Node \| Consensus \| Logo \| Logo (Reverse) \| Collections \| Matrix (Transfac format) \| IC \| Nb sites; Singleton \| **wwTCTGAAww** \| \| \| coexpression\_cluster1 : 1 motif \| PSSM \| 9.23 \| 216 **cluster\_9** Branch Motifs \| \| \| \| \| \| \|; Node \| Consensus \| Logo \| Logo (Reverse) \| Collections \| Matrix (Transfac format) \| IC \| Nb sites; Singleton \| **tGaaCaaaty** \| \| \| coexpression\_cluster1 : 1 motif \| PSSM \| 11.11 \| 227 **cluster\_10** Branch Motifs \| \| \| \| \| \| \|; Node \| Consensus \| Logo \| Logo (Reverse) \| Collections \| Matrix (Transfac format) \| IC \| Nb sites; Singleton \| **mbSssssyGCaCaCrCvSSSSS** \| \| \| coexpression\_cluster1 : 1 motif \| PSSM \| 10.85 \| 148 **cluster\_11** Branch Motifs \| \| \| \| \| \| \|; Node \| Consensus \| Logo \| Logo (Reverse) \| Collections \| Matrix (Transfac format) \| IC \| Nb sites; Singleton \| **GCatGC** \| \| \| coexpression\_cluster1 : 1 motif \| PSSM \| 6.26 \| 168 **cluster\_12** Branch Motifs \| \| \| \| \| \| \|; Node \| Consensus \| Logo \| Logo (Reverse) \| Collections \| Matrix (Transfac format) \| IC \| Nb sites; Singleton \| **CatttCatGaCCCat** \| \| \| coexpression\_cluster1 : 1 motif \| PSSM \| 20.79 \| 6 **cluster\_13** Branch Motifs \| \| \| \| \| \| \|; Node \| Consensus \| Logo \| Logo (Reverse) \| Collections \| Matrix (Transfac format) \| IC \| Nb sites; Singleton \| **MSSkMCCCCCMVtmy** \| \| \| coexpression\_cluster1 : 1 motif \| PSSM \| 9.94 \| 25 - **Each cluster has a different color.** - **Click on the upper buttons to display separately the information of each cluster.** - **Click on the circles at each tree to display its corresponding merged motifs.** Definitions - **Logo tree.** This results shows the hierarchical tree with its logo alignment of one cluster. The logos are shown in Forward (left logo) and Reverse (right logo) orientation. - **Branch-motifs.** On the trees are displayed the branch number that is the number in which the motifs were incorporated in the tree. The Branch-Motifs table shows the logo in both orientations and the link to the file in TRANSFAC format with the branch-motif. |
|  | **Individual Motif View** Individual Motif View Motif id \| Motif name \| Cluster \| Collection \| Width \| IC \| Number of sites \| Consensus \| Consensus (Rev) \| Logo \| Logo (Rev); coexpression\_cluster1\_m38\_cluster1\_motif38\_m \| cluster1\_motif38\_m \| cluster\_5 \| coexpression\_cluster1 \| 22 \| 9.77 \| 201 \| **yygtrwyAGCTAGCTvrckmtr** \| **YAKMGYBAGCTAGCTRWYACRR** \| \|; coexpression\_cluster1\_m8\_cluster1\_motif8\_r \| cluster1\_motif8\_r \| cluster\_1 \| coexpression\_cluster1 \| 15 \| 9.39 \| 1367 \| **WWTDYATGTCTTTWW** \| **wwAAAgACATrhAww** \| \|; coexpression\_cluster1\_m18\_cluster1\_motif18\_r \| cluster1\_motif18\_r \| cluster\_6 \| coexpression\_cluster1 \| 11 \| 7.22 \| 3137 \| **wwATTACACww** \| **WWGTGTAATWW** \| \|; coexpression\_cluster1\_m32\_cluster1\_motif32\_m \| cluster1\_motif32\_m \| cluster\_1 \| coexpression\_cluster1 \| 15 \| 14.02 \| 873 \| **RAAAAAAWAARAAAA** \| **TTTTYTTwTTTTTTy** \| \|; coexpression\_cluster1\_m31\_cluster1\_motif31\_m \| cluster1\_motif31\_m \| cluster\_4 \| coexpression\_cluster1 \| 15 \| 11.62 \| 128 \| **+++ATgCACACATrT** \| **AYATGTGTGCAT+++** \| \|; coexpression\_cluster1\_m15\_cluster1\_motif15\_r \| cluster1\_motif15\_r \| cluster\_1 \| coexpression\_cluster1 \| 15 \| 8.59 \| 1677 \| **wwwAAgACAATwww+** \| **+WWWATTGTCTTWWW** \| \|; coexpression\_cluster1\_m28\_cluster1\_motif28\_m \| cluster1\_motif28\_m \| cluster\_1 \| coexpression\_cluster1 \| 15 \| 11.68 \| 276 \| **+TYTTTCTCTTTTT+** \| **+AAAAAGAGAAArA+** \| \|; coexpression\_cluster1\_m14\_cluster1\_motif14\_r \| cluster1\_motif14\_r \| cluster\_3 \| coexpression\_cluster1 \| 24 \| 11.36 \| 7760 \| **+++WWWTTTAAAAATTTTTTWWWW** \| **wwwwAAAAAATTTTTAAAwww+++** \| \|; coexpression\_cluster1\_m17\_cluster1\_motif17\_r \| cluster1\_motif17\_r \| cluster\_3 \| coexpression\_cluster1 \| 24 \| 9.39 \| 9276 \| **++++++++WWWAATTTTTAAAWWW** \| **wwwTTTAAAAATTwww++++++++** \| \|; coexpression\_cluster1\_m36\_cluster1\_motif36\_m \| cluster1\_motif36\_m \| cluster\_9 \| coexpression\_cluster1 \| 10 \| 11.11 \| 227 \| **TGAACAAATT** \| **AATTTGTTCA** \| \|; coexpression\_cluster1\_m26\_cluster1\_motif26\_m \| cluster1\_motif26\_m \| cluster\_4 \| coexpression\_cluster1 \| 15 \| 13.08 \| 1000 \| **TATATATATryATry** \| **RYATRYATATATATA** \| \|; coexpression\_cluster1\_m5\_cluster1\_motif5\_r \| cluster1\_motif5\_r \| cluster\_2 \| coexpression\_cluster1 \| 12 \| 8.10 \| 836 \| **WWGTGTTGTAWW** \| **wwtACAACAcww** \| \|; coexpression\_cluster1\_m21\_cluster1\_motif21\_r \| cluster1\_motif21\_r \| cluster\_1 \| coexpression\_cluster1 \| 15 \| 9.29 \| 4491 \| **wwATrAAcAAAwww+** \| **+WWWTTTGTTYATWW** \| \|; coexpression\_cluster1\_m33\_cluster1\_motif33\_m \| cluster1\_motif33\_m \| cluster\_1 \| coexpression\_cluster1 \| 15 \| 13.42 \| 300 \| **+TATATATTTTTT++** \| **++AAAAAATATATA+** \| \|; coexpression\_cluster1\_m29\_cluster1\_motif29\_m \| cluster1\_motif29\_m \| cluster\_5 \| coexpression\_cluster1 \| 22 \| 11.34 \| 82 \| **+++++++AGCTAGCYAwwwwt+** \| **+AWWWWTRGCTAGCT+++++++** \| \|; coexpression\_cluster1\_m19\_cluster1\_motif19\_r \| cluster1\_motif19\_r \| cluster\_3 \| coexpression\_cluster1 \| 24 \| 12.38 \| 7463 \| **WWWTWTWWWWWTTTTTKTYTTTWW** \| **wwAAArAmAAAAAwwwwwAwAwww** \| \|; coexpression\_cluster1\_m12\_cluster1\_motif12\_r \| cluster1\_motif12\_r \| cluster\_1 \| coexpression\_cluster1 \| 15 \| 9.70 \| 2441 \| **WWWTKTTKTYTTTWW** \| **wwAAArAmAAmAwww** \| \|; coexpression\_cluster1\_m37\_cluster1\_motif37\_m \| cluster1\_motif37\_m \| cluster\_7 \| coexpression\_cluster1 \| 14 \| 15.20 \| 85 \| **TAATAATAATAAwA** \| **TWTTATTATTATTA** \| \|; coexpression\_cluster1\_m43\_cluster1\_motif43\_m \| cluster1\_motif43\_m \| cluster\_4 \| coexpression\_cluster1 \| 15 \| 12.92 \| 203 \| **++TATATATGCATGY** \| **rCATGCATATATA++** \| \|; coexpression\_cluster1\_m25\_cluster1\_motif25\_r \| cluster1\_motif25\_r \| cluster\_1 \| coexpression\_cluster1 \| 15 \| 7.85 \| 4724 \| **wwAAAgACAwww+++** \| **+++WWWTGTCTTTWW** \| \|; coexpression\_cluster1\_m11\_cluster1\_motif11\_r \| cluster1\_motif11\_r \| cluster\_1 \| coexpression\_cluster1 \| 15 \| 8.50 \| 1748 \| **WWWTAGACAAWWWW+** \| **+wwwwTTGTcTAwww** \| \|; coexpression\_cluster1\_m4\_cluster1\_motif4\_r \| cluster1\_motif4\_r \| cluster\_4 \| coexpression\_cluster1 \| 15 \| 8.93 \| 1058 \| **++WATGCATGCATW+** \| **+waTGCATGCAtw++** \| \|; coexpression\_cluster1\_m39\_cluster1\_motif39\_m \| cluster1\_motif39\_m \| cluster\_10 \| coexpression\_cluster1 \| 22 \| 10.85 \| 148 \| **wtswwttTGCACACACrhmvrg** \| **CYBKDYGTGTGTGCAAAWWSAW** \| \|; coexpression\_cluster1\_m23\_cluster1\_motif23\_r \| cluster1\_motif23\_r \| cluster\_2 \| coexpression\_cluster1 \| 12 \| 8.21 \| 1798 \| **wwATGTTGTAww** \| **WWTACAACATWW** \| \|; coexpression\_cluster1\_m30\_cluster1\_motif30\_m \| cluster1\_motif30\_m \| cluster\_4 \| coexpression\_cluster1 \| 15 \| 8.24 \| 544 \| **+++RYAYATRCAYA+** \| **+TrTGyATrTry+++** \| \|; coexpression\_cluster1\_m3\_cluster1\_motif3\_r \| cluster1\_motif3\_r \| cluster\_1 \| coexpression\_cluster1 \| 15 \| 8.87 \| 1735 \| **WWWATGTCTTTWW++** \| **++wwAAAgACATwww** \| \|; coexpression\_cluster1\_m42\_cluster1\_motif42\_m \| cluster1\_motif42\_m \| cluster\_13 \| coexpression\_cluster1 \| 15 \| 9.94 \| 25 \| **msvwmCCCCMmmTAt** \| **ATAKKKGGGGKWBSK** \| \|; coexpression\_cluster1\_m41\_cluster1\_motif41\_m \| cluster1\_motif41\_m \| cluster\_1 \| coexpression\_cluster1 \| 15 \| 14.58 \| 549 \| **+TTAWTTTWTYTT++** \| **++AARAwAAAwTAA+** \| \|; coexpression\_cluster1\_m2\_cluster1\_motif2\_r \| cluster1\_motif2\_r \| cluster\_1 \| coexpression\_cluster1 \| 15 \| 8.90 \| 1983 \| **wwATAgACAAwwww+** \| **+WWWWTTGTCTATWW** \| \|; coexpression\_cluster1\_m1\_cluster1\_motif1\_r \| cluster1\_motif1\_r \| cluster\_1 \| coexpression\_cluster1 \| 15 \| 9.10 \| 2614 \| **WWWWTTGTCTTTWW+** \| **+wwAAAgACAAwwww** \| \|; coexpression\_cluster1\_m40\_cluster1\_motif40\_m \| cluster1\_motif40\_m \| cluster\_1 \| coexpression\_cluster1 \| 15 \| 14.58 \| 198 \| **AAATAAATaAAATAA** \| **TTATTTTATTTATTT** \| \|; coexpression\_cluster1\_m35\_cluster1\_motif35\_m \| cluster1\_motif35\_m \| cluster\_11 \| coexpression\_cluster1 \| 6 \| 6.26 \| 168 \| **GCATGC** \| **GCATGC** \| \|; coexpression\_cluster1\_m16\_cluster1\_motif16\_r \| cluster1\_motif16\_r \| cluster\_3 \| coexpression\_cluster1 \| 24 \| 9.39 \| 5262 \| **+++++++++WWYATTTTTAAATWW** \| **wwATTTAAAAATrww+++++++++** \| \|; coexpression\_cluster1\_m10\_cluster1\_motif10\_r \| cluster1\_motif10\_r \| cluster\_4 \| coexpression\_cluster1 \| 15 \| 8.75 \| 1740 \| **+++atACACACATww** \| **WWATGTGTGTAT+++** \| \|; coexpression\_cluster1\_m20\_cluster1\_motif20\_r \| cluster1\_motif20\_r \| cluster\_2 \| coexpression\_cluster1 \| 12 \| 9.05 \| 201 \| **++WWTGCAACWW** \| **wwGTTGCAww++** \| \|; coexpression\_cluster1\_m13\_cluster1\_motif13\_r \| cluster1\_motif13\_r \| cluster\_8 \| coexpression\_cluster1 \| 10 \| 9.23 \| 216 \| **wwTCTGAAww** \| **WWTTCAGAWW** \| \|; coexpression\_cluster1\_m6\_cluster1\_motif6\_r \| cluster1\_motif6\_r \| cluster\_3 \| coexpression\_cluster1 \| 24 \| 13.05 \| 3274 \| **wwATrAAmAAAAATwTtTAAATww** \| **WWATTTAAAWATTTTTKTTYATWW** \| \|; coexpression\_cluster1\_m22\_cluster1\_motif22\_r \| cluster1\_motif22\_r \| cluster\_3 \| coexpression\_cluster1 \| 24 \| 13.12 \| 2727 \| **wwTAmAAmAAAAATwTtTAAAtww** \| **WWATTTAAAWATTTTTKTTKTAWW** \| \|; coexpression\_cluster1\_m9\_cluster1\_motif9\_r \| cluster1\_motif9\_r \| cluster\_1 \| coexpression\_cluster1 \| 15 \| 8.35 \| 943 \| **wwAAAGACAcww+++** \| **+++WWGTGTCTTTWW** \| \|; coexpression\_cluster1\_m34\_cluster1\_motif34\_m \| cluster1\_motif34\_m \| cluster\_1 \| coexpression\_cluster1 \| 15 \| 14.58 \| 264 \| **+AAAAAAAAAAAAAC** \| **GTTTTTTTTTTTTT+** \| \|; coexpression\_cluster1\_m7\_cluster1\_motif7\_r \| cluster1\_motif7\_r \| cluster\_1 \| coexpression\_cluster1 \| 15 \| 8.62 \| 2153 \| **WWWAAGACAAWWWW+** \| **+wwwwTTGTcTTwww** \| \|; coexpression\_cluster1\_m44\_cluster1\_motif44\_m \| cluster1\_motif44\_m \| cluster\_12 \| coexpression\_cluster1 \| 15 \| 20.79 \| 6 \| **CATTTCATGACCCAT** \| **ATGGGTCATGAAATG** \| \|; coexpression\_cluster1\_m24\_cluster1\_motif24\_r \| cluster1\_motif24\_r \| cluster\_2 \| coexpression\_cluster1 \| 12 \| 9.22 \| 2255 \| **+WWTGTTTTWW+** \| **+wwAAAACAww+** \| \|; coexpression\_cluster1\_m27\_cluster1\_motif27\_m \| cluster1\_motif27\_m \| cluster\_4 \| coexpression\_cluster1 \| 15 \| 15.49 \| 269 \| **TATATrTAyATATA+** \| **+TATATRTAYATATA** \| \| - Click on the column names to change the order of the data. - Write the name of one cluster, collection or a pattern in the *Search* window. **Heatmap View** Heatmap View Distance table \| PDF |

**Additional Files**

#### Exported files

| File | Description |
| --- | --- |
| **Pairwise comparison** | This table shows the pairwise comparison between all the input motifs using different metrics. This is the *compare-matrices* result. |
| **Matrix description** | This table shows information of each input motif. |
| **Cluster-wise Motif Files [zip]** | One TRANSFAC file is generated for each cluster including all the input motifs grouped in the corresponding cluster. |
| **Root motifs** | A file with the root matrix of each cluster. |
| **Motif collections per cluster** | A tab separated file containing the counts of motif of each collection on each cluster. |
| **Clusters** | A tab separated file containing the clusters and their correspondng motifs. |
| **Central motifs IDs** | A tab separated file containing the central motif, that one who is the most similar to all the others motif in the same cluster. |
| **Central motifs (in TRANSFAC format)** | A transfac file containing the central motif, that one who is the most similar to all the others motif within the same cluster. |
| **Input motifs** | The input motifs analyzed with matrix-clustering. |

- **References**
  1. Castro-Mondragon JA et al.  RSAT matrix-clustering: dynamic exploration and redundancy reduction of transcription factor binding motif collections. doi: 10.1093/nar/gkx314; Nucleic Acids Research (2017)
  2. **Last RSAT release:** Nguyen NTT et al. RSAT 2018: regulatory sequence analysis tools 20th anniversary. Nucleic Acids Research (2018).
- **Additional Information**
  - Jacques van Helden course
- **Contact**
  - Jaime Castro-Mondragon - ****
  - Morgane Thomas-Chollier - ****
  - Jacques van Helden - ****
