## Additional file 9 for "Transcriptome analysis reveals a novel DNA element that may interact with chromatin-associated proteins in *Plasmodium berghei* during erythrocytic development": matrix-clustering_SUMMARY_cluster2.html

matrix-clustering coexpression\_cluster2

### RSAT - matrix-clustering result

##### Analysis: coexpression\_cluster2 (09/09/2023 18:16)

##### Command

```
matrix-clustering  -v 1 -max_matrices 300 -matrix coexpression_cluster2 $RSAT/public_html/tmp/www-data/2023/09/09/matrix-clustering_2023-09-09.181618_SMplQh/matrix-clustering_query_matrices.transfac transfac -hclust_method average -calc sum -title coexpression_cluster2 -metric_build_tree Ncor -lth w 5 -lth cor 0.6 -lth Ncor 0.4 -quick -label_in_tree name -return json,heatmap -o $RSAT/public_html/tmp/www-data/2023/09/09/matrix-clustering_2023-09-09.181618_SMplQh/matrix-clustering
```

#### Results Summary

|  |  |  |  |  |  |
| --- | --- | --- | --- | --- | --- |
| Nb Input motifs | Nb Input collections | Nb Clusters Found | Download   Root motifs | Complete results [zip] | Parameters used Linkage method \| Similarity metric \| Thresholds to partition the tree |
| 32 | 1 | 9 | **Download** | Download | average \| Ncor \| **Ncor** = 0.4 **cor** = 0.6 |

|  |  |  |  |  |  |  |
| --- | --- | --- | --- | --- | --- | --- |
|  | **Logo Forest (dynamic browsing)**  **Logo Forest (rapid overview - low image quality)**  cluster\_1  cluster\_2  cluster\_3  cluster\_4  cluster\_5  cluster\_6  cluster\_7  cluster\_8  cluster\_9 |  |  |  |  |  |
|  | **Clusters Summary** Clusters Summary Table (Root Motif \| Root Motif (Reverse) \| Cluster ID \| # Motifs \| Motif\_Name::Collection \| Number of Motifs by Collection \| Root Motif (transfac Format)) \| | cluster\_1 | 8 | cluster2\_motif4\_r::coexpression\_cluster2:: cluster2\_motif8\_r::coexpression\_cluster2:: cluster2\_motif17\_r::coexpression\_cluster2:: cluster2\_motif5\_r::coexpression\_cluster2:: cluster2\_motif10\_r::coexpression\_cluster2:: cluster2\_motif2\_r::coexpression\_cluster2:: cluster2\_motif18\_r::coexpression\_cluster2:: cluster2\_motif1\_r::coexpression\_cluster2:: | coexpression\_cluster2::8\_motifs | Show matrix |
| cluster\_2 \| 3 \| cluster2\_motif24\_m::coexpression\_cluster2:: cluster2\_motif20\_m::coexpression\_cluster2:: cluster2\_motif26\_m::coexpression\_cluster2:: \| coexpression\_cluster2::3\_motifs \| Show matrix \| \| |  |  |  |  |  |  |
| cluster\_3 \| 9 \| cluster2\_motif12\_r::coexpression\_cluster2:: cluster2\_motif16\_r::coexpression\_cluster2:: cluster2\_motif3\_r::coexpression\_cluster2:: cluster2\_motif6\_r::coexpression\_cluster2:: cluster2\_motif19\_r::coexpression\_cluster2:: cluster2\_motif9\_r::coexpression\_cluster2:: cluster2\_motif27\_m::coexpression\_cluster2:: cluster2\_motif14\_r::coexpression\_cluster2:: cluster2\_motif7\_r::coexpression\_cluster2:: \| coexpression\_cluster2::9\_motifs \| Show matrix \| \| |  |  |  |  |  |  |
| cluster\_4 | 7 | cluster2\_motif30\_m::coexpression\_cluster2:: cluster2\_motif29\_m::coexpression\_cluster2:: cluster2\_motif21\_m::coexpression\_cluster2:: cluster2\_motif22\_m::coexpression\_cluster2:: cluster2\_motif25\_m::coexpression\_cluster2:: cluster2\_motif23\_m::coexpression\_cluster2:: cluster2\_motif28\_m::coexpression\_cluster2:: | coexpression\_cluster2::7\_motifs | Show matrix | | |
| cluster\_5 | 1 | cluster2\_motif15\_r::coexpression\_cluster2 | coexpression\_cluster2 : 1 motif | Show matrix | | |
| cluster\_6 | 1 | cluster2\_motif32\_m::coexpression\_cluster2 | coexpression\_cluster2 : 1 motif | Show matrix | | |
| cluster\_7 | 1 | cluster2\_motif11\_r::coexpression\_cluster2 | coexpression\_cluster2 : 1 motif | Show matrix | | |
| cluster\_8 | 1 | cluster2\_motif33\_m::coexpression\_cluster2 | coexpression\_cluster2 : 1 motif | Show matrix | | |
| cluster\_9 | 1 | cluster2\_motif31\_m::coexpression\_cluster2 | coexpression\_cluster2 : 1 motif | Show matrix | | |

- Note: by default the table displays the first 50 clusters. Click on the *Show X entries* button to display more (or less) clusters.
- Click on the column names to change the order of the data.
- Use the *Search* window to find your motif (s) of interest.

| **Individual Cluster View** Individual Cluster View **Show All** **Hide All** **cluster\_1** **cluster\_2** **cluster\_3** **cluster\_4** **cluster\_5** **cluster\_6** **cluster\_7** **cluster\_8** **cluster\_9** **Display Logo Trees** **cluster\_1** **cluster\_2** **cluster\_3** **cluster\_4** **cluster\_5** **cluster\_6** **cluster\_7** **cluster\_8** **cluster\_9** **Display Branch-Motifs** **cluster\_1** Branch Motifs \| \| \| \| \| \| \|; Node \| Consensus \| Logo \| Logo (Reverse) \| Collections \| Matrix (Transfac format) \| IC \| Nb sites; Node 1 \| **---------wwwATATTATATAATATAww----** \| \| \| coexpression\_cluster2::2\_motifs \| PSSM \| 11.23 \| 16744; Node 2 \| **wwwtwTATAATATTATTATATtwwwwwwww---** \| \| \| coexpression\_cluster2::2\_motifs \| PSSM \| 15.14 \| 7399; Node 3 \| **-------wwwwwwTATAATATAATww-------** \| \| \| coexpression\_cluster2::2\_motifs \| PSSM \| 10.36 \| 16758; Node 4 \| **------wwTwwwwTATTATATAATATAww----** \| \| \| coexpression\_cluster2::3\_motifs \| PSSM \| 12.04 \| 22358; Node 5 \| **------wwTwwwwTATTATATAATwwAwwTAww** \| \| \| coexpression\_cluster2::4\_motifs \| PSSM \| 13.32 \| 30226; Node 6 \| **------wwwwwwwTATwATATAATwwAwwTAww** \| \| \| coexpression\_cluster2::6\_motifs \| PSSM \| 12.86 \| 46984; Node 7 \| **wwwtwTwwwwwwwTATwATATwATwwwwwwAww** \| \| \| coexpression\_cluster2::8\_motifs \| PSSM \| 15.12 \| 54383 **cluster\_2** Branch Motifs \| \| \| \| \| \| \|; Node \| Consensus \| Logo \| Logo (Reverse) \| Collections \| Matrix (Transfac format) \| IC \| Nb sites; Node 1 \| **ATAtATAyATAyATA** \| \| \| coexpression\_cluster2::2\_motifs \| PSSM \| 13.56 \| 1235; Node 2 \| **ATATATAyATAyATA** \| \| \| coexpression\_cluster2::3\_motifs \| PSSM \| 13.08 \| 1378 **cluster\_3** Branch Motifs \| \| \| \| \| \| \|; Node \| Consensus \| Logo \| Logo (Reverse) \| Collections \| Matrix (Transfac format) \| IC \| Nb sites; Node 1 \| **wwTTTTAdAAwAww** \| \| \| coexpression\_cluster2::2\_motifs \| PSSM \| 8.65 \| 13623; Node 2 \| **wwTTTTAkAwwww-** \| \| \| coexpression\_cluster2::2\_motifs \| PSSM \| 8.37 \| 10832; Node 3 \| **wwTATtmTATww--** \| \| \| coexpression\_cluster2::2\_motifs \| PSSM \| 8.02 \| 6913; Node 4 \| **wwTTTTAdAwwaww** \| \| \| coexpression\_cluster2::4\_motifs \| PSSM \| 8.19 \| 24455; Node 5 \| **-wwATAmTAAww--** \| \| \| coexpression\_cluster2::2\_motifs \| PSSM \| 7.41 \| 8501; Node 6 \| **wwTATtyTATww--** \| \| \| coexpression\_cluster2::3\_motifs \| PSSM \| 7.96 \| 7327; Node 7 \| **wwTWTTAkAwwwww** \| \| \| coexpression\_cluster2::7\_motifs \| PSSM \| 7.35 \| 31782; Node 8 \| **wwTwTwakAwwwww** \| \| \| coexpression\_cluster2::9\_motifs \| PSSM \| 6.73 \| 40283 **cluster\_4** Branch Motifs \| \| \| \| \| \| \|; Node \| Consensus \| Logo \| Logo (Reverse) \| Collections \| Matrix (Transfac format) \| IC \| Nb sites; Node 1 \| **-rAAAAAArAGrrAAA-** \| \| \| coexpression\_cluster2::2\_motifs \| PSSM \| 11.91 \| 700; Node 2 \| **-rAAAAAaraGraAAAA** \| \| \| coexpression\_cluster2::3\_motifs \| PSSM \| 11.80 \| 933; Node 3 \| **-AAATwAAwAAAwAAA-** \| \| \| coexpression\_cluster2::2\_motifs \| PSSM \| 12.45 \| 850; Node 4 \| **AAAATwAAwAAwwAAA-** \| \| \| coexpression\_cluster2::3\_motifs \| PSSM \| 13.09 \| 1048; Node 5 \| **AAAAwwAAwarawAAAA** \| \| \| coexpression\_cluster2::6\_motifs \| PSSM \| 11.79 \| 1980; Node 6 \| **AAAAwwAAwarawAAAA** \| \| \| coexpression\_cluster2::7\_motifs \| PSSM \| 11.27 \| 2026 **cluster\_5** Branch Motifs \| \| \| \| \| \| \|; Node \| Consensus \| Logo \| Logo (Reverse) \| Collections \| Matrix (Transfac format) \| IC \| Nb sites; Singleton \| **wwAGGTAAww** \| \| \| coexpression\_cluster2 : 1 motif \| PSSM \| 9.25 \| 270 **cluster\_6** Branch Motifs \| \| \| \| \| \| \|; Node \| Consensus \| Logo \| Logo (Reverse) \| Collections \| Matrix (Transfac format) \| IC \| Nb sites; Singleton \| **CaStGCtaattGCYG** \| \| \| coexpression\_cluster2 : 1 motif \| PSSM \| 15.09 \| 18 **cluster\_7** Branch Motifs \| \| \| \| \| \| \|; Node \| Consensus \| Logo \| Logo (Reverse) \| Collections \| Matrix (Transfac format) \| IC \| Nb sites; Singleton \| **wwGCACTAww** \| \| \| coexpression\_cluster2 : 1 motif \| PSSM \| 9.06 \| 123 **cluster\_8** Branch Motifs \| \| \| \| \| \| \|; Node \| Consensus \| Logo \| Logo (Reverse) \| Collections \| Matrix (Transfac format) \| IC \| Nb sites; Singleton \| **tSGRaaaGrtGtGCa** \| \| \| coexpression\_cluster2 : 1 motif \| PSSM \| 16.85 \| 14 **cluster\_9** Branch Motifs \| \| \| \| \| \| \|; Node \| Consensus \| Logo \| Logo (Reverse) \| Collections \| Matrix (Transfac format) \| IC \| Nb sites; Singleton \| **KyktKGGGGSG** \| \| \| coexpression\_cluster2 : 1 motif \| PSSM \| 9.09 \| 71 - **Each cluster has a different color.** - **Click on the upper buttons to display separately the information of each cluster.** - **Click on the circles at each tree to display its corresponding merged motifs.** Definitions - **Logo tree.** This results shows the hierarchical tree with its logo alignment of one cluster. The logos are shown in Forward (left logo) and Reverse (right logo) orientation. - **Branch-motifs.** On the trees are displayed the branch number that is the number in which the motifs were incorporated in the tree. The Branch-Motifs table shows the logo in both orientations and the link to the file in TRANSFAC format with the branch-motif. |
|  | **Individual Motif View** Individual Motif View Motif id \| Motif name \| Cluster \| Collection \| Width \| IC \| Number of sites \| Consensus \| Consensus (Rev) \| Logo \| Logo (Rev); coexpression\_cluster2\_m19\_cluster2\_motif20\_m \| cluster2\_motif20\_m \| cluster\_2 \| coexpression\_cluster2 \| 15 \| 13.12 \| 942 \| **ATATATAYATRYATA** \| **TATryATrTATaTAT** \| \|; coexpression\_cluster2\_m4\_cluster2\_motif4\_r \| cluster2\_motif4\_r \| cluster\_1 \| coexpression\_cluster2 \| 33 \| 15.31 \| 4244 \| **wwwtwTATAATATTATTATATtwwwwwwww+++** \| **+++WWWWWWWWAATATAATAATATTATAWAWWW** \| \|; coexpression\_cluster2\_m22\_cluster2\_motif23\_m \| cluster2\_motif23\_m \| cluster\_4 \| coexpression\_cluster2 \| 17 \| 13.03 \| 481 \| **+AAATAAATAAATAAA+** \| **+TTTATTTATTTATTT+** \| \|; coexpression\_cluster2\_m21\_cluster2\_motif22\_m \| cluster2\_motif22\_m \| cluster\_4 \| coexpression\_cluster2 \| 17 \| 11.52 \| 399 \| **+++AAAAAraGrrAAA+** \| **+TTTYYCTYTTTTT+++** \| \|; coexpression\_cluster2\_m2\_cluster2\_motif2\_r \| cluster2\_motif2\_r \| cluster\_1 \| coexpression\_cluster2 \| 33 \| 12.97 \| 5614 \| **++++++WWTATATTWTTATATAATATAWW++++** \| **++++wwtATATTATATAAwAATATAww++++++** \| \|; coexpression\_cluster2\_m1\_cluster2\_motif1\_r \| cluster2\_motif1\_r \| cluster\_1 \| coexpression\_cluster2 \| 33 \| 11.53 \| 7590 \| **+++++++++WWTATATTATATAATATAWW++++** \| **++++wwTATATTATATAATATAww+++++++++** \| \|; coexpression\_cluster2\_m17\_cluster2\_motif18\_r \| cluster2\_motif18\_r \| cluster\_1 \| coexpression\_cluster2 \| 33 \| 11.07 \| 9154 \| **+++++++++WWWATATTATATAATATAWW++++** \| **++++wwtaTATTATATAATAtwww+++++++++** \| \|; coexpression\_cluster2\_m15\_cluster2\_motif16\_r \| cluster2\_motif16\_r \| cluster\_3 \| coexpression\_cluster2 \| 14 \| 9.43 \| 893 \| **++wwTAAGAAww++** \| **++WWTTCTTAWW++** \| \|; coexpression\_cluster2\_m8\_cluster2\_motif8\_r \| cluster2\_motif8\_r \| cluster\_1 \| coexpression\_cluster2 \| 33 \| 15.48 \| 3155 \| **WWWTATATAATATTATTATATWAWWAWWWW+++** \| **+++wwwwtwwtwATATAATAATATTATAtAwww** \| \|; coexpression\_cluster2\_m20\_cluster2\_motif21\_m \| cluster2\_motif21\_m \| cluster\_4 \| coexpression\_cluster2 \| 17 \| 12.22 \| 301 \| **+rAAAAAwrArarAAA+** \| **+TTTYTYTYWTTTTTY+** \| \|; coexpression\_cluster2\_m7\_cluster2\_motif7\_r \| cluster2\_motif7\_r \| cluster\_3 \| coexpression\_cluster2 \| 14 \| 9.25 \| 4484 \| **WWTATTHTATWW++** \| **++wwATAdAATAww** \| \|; coexpression\_cluster2\_m14\_cluster2\_motif15\_r \| cluster2\_motif15\_r \| cluster\_5 \| coexpression\_cluster2 \| 10 \| 9.25 \| 270 \| **wwAGGTAAww** \| **WWTTACCTWW** \| \|; coexpression\_cluster2\_m30\_cluster2\_motif31\_m \| cluster2\_motif31\_m \| cluster\_9 \| coexpression\_cluster2 \| 11 \| 9.09 \| 71 \| **KTtTkGrGGkG** \| **CMCCYCMAAAM** \| \|; coexpression\_cluster2\_m13\_cluster2\_motif14\_r \| cluster2\_motif14\_r \| cluster\_3 \| coexpression\_cluster2 \| 14 \| 7.33 \| 2429 \| **WWTATCCTAWW+++** \| **+++wwTAGgATAww** \| \|; coexpression\_cluster2\_m5\_cluster2\_motif5\_r \| cluster2\_motif5\_r \| cluster\_1 \| coexpression\_cluster2 \| 33 \| 10.55 \| 8507 \| **+++++++wwAwAwTATAATATAwTww+++++++** \| **+++++++WWAWTATATTATAWTWTWW+++++++** \| \|; coexpression\_cluster2\_m6\_cluster2\_motif6\_r \| cluster2\_motif6\_r \| cluster\_3 \| coexpression\_cluster2 \| 14 \| 8.95 \| 7557 \| **wwTTTTAdAAAAww** \| **WWTTTTHTAAAAWW** \| \|; coexpression\_cluster2\_m9\_cluster2\_motif9\_r \| cluster2\_motif9\_r \| cluster\_3 \| coexpression\_cluster2 \| 14 \| 9.28 \| 3289 \| **WWTTTTAKATWWW+** \| **+wwwATmTAAAAww** \| \|; coexpression\_cluster2\_m32\_cluster2\_motif33\_m \| cluster2\_motif33\_m \| cluster\_8 \| coexpression\_cluster2 \| 15 \| 16.85 \| 14 \| **TGGrAAAGATGTGCA** \| **TGCACATCTTTYCCA** \| \|; coexpression\_cluster2\_m26\_cluster2\_motif27\_m \| cluster2\_motif27\_m \| cluster\_3 \| coexpression\_cluster2 \| 14 \| 12.35 \| 414 \| **TATATTTTTTT+++** \| **+++AAAAAAATATA** \| \|; coexpression\_cluster2\_m27\_cluster2\_motif28\_m \| cluster2\_motif28\_m \| cluster\_4 \| coexpression\_cluster2 \| 17 \| 14.00 \| 369 \| **+AAATTAAAAAAAAA++** \| **++TTTTTTTTTAATTT+** \| \|; coexpression\_cluster2\_m3\_cluster2\_motif3\_r \| cluster2\_motif3\_r \| cluster\_3 \| coexpression\_cluster2 \| 14 \| 9.07 \| 6066 \| **wwTTTTAdAATAww** \| **WWTATTHTAAAAWW** \| \|; coexpression\_cluster2\_m31\_cluster2\_motif32\_m \| cluster2\_motif32\_m \| cluster\_6 \| coexpression\_cluster2 \| 15 \| 15.09 \| 18 \| **yAsTGCTAATTGmyG** \| **CRKCAATTAGCASTR** \| \|; coexpression\_cluster2\_m12\_cluster2\_motif12\_r \| cluster2\_motif12\_r \| cluster\_3 \| coexpression\_cluster2 \| 14 \| 7.70 \| 7608 \| **+wwATAmTAAww++** \| **++WWTTAKTATWW+** \| \|; coexpression\_cluster2\_m25\_cluster2\_motif26\_m \| cluster2\_motif26\_m \| cluster\_2 \| coexpression\_cluster2 \| 15 \| 15.81 \| 293 \| **ATATATATATAYAT+** \| **+ATRTATATATATAT** \| \|; coexpression\_cluster2\_m16\_cluster2\_motif17\_r \| cluster2\_motif17\_r \| cluster\_1 \| coexpression\_cluster2 \| 33 \| 9.99 \| 8251 \| **+++++++++wwwwTATAATATAATww+++++++** \| **+++++++WWATTATATTATAWWWW+++++++++** \| \|; coexpression\_cluster2\_m23\_cluster2\_motif24\_m \| cluster2\_motif24\_m \| cluster\_2 \| coexpression\_cluster2 \| 15 \| 12.48 \| 143 \| **ATATATGCATATAT+** \| **+ATATATGCATATAT** \| \|; coexpression\_cluster2\_m29\_cluster2\_motif30\_m \| cluster2\_motif30\_m \| cluster\_4 \| coexpression\_cluster2 \| 17 \| 14.13 \| 46 \| **++RTTWGTKTRYTTKTT** \| **AAMAArYAmACwAAy++** \| \|; coexpression\_cluster2\_m24\_cluster2\_motif25\_m \| cluster2\_motif25\_m \| cluster\_4 \| coexpression\_cluster2 \| 17 \| 15.59 \| 199 \| **TTTATTATTATTWTT++** \| **++AAwAATAATAATAAA** \| \|; coexpression\_cluster2\_m28\_cluster2\_motif29\_m \| cluster2\_motif29\_m \| cluster\_4 \| coexpression\_cluster2 \| 17 \| 11.79 \| 233 \| **++TTTTTYCMYYTTTTT** \| **AAAAArrkGrAAAAA++** \| \|; coexpression\_cluster2\_m10\_cluster2\_motif10\_r \| cluster2\_motif10\_r \| cluster\_1 \| coexpression\_cluster2 \| 33 \| 12.66 \| 7868 \| **+++++++++wwTATATTATATAwwwwATATAww** \| **WWTATATWWWWTATATAATATAWW+++++++++** \| \|; coexpression\_cluster2\_m11\_cluster2\_motif11\_r \| cluster2\_motif11\_r \| cluster\_7 \| coexpression\_cluster2 \| 10 \| 9.06 \| 123 \| **wwGCACTAww** \| **WWTAGTGCWW** \| \|; coexpression\_cluster2\_m18\_cluster2\_motif19\_r \| cluster2\_motif19\_r \| cluster\_3 \| coexpression\_cluster2 \| 14 \| 7.96 \| 7543 \| **WWTTTTAKAWWW++** \| **++wwwTmTAAAAww** \| \| - Click on the column names to change the order of the data. - Write the name of one cluster, collection or a pattern in the *Search* window. **Heatmap View** Heatmap View Distance table \| PDF |

**Additional Files**

#### Exported files

| File | Description |
| --- | --- |
| **Pairwise comparison** | This table shows the pairwise comparison between all the input motifs using different metrics. This is the *compare-matrices* result. |
| **Matrix description** | This table shows information of each input motif. |
| **Cluster-wise Motif Files [zip]** | One TRANSFAC file is generated for each cluster including all the input motifs grouped in the corresponding cluster. |
| **Root motifs** | A file with the root matrix of each cluster. |
| **Motif collections per cluster** | A tab separated file containing the counts of motif of each collection on each cluster. |
| **Clusters** | A tab separated file containing the clusters and their correspondng motifs. |
| **Central motifs IDs** | A tab separated file containing the central motif, that one who is the most similar to all the others motif in the same cluster. |
| **Central motifs (in TRANSFAC format)** | A transfac file containing the central motif, that one who is the most similar to all the others motif within the same cluster. |
| **Input motifs** | The input motifs analyzed with matrix-clustering. |

- **References**
  1. Castro-Mondragon JA et al.  RSAT matrix-clustering: dynamic exploration and redundancy reduction of transcription factor binding motif collections. doi: 10.1093/nar/gkx314; Nucleic Acids Research (2017)
  2. **Last RSAT release:** Nguyen NTT et al. RSAT 2018: regulatory sequence analysis tools 20th anniversary. Nucleic Acids Research (2018).
- **Additional Information**
  - Jacques van Helden course
- **Contact**
  - Jaime Castro-Mondragon - ****
  - Morgane Thomas-Chollier - ****
  - Jacques van Helden - ****
