## Additional file 9 for "Transcriptome analysis reveals a novel DNA element that may interact with chromatin-associated proteins in *Plasmodium berghei* during erythrocytic development": matrix-clustering_SUMMARY_cluster3.html

matrix-clustering coexpression\_cluster3

### RSAT - matrix-clustering result

##### Analysis: coexpression\_cluster3 (08/09/2023 12:48)

##### Command

```
matrix-clustering  -v 1 -max_matrices 300 -matrix coexpression_cluster3 $RSAT/public_html/tmp/www-data/2023/09/08/matrix-clustering_2023-09-08.124743_XczLfy/matrix-clustering_query_matrices.transfac transfac -hclust_method average -calc sum -title coexpression_cluster3 -metric_build_tree Ncor -lth w 5 -lth cor 0.6 -lth Ncor 0.4 -quick -label_in_tree name -return json,heatmap -o $RSAT/public_html/tmp/www-data/2023/09/08/matrix-clustering_2023-09-08.124743_XczLfy/matrix-clustering
```

#### Results Summary

|  |  |  |  |  |  |
| --- | --- | --- | --- | --- | --- |
| Nb Input motifs | Nb Input collections | Nb Clusters Found | Download   Root motifs | Complete results [zip] | Parameters used Linkage method \| Similarity metric \| Thresholds to partition the tree |
| 44 | 1 | 9 | **Download** | Download | average \| Ncor \| **Ncor** = 0.4 **cor** = 0.6 |

|  |  |  |  |  |  |  |
| --- | --- | --- | --- | --- | --- | --- |
|  | **Logo Forest (dynamic browsing)**  **Logo Forest (rapid overview - low image quality)**  cluster\_1  cluster\_2  cluster\_3  cluster\_4  cluster\_5  cluster\_6  cluster\_7  cluster\_8  cluster\_9 |  |  |  |  |  |
|  | **Clusters Summary** Clusters Summary Table (Root Motif \| Root Motif (Reverse) \| Cluster ID \| # Motifs \| Motif\_Name::Collection \| Number of Motifs by Collection \| Root Motif (transfac Format)) \| | cluster\_1 | 18 | cluster3\_motif19\_r::coexpression\_cluster3:: cluster3\_motif36\_m::coexpression\_cluster3:: cluster3\_motif8\_r::coexpression\_cluster3:: cluster3\_motif3\_r::coexpression\_cluster3:: cluster3\_motif18\_r::coexpression\_cluster3:: cluster3\_motif5\_r::coexpression\_cluster3:: cluster3\_motif21\_r::coexpression\_cluster3:: cluster3\_motif15\_r::coexpression\_cluster3:: cluster3\_motif10\_r::coexpression\_cluster3:: cluster3\_motif4\_r::coexpression\_cluster3:: cluster3\_motif14\_r::coexpression\_cluster3:: cluster3\_motif13\_r::coexpression\_cluster3:: cluster3\_motif1\_r::coexpression\_cluster3:: cluster3\_motif7\_r::coexpression\_cluster3:: cluster3\_motif9\_r::coexpression\_cluster3:: cluster3\_motif16\_r::coexpression\_cluster3:: cluster3\_motif22\_m::coexpression\_cluster3:: cluster3\_motif23\_m::coexpression\_cluster3:: | coexpression\_cluster3::18\_motifs | Show matrix |
| cluster\_2 \| 4 \| cluster3\_motif35\_m::coexpression\_cluster3:: cluster3\_motif44\_m::coexpression\_cluster3:: cluster3\_motif40\_m::coexpression\_cluster3:: cluster3\_motif42\_m::coexpression\_cluster3:: \| coexpression\_cluster3::4\_motifs \| Show matrix \| \| |  |  |  |  |  |  |
| cluster\_3 \| 2 \| cluster3\_motif33\_m::coexpression\_cluster3:: cluster3\_motif34\_m::coexpression\_cluster3:: \| coexpression\_cluster3::2\_motifs \| Show matrix \| \| |  |  |  |  |  |  |
| cluster\_4 | 6 | cluster3\_motif41\_m::coexpression\_cluster3:: cluster3\_motif30\_m::coexpression\_cluster3:: cluster3\_motif28\_m::coexpression\_cluster3:: cluster3\_motif6\_r::coexpression\_cluster3:: cluster3\_motif11\_r::coexpression\_cluster3:: cluster3\_motif12\_r::coexpression\_cluster3:: | coexpression\_cluster3::6\_motifs | Show matrix | | |
| cluster\_5 | 7 | cluster3\_motif32\_m::coexpression\_cluster3:: cluster3\_motif31\_m::coexpression\_cluster3:: cluster3\_motif2\_r::coexpression\_cluster3:: cluster3\_motif25\_m::coexpression\_cluster3:: cluster3\_motif29\_m::coexpression\_cluster3:: cluster3\_motif37\_m::coexpression\_cluster3:: cluster3\_motif39\_m::coexpression\_cluster3:: | coexpression\_cluster3::7\_motifs | Show matrix | | |
| cluster\_6 | 3 | cluster3\_motif27\_m::coexpression\_cluster3:: cluster3\_motif24\_m::coexpression\_cluster3:: cluster3\_motif26\_m::coexpression\_cluster3:: | coexpression\_cluster3::3\_motifs | Show matrix | | |
| cluster\_7 | 2 | cluster3\_motif17\_r::coexpression\_cluster3:: cluster3\_motif43\_m::coexpression\_cluster3:: | coexpression\_cluster3::2\_motifs | Show matrix | | |
| cluster\_8 | 1 | cluster3\_motif20\_r::coexpression\_cluster3 | coexpression\_cluster3 : 1 motif | Show matrix | | |
| cluster\_9 | 1 | cluster3\_motif38\_m::coexpression\_cluster3 | coexpression\_cluster3 : 1 motif | Show matrix | | |

- Note: by default the table displays the first 50 clusters. Click on the *Show X entries* button to display more (or less) clusters.
- Click on the column names to change the order of the data.
- Use the *Search* window to find your motif (s) of interest.

| **Individual Cluster View** Individual Cluster View **Show All** **Hide All** **cluster\_1** **cluster\_2** **cluster\_3** **cluster\_4** **cluster\_5** **cluster\_6** **cluster\_7** **cluster\_8** **cluster\_9** **Display Logo Trees** **cluster\_1** **cluster\_2** **cluster\_3** **cluster\_4** **cluster\_5** **cluster\_6** **cluster\_7** **cluster\_8** **cluster\_9** **Display Branch-Motifs** **cluster\_1** Branch Motifs \| \| \| \| \| \| \|; Node \| Consensus \| Logo \| Logo (Reverse) \| Collections \| Matrix (Transfac format) \| IC \| Nb sites; Node 1 \| **-wwTATAkACAtaww------** \| \| \| coexpression\_cluster3::2\_motifs \| PSSM \| 8.75 \| 6081; Node 2 \| **--wwaTrCACACAcwww----** \| \| \| coexpression\_cluster3::2\_motifs \| PSSM \| 9.69 \| 1566; Node 3 \| **-wwTATAgAcAAAwwww----** \| \| \| coexpression\_cluster3::2\_motifs \| PSSM \| 9.21 \| 7123; Node 4 \| **-wwTATAgAcAAAwwww----** \| \| \| coexpression\_cluster3::3\_motifs \| PSSM \| 8.97 \| 9166; Node 5 \| **-wwtATAkACAwaww------** \| \| \| coexpression\_cluster3::3\_motifs \| PSSM \| 8.50 \| 8249; Node 6 \| **--wwwwArAmAAAAAww----** \| \| \| coexpression\_cluster3::2\_motifs \| PSSM \| 9.23 \| 7775; Node 7 \| **-CATATryATATATATA----** \| \| \| coexpression\_cluster3::2\_motifs \| PSSM \| 14.89 \| 1170; Node 8 \| **-wwTATAkACAwwww------** \| \| \| coexpression\_cluster3::4\_motifs \| PSSM \| 8.23 \| 12408; Node 9 \| **-wwwwwAgAmAAAaaww----** \| \| \| coexpression\_cluster3::5\_motifs \| PSSM \| 8.69 \| 16941; Node 10 \| **--wwaTrCACACAhwww----** \| \| \| coexpression\_cluster3::3\_motifs \| PSSM \| 8.70 \| 4377; Node 11 \| **-wwwaTAcACACAhawww---** \| \| \| coexpression\_cluster3::4\_motifs \| PSSM \| 8.92 \| 6125; Node 12 \| **-wwwATAgAcAAAwwww----** \| \| \| coexpression\_cluster3::9\_motifs \| PSSM \| 8.22 \| 29349; Node 13 \| **aTwwATmTACAww--------** \| \| \| coexpression\_cluster3::2\_motifs \| PSSM \| 8.95 \| 4213; Node 14 \| **-wwwATAgACAaAwwwww---** \| \| \| coexpression\_cluster3::13\_motifs \| PSSM \| 8.03 \| 35474; Node 15 \| **-yATAyAyAyATrTrTrTAta** \| \| \| coexpression\_cluster3::3\_motifs \| PSSM \| 13.67 \| 1962; Node 16 \| **-wwwATAgAcAaAwawwwAta** \| \| \| coexpression\_cluster3::16\_motifs \| PSSM \| 9.26 \| 37436; Node 17 \| **awwwATAkACAwAwawwwAta** \| \| \| coexpression\_cluster3::18\_motifs \| PSSM \| 9.44 \| 41648 **cluster\_2** Branch Motifs \| \| \| \| \| \| \|; Node \| Consensus \| Logo \| Logo (Reverse) \| Collections \| Matrix (Transfac format) \| IC \| Nb sites; Node 1 \| **thwwrtgCACACACAytyrkaw-** \| \| \| coexpression\_cluster3::2\_motifs \| PSSM \| 7.86 \| 644; Node 2 \| **twwaawkCACACACaywwtgrwy** \| \| \| coexpression\_cluster3::3\_motifs \| PSSM \| 7.84 \| 1143; Node 3 \| **wwwaawtyrCACACAywwwgrwy** \| \| \| coexpression\_cluster3::4\_motifs \| PSSM \| 7.76 \| 1481 **cluster\_3** Branch Motifs \| \| \| \| \| \| \|; Node \| Consensus \| Logo \| Logo (Reverse) \| Collections \| Matrix (Transfac format) \| IC \| Nb sites; Node 1 \| **WTATKAACAaktMAKA** \| \| \| coexpression\_cluster3::2\_motifs \| PSSM \| 13.68 \| 258 **cluster\_4** Branch Motifs \| \| \| \| \| \| \|; Node \| Consensus \| Logo \| Logo (Reverse) \| Collections \| Matrix (Transfac format) \| IC \| Nb sites; Node 1 \| **-----wwAgCTAGCTAwwww--** \| \| \| coexpression\_cluster3::2\_motifs \| PSSM \| 8.76 \| 1495; Node 2 \| **----wwwAgCTAGCTAwwwww-** \| \| \| coexpression\_cluster3::3\_motifs \| PSSM \| 9.04 \| 1754; Node 3 \| **----wwwAgcTAGCTAwwwww-** \| \| \| coexpression\_cluster3::4\_motifs \| PSSM \| 9.05 \| 1933; Node 4 \| **----wwwAgCTAGCTAwwwww-** \| \| \| coexpression\_cluster3::5\_motifs \| PSSM \| 9.11 \| 2063; Node 5 \| **yygtwwwAgCTAGCTAwwwwwr** \| \| \| coexpression\_cluster3::6\_motifs \| PSSM \| 9.25 \| 2196 **cluster\_5** Branch Motifs \| \| \| \| \| \| \|; Node \| Consensus \| Logo \| Logo (Reverse) \| Collections \| Matrix (Transfac format) \| IC \| Nb sites; Node 1 \| **AAAAAAArarAAwAA-** \| \| \| coexpression\_cluster3::2\_motifs \| PSSM \| 12.02 \| 1593; Node 2 \| **aaAAAArAmAAAaAaw** \| \| \| coexpression\_cluster3::3\_motifs \| PSSM \| 10.39 \| 10168; Node 3 \| **aaAAAArAmAAAaAaw** \| \| \| coexpression\_cluster3::4\_motifs \| PSSM \| 10.28 \| 10539; Node 4 \| **aaAAAArAmAAAaAaw** \| \| \| coexpression\_cluster3::5\_motifs \| PSSM \| 10.21 \| 10882; Node 5 \| **-AAwTAAAAmAAAaw-** \| \| \| coexpression\_cluster3::2\_motifs \| PSSM \| 11.03 \| 1193; Node 6 \| **aaAAAArAmAAAaAww** \| \| \| coexpression\_cluster3::7\_motifs \| PSSM \| 9.80 \| 12075 **cluster\_6** Branch Motifs \| \| \| \| \| \| \|; Node \| Consensus \| Logo \| Logo (Reverse) \| Collections \| Matrix (Transfac format) \| IC \| Nb sites; Node 1 \| **---TrCAYACAy** \| \| \| coexpression\_cluster3::2\_motifs \| PSSM \| 8.99 \| 1459; Node 2 \| **rTATrCAyACAy** \| \| \| coexpression\_cluster3::3\_motifs \| PSSM \| 10.58 \| 1762 **cluster\_7** Branch Motifs \| \| \| \| \| \| \|; Node \| Consensus \| Logo \| Logo (Reverse) \| Collections \| Matrix (Transfac format) \| IC \| Nb sites; Node 1 \| **yrtwtATTCTAwwc** \| \| \| coexpression\_cluster3::2\_motifs \| PSSM \| 8.92 \| 772 **cluster\_8** Branch Motifs \| \| \| \| \| \| \|; Node \| Consensus \| Logo \| Logo (Reverse) \| Collections \| Matrix (Transfac format) \| IC \| Nb sites; Singleton \| **wtTCTTCAww** \| \| \| coexpression\_cluster3 : 1 motif \| PSSM \| 9.20 \| 326 **cluster\_9** Branch Motifs \| \| \| \| \| \| \|; Node \| Consensus \| Logo \| Logo (Reverse) \| Collections \| Matrix (Transfac format) \| IC \| Nb sites; Singleton \| **aaytratrr** \| \| \| coexpression\_cluster3 : 1 motif \| PSSM \| 9.47 \| 692 - **Each cluster has a different color.** - **Click on the upper buttons to display separately the information of each cluster.** - **Click on the circles at each tree to display its corresponding merged motifs.** Definitions - **Logo tree.** This results shows the hierarchical tree with its logo alignment of one cluster. The logos are shown in Forward (left logo) and Reverse (right logo) orientation. - **Branch-motifs.** On the trees are displayed the branch number that is the number in which the motifs were incorporated in the tree. The Branch-Motifs table shows the logo in both orientations and the link to the file in TRANSFAC format with the branch-motif. |
|  | **Individual Motif View** Individual Motif View Motif id \| Motif name \| Cluster \| Collection \| Width \| IC \| Number of sites \| Consensus \| Consensus (Rev) \| Logo \| Logo (Rev); coexpression\_cluster3\_m30\_cluster3\_motif30\_m \| cluster3\_motif30\_m \| cluster\_4 \| coexpression\_cluster3 \| 22 \| 10.18 \| 130 \| **+++++TRGCTAGYTRT++++++** \| **++++++ayARCTAGCyA+++++** \| \|; coexpression\_cluster3\_m41\_cluster3\_motif41\_m \| cluster3\_motif41\_m \| cluster\_4 \| coexpression\_cluster3 \| 22 \| 9.77 \| 133 \| **YAKMGYBAGCTAGCTRWYACRR** \| **yygtrwyAGCTAGCTvrckmtr** \| \|; coexpression\_cluster3\_m38\_cluster3\_motif38\_m \| cluster3\_motif38\_m \| cluster\_9 \| coexpression\_cluster3 \| 9 \| 9.47 \| 692 \| **AATTAATAA** \| **TTATTAATT** \| \|; coexpression\_cluster3\_m10\_cluster3\_motif10\_r \| cluster3\_motif10\_r \| cluster\_1 \| coexpression\_cluster3 \| 21 \| 8.93 \| 2783 \| **+wwTATAkACAwwww++++++** \| **++++++WWWWTGTMTATAWW+** \| \|; coexpression\_cluster3\_m17\_cluster3\_motif17\_r \| cluster3\_motif17\_r \| cluster\_7 \| coexpression\_cluster3 \| 14 \| 9.22 \| 684 \| **+++wwATTCTAww+** \| **+WWTAGAATWW+++** \| \|; coexpression\_cluster3\_m26\_cluster3\_motif26\_m \| cluster3\_motif26\_m \| cluster\_6 \| coexpression\_cluster3 \| 12 \| 8.68 \| 1000 \| **+++TrCAyACA+** \| **+TGTRTGYA+++** \| \|; coexpression\_cluster3\_m29\_cluster3\_motif29\_m \| cluster3\_motif29\_m \| cluster\_5 \| coexpression\_cluster3 \| 16 \| 16.21 \| 593 \| **AAAAAAAAAAAATAA+** \| **+TTATTTTTTTTTTTT** \| \|; coexpression\_cluster3\_m24\_cluster3\_motif24\_m \| cluster3\_motif24\_m \| cluster\_6 \| coexpression\_cluster3 \| 12 \| 9.68 \| 459 \| **+++TGCACACAY** \| **rTGTGTGCA+++** \| \|; coexpression\_cluster3\_m19\_cluster3\_motif19\_r \| cluster3\_motif19\_r \| cluster\_1 \| coexpression\_cluster3 \| 21 \| 7.77 \| 3909 \| **++wwATmTACAww++++++++** \| **++++++++WWTGTAKATWW++** \| \|; coexpression\_cluster3\_m33\_cluster3\_motif33\_m \| cluster3\_motif33\_m \| cluster\_3 \| coexpression\_cluster3 \| 16 \| 13.54 \| 111 \| **WTATKAACAaktCAK+** \| **+MTGAMTTGTTMATAW** \| \|; coexpression\_cluster3\_m43\_cluster3\_motif43\_m \| cluster3\_motif43\_m \| cluster\_7 \| coexpression\_cluster3 \| 14 \| 9.18 \| 88 \| **YRTTTGTTCTAYGC** \| **gcrTAGAACAAayr** \| \|; coexpression\_cluster3\_m16\_cluster3\_motif16\_r \| cluster3\_motif16\_r \| cluster\_1 \| coexpression\_cluster3 \| 21 \| 12.66 \| 792 \| **+TATAYAYAYATRTRTRTATA** \| **taTAyAyAyATrTrTrTAta+** \| \|; coexpression\_cluster3\_m32\_cluster3\_motif32\_m \| cluster3\_motif32\_m \| cluster\_5 \| coexpression\_cluster3 \| 16 \| 10.98 \| 344 \| **++TTTTTTCTCTTTT+** \| **+AAAAGAGAAaAAa++** \| \|; coexpression\_cluster3\_m31\_cluster3\_motif31\_m \| cluster3\_motif31\_m \| cluster\_5 \| coexpression\_cluster3 \| 16 \| 11.80 \| 372 \| **TTTTCCTTTTTTT+++** \| **+++AAAAAAAGGAAAA** \| \|; coexpression\_cluster3\_m23\_cluster3\_motif23\_m \| cluster3\_motif23\_m \| cluster\_1 \| coexpression\_cluster3 \| 21 \| 14.46 \| 687 \| **++ATATryATATATATA++++** \| **++++TATATATATRYATAT++** \| \|; coexpression\_cluster3\_m4\_cluster3\_motif4\_r \| cluster3\_motif4\_r \| cluster\_1 \| coexpression\_cluster3 \| 21 \| 8.85 \| 3298 \| **+wwtATAkACATAww++++++** \| **++++++WWTATGTMTATAWW+** \| \|; coexpression\_cluster3\_m18\_cluster3\_motif18\_r \| cluster3\_motif18\_r \| cluster\_1 \| coexpression\_cluster3 \| 21 \| 9.78 \| 628 \| **++WWWTGCACACACWWW++++** \| **++++wwwgTGTGTGcAwww++** \| \|; coexpression\_cluster3\_m25\_cluster3\_motif25\_m \| cluster3\_motif25\_m \| cluster\_5 \| coexpression\_cluster3 \| 16 \| 11.75 \| 1000 \| **aAAAAAArrrRAaAA+** \| **+TTTTYYYYTTTTTTT** \| \|; coexpression\_cluster3\_m36\_cluster3\_motif36\_m \| cluster3\_motif36\_m \| cluster\_1 \| coexpression\_cluster3 \| 21 \| 8.90 \| 304 \| **aTGTAyrTACAt+++++++++** \| **+++++++++ATGTAYRTACAT** \| \|; coexpression\_cluster3\_m20\_cluster3\_motif20\_r \| cluster3\_motif20\_r \| cluster\_8 \| coexpression\_cluster3 \| 10 \| 9.20 \| 326 \| **wtTCTTCAww** \| **WWTGAAGAAW** \| \|; coexpression\_cluster3\_m15\_cluster3\_motif15\_r \| cluster3\_motif15\_r \| cluster\_1 \| coexpression\_cluster3 \| 21 \| 8.98 \| 2168 \| **+WWWATAGACAAWWW++++++** \| **++++++wwwTTGTcTATwww+** \| \|; coexpression\_cluster3\_m9\_cluster3\_motif9\_r \| cluster3\_motif9\_r \| cluster\_1 \| coexpression\_cluster3 \| 21 \| 9.50 \| 3429 \| **++WWTTARAMAAAAAWW++++** \| **++++wwTTTTTkTyTAaww++** \| \|; coexpression\_cluster3\_m13\_cluster3\_motif13\_r \| cluster3\_motif13\_r \| cluster\_1 \| coexpression\_cluster3 \| 21 \| 9.34 \| 2336 \| **+wwwrTAgACAAAawww++++** \| **++++WWWTTTTGTCTAYWWW+** \| \|; coexpression\_cluster3\_m34\_cluster3\_motif34\_m \| cluster3\_motif34\_m \| cluster\_3 \| coexpression\_cluster3 \| 16 \| 12.86 \| 147 \| **+TATKAACAAKTMAKA** \| **TMTkamtTGTTmATA+** \| \|; coexpression\_cluster3\_m42\_cluster3\_motif42\_m \| cluster3\_motif42\_m \| cluster\_2 \| coexpression\_cluster3 \| 23 \| 7.11 \| 456 \| **thdwgwryACACACayyyrgrw+** \| **+WYCYRRRTGTGTGTRYWCWHDA** \| \|; coexpression\_cluster3\_m27\_cluster3\_motif27\_m \| cluster3\_motif27\_m \| cluster\_6 \| coexpression\_cluster3 \| 12 \| 8.94 \| 303 \| **RTATGCATAY++** \| **++rTATGCATAy** \| \|; coexpression\_cluster3\_m5\_cluster3\_motif5\_r \| cluster3\_motif5\_r \| cluster\_1 \| coexpression\_cluster3 \| 21 \| 9.83 \| 938 \| **++wwATrCACACAcwww++++** \| **++++WWWGTGTGTGYATWW++** \| \|; coexpression\_cluster3\_m37\_cluster3\_motif37\_m \| cluster3\_motif37\_m \| cluster\_5 \| coexpression\_cluster3 \| 16 \| 12.38 \| 731 \| **+AATKAAAAAAAAaw+** \| **+WTTTTTTTTTMATT+** \| \|; coexpression\_cluster3\_m1\_cluster3\_motif1\_r \| cluster3\_motif1\_r \| cluster\_1 \| coexpression\_cluster3 \| 21 \| 9.41 \| 4787 \| **+wwTATAkAcAAAwwww++++** \| **++++WWWWTTTGTMTATAWW+** \| \|; coexpression\_cluster3\_m21\_cluster3\_motif21\_r \| cluster3\_motif21\_r \| cluster\_1 \| coexpression\_cluster3 \| 21 \| 7.93 \| 4159 \| **+WWTATAKACAWW++++++++** \| **++++++++wwtGTmTATAww+** \| \|; coexpression\_cluster3\_m40\_cluster3\_motif40\_m \| cluster3\_motif40\_m \| cluster\_2 \| coexpression\_cluster3 \| 23 \| 11.81 \| 189 \| **TrtwwtgCRCACACACrwrkmw+** \| **+WKMYWYGTGTGTGYGCAWWAYA** \| \|; coexpression\_cluster3\_m8\_cluster3\_motif8\_r \| cluster3\_motif8\_r \| cluster\_1 \| coexpression\_cluster3 \| 21 \| 10.11 \| 1748 \| **+WWWTGTGTGTMTATAWW+++** \| **+++wwTATAkACAcAcAwww+** \| \|; coexpression\_cluster3\_m14\_cluster3\_motif14\_r \| cluster3\_motif14\_r \| cluster\_1 \| coexpression\_cluster3 \| 21 \| 9.11 \| 2043 \| **+WWWATAGACAAWDWWW++++** \| **++++wwwhwTTGTcTATwww+** \| \|; coexpression\_cluster3\_m6\_cluster3\_motif6\_r \| cluster3\_motif6\_r \| cluster\_4 \| coexpression\_cluster3 \| 22 \| 10.33 \| 259 \| **++++WWAAMTAGCTAGCYAWW+** \| **+wwTrgCTAGCTAktTww++++** \| \|; coexpression\_cluster3\_m28\_cluster3\_motif28\_m \| cluster3\_motif28\_m \| cluster\_4 \| coexpression\_cluster3 \| 22 \| 10.13 \| 179 \| **+++++++AAHTAGCTADTT+++** \| **+++AahTAGCTAdtT+++++++** \| \|; coexpression\_cluster3\_m35\_cluster3\_motif35\_m \| cluster3\_motif35\_m \| cluster\_2 \| coexpression\_cluster3 \| 23 \| 10.73 \| 339 \| **CYYKRYGTGTGTGCAAAWTSAW+** \| **+wtsawttTGCACACACrymrrg** \| \|; coexpression\_cluster3\_m12\_cluster3\_motif12\_r \| cluster3\_motif12\_r \| cluster\_4 \| coexpression\_cluster3 \| 22 \| 9.03 \| 682 \| **+++++wwAGCTAGCTAAwww++** \| **++WWWTTAGCTAGCTWW+++++** \| \|; coexpression\_cluster3\_m44\_cluster3\_motif44\_m \| cluster3\_motif44\_m \| cluster\_2 \| coexpression\_cluster3 \| 23 \| 9.69 \| 500 \| **MWWAMMTCACACACATWWTGRAY** \| **rtycawwatGTGTGTgakktwwk** \| \|; coexpression\_cluster3\_m11\_cluster3\_motif11\_r \| cluster3\_motif11\_r \| cluster\_4 \| coexpression\_cluster3 \| 22 \| 8.78 \| 813 \| **+++++wwAgCTAGCTATww+++** \| **+++WWATAGCTAGCTWW+++++** \| \|; coexpression\_cluster3\_m3\_cluster3\_motif3\_r \| cluster3\_motif3\_r \| cluster\_1 \| coexpression\_cluster3 \| 21 \| 8.32 \| 2811 \| **+++WWWTGTGTGTATW+++++** \| **+++++watACACACAwww+++** \| \|; coexpression\_cluster3\_m22\_cluster3\_motif22\_m \| cluster3\_motif22\_m \| cluster\_1 \| coexpression\_cluster3 \| 21 \| 14.36 \| 483 \| **+CATAYRYATATATAT+++++** \| **+++++ATATATATrYRTATG+** \| \|; coexpression\_cluster3\_m7\_cluster3\_motif7\_r \| cluster3\_motif7\_r \| cluster\_1 \| coexpression\_cluster3 \| 21 \| 9.79 \| 4346 \| **++WWWAARAMAAAAAWW++++** \| **++++wwTTTTTkTyTTwww++** \| \|; coexpression\_cluster3\_m2\_cluster3\_motif2\_r \| cluster3\_motif2\_r \| cluster\_5 \| coexpression\_cluster3 \| 16 \| 10.55 \| 8575 \| **WWTTTTTKTYTTTTTW** \| **waAAAArAmAAAaaww** \| \|; coexpression\_cluster3\_m39\_cluster3\_motif39\_m \| cluster3\_motif39\_m \| cluster\_5 \| coexpression\_cluster3 \| 16 \| 11.26 \| 462 \| **+AAATAAAACAA++++** \| **++++TTGTTTTATTT+** \| \| - Click on the column names to change the order of the data. - Write the name of one cluster, collection or a pattern in the *Search* window. **Heatmap View** Heatmap View Distance table \| PDF |

**Additional Files**

#### Exported files

| File | Description |
| --- | --- |
| **Pairwise comparison** | This table shows the pairwise comparison between all the input motifs using different metrics. This is the *compare-matrices* result. |
| **Matrix description** | This table shows information of each input motif. |
| **Cluster-wise Motif Files [zip]** | One TRANSFAC file is generated for each cluster including all the input motifs grouped in the corresponding cluster. |
| **Root motifs** | A file with the root matrix of each cluster. |
| **Motif collections per cluster** | A tab separated file containing the counts of motif of each collection on each cluster. |
| **Clusters** | A tab separated file containing the clusters and their correspondng motifs. |
| **Central motifs IDs** | A tab separated file containing the central motif, that one who is the most similar to all the others motif in the same cluster. |
| **Central motifs (in TRANSFAC format)** | A transfac file containing the central motif, that one who is the most similar to all the others motif within the same cluster. |
| **Input motifs** | The input motifs analyzed with matrix-clustering. |

- **References**
  1. Castro-Mondragon JA et al.  RSAT matrix-clustering: dynamic exploration and redundancy reduction of transcription factor binding motif collections. doi: 10.1093/nar/gkx314; Nucleic Acids Research (2017)
  2. **Last RSAT release:** Nguyen NTT et al. RSAT 2018: regulatory sequence analysis tools 20th anniversary. Nucleic Acids Research (2018).
- **Additional Information**
  - Jacques van Helden course
- **Contact**
  - Jaime Castro-Mondragon - ****
  - Morgane Thomas-Chollier - ****
  - Jacques van Helden - ****
