## Additional file 10 for "Transcriptome analysis reveals a novel DNA element that may interact with chromatin-associated proteins in *Plasmodium berghei* during erythrocytic development": matrix-clustering_SUMMARY_cluster1_1.html

matrix-clustering coexpression\_cluster1\_1

### RSAT - matrix-clustering result

##### Analysis: coexpression\_cluster1\_1 (08/09/2023 14:47)

##### Command

```
matrix-clustering  -v 1 -max_matrices 300 -matrix coexpression_cluster1_1 $RSAT/public_html/tmp/www-data/2023/09/08/matrix-clustering_2023-09-08.144739_G7IEX9/matrix-clustering_query_matrices.transfac transfac -hclust_method average -calc sum -title coexpression_cluster1_1 -metric_build_tree Ncor -lth w 5 -lth cor 0.6 -lth Ncor 0.4 -quick -label_in_tree name -return json,heatmap -o $RSAT/public_html/tmp/www-data/2023/09/08/matrix-clustering_2023-09-08.144739_G7IEX9/matrix-clustering
```

#### Results Summary

|  |  |  |  |  |  |
| --- | --- | --- | --- | --- | --- |
| Nb Input motifs | Nb Input collections | Nb Clusters Found | Download   Root motifs | Complete results [zip] | Parameters used Linkage method \| Similarity metric \| Thresholds to partition the tree |
| 23 | 1 | 12 | **Download** | Download | average \| Ncor \| **Ncor** = 0.4 **cor** = 0.6 |

|  |  |  |  |  |  |  |
| --- | --- | --- | --- | --- | --- | --- |
|  | **Logo Forest (dynamic browsing)**  **Logo Forest (rapid overview - low image quality)**  cluster\_1  cluster\_2  cluster\_3  cluster\_4  cluster\_5  cluster\_6  cluster\_7  cluster\_8  cluster\_9  cluster\_10  cluster\_11  cluster\_12 |  |  |  |  |  |
|  | **Clusters Summary** Clusters Summary Table (Root Motif \| Root Motif (Reverse) \| Cluster ID \| # Motifs \| Motif\_Name::Collection \| Number of Motifs by Collection \| Root Motif (transfac Format)) \| | cluster\_1 | 4 | cluster1\_1\_motif16\_m::coexpression\_cluster1\_1:: cluster1\_1\_motif6\_r::coexpression\_cluster1\_1:: cluster1\_1\_motif12\_r::coexpression\_cluster1\_1:: cluster1\_1\_motif2\_r::coexpression\_cluster1\_1:: | coexpression\_cluster1\_1::4\_motifs | Show matrix |
| cluster\_2 \| 4 \| cluster1\_1\_motif1\_r::coexpression\_cluster1\_1:: cluster1\_1\_motif11\_r::coexpression\_cluster1\_1:: cluster1\_1\_motif3\_r::coexpression\_cluster1\_1:: cluster1\_1\_motif4\_r::coexpression\_cluster1\_1:: \| coexpression\_cluster1\_1::4\_motifs \| Show matrix \| \| |  |  |  |  |  |  |
| cluster\_3 \| 4 \| cluster1\_1\_motif5\_r::coexpression\_cluster1\_1:: cluster1\_1\_motif8\_r::coexpression\_cluster1\_1:: cluster1\_1\_motif13\_m::coexpression\_cluster1\_1:: cluster1\_1\_motif14\_m::coexpression\_cluster1\_1:: \| coexpression\_cluster1\_1::4\_motifs \| Show matrix \| \| |  |  |  |  |  |  |
| cluster\_4 | 3 | cluster1\_1\_motif10\_r::coexpression\_cluster1\_1:: cluster1\_1\_motif7\_r::coexpression\_cluster1\_1:: cluster1\_1\_motif9\_r::coexpression\_cluster1\_1:: | coexpression\_cluster1\_1::3\_motifs | Show matrix | | |
| cluster\_5 | 1 | cluster1\_1\_motif15\_m::coexpression\_cluster1\_1 | coexpression\_cluster1\_1 : 1 motif | Show matrix | | |
| cluster\_6 | 1 | cluster1\_1\_motif19\_m::coexpression\_cluster1\_1 | coexpression\_cluster1\_1 : 1 motif | Show matrix | | |
| cluster\_7 | 1 | cluster1\_1\_motif22\_m::coexpression\_cluster1\_1 | coexpression\_cluster1\_1 : 1 motif | Show matrix | | |
| cluster\_8 | 1 | cluster1\_1\_motif23\_m::coexpression\_cluster1\_1 | coexpression\_cluster1\_1 : 1 motif | Show matrix | | |
| cluster\_9 | 1 | cluster1\_1\_motif17\_m::coexpression\_cluster1\_1 | coexpression\_cluster1\_1 : 1 motif | Show matrix | | |
| cluster\_10 | 1 | cluster1\_1\_motif20\_m::coexpression\_cluster1\_1 | coexpression\_cluster1\_1 : 1 motif | Show matrix | | |
| cluster\_11 | 1 | cluster1\_1\_motif21\_m::coexpression\_cluster1\_1 | coexpression\_cluster1\_1 : 1 motif | Show matrix | | |
| cluster\_12 | 1 | cluster1\_1\_motif18\_m::coexpression\_cluster1\_1 | coexpression\_cluster1\_1 : 1 motif | Show matrix | | |

- Note: by default the table displays the first 50 clusters. Click on the *Show X entries* button to display more (or less) clusters.
- Click on the column names to change the order of the data.
- Use the *Search* window to find your motif (s) of interest.

| **Individual Cluster View** Individual Cluster View **Show All** **Hide All** **cluster\_1** **cluster\_2** **cluster\_3** **cluster\_4** **cluster\_5** **cluster\_6** **cluster\_7** **cluster\_8** **cluster\_9** **cluster\_10** **cluster\_11** **cluster\_12** **Display Logo Trees** **cluster\_1** **cluster\_2** **cluster\_3** **cluster\_4** **cluster\_5** **cluster\_6** **cluster\_7** **cluster\_8** **cluster\_9** **cluster\_10** **cluster\_11** **cluster\_12** **Display Branch-Motifs** **cluster\_1** Branch Motifs \| \| \| \| \| \| \|; Node \| Consensus \| Logo \| Logo (Reverse) \| Collections \| Matrix (Transfac format) \| IC \| Nb sites; Node 1 \| **wwTGCATGCAww--** \| \| \| coexpression\_cluster1\_1::2\_motifs \| PSSM \| 8.21 \| 673; Node 2 \| **wwTGCATGCAtwww** \| \| \| coexpression\_cluster1\_1::3\_motifs \| PSSM \| 8.53 \| 1058; Node 3 \| **wwTGCATGCAtwww** \| \| \| coexpression\_cluster1\_1::4\_motifs \| PSSM \| 8.64 \| 1171 **cluster\_2** Branch Motifs \| \| \| \| \| \| \|; Node \| Consensus \| Logo \| Logo (Reverse) \| Collections \| Matrix (Transfac format) \| IC \| Nb sites; Node 1 \| **wwwwATTTAAATTATAATTTAwwww--** \| \| \| coexpression\_cluster1\_1::2\_motifs \| PSSM \| 13.54 \| 2263; Node 2 \| **wwwwATTTAAATTATAATTTAwwwwww** \| \| \| coexpression\_cluster1\_1::3\_motifs \| PSSM \| 14.06 \| 3341; Node 3 \| **wwwwwTwTAAwTTAwAwTwTawwwwww** \| \| \| coexpression\_cluster1\_1::4\_motifs \| PSSM \| 12.66 \| 5673 **cluster\_3** Branch Motifs \| \| \| \| \| \| \|; Node \| Consensus \| Logo \| Logo (Reverse) \| Collections \| Matrix (Transfac format) \| IC \| Nb sites; Node 1 \| **wwACATmTwww---** \| \| \| coexpression\_cluster1\_1::2\_motifs \| PSSM \| 7.06 \| 1545; Node 2 \| **ATAyATATATATrT** \| \| \| coexpression\_cluster1\_1::2\_motifs \| PSSM \| 14.39 \| 134; Node 3 \| **wwACATmTwtwTrT** \| \| \| coexpression\_cluster1\_1::4\_motifs \| PSSM \| 9.55 \| 1679 **cluster\_4** Branch Motifs \| \| \| \| \| \| \|; Node \| Consensus \| Logo \| Logo (Reverse) \| Collections \| Matrix (Transfac format) \| IC \| Nb sites; Node 1 \| **wwwTATTAmAww** \| \| \| coexpression\_cluster1\_1::2\_motifs \| PSSM \| 7.86 \| 3108; Node 2 \| **wwwtATwAaaww** \| \| \| coexpression\_cluster1\_1::3\_motifs \| PSSM \| 6.52 \| 5337 **cluster\_5** Branch Motifs \| \| \| \| \| \| \|; Node \| Consensus \| Logo \| Logo (Reverse) \| Collections \| Matrix (Transfac format) \| IC \| Nb sites; Singleton \| **aaaGaGaamaartSv** \| \| \| coexpression\_cluster1\_1 : 1 motif \| PSSM \| 12.30 \| 49 **cluster\_6** Branch Motifs \| \| \| \| \| \| \|; Node \| Consensus \| Logo \| Logo (Reverse) \| Collections \| Matrix (Transfac format) \| IC \| Nb sites; Singleton \| **aaaattarCrCaaaa** \| \| \| coexpression\_cluster1\_1 : 1 motif \| PSSM \| 15.03 \| 42 **cluster\_7** Branch Motifs \| \| \| \| \| \| \|; Node \| Consensus \| Logo \| Logo (Reverse) \| Collections \| Matrix (Transfac format) \| IC \| Nb sites; Singleton \| **CCtGSMtGaGSGGyK** \| \| \| coexpression\_cluster1\_1 : 1 motif \| PSSM \| 13.58 \| 13 **cluster\_8** Branch Motifs \| \| \| \| \| \| \|; Node \| Consensus \| Logo \| Logo (Reverse) \| Collections \| Matrix (Transfac format) \| IC \| Nb sites; Singleton \| **SrYttCMyGaCSSat** \| \| \| coexpression\_cluster1\_1 : 1 motif \| PSSM \| 13.95 \| 14 **cluster\_9** Branch Motifs \| \| \| \| \| \| \|; Node \| Consensus \| Logo \| Logo (Reverse) \| Collections \| Matrix (Transfac format) \| IC \| Nb sites; Singleton \| **tSrGttCraRtCy** \| \| \| coexpression\_cluster1\_1 : 1 motif \| PSSM \| 14.30 \| 11 **cluster\_10** Branch Motifs \| \| \| \| \| \| \|; Node \| Consensus \| Logo \| Logo (Reverse) \| Collections \| Matrix (Transfac format) \| IC \| Nb sites; Singleton \| **SSSGGSaVmYSBSSG** \| \| \| coexpression\_cluster1\_1 : 1 motif \| PSSM \| 10.45 \| 19 **cluster\_11** Branch Motifs \| \| \| \| \| \| \|; Node \| Consensus \| Logo \| Logo (Reverse) \| Collections \| Matrix (Transfac format) \| IC \| Nb sites; Singleton \| **CGSCCSKaMGGtwRS** \| \| \| coexpression\_cluster1\_1 : 1 motif \| PSSM \| 13.67 \| 14 **cluster\_12** Branch Motifs \| \| \| \| \| \| \|; Node \| Consensus \| Logo \| Logo (Reverse) \| Collections \| Matrix (Transfac format) \| IC \| Nb sites; Singleton \| **GCCSCC** \| \| \| coexpression\_cluster1\_1 : 1 motif \| PSSM \| 6.22 \| 60 - **Each cluster has a different color.** - **Click on the upper buttons to display separately the information of each cluster.** - **Click on the circles at each tree to display its corresponding merged motifs.** Definitions - **Logo tree.** This results shows the hierarchical tree with its logo alignment of one cluster. The logos are shown in Forward (left logo) and Reverse (right logo) orientation. - **Branch-motifs.** On the trees are displayed the branch number that is the number in which the motifs were incorporated in the tree. The Branch-Motifs table shows the logo in both orientations and the link to the file in TRANSFAC format with the branch-motif. |
|  | **Individual Motif View** Individual Motif View Motif id \| Motif name \| Cluster \| Collection \| Width \| IC \| Number of sites \| Consensus \| Consensus (Rev) \| Logo \| Logo (Rev); coexpression\_cluster1\_1\_m21\_cluster1\_1\_motif21\_m \| cluster1\_1\_motif21\_m \| cluster\_11 \| coexpression\_cluster1\_1 \| 15 \| 13.67 \| 14 \| **CGkyYSkAmGGTwrS** \| **SYWACCKTMSRRMCG** \| \|; coexpression\_cluster1\_1\_m7\_cluster1\_1\_motif7\_r \| cluster1\_1\_motif7\_r \| cluster\_4 \| coexpression\_cluster1\_1 \| 12 \| 7.65 \| 2194 \| **+wwTATTAyAww** \| **WWTRTAATAWW+** \| \|; coexpression\_cluster1\_1\_m20\_cluster1\_1\_motif20\_m \| cluster1\_1\_motif20\_m \| cluster\_10 \| coexpression\_cluster1\_1 \| 15 \| 10.45 \| 19 \| **sysGGswvAYCkscG** \| **CGSMGRTBWSCCSRS** \| \|; coexpression\_cluster1\_1\_m16\_cluster1\_1\_motif16\_m \| cluster1\_1\_motif16\_m \| cluster\_1 \| coexpression\_cluster1\_1 \| 14 \| 8.56 \| 113 \| **+++ATGCATGY+++** \| **+++rCATGCAT+++** \| \|; coexpression\_cluster1\_1\_m3\_cluster1\_1\_motif3\_r \| cluster1\_1\_motif3\_r \| cluster\_2 \| coexpression\_cluster1\_1 \| 27 \| 13.50 \| 1169 \| **+wwTATTTAAATTwwAATTTAwwww++** \| **++WWWWTAAATTWWAATTTAAATAWW+** \| \|; coexpression\_cluster1\_1\_m12\_cluster1\_1\_motif12\_r \| cluster1\_1\_motif12\_r \| cluster\_1 \| coexpression\_cluster1\_1 \| 14 \| 7.86 \| 409 \| **wwtGCATGCAww++** \| **++WWTGCATGCAWW** \| \|; coexpression\_cluster1\_1\_m1\_cluster1\_1\_motif1\_r \| cluster1\_1\_motif1\_r \| cluster\_2 \| coexpression\_cluster1\_1 \| 27 \| 11.66 \| 2332 \| **++WWWWATAATTTAAATTATWWWW+++** \| **+++wwwwATAATTTAAATTATwwww++** \| \|; coexpression\_cluster1\_1\_m5\_cluster1\_1\_motif5\_r \| cluster1\_1\_motif5\_r \| cluster\_3 \| coexpression\_cluster1\_1 \| 14 \| 7.52 \| 767 \| **wwACATyTTww+++** \| **+++WWAARATGTWW** \| \|; coexpression\_cluster1\_1\_m19\_cluster1\_1\_motif19\_m \| cluster1\_1\_motif19\_m \| cluster\_6 \| coexpression\_cluster1\_1 \| 15 \| 15.03 \| 42 \| **AAAATwAAmamAWAA** \| **TTWTKTKTTWATTTT** \| \|; coexpression\_cluster1\_1\_m13\_cluster1\_1\_motif13\_m \| cluster1\_1\_motif13\_m \| cluster\_3 \| coexpression\_cluster1\_1 \| 14 \| 15.52 \| 66 \| **ATATATATATAT++** \| **++ATATATATATAT** \| \|; coexpression\_cluster1\_1\_m15\_cluster1\_1\_motif15\_m \| cluster1\_1\_motif15\_m \| cluster\_5 \| coexpression\_cluster1\_1 \| 15 \| 12.30 \| 49 \| **AAArAGAAAAAATga** \| **TCATTTTTTCTYTTT** \| \|; coexpression\_cluster1\_1\_m11\_cluster1\_1\_motif11\_r \| cluster1\_1\_motif11\_r \| cluster\_2 \| coexpression\_cluster1\_1 \| 27 \| 14.38 \| 1078 \| **WWWWATWTAAATTATAATTTAAWWWWW** \| **wwwwwTTAAATTATAATTTAWATwwww** \| \|; coexpression\_cluster1\_1\_m23\_cluster1\_1\_motif23\_m \| cluster1\_1\_motif23\_m \| cluster\_8 \| coexpression\_cluster1\_1 \| 15 \| 13.95 \| 14 \| **cAYTTCMTGACcCAT** \| **ATGGGTCAKGAARTG** \| \|; coexpression\_cluster1\_1\_m2\_cluster1\_1\_motif2\_r \| cluster1\_1\_motif2\_r \| cluster\_1 \| coexpression\_cluster1\_1 \| 14 \| 8.95 \| 264 \| **WWTGCATGCAWW++** \| **++wwTGCATGCAww** \| \|; coexpression\_cluster1\_1\_m8\_cluster1\_1\_motif8\_r \| cluster1\_1\_motif8\_r \| cluster\_3 \| coexpression\_cluster1\_1 \| 14 \| 7.79 \| 778 \| **wwACATmTAww+++** \| **+++WWTAKATGTWW** \| \|; coexpression\_cluster1\_1\_m4\_cluster1\_1\_motif4\_r \| cluster1\_1\_motif4\_r \| cluster\_2 \| coexpression\_cluster1\_1 \| 27 \| 13.58 \| 1094 \| **wwwwATTTAAATTATAATTTAwww+++** \| **+++WWWTAAATTATAATTTAAATWWWW** \| \|; coexpression\_cluster1\_1\_m18\_cluster1\_1\_motif18\_m \| cluster1\_1\_motif18\_m \| cluster\_12 \| coexpression\_cluster1\_1 \| 6 \| 6.22 \| 60 \| **dCCCCC** \| **GGGGGH** \| \|; coexpression\_cluster1\_1\_m10\_cluster1\_1\_motif10\_r \| cluster1\_1\_motif10\_r \| cluster\_4 \| coexpression\_cluster1\_1 \| 12 \| 7.57 \| 2229 \| **WWTTTATCAWW+** \| **+wwTgATAAAww** \| \|; coexpression\_cluster1\_1\_m9\_cluster1\_1\_motif9\_r \| cluster1\_1\_motif9\_r \| cluster\_4 \| coexpression\_cluster1\_1 \| 12 \| 9.66 \| 914 \| **wwwTATTAAww+** \| **+WWTTAATAWWW** \| \|; coexpression\_cluster1\_1\_m6\_cluster1\_1\_motif6\_r \| cluster1\_1\_motif6\_r \| cluster\_1 \| coexpression\_cluster1\_1 \| 14 \| 8.87 \| 385 \| **WWTGCATGCATTWW** \| **wwAATGCATGCAww** \| \|; coexpression\_cluster1\_1\_m22\_cluster1\_1\_motif22\_m \| cluster1\_1\_motif22\_m \| cluster\_7 \| coexpression\_cluster1\_1 \| 15 \| 13.58 \| 13 \| **mmTGCmTGArcGRTK** \| **MAYCGYTCAKGCAKK** \| \|; coexpression\_cluster1\_1\_m14\_cluster1\_1\_motif14\_m \| cluster1\_1\_motif14\_m \| cluster\_3 \| coexpression\_cluster1\_1 \| 14 \| 14.20 \| 68 \| **ATACATATATATrT** \| **AYATATATATGTAT** \| \|; coexpression\_cluster1\_1\_m17\_cluster1\_1\_motif17\_m \| cluster1\_1\_motif17\_m \| cluster\_9 \| coexpression\_cluster1\_1 \| 13 \| 14.30 \| 11 \| **WsRGTTCRArTCY** \| **RGAYTYGAACYSW** \| \| - Click on the column names to change the order of the data. - Write the name of one cluster, collection or a pattern in the *Search* window. **Heatmap View** Heatmap View Distance table \| PDF |

**Additional Files**

#### Exported files

| File | Description |
| --- | --- |
| **Pairwise comparison** | This table shows the pairwise comparison between all the input motifs using different metrics. This is the *compare-matrices* result. |
| **Matrix description** | This table shows information of each input motif. |
| **Cluster-wise Motif Files [zip]** | One TRANSFAC file is generated for each cluster including all the input motifs grouped in the corresponding cluster. |
| **Root motifs** | A file with the root matrix of each cluster. |
| **Motif collections per cluster** | A tab separated file containing the counts of motif of each collection on each cluster. |
| **Clusters** | A tab separated file containing the clusters and their correspondng motifs. |
| **Central motifs IDs** | A tab separated file containing the central motif, that one who is the most similar to all the others motif in the same cluster. |
| **Central motifs (in TRANSFAC format)** | A transfac file containing the central motif, that one who is the most similar to all the others motif within the same cluster. |
| **Input motifs** | The input motifs analyzed with matrix-clustering. |

- **References**
  1. Castro-Mondragon JA et al.  RSAT matrix-clustering: dynamic exploration and redundancy reduction of transcription factor binding motif collections. doi: 10.1093/nar/gkx314; Nucleic Acids Research (2017)
  2. **Last RSAT release:** Nguyen NTT et al. RSAT 2018: regulatory sequence analysis tools 20th anniversary. Nucleic Acids Research (2018).
- **Additional Information**
  - Jacques van Helden course
- **Contact**
  - Jaime Castro-Mondragon - ****
  - Morgane Thomas-Chollier - ****
  - Jacques van Helden - ****
