## Additional file 10 for "Transcriptome analysis reveals a novel DNA element that may interact with chromatin-associated proteins in *Plasmodium berghei* during erythrocytic development": matrix-clustering_SUMMARY_cluster1_3.html

matrix-clustering coexpression\_cluster1\_3

### RSAT - matrix-clustering result

##### Analysis: coexpression\_cluster1\_3 (08/09/2023 14:59)

##### Command

```
matrix-clustering  -v 1 -max_matrices 300 -matrix coexpression_cluster1_3 $RSAT/public_html/tmp/www-data/2023/09/08/matrix-clustering_2023-09-08.145929_1pV1Ah/matrix-clustering_query_matrices.transfac transfac -hclust_method average -calc sum -title coexpression_cluster1_3 -metric_build_tree Ncor -lth w 5 -lth cor 0.6 -lth Ncor 0.4 -quick -label_in_tree name -return json,heatmap -o $RSAT/public_html/tmp/www-data/2023/09/08/matrix-clustering_2023-09-08.145929_1pV1Ah/matrix-clustering
```

#### Results Summary

|  |  |  |  |  |  |
| --- | --- | --- | --- | --- | --- |
| Nb Input motifs | Nb Input collections | Nb Clusters Found | Download   Root motifs | Complete results [zip] | Parameters used Linkage method \| Similarity metric \| Thresholds to partition the tree |
| 30 | 1 | 10 | **Download** | Download | average \| Ncor \| **Ncor** = 0.4 **cor** = 0.6 |

|  |  |  |  |  |  |  |
| --- | --- | --- | --- | --- | --- | --- |
|  | **Logo Forest (dynamic browsing)**  **Logo Forest (rapid overview - low image quality)**  cluster\_1  cluster\_2  cluster\_3  cluster\_4  cluster\_5  cluster\_6  cluster\_7  cluster\_8  cluster\_9  cluster\_10 |  |  |  |  |  |
|  | **Clusters Summary** Clusters Summary Table (Root Motif \| Root Motif (Reverse) \| Cluster ID \| # Motifs \| Motif\_Name::Collection \| Number of Motifs by Collection \| Root Motif (transfac Format)) \| | cluster\_1 | 11 | cluster1\_3\_motif10\_r::coexpression\_cluster1\_3:: cluster1\_3\_motif4\_r::coexpression\_cluster1\_3:: cluster1\_3\_motif23\_m::coexpression\_cluster1\_3:: cluster1\_3\_motif1\_r::coexpression\_cluster1\_3:: cluster1\_3\_motif12\_r::coexpression\_cluster1\_3:: cluster1\_3\_motif17\_r::coexpression\_cluster1\_3:: cluster1\_3\_motif15\_r::coexpression\_cluster1\_3:: cluster1\_3\_motif6\_r::coexpression\_cluster1\_3:: cluster1\_3\_motif26\_m::coexpression\_cluster1\_3:: cluster1\_3\_motif21\_m::coexpression\_cluster1\_3:: cluster1\_3\_motif24\_m::coexpression\_cluster1\_3:: | coexpression\_cluster1\_3::11\_motifs | Show matrix |
| cluster\_2 \| 7 \| cluster1\_3\_motif27\_m::coexpression\_cluster1\_3:: cluster1\_3\_motif20\_m::coexpression\_cluster1\_3:: cluster1\_3\_motif25\_m::coexpression\_cluster1\_3:: cluster1\_3\_motif16\_r::coexpression\_cluster1\_3:: cluster1\_3\_motif7\_r::coexpression\_cluster1\_3:: cluster1\_3\_motif2\_r::coexpression\_cluster1\_3:: cluster1\_3\_motif3\_r::coexpression\_cluster1\_3:: \| coexpression\_cluster1\_3::7\_motifs \| Show matrix \| \| |  |  |  |  |  |  |
| cluster\_3 \| 5 \| cluster1\_3\_motif13\_r::coexpression\_cluster1\_3:: cluster1\_3\_motif9\_r::coexpression\_cluster1\_3:: cluster1\_3\_motif14\_r::coexpression\_cluster1\_3:: cluster1\_3\_motif22\_m::coexpression\_cluster1\_3:: cluster1\_3\_motif5\_r::coexpression\_cluster1\_3:: \| coexpression\_cluster1\_3::5\_motifs \| Show matrix \| \| |  |  |  |  |  |  |
| cluster\_4 | 1 | cluster1\_3\_motif19\_r::coexpression\_cluster1\_3 | coexpression\_cluster1\_3 : 1 motif | Show matrix | | |
| cluster\_5 | 1 | cluster1\_3\_motif8\_r::coexpression\_cluster1\_3 | coexpression\_cluster1\_3 : 1 motif | Show matrix | | |
| cluster\_6 | 1 | cluster1\_3\_motif28\_m::coexpression\_cluster1\_3 | coexpression\_cluster1\_3 : 1 motif | Show matrix | | |
| cluster\_7 | 1 | cluster1\_3\_motif11\_r::coexpression\_cluster1\_3 | coexpression\_cluster1\_3 : 1 motif | Show matrix | | |
| cluster\_8 | 1 | cluster1\_3\_motif18\_r::coexpression\_cluster1\_3 | coexpression\_cluster1\_3 : 1 motif | Show matrix | | |
| cluster\_9 | 1 | cluster1\_3\_motif30\_m::coexpression\_cluster1\_3 | coexpression\_cluster1\_3 : 1 motif | Show matrix | | |
| cluster\_10 | 1 | cluster1\_3\_motif29\_m::coexpression\_cluster1\_3 | coexpression\_cluster1\_3 : 1 motif | Show matrix | | |

- Note: by default the table displays the first 50 clusters. Click on the *Show X entries* button to display more (or less) clusters.
- Click on the column names to change the order of the data.
- Use the *Search* window to find your motif (s) of interest.

| **Individual Cluster View** Individual Cluster View **Show All** **Hide All** **cluster\_1** **cluster\_2** **cluster\_3** **cluster\_4** **cluster\_5** **cluster\_6** **cluster\_7** **cluster\_8** **cluster\_9** **cluster\_10** **Display Logo Trees** **cluster\_1** **cluster\_2** **cluster\_3** **cluster\_4** **cluster\_5** **cluster\_6** **cluster\_7** **cluster\_8** **cluster\_9** **cluster\_10** **Display Branch-Motifs** **cluster\_1** Branch Motifs \| \| \| \| \| \| \|; Node \| Consensus \| Logo \| Logo (Reverse) \| Collections \| Matrix (Transfac format) \| IC \| Nb sites; Node 1 \| **wwaAAAAArAmAAAww--** \| \| \| coexpression\_cluster1\_3::2\_motifs \| PSSM \| 10.16 \| 4023; Node 2 \| **wwwAAAAArAcAwwwww-** \| \| \| coexpression\_cluster1\_3::2\_motifs \| PSSM \| 9.55 \| 2079; Node 3 \| **-AAAAAAtAAAAAAAa--** \| \| \| coexpression\_cluster1\_3::2\_motifs \| PSSM \| 12.98 \| 809; Node 4 \| **wwaAAAAArAmAwawww-** \| \| \| coexpression\_cluster1\_3::4\_motifs \| PSSM \| 9.77 \| 6102; Node 5 \| **wwaAAAAArAmAwawwww** \| \| \| coexpression\_cluster1\_3::5\_motifs \| PSSM \| 9.95 \| 6780; Node 6 \| **-AAAAAAtRAAAAAAA--** \| \| \| coexpression\_cluster1\_3::3\_motifs \| PSSM \| 12.59 \| 931; Node 7 \| **wwaAAAAArAmAaaawww** \| \| \| coexpression\_cluster1\_3::8\_motifs \| PSSM \| 9.83 \| 7711; Node 8 \| **---wwwwwTACAACAwww** \| \| \| coexpression\_cluster1\_3::2\_motifs \| PSSM \| 8.65 \| 725; Node 9 \| **wwaAAAAArAmAaaawww** \| \| \| coexpression\_cluster1\_3::9\_motifs \| PSSM \| 9.82 \| 7830; Node 10 \| **wwaAAAAArAmAAaawww** \| \| \| coexpression\_cluster1\_3::11\_motifs \| PSSM \| 9.42 \| 8555 **cluster\_2** Branch Motifs \| \| \| \| \| \| \|; Node \| Consensus \| Logo \| Logo (Reverse) \| Collections \| Matrix (Transfac format) \| IC \| Nb sites; Node 1 \| **---ATATATATrYATATA** \| \| \| coexpression\_cluster1\_3::2\_motifs \| PSSM \| 13.23 \| 707; Node 2 \| **watAyATGCATryAwaww** \| \| \| coexpression\_cluster1\_3::2\_motifs \| PSSM \| 10.53 \| 857; Node 3 \| **--wwtwTrCATGCAwAww** \| \| \| coexpression\_cluster1\_3::2\_motifs \| PSSM \| 9.14 \| 1205; Node 4 \| **wawAyaTrCATGyAwAww** \| \| \| coexpression\_cluster1\_3::4\_motifs \| PSSM \| 9.40 \| 2062; Node 5 \| **wawAtATryATryAwAta** \| \| \| coexpression\_cluster1\_3::6\_motifs \| PSSM \| 9.67 \| 2769; Node 6 \| **wawAtATryATryAwAta** \| \| \| coexpression\_cluster1\_3::7\_motifs \| PSSM \| 9.74 \| 2872 **cluster\_3** Branch Motifs \| \| \| \| \| \| \|; Node \| Consensus \| Logo \| Logo (Reverse) \| Collections \| Matrix (Transfac format) \| IC \| Nb sites; Node 1 \| **AwwAACAAAww---** \| \| \| coexpression\_cluster1\_3::2\_motifs \| PSSM \| 10.34 \| 948; Node 2 \| **-wwAAmAAAATTww** \| \| \| coexpression\_cluster1\_3::2\_motifs \| PSSM \| 9.53 \| 3037; Node 3 \| **AwwAACwmAAww--** \| \| \| coexpression\_cluster1\_3::3\_motifs \| PSSM \| 8.38 \| 1687; Node 4 \| **AwwAAmAAAATTww** \| \| \| coexpression\_cluster1\_3::5\_motifs \| PSSM \| 9.42 \| 4724 **cluster\_4** Branch Motifs \| \| \| \| \| \| \|; Node \| Consensus \| Logo \| Logo (Reverse) \| Collections \| Matrix (Transfac format) \| IC \| Nb sites; Singleton \| **aaAAGAGAaa** \| \| \| coexpression\_cluster1\_3 : 1 motif \| PSSM \| 9.68 \| 122 **cluster\_5** Branch Motifs \| \| \| \| \| \| \|; Node \| Consensus \| Logo \| Logo (Reverse) \| Collections \| Matrix (Transfac format) \| IC \| Nb sites; Singleton \| **wwAAAACAww** \| \| \| coexpression\_cluster1\_3 : 1 motif \| PSSM \| 9.17 \| 821 **cluster\_6** Branch Motifs \| \| \| \| \| \| \|; Node \| Consensus \| Logo \| Logo (Reverse) \| Collections \| Matrix (Transfac format) \| IC \| Nb sites; Singleton \| **bVYkYSSmtkyCCC** \| \| \| coexpression\_cluster1\_3 : 1 motif \| PSSM \| 7.59 \| 66 **cluster\_7** Branch Motifs \| \| \| \| \| \| \|; Node \| Consensus \| Logo \| Logo (Reverse) \| Collections \| Matrix (Transfac format) \| IC \| Nb sites; Singleton \| **wwCAAATGww** \| \| \| coexpression\_cluster1\_3 : 1 motif \| PSSM \| 8.75 \| 205 **cluster\_8** Branch Motifs \| \| \| \| \| \| \|; Node \| Consensus \| Logo \| Logo (Reverse) \| Collections \| Matrix (Transfac format) \| IC \| Nb sites; Singleton \| **wwAACGTCww** \| \| \| coexpression\_cluster1\_3 : 1 motif \| PSSM \| 9.04 \| 31 **cluster\_9** Branch Motifs \| \| \| \| \| \| \|; Node \| Consensus \| Logo \| Logo (Reverse) \| Collections \| Matrix (Transfac format) \| IC \| Nb sites; Singleton \| **aaaCmttGbaaGttG** \| \| \| coexpression\_cluster1\_3 : 1 motif \| PSSM \| 16.37 \| 11 **cluster\_10** Branch Motifs \| \| \| \| \| \| \|; Node \| Consensus \| Logo \| Logo (Reverse) \| Collections \| Matrix (Transfac format) \| IC \| Nb sites; Singleton \| **tGKtmGGGRCayCam** \| \| \| coexpression\_cluster1\_3 : 1 motif \| PSSM \| 16.04 \| 10 - **Each cluster has a different color.** - **Click on the upper buttons to display separately the information of each cluster.** - **Click on the circles at each tree to display its corresponding merged motifs.** Definitions - **Logo tree.** This results shows the hierarchical tree with its logo alignment of one cluster. The logos are shown in Forward (left logo) and Reverse (right logo) orientation. - **Branch-motifs.** On the trees are displayed the branch number that is the number in which the motifs were incorporated in the tree. The Branch-Motifs table shows the logo in both orientations and the link to the file in TRANSFAC format with the branch-motif. |
|  | **Individual Motif View** Individual Motif View Motif id \| Motif name \| Cluster \| Collection \| Width \| IC \| Number of sites \| Consensus \| Consensus (Rev) \| Logo \| Logo (Rev); coexpression\_cluster1\_3\_m28\_cluster1\_3\_motif28\_m \| cluster1\_3\_motif28\_m \| cluster\_6 \| coexpression\_cluster1\_3 \| 14 \| 7.59 \| 66 \| **tmywyscaTtTyCC** \| **GGRAAATGSRWRKA** \| \|; coexpression\_cluster1\_3\_m15\_cluster1\_3\_motif15\_r \| cluster1\_3\_motif15\_r \| cluster\_1 \| coexpression\_cluster1\_3 \| 18 \| 9.36 \| 1342 \| **wwwAAAAArAcAtwww++** \| **++WWWATGTYTTTTTWWW** \| \|; coexpression\_cluster1\_3\_m5\_cluster1\_3\_motif5\_r \| cluster1\_3\_motif5\_r \| cluster\_3 \| coexpression\_cluster1\_3 \| 14 \| 9.26 \| 849 \| **+waAACAAAww+++** \| **+++WWTTTGTTTW+** \| \|; coexpression\_cluster1\_3\_m21\_cluster1\_3\_motif21\_m \| cluster1\_3\_motif21\_m \| cluster\_1 \| coexpression\_cluster1\_3 \| 18 \| 14.60 \| 335 \| **+AAAAAATAAaWAAA+++** \| **+++TTTWTTTATTTTTT+** \| \|; coexpression\_cluster1\_3\_m8\_cluster1\_3\_motif8\_r \| cluster1\_3\_motif8\_r \| cluster\_5 \| coexpression\_cluster1\_3 \| 10 \| 9.17 \| 821 \| **wwAAAACAww** \| **WWTGTTTTWW** \| \|; coexpression\_cluster1\_3\_m24\_cluster1\_3\_motif24\_m \| cluster1\_3\_motif24\_m \| cluster\_1 \| coexpression\_cluster1\_3 \| 18 \| 12.81 \| 474 \| **+AAAAAAdAAAAAAAa++** \| **++TTTTTTTTHTTTTTT+** \| \|; coexpression\_cluster1\_3\_m19\_cluster1\_3\_motif19\_r \| cluster1\_3\_motif19\_r \| cluster\_4 \| coexpression\_cluster1\_3 \| 10 \| 9.68 \| 122 \| **aaAAGAGAaa** \| **TTTCTCTTTT** \| \|; coexpression\_cluster1\_3\_m7\_cluster1\_3\_motif7\_r \| cluster1\_3\_motif7\_r \| cluster\_2 \| coexpression\_cluster1\_3 \| 18 \| 10.56 \| 530 \| **WATAYATGCATRTATW++** \| **++watAyATGCATrTatw** \| \|; coexpression\_cluster1\_3\_m6\_cluster1\_3\_motif6\_r \| cluster1\_3\_motif6\_r \| cluster\_1 \| coexpression\_cluster1\_3 \| 18 \| 9.93 \| 737 \| **wwaaAAAAgAcAwmaww+** \| **+WWTKWTGTCTTTTTTWW** \| \|; coexpression\_cluster1\_3\_m23\_cluster1\_3\_motif23\_m \| cluster1\_3\_motif23\_m \| cluster\_1 \| coexpression\_cluster1\_3 \| 18 \| 14.39 \| 120 \| **TTTTTTTTTTTAATT+++** \| **+++AATTAAAAAAAAAAA** \| \|; coexpression\_cluster1\_3\_m13\_cluster1\_3\_motif13\_r \| cluster1\_3\_motif13\_r \| cluster\_3 \| coexpression\_cluster1\_3 \| 14 \| 9.45 \| 1236 \| **+wwAAcAAAATTww** \| **WWAATTTTGTTWW+** \| \|; coexpression\_cluster1\_3\_m17\_cluster1\_3\_motif17\_r \| cluster1\_3\_motif17\_r \| cluster\_1 \| coexpression\_cluster1\_3 \| 18 \| 10.78 \| 2921 \| **wwaAAAAArAmAAAww++** \| **++WWTTTKTYTTTTTTWW** \| \|; coexpression\_cluster1\_3\_m12\_cluster1\_3\_motif12\_r \| cluster1\_3\_motif12\_r \| cluster\_1 \| coexpression\_cluster1\_3 \| 18 \| 9.42 \| 1102 \| **wwwAAAAArAcAmAww++** \| **++WWTKTGTYTTTTTWWW** \| \|; coexpression\_cluster1\_3\_m25\_cluster1\_3\_motif25\_m \| cluster1\_3\_motif25\_m \| cluster\_2 \| coexpression\_cluster1\_3 \| 18 \| 15.44 \| 182 \| **+++ATATRTATAYATAT+** \| **+ATATRTATAYATAT+++** \| \|; coexpression\_cluster1\_3\_m20\_cluster1\_3\_motif20\_m \| cluster1\_3\_motif20\_m \| cluster\_2 \| coexpression\_cluster1\_3 \| 18 \| 12.68 \| 525 \| **+++ATATAYATRYATRYA** \| **TRYATRyATRTATAT+++** \| \|; coexpression\_cluster1\_3\_m29\_cluster1\_3\_motif29\_m \| cluster1\_3\_motif29\_m \| cluster\_10 \| coexpression\_cluster1\_3 \| 15 \| 16.04 \| 10 \| **TGKTAGGGRCATCAA** \| **TTGATGYCCCTAMCA** \| \|; coexpression\_cluster1\_3\_m27\_cluster1\_3\_motif27\_m \| cluster1\_3\_motif27\_m \| cluster\_2 \| coexpression\_cluster1\_3 \| 18 \| 11.25 \| 103 \| **++++++TATGYATGTA++** \| **++TACATRCATA++++++** \| \|; coexpression\_cluster1\_3\_m9\_cluster1\_3\_motif9\_r \| cluster1\_3\_motif9\_r \| cluster\_3 \| coexpression\_cluster1\_3 \| 14 \| 9.09 \| 1801 \| **++++wwAAAATTww** \| **WWAATTTTWW++++** \| \|; coexpression\_cluster1\_3\_m16\_cluster1\_3\_motif16\_r \| cluster1\_3\_motif16\_r \| cluster\_2 \| coexpression\_cluster1\_3 \| 18 \| 10.89 \| 327 \| **wawAyATGCATrcAwAww** \| **WWTWTGYATGCATRTWTW** \| \|; coexpression\_cluster1\_3\_m2\_cluster1\_3\_motif2\_r \| cluster1\_3\_motif2\_r \| cluster\_2 \| coexpression\_cluster1\_3 \| 18 \| 9.20 \| 655 \| **++++waTACATGCAwAww** \| **WWTWTGCATGTATW++++** \| \|; coexpression\_cluster1\_3\_m22\_cluster1\_3\_motif22\_m \| cluster1\_3\_motif22\_m \| cluster\_3 \| coexpression\_cluster1\_3 \| 14 \| 12.14 \| 99 \| **ATTAACAAAAT+++** \| **+++ATTTTGTTAAT** \| \|; coexpression\_cluster1\_3\_m26\_cluster1\_3\_motif26\_m \| cluster1\_3\_motif26\_m \| cluster\_1 \| coexpression\_cluster1\_3 \| 18 \| 11.65 \| 122 \| **+++TYTTTTTCATTTT++** \| **++AAAAtGAAAAarA+++** \| \|; coexpression\_cluster1\_3\_m3\_cluster1\_3\_motif3\_r \| cluster1\_3\_motif3\_r \| cluster\_2 \| coexpression\_cluster1\_3 \| 18 \| 9.36 \| 550 \| **++wwTwTGCATGCAwAww** \| **WWTWTGCATGCAWAWW++** \| \|; coexpression\_cluster1\_3\_m10\_cluster1\_3\_motif10\_r \| cluster1\_3\_motif10\_r \| cluster\_1 \| coexpression\_cluster1\_3 \| 18 \| 9.71 \| 204 \| **+++wwAwAgACAAcAdww** \| **WWHTGTTGTCTWTWW+++** \| \|; coexpression\_cluster1\_3\_m30\_cluster1\_3\_motif30\_m \| cluster1\_3\_motif30\_m \| cluster\_9 \| coexpression\_cluster1\_3 \| 15 \| 16.37 \| 11 \| **AAACATTGyAAGTTG** \| **CAACTTRCAATGTTT** \| \|; coexpression\_cluster1\_3\_m11\_cluster1\_3\_motif11\_r \| cluster1\_3\_motif11\_r \| cluster\_7 \| coexpression\_cluster1\_3 \| 10 \| 8.75 \| 205 \| **wwCAAATGww** \| **WWCATTTGWW** \| \|; coexpression\_cluster1\_3\_m14\_cluster1\_3\_motif14\_r \| cluster1\_3\_motif14\_r \| cluster\_3 \| coexpression\_cluster1\_3 \| 14 \| 7.24 \| 739 \| **+WWTTGAGTTWW++** \| **++wwAAcTCAAww+** \| \|; coexpression\_cluster1\_3\_m4\_cluster1\_3\_motif4\_r \| cluster1\_3\_motif4\_r \| cluster\_1 \| coexpression\_cluster1\_3 \| 18 \| 8.86 \| 521 \| **+++++wwTTACAACAwww** \| **WWWTGTTGTAAWW+++++** \| \|; coexpression\_cluster1\_3\_m1\_cluster1\_3\_motif1\_r \| cluster1\_3\_motif1\_r \| cluster\_1 \| coexpression\_cluster1\_3 \| 18 \| 10.86 \| 678 \| **WWYTKTTKTYTTTTTWWW** \| **wwwAAAAArAmAAmArww** \| \|; coexpression\_cluster1\_3\_m18\_cluster1\_3\_motif18\_r \| cluster1\_3\_motif18\_r \| cluster\_8 \| coexpression\_cluster1\_3 \| 10 \| 9.04 \| 31 \| **wwAACGTCww** \| **WWGACGTTWW** \| \| - Click on the column names to change the order of the data. - Write the name of one cluster, collection or a pattern in the *Search* window. **Heatmap View** Heatmap View Distance table \| PDF |

**Additional Files**

#### Exported files

| File | Description |
| --- | --- |
| **Pairwise comparison** | This table shows the pairwise comparison between all the input motifs using different metrics. This is the *compare-matrices* result. |
| **Matrix description** | This table shows information of each input motif. |
| **Cluster-wise Motif Files [zip]** | One TRANSFAC file is generated for each cluster including all the input motifs grouped in the corresponding cluster. |
| **Root motifs** | A file with the root matrix of each cluster. |
| **Motif collections per cluster** | A tab separated file containing the counts of motif of each collection on each cluster. |
| **Clusters** | A tab separated file containing the clusters and their correspondng motifs. |
| **Central motifs IDs** | A tab separated file containing the central motif, that one who is the most similar to all the others motif in the same cluster. |
| **Central motifs (in TRANSFAC format)** | A transfac file containing the central motif, that one who is the most similar to all the others motif within the same cluster. |
| **Input motifs** | The input motifs analyzed with matrix-clustering. |

- **References**
  1. Castro-Mondragon JA et al.  RSAT matrix-clustering: dynamic exploration and redundancy reduction of transcription factor binding motif collections. doi: 10.1093/nar/gkx314; Nucleic Acids Research (2017)
  2. **Last RSAT release:** Nguyen NTT et al. RSAT 2018: regulatory sequence analysis tools 20th anniversary. Nucleic Acids Research (2018).
- **Additional Information**
  - Jacques van Helden course
- **Contact**
  - Jaime Castro-Mondragon - ****
  - Morgane Thomas-Chollier - ****
  - Jacques van Helden - ****
