## Additional file 10 for "Transcriptome analysis reveals a novel DNA element that may interact with chromatin-associated proteins in *Plasmodium berghei* during erythrocytic development": matrix-clustering_SUMMARY_cluster1_6.html

matrix-clustering coexpression\_cluster1\_6

### RSAT - matrix-clustering result

##### Analysis: coexpression\_cluster1\_6 (08/09/2023 15:09)

##### Command

```
matrix-clustering  -v 1 -max_matrices 300 -matrix coexpression_cluster1_6 $RSAT/public_html/tmp/www-data/2023/09/08/matrix-clustering_2023-09-08.150848_ydhDNs/matrix-clustering_query_matrices.transfac transfac -hclust_method average -calc sum -title coexpression_cluster1_6 -metric_build_tree Ncor -lth w 5 -lth cor 0.6 -lth Ncor 0.4 -quick -label_in_tree name -return json,heatmap -o $RSAT/public_html/tmp/www-data/2023/09/08/matrix-clustering_2023-09-08.150848_ydhDNs/matrix-clustering
```

#### Results Summary

|  |  |  |  |  |  |
| --- | --- | --- | --- | --- | --- |
| Nb Input motifs | Nb Input collections | Nb Clusters Found | Download   Root motifs | Complete results [zip] | Parameters used Linkage method \| Similarity metric \| Thresholds to partition the tree |
| 32 | 1 | 14 | **Download** | Download | average \| Ncor \| **Ncor** = 0.4 **cor** = 0.6 |

|  |  |  |  |  |  |  |
| --- | --- | --- | --- | --- | --- | --- |
|  | **Logo Forest (dynamic browsing)**  **Logo Forest (rapid overview - low image quality)**  cluster\_1  cluster\_2  cluster\_3  cluster\_4  cluster\_5  cluster\_6  cluster\_7  cluster\_8  cluster\_9  cluster\_10  cluster\_11  cluster\_12  cluster\_13  cluster\_14 |  |  |  |  |  |
|  | **Clusters Summary** Clusters Summary Table (Root Motif \| Root Motif (Reverse) \| Cluster ID \| # Motifs \| Motif\_Name::Collection \| Number of Motifs by Collection \| Root Motif (transfac Format)) \| | cluster\_1 | 7 | cluster1\_6\_motif6\_r::coexpression\_cluster1\_6:: cluster1\_6\_motif2\_r::coexpression\_cluster1\_6:: cluster1\_6\_motif13\_r::coexpression\_cluster1\_6:: cluster1\_6\_motif8\_r::coexpression\_cluster1\_6:: cluster1\_6\_motif25\_m::coexpression\_cluster1\_6:: cluster1\_6\_motif24\_m::coexpression\_cluster1\_6:: cluster1\_6\_motif26\_m::coexpression\_cluster1\_6:: | coexpression\_cluster1\_6::7\_motifs | Show matrix |
| cluster\_2 \| 2 \| cluster1\_6\_motif10\_r::coexpression\_cluster1\_6:: cluster1\_6\_motif3\_r::coexpression\_cluster1\_6:: \| coexpression\_cluster1\_6::2\_motifs \| Show matrix \| \| |  |  |  |  |  |  |
| cluster\_3 \| 2 \| cluster1\_6\_motif20\_m::coexpression\_cluster1\_6:: cluster1\_6\_motif21\_m::coexpression\_cluster1\_6:: \| coexpression\_cluster1\_6::2\_motifs \| Show matrix \| \| |  |  |  |  |  |  |
| cluster\_4 | 2 | cluster1\_6\_motif12\_r::coexpression\_cluster1\_6:: cluster1\_6\_motif18\_r::coexpression\_cluster1\_6:: | coexpression\_cluster1\_6::2\_motifs | Show matrix | | |
| cluster\_5 | 3 | cluster1\_6\_motif14\_r::coexpression\_cluster1\_6:: cluster1\_6\_motif15\_r::coexpression\_cluster1\_6:: cluster1\_6\_motif31\_m::coexpression\_cluster1\_6:: | coexpression\_cluster1\_6::3\_motifs | Show matrix | | |
| cluster\_6 | 5 | cluster1\_6\_motif22\_m::coexpression\_cluster1\_6:: cluster1\_6\_motif1\_r::coexpression\_cluster1\_6:: cluster1\_6\_motif7\_r::coexpression\_cluster1\_6:: cluster1\_6\_motif17\_r::coexpression\_cluster1\_6:: cluster1\_6\_motif27\_m::coexpression\_cluster1\_6:: | coexpression\_cluster1\_6::5\_motifs | Show matrix | | |
| cluster\_7 | 2 | cluster1\_6\_motif19\_r::coexpression\_cluster1\_6:: cluster1\_6\_motif4\_r::coexpression\_cluster1\_6:: | coexpression\_cluster1\_6::2\_motifs | Show matrix | | |
| cluster\_8 | 1 | cluster1\_6\_motif5\_r::coexpression\_cluster1\_6 | coexpression\_cluster1\_6 : 1 motif | Show matrix | | |
| cluster\_9 | 2 | cluster1\_6\_motif23\_m::coexpression\_cluster1\_6:: cluster1\_6\_motif28\_m::coexpression\_cluster1\_6:: | coexpression\_cluster1\_6::2\_motifs | Show matrix | | |
| cluster\_10 | 1 | cluster1\_6\_motif30\_m::coexpression\_cluster1\_6 | coexpression\_cluster1\_6 : 1 motif | Show matrix | | |
| cluster\_11 | 2 | cluster1\_6\_motif11\_r::coexpression\_cluster1\_6:: cluster1\_6\_motif16\_r::coexpression\_cluster1\_6:: | coexpression\_cluster1\_6::2\_motifs | Show matrix | | |
| cluster\_12 | 1 | cluster1\_6\_motif9\_r::coexpression\_cluster1\_6 | coexpression\_cluster1\_6 : 1 motif | Show matrix | | |
| cluster\_13 | 1 | cluster1\_6\_motif29\_m::coexpression\_cluster1\_6 | coexpression\_cluster1\_6 : 1 motif | Show matrix | | |
| cluster\_14 | 1 | cluster1\_6\_motif32\_m::coexpression\_cluster1\_6 | coexpression\_cluster1\_6 : 1 motif | Show matrix | | |

- Note: by default the table displays the first 50 clusters. Click on the *Show X entries* button to display more (or less) clusters.
- Click on the column names to change the order of the data.
- Use the *Search* window to find your motif (s) of interest.

| **Individual Cluster View** Individual Cluster View **Show All** **Hide All** **cluster\_1** **cluster\_2** **cluster\_3** **cluster\_4** **cluster\_5** **cluster\_6** **cluster\_7** **cluster\_8** **cluster\_9** **cluster\_10** **cluster\_11** **cluster\_12** **cluster\_13** **cluster\_14** **Display Logo Trees** **cluster\_1** **cluster\_2** **cluster\_3** **cluster\_4** **cluster\_5** **cluster\_6** **cluster\_7** **cluster\_8** **cluster\_9** **cluster\_10** **cluster\_11** **cluster\_12** **cluster\_13** **cluster\_14** **Display Branch-Motifs** **cluster\_1** Branch Motifs \| \| \| \| \| \| \|; Node \| Consensus \| Logo \| Logo (Reverse) \| Collections \| Matrix (Transfac format) \| IC \| Nb sites; Node 1 \| **-wwAAAAgACAwww--** \| \| \| coexpression\_cluster1\_6::2\_motifs \| PSSM \| 8.46 \| 802; Node 2 \| **-wwAAAAgACAwwww-** \| \| \| coexpression\_cluster1\_6::3\_motifs \| PSSM \| 8.60 \| 1379; Node 3 \| **AAAAAAAAAAAAwAA-** \| \| \| coexpression\_cluster1\_6::2\_motifs \| PSSM \| 12.87 \| 416; Node 4 \| **-wwAAAAgACAawww-** \| \| \| coexpression\_cluster1\_6::4\_motifs \| PSSM \| 8.27 \| 1663; Node 5 \| **AAAAAAAaAAAAwAAA** \| \| \| coexpression\_cluster1\_6::3\_motifs \| PSSM \| 13.43 \| 524; Node 6 \| **AwaAAAArAmAawwaA** \| \| \| coexpression\_cluster1\_6::7\_motifs \| PSSM \| 9.92 \| 2187 **cluster\_2** Branch Motifs \| \| \| \| \| \| \|; Node \| Consensus \| Logo \| Logo (Reverse) \| Collections \| Matrix (Transfac format) \| IC \| Nb sites; Node 1 \| **wwwwAAAAAAATTTTTTTwwww** \| \| \| coexpression\_cluster1\_6::2\_motifs \| PSSM \| 12.62 \| 1479 **cluster\_3** Branch Motifs \| \| \| \| \| \| \|; Node \| Consensus \| Logo \| Logo (Reverse) \| Collections \| Matrix (Transfac format) \| IC \| Nb sites; Node 1 \| **TATATRyATATaTAT** \| \| \| coexpression\_cluster1\_6::2\_motifs \| PSSM \| 12.64 \| 349 **cluster\_4** Branch Motifs \| \| \| \| \| \| \|; Node \| Consensus \| Logo \| Logo (Reverse) \| Collections \| Matrix (Transfac format) \| IC \| Nb sites; Node 1 \| **wwyAACCAwa** \| \| \| coexpression\_cluster1\_6::2\_motifs \| PSSM \| 8.64 \| 109 **cluster\_5** Branch Motifs \| \| \| \| \| \| \|; Node \| Consensus \| Logo \| Logo (Reverse) \| Collections \| Matrix (Transfac format) \| IC \| Nb sites; Node 1 \| **whTAGCTAwwT** \| \| \| coexpression\_cluster1\_6::2\_motifs \| PSSM \| 9.64 \| 202; Node 2 \| **waTAGCyAaww** \| \| \| coexpression\_cluster1\_6::3\_motifs \| PSSM \| 7.31 \| 513 **cluster\_6** Branch Motifs \| \| \| \| \| \| \|; Node \| Consensus \| Logo \| Logo (Reverse) \| Collections \| Matrix (Transfac format) \| IC \| Nb sites; Node 1 \| **wwTTATGAAcAAaww---** \| \| \| coexpression\_cluster1\_6::2\_motifs \| PSSM \| 9.19 \| 717; Node 2 \| **wwTTATGAAcAAawwAkr** \| \| \| coexpression\_cluster1\_6::3\_motifs \| PSSM \| 11.78 \| 776; Node 3 \| **---wwTkAACAaw-----** \| \| \| coexpression\_cluster1\_6::2\_motifs \| PSSM \| 8.08 \| 351; Node 4 \| **wwTTATkAACAAwwwAkr** \| \| \| coexpression\_cluster1\_6::5\_motifs \| PSSM \| 11.60 \| 1127 **cluster\_7** Branch Motifs \| \| \| \| \| \| \|; Node \| Consensus \| Logo \| Logo (Reverse) \| Collections \| Matrix (Transfac format) \| IC \| Nb sites; Node 1 \| **wwTGTTGTAAww** \| \| \| coexpression\_cluster1\_6::2\_motifs \| PSSM \| 8.17 \| 335 **cluster\_8** Branch Motifs \| \| \| \| \| \| \|; Node \| Consensus \| Logo \| Logo (Reverse) \| Collections \| Matrix (Transfac format) \| IC \| Nb sites; Singleton \| **wwACACACAaww** \| \| \| coexpression\_cluster1\_6 : 1 motif \| PSSM \| 8.37 \| 361 **cluster\_9** Branch Motifs \| \| \| \| \| \| \|; Node \| Consensus \| Logo \| Logo (Reverse) \| Collections \| Matrix (Transfac format) \| IC \| Nb sites; Node 1 \| **yystrwTAGCTAkyTrwytatg** \| \| \| coexpression\_cluster1\_6::2\_motifs \| PSSM \| 9.45 \| 109 **cluster\_10** Branch Motifs \| \| \| \| \| \| \|; Node \| Consensus \| Logo \| Logo (Reverse) \| Collections \| Matrix (Transfac format) \| IC \| Nb sites; Singleton \| **aaaaaSGrrra** \| \| \| coexpression\_cluster1\_6 : 1 motif \| PSSM \| 10.37 \| 79 **cluster\_11** Branch Motifs \| \| \| \| \| \| \|; Node \| Consensus \| Logo \| Logo (Reverse) \| Collections \| Matrix (Transfac format) \| IC \| Nb sites; Node 1 \| **wwAATTCAGwwww** \| \| \| coexpression\_cluster1\_6::2\_motifs \| PSSM \| 8.06 \| 469 **cluster\_12** Branch Motifs \| \| \| \| \| \| \|; Node \| Consensus \| Logo \| Logo (Reverse) \| Collections \| Matrix (Transfac format) \| IC \| Nb sites; Singleton \| **wwGAGTTAww** \| \| \| coexpression\_cluster1\_6 : 1 motif \| PSSM \| 9.09 \| 58 **cluster\_13** Branch Motifs \| \| \| \| \| \| \|; Node \| Consensus \| Logo \| Logo (Reverse) \| Collections \| Matrix (Transfac format) \| IC \| Nb sites; Singleton \| **mbSssssyGCaCaCrCvSSSSS** \| \| \| coexpression\_cluster1\_6 : 1 motif \| PSSM \| 11.24 \| 51 **cluster\_14** Branch Motifs \| \| \| \| \| \| \|; Node \| Consensus \| Logo \| Logo (Reverse) \| Collections \| Matrix (Transfac format) \| IC \| Nb sites; Singleton \| **CVCCVC** \| \| \| coexpression\_cluster1\_6 : 1 motif \| PSSM \| 6.35 \| 49 - **Each cluster has a different color.** - **Click on the upper buttons to display separately the information of each cluster.** - **Click on the circles at each tree to display its corresponding merged motifs.** Definitions - **Logo tree.** This results shows the hierarchical tree with its logo alignment of one cluster. The logos are shown in Forward (left logo) and Reverse (right logo) orientation. - **Branch-motifs.** On the trees are displayed the branch number that is the number in which the motifs were incorporated in the tree. The Branch-Motifs table shows the logo in both orientations and the link to the file in TRANSFAC format with the branch-motif. |
|  | **Individual Motif View** Individual Motif View Motif id \| Motif name \| Cluster \| Collection \| Width \| IC \| Number of sites \| Consensus \| Consensus (Rev) \| Logo \| Logo (Rev); coexpression\_cluster1\_6\_m22\_cluster1\_6\_motif22\_m \| cluster1\_6\_motif22\_m \| cluster\_6 \| coexpression\_cluster1\_6 \| 18 \| 12.48 \| 59 \| **+++YMTKRMWTGTTCATA** \| **TATGAACAwkymAkr+++** \| \|; coexpression\_cluster1\_6\_m31\_cluster1\_6\_motif31\_m \| cluster1\_6\_motif31\_m \| cluster\_5 \| coexpression\_cluster1\_6 \| 11 \| 10.50 \| 52 \| **rcTAGCyAWTT** \| **AAWTRGCTAGY** \| \|; coexpression\_cluster1\_6\_m3\_cluster1\_6\_motif3\_r \| cluster1\_6\_motif3\_r \| cluster\_2 \| coexpression\_cluster1\_6 \| 22 \| 12.78 \| 570 \| **wwwmAAAAAAATTTTTTTkwww** \| **WWWMAAAAAAATTTTTTTKWWW** \| \|; coexpression\_cluster1\_6\_m27\_cluster1\_6\_motif27\_m \| cluster1\_6\_motif27\_m \| cluster\_6 \| coexpression\_cluster1\_6 \| 18 \| 7.75 \| 134 \| **+++++TGAACAA++++++** \| **++++++TTGTTCA+++++** \| \|; coexpression\_cluster1\_6\_m24\_cluster1\_6\_motif24\_m \| cluster1\_6\_motif24\_m \| cluster\_1 \| coexpression\_cluster1\_6 \| 16 \| 15.78 \| 107 \| **+AAAAAAAAAAATAA+** \| **+TTATTTTTTTTTTT+** \| \|; coexpression\_cluster1\_6\_m18\_cluster1\_6\_motif18\_r \| cluster1\_6\_motif18\_r \| cluster\_4 \| coexpression\_cluster1\_6 \| 10 \| 9.24 \| 73 \| **wwTAACCAww** \| **WWTGGTTAWW** \| \|; coexpression\_cluster1\_6\_m32\_cluster1\_6\_motif32\_m \| cluster1\_6\_motif32\_m \| cluster\_14 \| coexpression\_cluster1\_6 \| 6 \| 6.35 \| 49 \| **CmCCmC** \| **GKGGKG** \| \|; coexpression\_cluster1\_6\_m12\_cluster1\_6\_motif12\_r \| cluster1\_6\_motif12\_r \| cluster\_4 \| coexpression\_cluster1\_6 \| 10 \| 9.50 \| 36 \| **waCAACCAwa** \| **TWTGGTTGTW** \| \|; coexpression\_cluster1\_6\_m19\_cluster1\_6\_motif19\_r \| cluster1\_6\_motif19\_r \| cluster\_7 \| coexpression\_cluster1\_6 \| 12 \| 8.86 \| 52 \| **+wwGTTGCAww+** \| **+WWTGCAACWW+** \| \|; coexpression\_cluster1\_6\_m11\_cluster1\_6\_motif11\_r \| cluster1\_6\_motif11\_r \| cluster\_11 \| coexpression\_cluster1\_6 \| 13 \| 9.70 \| 59 \| **+++rtTCAGAAww** \| **WWTTCTGAAY+++** \| \|; coexpression\_cluster1\_6\_m9\_cluster1\_6\_motif9\_r \| cluster1\_6\_motif9\_r \| cluster\_12 \| coexpression\_cluster1\_6 \| 10 \| 9.09 \| 58 \| **wwGAGTTAww** \| **WWTAACTCWW** \| \|; coexpression\_cluster1\_6\_m10\_cluster1\_6\_motif10\_r \| cluster1\_6\_motif10\_r \| cluster\_2 \| coexpression\_cluster1\_6 \| 22 \| 12.48 \| 909 \| **WWWWAAAAAAATTTTTTTWWW+** \| **+wwwAAAAAAATTTTTTTwwww** \| \|; coexpression\_cluster1\_6\_m23\_cluster1\_6\_motif23\_m \| cluster1\_6\_motif23\_m \| cluster\_9 \| coexpression\_cluster1\_6 \| 22 \| 11.68 \| 51 \| **+++++wTrGCTAkYTwwkha++** \| **++TDMWWARMTAGCYAW+++++** \| \|; coexpression\_cluster1\_6\_m2\_cluster1\_6\_motif2\_r \| cluster1\_6\_motif2\_r \| cluster\_1 \| coexpression\_cluster1\_6 \| 16 \| 8.89 \| 577 \| **+WWWTTGTCTTTTWW+** \| **+wwaAAAgACAAwww+** \| \|; coexpression\_cluster1\_6\_m28\_cluster1\_6\_motif28\_m \| cluster1\_6\_motif28\_m \| cluster\_9 \| coexpression\_cluster1\_6 \| 22 \| 9.85 \| 58 \| **yystrwyAGCTAGCTvrckmtg** \| **CAKMGYBAGCTAGCTRWYASRR** \| \|; coexpression\_cluster1\_6\_m17\_cluster1\_6\_motif17\_r \| cluster1\_6\_motif17\_r \| cluster\_6 \| coexpression\_cluster1\_6 \| 18 \| 9.22 \| 217 \| **+++WWTTAACAWW+++++** \| **+++++wwTGTTAAww+++** \| \|; coexpression\_cluster1\_6\_m16\_cluster1\_6\_motif16\_r \| cluster1\_6\_motif16\_r \| cluster\_11 \| coexpression\_cluster1\_6 \| 13 \| 7.16 \| 410 \| **wwAATTCAgww++** \| **++WWCTGAATTWW** \| \|; coexpression\_cluster1\_6\_m13\_cluster1\_6\_motif13\_r \| cluster1\_6\_motif13\_r \| cluster\_1 \| coexpression\_cluster1\_6 \| 16 \| 8.57 \| 631 \| **+wwAAAAgACAwww++** \| **++WWWTGTCTTTTWW+** \| \|; coexpression\_cluster1\_6\_m6\_cluster1\_6\_motif6\_r \| cluster1\_6\_motif6\_r \| cluster\_1 \| coexpression\_cluster1\_6 \| 16 \| 8.90 \| 284 \| **+WWWTTGTCTATTWW+** \| **+wwAATAgACAAwww+** \| \|; coexpression\_cluster1\_6\_m4\_cluster1\_6\_motif4\_r \| cluster1\_6\_motif4\_r \| cluster\_7 \| coexpression\_cluster1\_6 \| 12 \| 8.77 \| 283 \| **wwTGTTGTAAww** \| **WWTTACAACAWW** \| \|; coexpression\_cluster1\_6\_m25\_cluster1\_6\_motif25\_m \| cluster1\_6\_motif25\_m \| cluster\_1 \| coexpression\_cluster1\_6 \| 16 \| 13.97 \| 108 \| **+++TTTATTTTATTTT** \| **AAAATAAAATAAA+++** \| \|; coexpression\_cluster1\_6\_m8\_cluster1\_6\_motif8\_r \| cluster1\_6\_motif8\_r \| cluster\_1 \| coexpression\_cluster1\_6 \| 16 \| 8.95 \| 171 \| **+wwwAAAgACAcaw++** \| **++WTGTGTCTTTWWW+** \| \|; coexpression\_cluster1\_6\_m26\_cluster1\_6\_motif26\_m \| cluster1\_6\_motif26\_m \| cluster\_1 \| coexpression\_cluster1\_6 \| 16 \| 12.64 \| 309 \| **AAAAAaAAAaAAAAA+** \| **+TTTTTTTTTTTTTTT** \| \|; coexpression\_cluster1\_6\_m21\_cluster1\_6\_motif21\_m \| cluster1\_6\_motif21\_m \| cluster\_3 \| coexpression\_cluster1\_6 \| 15 \| 13.90 \| 99 \| **TrTATATATATAyA+** \| **+TRTATATATATAYA** \| \|; coexpression\_cluster1\_6\_m1\_cluster1\_6\_motif1\_r \| cluster1\_6\_motif1\_r \| cluster\_6 \| coexpression\_cluster1\_6 \| 18 \| 9.56 \| 566 \| **WWTTATGAAMAAAWW+++** \| **+++wwtTTkTTcATAAww** \| \|; coexpression\_cluster1\_6\_m14\_cluster1\_6\_motif14\_r \| cluster1\_6\_motif14\_r \| cluster\_5 \| coexpression\_cluster1\_6 \| 11 \| 7.27 \| 311 \| **WWTTGGCTATW** \| **waTAGCCAAww** \| \|; coexpression\_cluster1\_6\_m15\_cluster1\_6\_motif15\_r \| cluster1\_6\_motif15\_r \| cluster\_5 \| coexpression\_cluster1\_6 \| 11 \| 9.27 \| 150 \| **WWTAGCTAWW+** \| **+wwTAGCTAww** \| \|; coexpression\_cluster1\_6\_m20\_cluster1\_6\_motif20\_m \| cluster1\_6\_motif20\_m \| cluster\_3 \| coexpression\_cluster1\_6 \| 15 \| 12.52 \| 250 \| **TATATryATATrTrT** \| **AYAYATATRYATATA** \| \|; coexpression\_cluster1\_6\_m5\_cluster1\_6\_motif5\_r \| cluster1\_6\_motif5\_r \| cluster\_8 \| coexpression\_cluster1\_6 \| 12 \| 8.37 \| 361 \| **wwACACACAaww** \| **WWTTGTGTGTWW** \| \|; coexpression\_cluster1\_6\_m7\_cluster1\_6\_motif7\_r \| cluster1\_6\_motif7\_r \| cluster\_6 \| coexpression\_cluster1\_6 \| 18 \| 9.12 \| 151 \| **++wwmTGAACAAwww+++** \| **+++WWWTTGTTCAKWW++** \| \|; coexpression\_cluster1\_6\_m29\_cluster1\_6\_motif29\_m \| cluster1\_6\_motif29\_m \| cluster\_13 \| coexpression\_cluster1\_6 \| 22 \| 11.24 \| 51 \| **wtsawttTGCACACACrymrrg** \| **CYYKRYGTGTGTGCAAAWTSAW** \| \|; coexpression\_cluster1\_6\_m30\_cluster1\_6\_motif30\_m \| cluster1\_6\_motif30\_m \| cluster\_10 \| coexpression\_cluster1\_6 \| 11 \| 10.37 \| 79 \| **AAAAACGAAaA** \| **TTTTCGTTTTT** \| \| - Click on the column names to change the order of the data. - Write the name of one cluster, collection or a pattern in the *Search* window. **Heatmap View** Heatmap View Distance table \| PDF |

**Additional Files**

#### Exported files

| File | Description |
| --- | --- |
| **Pairwise comparison** | This table shows the pairwise comparison between all the input motifs using different metrics. This is the *compare-matrices* result. |
| **Matrix description** | This table shows information of each input motif. |
| **Cluster-wise Motif Files [zip]** | One TRANSFAC file is generated for each cluster including all the input motifs grouped in the corresponding cluster. |
| **Root motifs** | A file with the root matrix of each cluster. |
| **Motif collections per cluster** | A tab separated file containing the counts of motif of each collection on each cluster. |
| **Clusters** | A tab separated file containing the clusters and their correspondng motifs. |
| **Central motifs IDs** | A tab separated file containing the central motif, that one who is the most similar to all the others motif in the same cluster. |
| **Central motifs (in TRANSFAC format)** | A transfac file containing the central motif, that one who is the most similar to all the others motif within the same cluster. |
| **Input motifs** | The input motifs analyzed with matrix-clustering. |

- **References**
  1. Castro-Mondragon JA et al.  RSAT matrix-clustering: dynamic exploration and redundancy reduction of transcription factor binding motif collections. doi: 10.1093/nar/gkx314; Nucleic Acids Research (2017)
  2. **Last RSAT release:** Nguyen NTT et al. RSAT 2018: regulatory sequence analysis tools 20th anniversary. Nucleic Acids Research (2018).
- **Additional Information**
  - Jacques van Helden course
- **Contact**
  - Jaime Castro-Mondragon - ****
  - Morgane Thomas-Chollier - ****
  - Jacques van Helden - ****
