## Additional file 10 for "Transcriptome analysis reveals a novel DNA element that may interact with chromatin-associated proteins in *Plasmodium berghei* during erythrocytic development": matrix-clustering_SUMMARY_cluster1_12.html

matrix-clustering coexpression\_cluster1\_12

### RSAT - matrix-clustering result

##### Analysis: coexpression\_cluster1\_12 (08/09/2023 15:17)

##### Command

```
matrix-clustering  -v 1 -max_matrices 300 -matrix coexpression_cluster1_12 $RSAT/public_html/tmp/www-data/2023/09/08/matrix-clustering_2023-09-08.151652_SAQQkK/matrix-clustering_query_matrices.transfac transfac -hclust_method average -calc sum -title coexpression_cluster1_12 -metric_build_tree Ncor -lth w 5 -lth cor 0.6 -lth Ncor 0.4 -quick -label_in_tree name -return json,heatmap -o $RSAT/public_html/tmp/www-data/2023/09/08/matrix-clustering_2023-09-08.151652_SAQQkK/matrix-clustering
```

#### Results Summary

|  |  |  |  |  |  |
| --- | --- | --- | --- | --- | --- |
| Nb Input motifs | Nb Input collections | Nb Clusters Found | Download   Root motifs | Complete results [zip] | Parameters used Linkage method \| Similarity metric \| Thresholds to partition the tree |
| 29 | 1 | 8 | **Download** | Download | average \| Ncor \| **Ncor** = 0.4 **cor** = 0.6 |

|  |  |  |  |  |  |  |
| --- | --- | --- | --- | --- | --- | --- |
|  | **Logo Forest (dynamic browsing)**  **Logo Forest (rapid overview - low image quality)**  cluster\_1  cluster\_2  cluster\_3  cluster\_4  cluster\_5  cluster\_6  cluster\_7  cluster\_8 |  |  |  |  |  |
|  | **Clusters Summary** Clusters Summary Table (Root Motif \| Root Motif (Reverse) \| Cluster ID \| # Motifs \| Motif\_Name::Collection \| Number of Motifs by Collection \| Root Motif (transfac Format)) \| | cluster\_1 | 14 | cluster1\_12\_motif25\_m::coexpression\_cluster1\_12:: cluster1\_12\_motif19\_m::coexpression\_cluster1\_12:: cluster1\_12\_motif12\_r::coexpression\_cluster1\_12:: cluster1\_12\_motif3\_r::coexpression\_cluster1\_12:: cluster1\_12\_motif8\_r::coexpression\_cluster1\_12:: cluster1\_12\_motif15\_r::coexpression\_cluster1\_12:: cluster1\_12\_motif6\_r::coexpression\_cluster1\_12:: cluster1\_12\_motif4\_r::coexpression\_cluster1\_12:: cluster1\_12\_motif11\_r::coexpression\_cluster1\_12:: cluster1\_12\_motif1\_r::coexpression\_cluster1\_12:: cluster1\_12\_motif10\_r::coexpression\_cluster1\_12:: cluster1\_12\_motif2\_r::coexpression\_cluster1\_12:: cluster1\_12\_motif24\_m::coexpression\_cluster1\_12:: cluster1\_12\_motif7\_r::coexpression\_cluster1\_12:: | coexpression\_cluster1\_12::14\_motifs | Show matrix |
| cluster\_2 \| 8 \| cluster1\_12\_motif13\_r::coexpression\_cluster1\_12:: cluster1\_12\_motif16\_r::coexpression\_cluster1\_12:: cluster1\_12\_motif14\_r::coexpression\_cluster1\_12:: cluster1\_12\_motif9\_r::coexpression\_cluster1\_12:: cluster1\_12\_motif23\_m::coexpression\_cluster1\_12:: cluster1\_12\_motif5\_r::coexpression\_cluster1\_12:: cluster1\_12\_motif17\_m::coexpression\_cluster1\_12:: cluster1\_12\_motif18\_m::coexpression\_cluster1\_12:: \| coexpression\_cluster1\_12::8\_motifs \| Show matrix \| \| |  |  |  |  |  |  |
| cluster\_3 \| 2 \| cluster1\_12\_motif20\_m::coexpression\_cluster1\_12:: cluster1\_12\_motif27\_m::coexpression\_cluster1\_12:: \| coexpression\_cluster1\_12::2\_motifs \| Show matrix \| \| |  |  |  |  |  |  |
| cluster\_4 | 1 | cluster1\_12\_motif21\_m::coexpression\_cluster1\_12 | coexpression\_cluster1\_12 : 1 motif | Show matrix | | |
| cluster\_5 | 1 | cluster1\_12\_motif26\_m::coexpression\_cluster1\_12 | coexpression\_cluster1\_12 : 1 motif | Show matrix | | |
| cluster\_6 | 1 | cluster1\_12\_motif28\_m::coexpression\_cluster1\_12 | coexpression\_cluster1\_12 : 1 motif | Show matrix | | |
| cluster\_7 | 1 | cluster1\_12\_motif29\_m::coexpression\_cluster1\_12 | coexpression\_cluster1\_12 : 1 motif | Show matrix | | |
| cluster\_8 | 1 | cluster1\_12\_motif22\_m::coexpression\_cluster1\_12 | coexpression\_cluster1\_12 : 1 motif | Show matrix | | |

- Note: by default the table displays the first 50 clusters. Click on the *Show X entries* button to display more (or less) clusters.
- Click on the column names to change the order of the data.
- Use the *Search* window to find your motif (s) of interest.

| **Individual Cluster View** Individual Cluster View **Show All** **Hide All** **cluster\_1** **cluster\_2** **cluster\_3** **cluster\_4** **cluster\_5** **cluster\_6** **cluster\_7** **cluster\_8** **Display Logo Trees** **cluster\_1** **cluster\_2** **cluster\_3** **cluster\_4** **cluster\_5** **cluster\_6** **cluster\_7** **cluster\_8** **Display Branch-Motifs** **cluster\_1** Branch Motifs \| \| \| \| \| \| \|; Node \| Consensus \| Logo \| Logo (Reverse) \| Collections \| Matrix (Transfac format) \| IC \| Nb sites; Node 1 \| **wawAAArAmAAAAAAwww-** \| \| \| coexpression\_cluster1\_12::2\_motifs \| PSSM \| 11.09 \| 4358; Node 2 \| **wwAAAArAmAAAAAAwww-** \| \| \| coexpression\_cluster1\_12::3\_motifs \| PSSM \| 10.99 \| 5938; Node 3 \| **wwwwTArAmAAAAAAwww-** \| \| \| coexpression\_cluster1\_12::2\_motifs \| PSSM \| 10.23 \| 2447; Node 4 \| **----waAAAArAmATwww-** \| \| \| coexpression\_cluster1\_12::2\_motifs \| PSSM \| 9.12 \| 2560; Node 5 \| **----waAAAArAmAAwwww** \| \| \| coexpression\_cluster1\_12::2\_motifs \| PSSM \| 9.41 \| 1984; Node 6 \| **wwwAWArAmAAAAAAwww-** \| \| \| coexpression\_cluster1\_12::5\_motifs \| PSSM \| 10.38 \| 8385; Node 7 \| **----waAAAArAmAAwwww** \| \| \| coexpression\_cluster1\_12::3\_motifs \| PSSM \| 9.20 \| 2691; Node 8 \| **----waAAAArAmAwwwww** \| \| \| coexpression\_cluster1\_12::5\_motifs \| PSSM \| 8.89 \| 5251; Node 9 \| **---wwAAAmAAAAAAwww-** \| \| \| coexpression\_cluster1\_12::2\_motifs \| PSSM \| 9.72 \| 4001; Node 10 \| **wwwawAaAmAAAAAAwww-** \| \| \| coexpression\_cluster1\_12::7\_motifs \| PSSM \| 10.22 \| 12386; Node 11 \| **wwwawAaAmAAAAAawwww** \| \| \| coexpression\_cluster1\_12::12\_motifs \| PSSM \| 9.65 \| 17637; Node 12 \| **wwwawAaAmAAAAAawwww** \| \| \| coexpression\_cluster1\_12::13\_motifs \| PSSM \| 9.65 \| 17726; Node 13 \| **wwwawAaAmAAAAAawwww** \| \| \| coexpression\_cluster1\_12::14\_motifs \| PSSM \| 9.64 \| 17836 **cluster\_2** Branch Motifs \| \| \| \| \| \| \|; Node \| Consensus \| Logo \| Logo (Reverse) \| Collections \| Matrix (Transfac format) \| IC \| Nb sites; Node 1 \| **awATRCACAcAtaw---** \| \| \| coexpression\_cluster1\_12::2\_motifs \| PSSM \| 9.03 \| 901; Node 2 \| **-tATrTATryATATATd** \| \| \| coexpression\_cluster1\_12::2\_motifs \| PSSM \| 12.56 \| 414; Node 3 \| **awATGCACACATaw---** \| \| \| coexpression\_cluster1\_12::3\_motifs \| PSSM \| 9.04 \| 1125; Node 4 \| **-taTATrTaTATATaTd** \| \| \| coexpression\_cluster1\_12::3\_motifs \| PSSM \| 11.56 \| 2273; Node 5 \| **-taTATrTaTATATaTd** \| \| \| coexpression\_cluster1\_12::4\_motifs \| PSSM \| 11.32 \| 2372; Node 6 \| **atATryryAyATAtaTd** \| \| \| coexpression\_cluster1\_12::7\_motifs \| PSSM \| 10.00 \| 3497; Node 7 \| **ataTATrTrTATAtaTd** \| \| \| coexpression\_cluster1\_12::8\_motifs \| PSSM \| 9.32 \| 5019 **cluster\_3** Branch Motifs \| \| \| \| \| \| \|; Node \| Consensus \| Logo \| Logo (Reverse) \| Collections \| Matrix (Transfac format) \| IC \| Nb sites; Node 1 \| **wwywwtkyrCACACACrhvgrk** \| \| \| coexpression\_cluster1\_12::2\_motifs \| PSSM \| 10.37 \| 136 **cluster\_4** Branch Motifs \| \| \| \| \| \| \|; Node \| Consensus \| Logo \| Logo (Reverse) \| Collections \| Matrix (Transfac format) \| IC \| Nb sites; Singleton \| **aGMtaGCYa** \| \| \| coexpression\_cluster1\_12 : 1 motif \| PSSM \| 8.17 \| 46 **cluster\_5** Branch Motifs \| \| \| \| \| \| \|; Node \| Consensus \| Logo \| Logo (Reverse) \| Collections \| Matrix (Transfac format) \| IC \| Nb sites; Singleton \| **SSSbSsBaGCtaGCtSVCSsbS** \| \| \| coexpression\_cluster1\_12 : 1 motif \| PSSM \| 9.86 \| 59 **cluster\_6** Branch Motifs \| \| \| \| \| \| \|; Node \| Consensus \| Logo \| Logo (Reverse) \| Collections \| Matrix (Transfac format) \| IC \| Nb sites; Singleton \| **GBGkGSGkGdG** \| \| \| coexpression\_cluster1\_12 : 1 motif \| PSSM \| 7.73 \| 34 **cluster\_7** Branch Motifs \| \| \| \| \| \| \|; Node \| Consensus \| Logo \| Logo (Reverse) \| Collections \| Matrix (Transfac format) \| IC \| Nb sites; Singleton \| **SsssvsswyraGCCSrGvvssG** \| \| \| coexpression\_cluster1\_12 : 1 motif \| PSSM \| 7.56 \| 99 **cluster\_8** Branch Motifs \| \| \| \| \| \| \|; Node \| Consensus \| Logo \| Logo (Reverse) \| Collections \| Matrix (Transfac format) \| IC \| Nb sites; Singleton \| **aaatCGGaa** \| \| \| coexpression\_cluster1\_12 : 1 motif \| PSSM \| 8.61 \| 110 - **Each cluster has a different color.** - **Click on the upper buttons to display separately the information of each cluster.** - **Click on the circles at each tree to display its corresponding merged motifs.** Definitions - **Logo tree.** This results shows the hierarchical tree with its logo alignment of one cluster. The logos are shown in Forward (left logo) and Reverse (right logo) orientation. - **Branch-motifs.** On the trees are displayed the branch number that is the number in which the motifs were incorporated in the tree. The Branch-Motifs table shows the logo in both orientations and the link to the file in TRANSFAC format with the branch-motif. |
|  | **Individual Motif View** Individual Motif View Motif id \| Motif name \| Cluster \| Collection \| Width \| IC \| Number of sites \| Consensus \| Consensus (Rev) \| Logo \| Logo (Rev); coexpression\_cluster1\_12\_m5\_cluster1\_12\_motif5\_r \| cluster1\_12\_motif5\_r \| cluster\_2 \| coexpression\_cluster1\_12 \| 17 \| 10.62 \| 1859 \| **+waTATrTaTATAtw++** \| **++WATATATAYATATW+** \| \|; coexpression\_cluster1\_12\_m12\_cluster1\_12\_motif12\_r \| cluster1\_12\_motif12\_r \| cluster\_1 \| coexpression\_cluster1\_12 \| 19 \| 9.19 \| 957 \| **++++WWAAAARACATTWW+** \| **+wwAATgTyTTTTww++++** \| \|; coexpression\_cluster1\_12\_m14\_cluster1\_12\_motif14\_r \| cluster1\_12\_motif14\_r \| cluster\_2 \| coexpression\_cluster1\_12 \| 17 \| 9.44 \| 320 \| **awATrCACACAyaw+++** \| **+++WTRTGTGTGYATWT** \| \|; coexpression\_cluster1\_12\_m6\_cluster1\_12\_motif6\_r \| cluster1\_12\_motif6\_r \| cluster\_1 \| coexpression\_cluster1\_12 \| 19 \| 9.59 \| 1326 \| **++++waAAAArAmAATww+** \| **+WWATTKTYTTTTTW++++** \| \|; coexpression\_cluster1\_12\_m28\_cluster1\_12\_motif28\_m \| cluster1\_12\_motif28\_m \| cluster\_6 \| coexpression\_cluster1\_12 \| 11 \| 7.73 \| 34 \| **GkrkGsrtGwG** \| **CWCAYSCMYMC** \| \|; coexpression\_cluster1\_12\_m16\_cluster1\_12\_motif16\_r \| cluster1\_12\_motif16\_r \| cluster\_2 \| coexpression\_cluster1\_12 \| 17 \| 9.59 \| 224 \| **+WTATGTGTGCAWW+++** \| **+++wwTGCACACATaw+** \| \|; coexpression\_cluster1\_12\_m24\_cluster1\_12\_motif24\_m \| cluster1\_12\_motif24\_m \| cluster\_1 \| coexpression\_cluster1\_12 \| 19 \| 12.56 \| 265 \| **+++AAAAARAmrAAAAAA+** \| **+TTTTTTYKTYTTTTT+++** \| \|; coexpression\_cluster1\_12\_m7\_cluster1\_12\_motif7\_r \| cluster1\_12\_motif7\_r \| cluster\_1 \| coexpression\_cluster1\_12 \| 19 \| 9.98 \| 3736 \| **+++wwAAAmAAAAAAwww+** \| **+WWWTTTTTTKTTTWW+++** \| \|; coexpression\_cluster1\_12\_m15\_cluster1\_12\_motif15\_r \| cluster1\_12\_motif15\_r \| cluster\_1 \| coexpression\_cluster1\_12 \| 19 \| 9.53 \| 658 \| **++++wawAAAgACAAawww** \| **WWWTTTGTCTTTWTW++++** \| \|; coexpression\_cluster1\_12\_m27\_cluster1\_12\_motif27\_m \| cluster1\_12\_motif27\_m \| cluster\_3 \| coexpression\_cluster1\_12 \| 22 \| 12.34 \| 64 \| **TrtwwtgCRCACACACrwrkmw** \| **WKMYWYGTGTGTGYGCAWWAYA** \| \|; coexpression\_cluster1\_12\_m13\_cluster1\_12\_motif13\_r \| cluster1\_12\_motif13\_r \| cluster\_2 \| coexpression\_cluster1\_12 \| 17 \| 7.54 \| 1522 \| **++WWATGTGTAWW++++** \| **++++wwTACACATww++** \| \|; coexpression\_cluster1\_12\_m4\_cluster1\_12\_motif4\_r \| cluster1\_12\_motif4\_r \| cluster\_1 \| coexpression\_cluster1\_12 \| 19 \| 11.20 \| 1580 \| **WWWTTTTTTKTYTTWTWW+** \| **+wwAwAArAmAAAAAAwww** \| \|; coexpression\_cluster1\_12\_m8\_cluster1\_12\_motif8\_r \| cluster1\_12\_motif8\_r \| cluster\_1 \| coexpression\_cluster1\_12 \| 19 \| 8.89 \| 707 \| **++++WWWTGTCTTTTTW++** \| **++waAAAAgACAwww++++** \| \|; coexpression\_cluster1\_12\_m20\_cluster1\_12\_motif20\_m \| cluster1\_12\_motif20\_m \| cluster\_3 \| coexpression\_cluster1\_12 \| 22 \| 11.05 \| 72 \| **wtswwttTGCACACACrhmvrg** \| **CYBKDYGTGTGTGCAAAWWSAW** \| \|; coexpression\_cluster1\_12\_m29\_cluster1\_12\_motif29\_m \| cluster1\_12\_motif29\_m \| cluster\_7 \| coexpression\_cluster1\_12 \| 22 \| 7.56 \| 99 \| **rwawwwwwTAAGCCcArawwar** \| **YTWWTYTGGGCTTAWWWWWTWY** \| \|; coexpression\_cluster1\_12\_m9\_cluster1\_12\_motif9\_r \| cluster1\_12\_motif9\_r \| cluster\_2 \| coexpression\_cluster1\_12 \| 17 \| 9.21 \| 581 \| **awATGCAcAyATww+++** \| **+++WWATRTGTGCATWT** \| \|; coexpression\_cluster1\_12\_m11\_cluster1\_12\_motif11\_r \| cluster1\_12\_motif11\_r \| cluster\_1 \| coexpression\_cluster1\_12 \| 19 \| 11.10 \| 1932 \| **WWWAAARAMAAAAAAWWW+** \| **+wwwTTTTTTkTyTTTwww** \| \|; coexpression\_cluster1\_12\_m23\_cluster1\_12\_motif23\_m \| cluster1\_12\_motif23\_m \| cluster\_2 \| coexpression\_cluster1\_12 \| 17 \| 13.63 \| 99 \| **++ATrCAYRYATATAT+** \| **+ATATATRYRTGYAT++** \| \|; coexpression\_cluster1\_12\_m10\_cluster1\_12\_motif10\_r \| cluster1\_12\_motif10\_r \| cluster\_1 \| coexpression\_cluster1\_12 \| 19 \| 10.13 \| 1149 \| **+WWWTARAMAAAAAAWWW+** \| **+wwwTTTTTTkTyTAwww+** \| \|; coexpression\_cluster1\_12\_m19\_cluster1\_12\_motif19\_m \| cluster1\_12\_motif19\_m \| cluster\_1 \| coexpression\_cluster1\_12 \| 19 \| 14.06 \| 89 \| **+++++TTATTAWTTTTTT+** \| **+AAAaAAwTAATAA+++++** \| \|; coexpression\_cluster1\_12\_m3\_cluster1\_12\_motif3\_r \| cluster1\_12\_motif3\_r \| cluster\_1 \| coexpression\_cluster1\_12 \| 19 \| 9.72 \| 1603 \| **++++waAAAArAhATAww+** \| **+WWTATDTYTTTTTW++++** \| \|; coexpression\_cluster1\_12\_m2\_cluster1\_12\_motif2\_r \| cluster1\_12\_motif2\_r \| cluster\_1 \| coexpression\_cluster1\_12 \| 19 \| 10.39 \| 1298 \| **wwwATAdAmAAAAAAwww+** \| **+WWWTTTTTTKTHTATWWW** \| \|; coexpression\_cluster1\_12\_m25\_cluster1\_12\_motif25\_m \| cluster1\_12\_motif25\_m \| cluster\_1 \| coexpression\_cluster1\_12 \| 19 \| 12.81 \| 110 \| **+++CWTTTTTTTTAATT++** \| **++AATTAAAAaAAAwg+++** \| \|; coexpression\_cluster1\_12\_m26\_cluster1\_12\_motif26\_m \| cluster1\_12\_motif26\_m \| cluster\_5 \| coexpression\_cluster1\_12 \| 22 \| 9.86 \| 59 \| **yygtrwyAGCTAGCTvrckmtg** \| **CAKMGYBAGCTAGCTRWYACRR** \| \|; coexpression\_cluster1\_12\_m21\_cluster1\_12\_motif21\_m \| cluster1\_12\_motif21\_m \| cluster\_4 \| coexpression\_cluster1\_12 \| 9 \| 8.17 \| 46 \| **AGmTAGcyA** \| **TRGCTAKCT** \| \|; coexpression\_cluster1\_12\_m22\_cluster1\_12\_motif22\_m \| cluster1\_12\_motif22\_m \| cluster\_8 \| coexpression\_cluster1\_12 \| 9 \| 8.61 \| 110 \| **AAATCGGAA** \| **TTCCGATTT** \| \|; coexpression\_cluster1\_12\_m1\_cluster1\_12\_motif1\_r \| cluster1\_12\_motif1\_r \| cluster\_1 \| coexpression\_cluster1\_12 \| 19 \| 11.26 \| 2426 \| **waAAAArAmAAAAAAwww+** \| **+WWWTTTTTTKTYTTTTTW** \| \|; coexpression\_cluster1\_12\_m18\_cluster1\_12\_motif18\_m \| cluster1\_12\_motif18\_m \| cluster\_2 \| coexpression\_cluster1\_12 \| 17 \| 13.36 \| 79 \| **+HATATATRYATATATD** \| **hATATATryATATATd+** \| \|; coexpression\_cluster1\_12\_m17\_cluster1\_12\_motif17\_m \| cluster1\_12\_motif17\_m \| cluster\_2 \| coexpression\_cluster1\_12 \| 17 \| 12.84 \| 335 \| **+TATRTATRYATATAT+** \| **+ATATATRyATAyATA+** \| \| - Click on the column names to change the order of the data. - Write the name of one cluster, collection or a pattern in the *Search* window. **Heatmap View** Heatmap View Distance table \| PDF |

**Additional Files**

#### Exported files

| File | Description |
| --- | --- |
| **Pairwise comparison** | This table shows the pairwise comparison between all the input motifs using different metrics. This is the *compare-matrices* result. |
| **Matrix description** | This table shows information of each input motif. |
| **Cluster-wise Motif Files [zip]** | One TRANSFAC file is generated for each cluster including all the input motifs grouped in the corresponding cluster. |
| **Root motifs** | A file with the root matrix of each cluster. |
| **Motif collections per cluster** | A tab separated file containing the counts of motif of each collection on each cluster. |
| **Clusters** | A tab separated file containing the clusters and their correspondng motifs. |
| **Central motifs IDs** | A tab separated file containing the central motif, that one who is the most similar to all the others motif in the same cluster. |
| **Central motifs (in TRANSFAC format)** | A transfac file containing the central motif, that one who is the most similar to all the others motif within the same cluster. |
| **Input motifs** | The input motifs analyzed with matrix-clustering. |

- **References**
  1. Castro-Mondragon JA et al.  RSAT matrix-clustering: dynamic exploration and redundancy reduction of transcription factor binding motif collections. doi: 10.1093/nar/gkx314; Nucleic Acids Research (2017)
  2. **Last RSAT release:** Nguyen NTT et al. RSAT 2018: regulatory sequence analysis tools 20th anniversary. Nucleic Acids Research (2018).
- **Additional Information**
  - Jacques van Helden course
- **Contact**
  - Jaime Castro-Mondragon - ****
  - Morgane Thomas-Chollier - ****
  - Jacques van Helden - ****
