## Additional file 10 for "Transcriptome analysis reveals a novel DNA element that may interact with chromatin-associated proteins in *Plasmodium berghei* during erythrocytic development": matrix-clustering_SUMMARY_cluster2_2.html

matrix-clustering coexpression\_cluster2\_2

### RSAT - matrix-clustering result

##### Analysis: coexpression\_cluster2\_2 (08/09/2023 15:27)

##### Command

```
matrix-clustering  -v 1 -max_matrices 300 -matrix coexpression_cluster2_2 $RSAT/public_html/tmp/www-data/2023/09/08/matrix-clustering_2023-09-08.152703_KrMovj/matrix-clustering_query_matrices.transfac transfac -hclust_method average -calc sum -title coexpression_cluster2_2 -metric_build_tree Ncor -lth w 5 -lth cor 0.6 -lth Ncor 0.4 -quick -label_in_tree name -return json,heatmap -o $RSAT/public_html/tmp/www-data/2023/09/08/matrix-clustering_2023-09-08.152703_KrMovj/matrix-clustering
```

#### Results Summary

|  |  |  |  |  |  |
| --- | --- | --- | --- | --- | --- |
| Nb Input motifs | Nb Input collections | Nb Clusters Found | Download   Root motifs | Complete results [zip] | Parameters used Linkage method \| Similarity metric \| Thresholds to partition the tree |
| 38 | 1 | 17 | **Download** | Download | average \| Ncor \| **Ncor** = 0.4 **cor** = 0.6 |

|  |  |  |  |  |  |  |
| --- | --- | --- | --- | --- | --- | --- |
|  | **Logo Forest (dynamic browsing)**  **Logo Forest (rapid overview - low image quality)**  cluster\_1  cluster\_2  cluster\_3  cluster\_4  cluster\_5  cluster\_6  cluster\_7  cluster\_8  cluster\_9  cluster\_10  cluster\_11  cluster\_12  cluster\_13  cluster\_14  cluster\_15  cluster\_16  cluster\_17 |  |  |  |  |  |
|  | **Clusters Summary** Clusters Summary Table (Root Motif \| Root Motif (Reverse) \| Cluster ID \| # Motifs \| Motif\_Name::Collection \| Number of Motifs by Collection \| Root Motif (transfac Format)) \| | cluster\_1 | 13 | cluster2\_2\_motif14\_r::coexpression\_cluster2\_2:: cluster2\_2\_motif1\_r::coexpression\_cluster2\_2:: cluster2\_2\_motif6\_r::coexpression\_cluster2\_2:: cluster2\_2\_motif10\_r::coexpression\_cluster2\_2:: cluster2\_2\_motif9\_r::coexpression\_cluster2\_2:: cluster2\_2\_motif8\_r::coexpression\_cluster2\_2:: cluster2\_2\_motif13\_r::coexpression\_cluster2\_2:: cluster2\_2\_motif5\_r::coexpression\_cluster2\_2:: cluster2\_2\_motif12\_r::coexpression\_cluster2\_2:: cluster2\_2\_motif16\_r::coexpression\_cluster2\_2:: cluster2\_2\_motif7\_r::coexpression\_cluster2\_2:: cluster2\_2\_motif15\_r::coexpression\_cluster2\_2:: cluster2\_2\_motif2\_r::coexpression\_cluster2\_2:: | coexpression\_cluster2\_2::13\_motifs | Show matrix |
| cluster\_2 \| 5 \| cluster2\_2\_motif18\_r::coexpression\_cluster2\_2:: cluster2\_2\_motif19\_r::coexpression\_cluster2\_2:: cluster2\_2\_motif4\_r::coexpression\_cluster2\_2:: cluster2\_2\_motif11\_r::coexpression\_cluster2\_2:: cluster2\_2\_motif3\_r::coexpression\_cluster2\_2:: \| coexpression\_cluster2\_2::5\_motifs \| Show matrix \| \| |  |  |  |  |  |  |
| cluster\_3 \| 2 \| cluster2\_2\_motif22\_m::coexpression\_cluster2\_2:: cluster2\_2\_motif23\_m::coexpression\_cluster2\_2:: \| coexpression\_cluster2\_2::2\_motifs \| Show matrix \| \| |  |  |  |  |  |  |
| cluster\_4 | 3 | cluster2\_2\_motif37\_m::coexpression\_cluster2\_2:: cluster2\_2\_motif21\_m::coexpression\_cluster2\_2:: cluster2\_2\_motif32\_m::coexpression\_cluster2\_2:: | coexpression\_cluster2\_2::3\_motifs | Show matrix | | |
| cluster\_5 | 2 | cluster2\_2\_motif17\_r::coexpression\_cluster2\_2:: cluster2\_2\_motif36\_m::coexpression\_cluster2\_2:: | coexpression\_cluster2\_2::2\_motifs | Show matrix | | |
| cluster\_6 | 1 | cluster2\_2\_motif25\_m::coexpression\_cluster2\_2 | coexpression\_cluster2\_2 : 1 motif | Show matrix | | |
| cluster\_7 | 1 | cluster2\_2\_motif33\_m::coexpression\_cluster2\_2 | coexpression\_cluster2\_2 : 1 motif | Show matrix | | |
| cluster\_8 | 1 | cluster2\_2\_motif28\_m::coexpression\_cluster2\_2 | coexpression\_cluster2\_2 : 1 motif | Show matrix | | |
| cluster\_9 | 1 | cluster2\_2\_motif31\_m::coexpression\_cluster2\_2 | coexpression\_cluster2\_2 : 1 motif | Show matrix | | |
| cluster\_10 | 2 | cluster2\_2\_motif20\_m::coexpression\_cluster2\_2:: cluster2\_2\_motif29\_m::coexpression\_cluster2\_2:: | coexpression\_cluster2\_2::2\_motifs | Show matrix | | |
| cluster\_11 | 1 | cluster2\_2\_motif30\_m::coexpression\_cluster2\_2 | coexpression\_cluster2\_2 : 1 motif | Show matrix | | |
| cluster\_12 | 1 | cluster2\_2\_motif26\_m::coexpression\_cluster2\_2 | coexpression\_cluster2\_2 : 1 motif | Show matrix | | |
| cluster\_13 | 1 | cluster2\_2\_motif24\_m::coexpression\_cluster2\_2 | coexpression\_cluster2\_2 : 1 motif | Show matrix | | |
| cluster\_14 | 1 | cluster2\_2\_motif38\_m::coexpression\_cluster2\_2 | coexpression\_cluster2\_2 : 1 motif | Show matrix | | |
| cluster\_15 | 1 | cluster2\_2\_motif34\_m::coexpression\_cluster2\_2 | coexpression\_cluster2\_2 : 1 motif | Show matrix | | |
| cluster\_16 | 1 | cluster2\_2\_motif35\_m::coexpression\_cluster2\_2 | coexpression\_cluster2\_2 : 1 motif | Show matrix | | |
| cluster\_17 | 1 | cluster2\_2\_motif27\_m::coexpression\_cluster2\_2 | coexpression\_cluster2\_2 : 1 motif | Show matrix | | |

- Note: by default the table displays the first 50 clusters. Click on the *Show X entries* button to display more (or less) clusters.
- Click on the column names to change the order of the data.
- Use the *Search* window to find your motif (s) of interest.

| **Individual Cluster View** Individual Cluster View **Show All** **Hide All** **cluster\_1** **cluster\_2** **cluster\_3** **cluster\_4** **cluster\_5** **cluster\_6** **cluster\_7** **cluster\_8** **cluster\_9** **cluster\_10** **cluster\_11** **cluster\_12** **cluster\_13** **cluster\_14** **cluster\_15** **cluster\_16** **cluster\_17** **Display Logo Trees** **cluster\_1** **cluster\_2** **cluster\_3** **cluster\_4** **cluster\_5** **cluster\_6** **cluster\_7** **cluster\_8** **cluster\_9** **cluster\_10** **cluster\_11** **cluster\_12** **cluster\_13** **cluster\_14** **cluster\_15** **cluster\_16** **cluster\_17** **Display Branch-Motifs** **cluster\_1** Branch Motifs \| \| \| \| \| \| \|; Node \| Consensus \| Logo \| Logo (Reverse) \| Collections \| Matrix (Transfac format) \| IC \| Nb sites; Node 1 \| **-----wwwwwwTAATAATATAawTATAwAwaww-----------** \| \| \| coexpression\_cluster2\_2::2\_motifs \| PSSM \| 14.33 \| 3701; Node 2 \| **--wwwwwwwwwwwAtwATAATAAWwwAATATawAwwww------** \| \| \| coexpression\_cluster2\_2::2\_motifs \| PSSM \| 17.21 \| 2265; Node 3 \| **--wwwwTwTwATAAwTATwATAATAwwAwwawwwww--------** \| \| \| coexpression\_cluster2\_2::2\_motifs \| PSSM \| 16.53 \| 2270; Node 4 \| **--wwwwTwTwATAATTATwATAATAwwwwwawwwww--------** \| \| \| coexpression\_cluster2\_2::3\_motifs \| PSSM \| 16.65 \| 3200; Node 5 \| **wwtwwwwwwwwTwwTwwTwwTAwwwwtwTwwTAATATAwwwwww** \| \| \| coexpression\_cluster2\_2::2\_motifs \| PSSM \| 19.90 \| 1537; Node 6 \| **-----wwwwwwTAATwATAwwAwTwTAwAwwww-----------** \| \| \| coexpression\_cluster2\_2::3\_motifs \| PSSM \| 13.42 \| 5074; Node 7 \| **wwtwwwwwtwwwwwTwwTwwTAwwwwTwTwwwwwwwTAwwwwww** \| \| \| coexpression\_cluster2\_2::3\_motifs \| PSSM \| 19.83 \| 2755; Node 8 \| **--wwwwwwTwwTAATTATwATAATwwwawwwwwawwww------** \| \| \| coexpression\_cluster2\_2::5\_motifs \| PSSM \| 16.57 \| 5465; Node 9 \| **----wwwwTwwwTAwTATwATAwwww------------------** \| \| \| coexpression\_cluster2\_2::2\_motifs \| PSSM \| 11.09 \| 4955; Node 10 \| **--wwwwwwwwwTAATwATwwwAwTwwwwwwwwwawwww------** \| \| \| coexpression\_cluster2\_2::8\_motifs \| PSSM \| 15.58 \| 10539; Node 11 \| **wwwwwwwwwwwTwATwWTwwwAwTwwwwwwwwwawwwwwwwwww** \| \| \| coexpression\_cluster2\_2::11\_motifs \| PSSM \| 18.47 \| 13294; Node 12 \| **wwwwwwwwwwwwwAwwATwwwAwtwwwwwwwwwawwwwwwwwww** \| \| \| coexpression\_cluster2\_2::13\_motifs \| PSSM \| 18.26 \| 18249 **cluster\_2** Branch Motifs \| \| \| \| \| \| \|; Node \| Consensus \| Logo \| Logo (Reverse) \| Collections \| Matrix (Transfac format) \| IC \| Nb sites; Node 1 \| **wwTATmTAAAATww** \| \| \| coexpression\_cluster2\_2::2\_motifs \| PSSM \| 9.42 \| 2202; Node 2 \| **wwwTThTAAAATww** \| \| \| coexpression\_cluster2\_2::2\_motifs \| PSSM \| 8.88 \| 3492; Node 3 \| **wwwwThTAAAATww** \| \| \| coexpression\_cluster2\_2::4\_motifs \| PSSM \| 8.68 \| 5694; Node 4 \| **wwTwTwTAAwATww** \| \| \| coexpression\_cluster2\_2::5\_motifs \| PSSM \| 7.92 \| 8299 **cluster\_3** Branch Motifs \| \| \| \| \| \| \|; Node \| Consensus \| Logo \| Logo (Reverse) \| Collections \| Matrix (Transfac format) \| IC \| Nb sites; Node 1 \| **GSAKAGAwAAgGrAA** \| \| \| coexpression\_cluster2\_2::2\_motifs \| PSSM \| 14.33 \| 68 **cluster\_4** Branch Motifs \| \| \| \| \| \| \|; Node \| Consensus \| Logo \| Logo (Reverse) \| Collections \| Matrix (Transfac format) \| IC \| Nb sites; Node 1 \| **--mCCCAATA** \| \| \| coexpression\_cluster2\_2::2\_motifs \| PSSM \| 7.62 \| 143; Node 2 \| **CrmCCCAATA** \| \| \| coexpression\_cluster2\_2::3\_motifs \| PSSM \| 8.61 \| 181 **cluster\_5** Branch Motifs \| \| \| \| \| \| \|; Node \| Consensus \| Logo \| Logo (Reverse) \| Collections \| Matrix (Transfac format) \| IC \| Nb sites; Node 1 \| **AwwAAACTAwwCGTA** \| \| \| coexpression\_cluster2\_2::2\_motifs \| PSSM \| 15.24 \| 217 **cluster\_6** Branch Motifs \| \| \| \| \| \| \|; Node \| Consensus \| Logo \| Logo (Reverse) \| Collections \| Matrix (Transfac format) \| IC \| Nb sites; Singleton \| **tryrtrYryrYrtaY** \| \| \| coexpression\_cluster2\_2 : 1 motif \| PSSM \| 12.51 \| 179 **cluster\_7** Branch Motifs \| \| \| \| \| \| \|; Node \| Consensus \| Logo \| Logo (Reverse) \| Collections \| Matrix (Transfac format) \| IC \| Nb sites; Singleton \| **taYataSGGYyaaaK** \| \| \| coexpression\_cluster2\_2 : 1 motif \| PSSM \| 16.40 \| 22 **cluster\_8** Branch Motifs \| \| \| \| \| \| \|; Node \| Consensus \| Logo \| Logo (Reverse) \| Collections \| Matrix (Transfac format) \| IC \| Nb sites; Singleton \| **RaaaGatGyGCamaa** \| \| \| coexpression\_cluster2\_2 : 1 motif \| PSSM \| 15.93 \| 18 **cluster\_9** Branch Motifs \| \| \| \| \| \| \|; Node \| Consensus \| Logo \| Logo (Reverse) \| Collections \| Matrix (Transfac format) \| IC \| Nb sites; Singleton \| **GCRCRCVtBSS** \| \| \| coexpression\_cluster2\_2 : 1 motif \| PSSM \| 8.24 \| 20 **cluster\_10** Branch Motifs \| \| \| \| \| \| \|; Node \| Consensus \| Logo \| Logo (Reverse) \| Collections \| Matrix (Transfac format) \| IC \| Nb sites; Node 1 \| **GcATyAwTACTAATT** \| \| \| coexpression\_cluster2\_2::2\_motifs \| PSSM \| 15.26 \| 78 **cluster\_11** Branch Motifs \| \| \| \| \| \| \|; Node \| Consensus \| Logo \| Logo (Reverse) \| Collections \| Matrix (Transfac format) \| IC \| Nb sites; Singleton \| **aGRCaaSRatRYaGC** \| \| \| coexpression\_cluster2\_2 : 1 motif \| PSSM \| 14.32 \| 12 **cluster\_12** Branch Motifs \| \| \| \| \| \| \|; Node \| Consensus \| Logo \| Logo (Reverse) \| Collections \| Matrix (Transfac format) \| IC \| Nb sites; Singleton \| **tGCatyCaaaGRCtt** \| \| \| coexpression\_cluster2\_2 : 1 motif \| PSSM \| 16.73 \| 18 **cluster\_13** Branch Motifs \| \| \| \| \| \| \|; Node \| Consensus \| Logo \| Logo (Reverse) \| Collections \| Matrix (Transfac format) \| IC \| Nb sites; Singleton \| **aatCaCCCaataaGG** \| \| \| coexpression\_cluster2\_2 : 1 motif \| PSSM \| 17.50 \| 13 **cluster\_14** Branch Motifs \| \| \| \| \| \| \|; Node \| Consensus \| Logo \| Logo (Reverse) \| Collections \| Matrix (Transfac format) \| IC \| Nb sites; Singleton \| **CtattaaaG** \| \| \| coexpression\_cluster2\_2 : 1 motif \| PSSM \| 10.22 \| 34 **cluster\_15** Branch Motifs \| \| \| \| \| \| \|; Node \| Consensus \| Logo \| Logo (Reverse) \| Collections \| Matrix (Transfac format) \| IC \| Nb sites; Singleton \| **tGCtCatatGaGtGt** \| \| \| coexpression\_cluster2\_2 : 1 motif \| PSSM \| 20.79 \| 5 **cluster\_16** Branch Motifs \| \| \| \| \| \| \|; Node \| Consensus \| Logo \| Logo (Reverse) \| Collections \| Matrix (Transfac format) \| IC \| Nb sites; Singleton \| **CCCCttaaCCtaGaR** \| \| \| coexpression\_cluster2\_2 : 1 motif \| PSSM \| 18.95 \| 5 **cluster\_17** Branch Motifs \| \| \| \| \| \| \|; Node \| Consensus \| Logo \| Logo (Reverse) \| Collections \| Matrix (Transfac format) \| IC \| Nb sites; Singleton \| **SGktkyaarCCSyGG** \| \| \| coexpression\_cluster2\_2 : 1 motif \| PSSM \| 13.56 \| 25 - **Each cluster has a different color.** - **Click on the upper buttons to display separately the information of each cluster.** - **Click on the circles at each tree to display its corresponding merged motifs.** Definitions - **Logo tree.** This results shows the hierarchical tree with its logo alignment of one cluster. The logos are shown in Forward (left logo) and Reverse (right logo) orientation. - **Branch-motifs.** On the trees are displayed the branch number that is the number in which the motifs were incorporated in the tree. The Branch-Motifs table shows the logo in both orientations and the link to the file in TRANSFAC format with the branch-motif. |
|  | **Individual Motif View** Individual Motif View Motif id \| Motif name \| Cluster \| Collection \| Width \| IC \| Number of sites \| Consensus \| Consensus (Rev) \| Logo \| Logo (Rev); coexpression\_cluster2\_2\_m13\_cluster2\_2\_motif13\_r \| cluster2\_2\_motif13\_r \| cluster\_1 \| coexpression\_cluster2\_2 \| 44 \| 14.30 \| 2135 \| **+++++WWWWWWTAATAATATAWWTATAWAWAWW+++++++++++** \| **+++++++++++wwtwTwTATAwwTATATTATTAwwwwww+++++** \| \|; coexpression\_cluster2\_2\_m8\_cluster2\_2\_motif8\_r \| cluster2\_2\_motif8\_r \| cluster\_1 \| coexpression\_cluster2\_2 \| 44 \| 14.22 \| 1373 \| **+++++WWTWTWTWATTATAATAATWTAWAAWWW+++++++++++** \| **+++++++++++wwwttwtAwATTATTATAATwAwAwAww+++++** \| \|; coexpression\_cluster2\_2\_m36\_cluster2\_2\_motif36\_m \| cluster2\_2\_motif36\_m \| cluster\_5 \| coexpression\_cluster2\_2 \| 15 \| 19.14 \| 11 \| **ARAAAACTATGCGTA** \| **TACGCATAGTTTTYT** \| \|; coexpression\_cluster2\_2\_m29\_cluster2\_2\_motif29\_m \| cluster2\_2\_motif29\_m \| cluster\_10 \| coexpression\_cluster2\_2 \| 15 \| 15.86 \| 29 \| **GcwTCArTrCTwATT** \| **AATWAGYAYTGAWGC** \| \|; coexpression\_cluster2\_2\_m22\_cluster2\_2\_motif22\_m \| cluster2\_2\_motif22\_m \| cluster\_3 \| coexpression\_cluster2\_2 \| 15 \| 10.46 \| 46 \| **++++AGAAAAgGrAA** \| **TTYCCTTTTCT++++** \| \|; coexpression\_cluster2\_2\_m21\_cluster2\_2\_motif21\_m \| cluster2\_2\_motif21\_m \| cluster\_4 \| coexpression\_cluster2\_2 \| 10 \| 8.39 \| 99 \| **++ACCCAATA** \| **TATTGGGT++** \| \|; coexpression\_cluster2\_2\_m14\_cluster2\_2\_motif14\_r \| cluster2\_2\_motif14\_r \| cluster\_1 \| coexpression\_cluster2\_2 \| 44 \| 11.87 \| 3027 \| **++++wwtwTwTATAATATTATAwwww++++++++++++++++++** \| **++++++++++++++++++WWWWTATAATATTATAWAWAWW++++** \| \|; coexpression\_cluster2\_2\_m16\_cluster2\_2\_motif16\_r \| cluster2\_2\_motif16\_r \| cluster\_1 \| coexpression\_cluster2\_2 \| 44 \| 17.45 \| 1165 \| **++WWWWWWTWWWWWTWATAATAAWWWAATATWWAWWWW++++++** \| **++++++wwwwTwwaTATTwwwTTATTATwawwwwwawwwwww++** \| \|; coexpression\_cluster2\_2\_m37\_cluster2\_2\_motif37\_m \| cluster2\_2\_motif37\_m \| cluster\_4 \| coexpression\_cluster2\_2 \| 10 \| 6.63 \| 38 \| **KRGGGKYG++** \| **++CrmCCCym** \| \|; coexpression\_cluster2\_2\_m27\_cluster2\_2\_motif27\_m \| cluster2\_2\_motif27\_m \| cluster\_17 \| coexpression\_cluster2\_2 \| 15 \| 13.56 \| 25 \| **CRTTTTAAACCGTGG** \| **CCACGGTTTAAAAYG** \| \|; coexpression\_cluster2\_2\_m9\_cluster2\_2\_motif9\_r \| cluster2\_2\_motif9\_r \| cluster\_1 \| coexpression\_cluster2\_2 \| 44 \| 19.62 \| 864 \| **+++wwwwtwTATwwwTATTATAwwwwtwTAATaATATAwwwwww** \| **WWWWWWTATATTATTAWAWWWWTATAATAWWWATAWAWWWW+++** \| \|; coexpression\_cluster2\_2\_m34\_cluster2\_2\_motif34\_m \| cluster2\_2\_motif34\_m \| cluster\_15 \| coexpression\_cluster2\_2 \| 15 \| 20.79 \| 5 \| **TGCTCATATGAGTGT** \| **ACACTCATATGAGCA** \| \|; coexpression\_cluster2\_2\_m1\_cluster2\_2\_motif1\_r \| cluster2\_2\_motif1\_r \| cluster\_1 \| coexpression\_cluster2\_2 \| 44 \| 12.13 \| 1928 \| **+++++wwwTwATTATTATAATAATww++++++++++++++++++** \| **++++++++++++++++++WWATTATTATAATAATWAWWW+++++** \| \|; coexpression\_cluster2\_2\_m18\_cluster2\_2\_motif18\_r \| cluster2\_2\_motif18\_r \| cluster\_2 \| coexpression\_cluster2\_2 \| 14 \| 8.81 \| 2605 \| **WWTATTTAATWWW+** \| **+wwwATTAAATAww** \| \|; coexpression\_cluster2\_2\_m26\_cluster2\_2\_motif26\_m \| cluster2\_2\_motif26\_m \| cluster\_12 \| coexpression\_cluster2\_2 \| 15 \| 16.73 \| 18 \| **TRCATTCAAAGrCTT** \| **AAGYCTTTGAATGYA** \| \|; coexpression\_cluster2\_2\_m11\_cluster2\_2\_motif11\_r \| cluster2\_2\_motif11\_r \| cluster\_2 \| coexpression\_cluster2\_2 \| 14 \| 9.15 \| 1801 \| **WWTTTHTAAAATWW** \| **wwATTTTAdAAAww** \| \|; coexpression\_cluster2\_2\_m3\_cluster2\_2\_motif3\_r \| cluster2\_2\_motif3\_r \| cluster\_2 \| coexpression\_cluster2\_2 \| 14 \| 9.33 \| 1691 \| **wwATThTAAAATww** \| **WWATTTTADAATWW** \| \|; coexpression\_cluster2\_2\_m5\_cluster2\_2\_motif5\_r \| cluster2\_2\_motif5\_r \| cluster\_1 \| coexpression\_cluster2\_2 \| 44 \| 14.50 \| 1566 \| **+++++WWWWWWTAATAATATAAWTATAWAWAWW+++++++++++** \| **+++++++++++wwtwTwTATAwTTATATTATTAwwwwww+++++** \| \|; coexpression\_cluster2\_2\_m19\_cluster2\_2\_motif19\_r \| cluster2\_2\_motif19\_r \| cluster\_2 \| coexpression\_cluster2\_2 \| 14 \| 10.55 \| 403 \| **wwwATmTAAAATww** \| **WWATTTTAKATWWW** \| \|; coexpression\_cluster2\_2\_m4\_cluster2\_2\_motif4\_r \| cluster2\_2\_motif4\_r \| cluster\_2 \| coexpression\_cluster2\_2 \| 14 \| 9.36 \| 1799 \| **wwTAThTAAAATww** \| **WWATTTTADATAWW** \| \|; coexpression\_cluster2\_2\_m32\_cluster2\_2\_motif32\_m \| cluster2\_2\_motif32\_m \| cluster\_4 \| coexpression\_cluster2\_2 \| 10 \| 6.48 \| 44 \| **++CCCCAA++** \| **++TTGGGG++** \| \|; coexpression\_cluster2\_2\_m20\_cluster2\_2\_motif20\_m \| cluster2\_2\_motif20\_m \| cluster\_10 \| coexpression\_cluster2\_2 \| 15 \| 12.02 \| 49 \| **++AtTATTACTA+++** \| **+++TAGTAATAAT++** \| \|; coexpression\_cluster2\_2\_m35\_cluster2\_2\_motif35\_m \| cluster2\_2\_motif35\_m \| cluster\_16 \| coexpression\_cluster2\_2 \| 15 \| 18.95 \| 5 \| **CCCCTTAACyTAGAr** \| **YTCTARGTTAAGGGG** \| \|; coexpression\_cluster2\_2\_m25\_cluster2\_2\_motif25\_m \| cluster2\_2\_motif25\_m \| cluster\_6 \| coexpression\_cluster2\_2 \| 15 \| 12.51 \| 179 \| **TATATAyATRyATAy** \| **RTATRYATRTATATA** \| \|; coexpression\_cluster2\_2\_m2\_cluster2\_2\_motif2\_r \| cluster2\_2\_motif2\_r \| cluster\_1 \| coexpression\_cluster2\_2 \| 44 \| 15.85 \| 1249 \| **++WWWWTWTWATAAWTATWATAATAWAWAWAWW+++++++++++** \| **+++++++++++wwtwTwTwTATTATwATAwTTATwAwAwwww++** \| \|; coexpression\_cluster2\_2\_m38\_cluster2\_2\_motif38\_m \| cluster2\_2\_motif38\_m \| cluster\_14 \| coexpression\_cluster2\_2 \| 9 \| 10.22 \| 34 \| **CTATTAAAG** \| **CTTTAATAG** \| \|; coexpression\_cluster2\_2\_m23\_cluster2\_2\_motif23\_m \| cluster2\_2\_motif23\_m \| cluster\_3 \| coexpression\_cluster2\_2 \| 15 \| 16.13 \| 22 \| **GSAKAGAWAAKGGAA** \| **TTCCmTTwTCTMTSC** \| \|; coexpression\_cluster2\_2\_m6\_cluster2\_2\_motif6\_r \| cluster2\_2\_motif6\_r \| cluster\_1 \| coexpression\_cluster2\_2 \| 44 \| 18.24 \| 1218 \| **WWWWWWTATATWWWTAWWAATATWWWTATAWAWAWW++++++++** \| **++++++++wwtwtwTATAwwwATATTWwTAWwWATATawwwwww** \| \|; coexpression\_cluster2\_2\_m17\_cluster2\_2\_motif17\_r \| cluster2\_2\_motif17\_r \| cluster\_5 \| coexpression\_cluster2\_2 \| 15 \| 9.38 \| 206 \| **+wwAAACTAww++++** \| **++++WWTAGTTTWW+** \| \|; coexpression\_cluster2\_2\_m33\_cluster2\_2\_motif33\_m \| cluster2\_2\_motif33\_m \| cluster\_7 \| coexpression\_cluster2\_2 \| 15 \| 16.40 \| 22 \| **TAyATAGGGyTAAAk** \| **MTTTARCCCTATRTA** \| \|; coexpression\_cluster2\_2\_m12\_cluster2\_2\_motif12\_r \| cluster2\_2\_motif12\_r \| cluster\_1 \| coexpression\_cluster2\_2 \| 44 \| 16.42 \| 1100 \| **+++++WWWWTWTWAWTATAATAATWWAATATAWAWAWW++++++** \| **++++++wwtwTwtATATtwwATTATTATAwTwawawwww+++++** \| \|; coexpression\_cluster2\_2\_m30\_cluster2\_2\_motif30\_m \| cluster2\_2\_motif30\_m \| cluster\_11 \| coexpression\_cluster2\_2 \| 15 \| 14.32 \| 12 \| **AGryAAsrATryAGC** \| **GCTRYATYSTTRYCT** \| \|; coexpression\_cluster2\_2\_m28\_cluster2\_2\_motif28\_m \| cluster2\_2\_motif28\_m \| cluster\_8 \| coexpression\_cluster2\_2 \| 15 \| 15.93 \| 18 \| **rAAAKATGTGCAMAA** \| **TTKTGCACATMTTTY** \| \|; coexpression\_cluster2\_2\_m31\_cluster2\_2\_motif31\_m \| cluster2\_2\_motif31\_m \| cluster\_9 \| coexpression\_cluster2\_2 \| 11 \| 8.24 \| 20 \| **GCrYrCrWkgm** \| **KCMWYGYRYGC** \| \|; coexpression\_cluster2\_2\_m24\_cluster2\_2\_motif24\_m \| cluster2\_2\_motif24\_m \| cluster\_13 \| coexpression\_cluster2\_2 \| 15 \| 17.50 \| 13 \| **AATCACCCAATAAGG** \| **CCTTATTGGGTGATT** \| \|; coexpression\_cluster2\_2\_m15\_cluster2\_2\_motif15\_r \| cluster2\_2\_motif15\_r \| cluster\_1 \| coexpression\_cluster2\_2 \| 44 \| 17.13 \| 1021 \| **++WWWWWWTWATAATTATWATAATAWTAWAWAWWWW++++++++** \| **++++++++wwwwtwtwTawTATTATwATAATTATwAwwwwww++** \| \|; coexpression\_cluster2\_2\_m7\_cluster2\_2\_motif7\_r \| cluster2\_2\_motif7\_r \| cluster\_1 \| coexpression\_cluster2\_2 \| 44 \| 14.38 \| 930 \| **++WWWWTTTWATAATTATWATAATAWWW++++++++++++++++** \| **++++++++++++++++wwwTATTATWATAATTATwAAAwwww++** \| \|; coexpression\_cluster2\_2\_m10\_cluster2\_2\_motif10\_r \| cluster2\_2\_motif10\_r \| cluster\_1 \| coexpression\_cluster2\_2 \| 44 \| 21.48 \| 673 \| **WWTWWWTATAWWTATATWWTTAWWWWTWTWWTWATAWAWWWWWW** \| **wwwwwwtWTATWAwwAwawwwwTAawWATATawwTATAwwwaww** \| \| - Click on the column names to change the order of the data. - Write the name of one cluster, collection or a pattern in the *Search* window. **Heatmap View** Heatmap View Distance table \| PDF |

**Additional Files**

#### Exported files

| File | Description |
| --- | --- |
| **Pairwise comparison** | This table shows the pairwise comparison between all the input motifs using different metrics. This is the *compare-matrices* result. |
| **Matrix description** | This table shows information of each input motif. |
| **Cluster-wise Motif Files [zip]** | One TRANSFAC file is generated for each cluster including all the input motifs grouped in the corresponding cluster. |
| **Root motifs** | A file with the root matrix of each cluster. |
| **Motif collections per cluster** | A tab separated file containing the counts of motif of each collection on each cluster. |
| **Clusters** | A tab separated file containing the clusters and their correspondng motifs. |
| **Central motifs IDs** | A tab separated file containing the central motif, that one who is the most similar to all the others motif in the same cluster. |
| **Central motifs (in TRANSFAC format)** | A transfac file containing the central motif, that one who is the most similar to all the others motif within the same cluster. |
| **Input motifs** | The input motifs analyzed with matrix-clustering. |

- **References**
  1. Castro-Mondragon JA et al.  RSAT matrix-clustering: dynamic exploration and redundancy reduction of transcription factor binding motif collections. doi: 10.1093/nar/gkx314; Nucleic Acids Research (2017)
  2. **Last RSAT release:** Nguyen NTT et al. RSAT 2018: regulatory sequence analysis tools 20th anniversary. Nucleic Acids Research (2018).
- **Additional Information**
  - Jacques van Helden course
- **Contact**
  - Jaime Castro-Mondragon - ****
  - Morgane Thomas-Chollier - ****
  - Jacques van Helden - ****
