## Additional file 10 for "Transcriptome analysis reveals a novel DNA element that may interact with chromatin-associated proteins in *Plasmodium berghei* during erythrocytic development": matrix-clustering_SUMMARY_cluster2_4.html

matrix-clustering coexpression\_cluster2\_4

### RSAT - matrix-clustering result

##### Analysis: coexpression\_cluster2\_4 (08/09/2023 15:33)

##### Command

```
matrix-clustering  -v 1 -max_matrices 300 -matrix coexpression_cluster2_4 $RSAT/public_html/tmp/www-data/2023/09/08/matrix-clustering_2023-09-08.153321_0k34kN/matrix-clustering_query_matrices.transfac transfac -hclust_method average -calc sum -title coexpression_cluster2_4 -metric_build_tree Ncor -lth w 5 -lth cor 0.6 -lth Ncor 0.4 -quick -label_in_tree name -return json,heatmap -o $RSAT/public_html/tmp/www-data/2023/09/08/matrix-clustering_2023-09-08.153321_0k34kN/matrix-clustering
```

#### Results Summary

|  |  |  |  |  |  |
| --- | --- | --- | --- | --- | --- |
| Nb Input motifs | Nb Input collections | Nb Clusters Found | Download   Root motifs | Complete results [zip] | Parameters used Linkage method \| Similarity metric \| Thresholds to partition the tree |
| 15 | 1 | 11 | **Download** | Download | average \| Ncor \| **Ncor** = 0.4 **cor** = 0.6 |

|  |  |  |  |  |  |  |
| --- | --- | --- | --- | --- | --- | --- |
|  | **Logo Forest (dynamic browsing)**  **Logo Forest (rapid overview - low image quality)**  cluster\_1  cluster\_2  cluster\_3  cluster\_4  cluster\_5  cluster\_6  cluster\_7  cluster\_8  cluster\_9  cluster\_10  cluster\_11 |  |  |  |  |  |
|  | **Clusters Summary** Clusters Summary Table (Root Motif \| Root Motif (Reverse) \| Cluster ID \| # Motifs \| Motif\_Name::Collection \| Number of Motifs by Collection \| Root Motif (transfac Format)) \| | cluster\_1 | 2 | cluster2\_4\_motif8\_m::coexpression\_cluster2\_4:: cluster2\_4\_motif9\_m::coexpression\_cluster2\_4:: | coexpression\_cluster2\_4::2\_motifs | Show matrix |
| cluster\_2 \| 3 \| cluster2\_4\_motif15\_m::coexpression\_cluster2\_4:: cluster2\_4\_motif10\_m::coexpression\_cluster2\_4:: cluster2\_4\_motif7\_m::coexpression\_cluster2\_4:: \| coexpression\_cluster2\_4::3\_motifs \| Show matrix \| \| |  |  |  |  |  |  |
| cluster\_3 \| 2 \| cluster2\_4\_motif11\_m::coexpression\_cluster2\_4:: cluster2\_4\_motif12\_m::coexpression\_cluster2\_4:: \| coexpression\_cluster2\_4::2\_motifs \| Show matrix \| \| |  |  |  |  |  |  |
| cluster\_4 | 1 | cluster2\_4\_motif2\_r::coexpression\_cluster2\_4 | coexpression\_cluster2\_4 : 1 motif | Show matrix | | |
| cluster\_5 | 1 | cluster2\_4\_motif5\_r::coexpression\_cluster2\_4 | coexpression\_cluster2\_4 : 1 motif | Show matrix | | |
| cluster\_6 | 1 | cluster2\_4\_motif3\_r::coexpression\_cluster2\_4 | coexpression\_cluster2\_4 : 1 motif | Show matrix | | |
| cluster\_7 | 1 | cluster2\_4\_motif4\_r::coexpression\_cluster2\_4 | coexpression\_cluster2\_4 : 1 motif | Show matrix | | |
| cluster\_8 | 1 | cluster2\_4\_motif6\_r::coexpression\_cluster2\_4 | coexpression\_cluster2\_4 : 1 motif | Show matrix | | |
| cluster\_9 | 1 | cluster2\_4\_motif1\_r::coexpression\_cluster2\_4 | coexpression\_cluster2\_4 : 1 motif | Show matrix | | |
| cluster\_10 | 1 | cluster2\_4\_motif13\_m::coexpression\_cluster2\_4 | coexpression\_cluster2\_4 : 1 motif | Show matrix | | |
| cluster\_11 | 1 | cluster2\_4\_motif14\_m::coexpression\_cluster2\_4 | coexpression\_cluster2\_4 : 1 motif | Show matrix | | |

- Note: by default the table displays the first 50 clusters. Click on the *Show X entries* button to display more (or less) clusters.
- Click on the column names to change the order of the data.
- Use the *Search* window to find your motif (s) of interest.

| **Individual Cluster View** Individual Cluster View **Show All** **Hide All** **cluster\_1** **cluster\_2** **cluster\_3** **cluster\_4** **cluster\_5** **cluster\_6** **cluster\_7** **cluster\_8** **cluster\_9** **cluster\_10** **cluster\_11** **Display Logo Trees** **cluster\_1** **cluster\_2** **cluster\_3** **cluster\_4** **cluster\_5** **cluster\_6** **cluster\_7** **cluster\_8** **cluster\_9** **cluster\_10** **cluster\_11** **Display Branch-Motifs** **cluster\_1** Branch Motifs \| \| \| \| \| \| \|; Node \| Consensus \| Logo \| Logo (Reverse) \| Collections \| Matrix (Transfac format) \| IC \| Nb sites; Node 1 \| **ATATATAyATryATR** \| \| \| coexpression\_cluster2\_4::2\_motifs \| PSSM \| 12.83 \| 373 **cluster\_2** Branch Motifs \| \| \| \| \| \| \|; Node \| Consensus \| Logo \| Logo (Reverse) \| Collections \| Matrix (Transfac format) \| IC \| Nb sites; Node 1 \| **AAAAAAwaAdAaAAAA** \| \| \| coexpression\_cluster2\_4::2\_motifs \| PSSM \| 12.77 \| 326; Node 2 \| **AAAAAAwAAdAaAAAA** \| \| \| coexpression\_cluster2\_4::3\_motifs \| PSSM \| 12.48 \| 369 **cluster\_3** Branch Motifs \| \| \| \| \| \| \|; Node \| Consensus \| Logo \| Logo (Reverse) \| Collections \| Matrix (Transfac format) \| IC \| Nb sites; Node 1 \| **yTcrTCAyTT** \| \| \| coexpression\_cluster2\_4::2\_motifs \| PSSM \| 8.02 \| 145 **cluster\_4** Branch Motifs \| \| \| \| \| \| \|; Node \| Consensus \| Logo \| Logo (Reverse) \| Collections \| Matrix (Transfac format) \| IC \| Nb sites; Singleton \| **wwCAAATGww** \| \| \| coexpression\_cluster2\_4 : 1 motif \| PSSM \| 8.83 \| 138 **cluster\_5** Branch Motifs \| \| \| \| \| \| \|; Node \| Consensus \| Logo \| Logo (Reverse) \| Collections \| Matrix (Transfac format) \| IC \| Nb sites; Singleton \| **twCAATCGtt** \| \| \| coexpression\_cluster2\_4 : 1 motif \| PSSM \| 8.83 \| 22 **cluster\_6** Branch Motifs \| \| \| \| \| \| \|; Node \| Consensus \| Logo \| Logo (Reverse) \| Collections \| Matrix (Transfac format) \| IC \| Nb sites; Singleton \| **wwGATGAArAww** \| \| \| coexpression\_cluster2\_4 : 1 motif \| PSSM \| 9.16 \| 105 **cluster\_7** Branch Motifs \| \| \| \| \| \| \|; Node \| Consensus \| Logo \| Logo (Reverse) \| Collections \| Matrix (Transfac format) \| IC \| Nb sites; Singleton \| **wwgAAGCAAww** \| \| \| coexpression\_cluster2\_4 : 1 motif \| PSSM \| 7.19 \| 323 **cluster\_8** Branch Motifs \| \| \| \| \| \| \|; Node \| Consensus \| Logo \| Logo (Reverse) \| Collections \| Matrix (Transfac format) \| IC \| Nb sites; Singleton \| **wwCCTTATAww** \| \| \| coexpression\_cluster2\_4 : 1 motif \| PSSM \| 7.05 \| 701 **cluster\_9** Branch Motifs \| \| \| \| \| \| \|; Node \| Consensus \| Logo \| Logo (Reverse) \| Collections \| Matrix (Transfac format) \| IC \| Nb sites; Singleton \| **awAGGATCrw** \| \| \| coexpression\_cluster2\_4 : 1 motif \| PSSM \| 8.88 \| 21 **cluster\_10** Branch Motifs \| \| \| \| \| \| \|; Node \| Consensus \| Logo \| Logo (Reverse) \| Collections \| Matrix (Transfac format) \| IC \| Nb sites; Singleton \| **SSSCCMCm** \| \| \| coexpression\_cluster2\_4 : 1 motif \| PSSM \| 7.17 \| 33 **cluster\_11** Branch Motifs \| \| \| \| \| \| \|; Node \| Consensus \| Logo \| Logo (Reverse) \| Collections \| Matrix (Transfac format) \| IC \| Nb sites; Singleton \| **SaYCCCCyCcSamkY** \| \| \| coexpression\_cluster2\_4 : 1 motif \| PSSM \| 11.20 \| 18 - **Each cluster has a different color.** - **Click on the upper buttons to display separately the information of each cluster.** - **Click on the circles at each tree to display its corresponding merged motifs.** Definitions - **Logo tree.** This results shows the hierarchical tree with its logo alignment of one cluster. The logos are shown in Forward (left logo) and Reverse (right logo) orientation. - **Branch-motifs.** On the trees are displayed the branch number that is the number in which the motifs were incorporated in the tree. The Branch-Motifs table shows the logo in both orientations and the link to the file in TRANSFAC format with the branch-motif. |
|  | **Individual Motif View** Individual Motif View Motif id \| Motif name \| Cluster \| Collection \| Width \| IC \| Number of sites \| Consensus \| Consensus (Rev) \| Logo \| Logo (Rev); coexpression\_cluster2\_4\_m13\_cluster2\_4\_motif13\_m \| cluster2\_4\_motif13\_m \| cluster\_10 \| coexpression\_cluster2\_4 \| 8 \| 7.17 \| 33 \| **ssCCCmMA** \| **TKKGGGSS** \| \|; coexpression\_cluster2\_4\_m9\_cluster2\_4\_motif9\_m \| cluster2\_4\_motif9\_m \| cluster\_1 \| coexpression\_cluster2\_4 \| 15 \| 12.37 \| 253 \| **ATATATAyATryATr** \| **YATRYATRTATATAT** \| \|; coexpression\_cluster2\_4\_m14\_cluster2\_4\_motif14\_m \| cluster2\_4\_motif14\_m \| cluster\_11 \| coexpression\_cluster2\_4 \| 15 \| 11.20 \| 18 \| **CAymCyCTChsAwTy** \| **RAWTSDGAGRGKRTG** \| \|; coexpression\_cluster2\_4\_m6\_cluster2\_4\_motif6\_r \| cluster2\_4\_motif6\_r \| cluster\_8 \| coexpression\_cluster2\_4 \| 11 \| 7.05 \| 701 \| **wwCCTTATAww** \| **WWTATAAGGWW** \| \|; coexpression\_cluster2\_4\_m11\_cluster2\_4\_motif11\_m \| cluster2\_4\_motif11\_m \| cluster\_3 \| coexpression\_cluster2\_4 \| 10 \| 8.42 \| 102 \| **+TCATCAcTT** \| **AAGTGATGA+** \| \|; coexpression\_cluster2\_4\_m15\_cluster2\_4\_motif15\_m \| cluster2\_4\_motif15\_m \| cluster\_2 \| coexpression\_cluster2\_4 \| 16 \| 15.78 \| 43 \| **+TTTTTATTTTATTW+** \| **+WAATAAAATAAAAA+** \| \|; coexpression\_cluster2\_4\_m10\_cluster2\_4\_motif10\_m \| cluster2\_4\_motif10\_m \| cluster\_2 \| coexpression\_cluster2\_4 \| 16 \| 12.79 \| 223 \| **AAAAAAwrArAwAAA+** \| **+TTTWTYTYWTTTTTT** \| \|; coexpression\_cluster2\_4\_m8\_cluster2\_4\_motif8\_m \| cluster2\_4\_motif8\_m \| cluster\_1 \| coexpression\_cluster2\_4 \| 15 \| 14.72 \| 120 \| **+TATrTATATAyATA** \| **TATRTATATAYATA+** \| \|; coexpression\_cluster2\_4\_m5\_cluster2\_4\_motif5\_r \| cluster2\_4\_motif5\_r \| cluster\_5 \| coexpression\_cluster2\_4 \| 10 \| 8.83 \| 22 \| **twCAATCGtt** \| **AACGATTGWA** \| \|; coexpression\_cluster2\_4\_m1\_cluster2\_4\_motif1\_r \| cluster2\_4\_motif1\_r \| cluster\_9 \| coexpression\_cluster2\_4 \| 10 \| 8.88 \| 21 \| **awAGGATCrw** \| **WYGATCCTWT** \| \|; coexpression\_cluster2\_4\_m2\_cluster2\_4\_motif2\_r \| cluster2\_4\_motif2\_r \| cluster\_4 \| coexpression\_cluster2\_4 \| 10 \| 8.83 \| 138 \| **wwCAAATGww** \| **WWCATTTGWW** \| \|; coexpression\_cluster2\_4\_m4\_cluster2\_4\_motif4\_r \| cluster2\_4\_motif4\_r \| cluster\_7 \| coexpression\_cluster2\_4 \| 11 \| 7.19 \| 323 \| **wwgAAGCAAww** \| **WWTTGCTTCWW** \| \|; coexpression\_cluster2\_4\_m7\_cluster2\_4\_motif7\_m \| cluster2\_4\_motif7\_m \| cluster\_2 \| coexpression\_cluster2\_4 \| 16 \| 14.86 \| 103 \| **+AAAAATAATAAAahA** \| **TDTTTTATTATTTTT+** \| \|; coexpression\_cluster2\_4\_m3\_cluster2\_4\_motif3\_r \| cluster2\_4\_motif3\_r \| cluster\_6 \| coexpression\_cluster2\_4 \| 12 \| 9.16 \| 105 \| **wwGATGAArAww** \| **WWTYTTCATCWW** \| \|; coexpression\_cluster2\_4\_m12\_cluster2\_4\_motif12\_m \| cluster2\_4\_motif12\_m \| cluster\_3 \| coexpression\_cluster2\_4 \| 10 \| 9.89 \| 43 \| **YTKGTCATTT** \| **AAaTGACmAr** \| \| - Click on the column names to change the order of the data. - Write the name of one cluster, collection or a pattern in the *Search* window. **Heatmap View** Heatmap View Distance table \| PDF |

**Additional Files**

#### Exported files

| File | Description |
| --- | --- |
| **Pairwise comparison** | This table shows the pairwise comparison between all the input motifs using different metrics. This is the *compare-matrices* result. |
| **Matrix description** | This table shows information of each input motif. |
| **Cluster-wise Motif Files [zip]** | One TRANSFAC file is generated for each cluster including all the input motifs grouped in the corresponding cluster. |
| **Root motifs** | A file with the root matrix of each cluster. |
| **Motif collections per cluster** | A tab separated file containing the counts of motif of each collection on each cluster. |
| **Clusters** | A tab separated file containing the clusters and their correspondng motifs. |
| **Central motifs IDs** | A tab separated file containing the central motif, that one who is the most similar to all the others motif in the same cluster. |
| **Central motifs (in TRANSFAC format)** | A transfac file containing the central motif, that one who is the most similar to all the others motif within the same cluster. |
| **Input motifs** | The input motifs analyzed with matrix-clustering. |

- **References**
  1. Castro-Mondragon JA et al.  RSAT matrix-clustering: dynamic exploration and redundancy reduction of transcription factor binding motif collections. doi: 10.1093/nar/gkx314; Nucleic Acids Research (2017)
  2. **Last RSAT release:** Nguyen NTT et al. RSAT 2018: regulatory sequence analysis tools 20th anniversary. Nucleic Acids Research (2018).
- **Additional Information**
  - Jacques van Helden course
- **Contact**
  - Jaime Castro-Mondragon - ****
  - Morgane Thomas-Chollier - ****
  - Jacques van Helden - ****
