## Additional file 10 for "Transcriptome analysis reveals a novel DNA element that may interact with chromatin-associated proteins in *Plasmodium berghei* during erythrocytic development": matrix-clustering_SUMMARY_cluster2_11.html

matrix-clustering coexpression\_cluster2\_11

### RSAT - matrix-clustering result

##### Analysis: coexpression\_cluster2\_11 (08/09/2023 15:46)

##### Command

```
matrix-clustering  -v 1 -max_matrices 300 -matrix coexpression_cluster2_11 $RSAT/public_html/tmp/www-data/2023/09/08/matrix-clustering_2023-09-08.154551_5ZxetT/matrix-clustering_query_matrices.transfac transfac -hclust_method average -calc sum -title coexpression_cluster2_11 -metric_build_tree Ncor -lth w 5 -lth cor 0.6 -lth Ncor 0.4 -quick -label_in_tree name -return json,heatmap -o $RSAT/public_html/tmp/www-data/2023/09/08/matrix-clustering_2023-09-08.154551_5ZxetT/matrix-clustering
```

#### Results Summary

|  |  |  |  |  |  |
| --- | --- | --- | --- | --- | --- |
| Nb Input motifs | Nb Input collections | Nb Clusters Found | Download   Root motifs | Complete results [zip] | Parameters used Linkage method \| Similarity metric \| Thresholds to partition the tree |
| 14 | 1 | 8 | **Download** | Download | average \| Ncor \| **Ncor** = 0.4 **cor** = 0.6 |

|  |  |  |  |  |  |  |
| --- | --- | --- | --- | --- | --- | --- |
|  | **Logo Forest (dynamic browsing)**  **Logo Forest (rapid overview - low image quality)**  cluster\_1  cluster\_2  cluster\_3  cluster\_4  cluster\_5  cluster\_6  cluster\_7  cluster\_8 |  |  |  |  |  |
|  | **Clusters Summary** Clusters Summary Table (Root Motif \| Root Motif (Reverse) \| Cluster ID \| # Motifs \| Motif\_Name::Collection \| Number of Motifs by Collection \| Root Motif (transfac Format)) \| | cluster\_1 | 5 | cluster2\_11\_motif10\_m::coexpression\_cluster2\_11:: cluster2\_11\_motif1\_r::coexpression\_cluster2\_11:: cluster2\_11\_motif2\_r::coexpression\_cluster2\_11:: cluster2\_11\_motif5\_m::coexpression\_cluster2\_11:: cluster2\_11\_motif6\_m::coexpression\_cluster2\_11:: | coexpression\_cluster2\_11::5\_motifs | Show matrix |
| cluster\_2 \| 2 \| cluster2\_11\_motif11\_m::coexpression\_cluster2\_11:: cluster2\_11\_motif8\_m::coexpression\_cluster2\_11:: \| coexpression\_cluster2\_11::2\_motifs \| Show matrix \| \| |  |  |  |  |  |  |
| cluster\_3 \| 2 \| cluster2\_11\_motif7\_m::coexpression\_cluster2\_11:: cluster2\_11\_motif9\_m::coexpression\_cluster2\_11:: \| coexpression\_cluster2\_11::2\_motifs \| Show matrix \| \| |  |  |  |  |  |  |
| cluster\_4 | 1 | cluster2\_11\_motif4\_r::coexpression\_cluster2\_11 | coexpression\_cluster2\_11 : 1 motif | Show matrix | | |
| cluster\_5 | 1 | cluster2\_11\_motif3\_r::coexpression\_cluster2\_11 | coexpression\_cluster2\_11 : 1 motif | Show matrix | | |
| cluster\_6 | 1 | cluster2\_11\_motif12\_m::coexpression\_cluster2\_11 | coexpression\_cluster2\_11 : 1 motif | Show matrix | | |
| cluster\_7 | 1 | cluster2\_11\_motif13\_m::coexpression\_cluster2\_11 | coexpression\_cluster2\_11 : 1 motif | Show matrix | | |
| cluster\_8 | 1 | cluster2\_11\_motif14\_m::coexpression\_cluster2\_11 | coexpression\_cluster2\_11 : 1 motif | Show matrix | | |

- Note: by default the table displays the first 50 clusters. Click on the *Show X entries* button to display more (or less) clusters.
- Click on the column names to change the order of the data.
- Use the *Search* window to find your motif (s) of interest.

| **Individual Cluster View** Individual Cluster View **Show All** **Hide All** **cluster\_1** **cluster\_2** **cluster\_3** **cluster\_4** **cluster\_5** **cluster\_6** **cluster\_7** **cluster\_8** **Display Logo Trees** **cluster\_1** **cluster\_2** **cluster\_3** **cluster\_4** **cluster\_5** **cluster\_6** **cluster\_7** **cluster\_8** **Display Branch-Motifs** **cluster\_1** Branch Motifs \| \| \| \| \| \| \|; Node \| Consensus \| Logo \| Logo (Reverse) \| Collections \| Matrix (Transfac format) \| IC \| Nb sites; Node 1 \| **--wwAAAgTGAaaww-** \| \| \| coexpression\_cluster2\_11::2\_motifs \| PSSM \| 8.01 \| 1192; Node 2 \| **-AAAAAarRawAAwAA** \| \| \| coexpression\_cluster2\_11::2\_motifs \| PSSM \| 10.89 \| 400; Node 3 \| **-AaaAAArtgAAawaA** \| \| \| coexpression\_cluster2\_11::4\_motifs \| PSSM \| 8.87 \| 1591; Node 4 \| **GAaaAAArtgAAawaA** \| \| \| coexpression\_cluster2\_11::5\_motifs \| PSSM \| 9.91 \| 1645 **cluster\_2** Branch Motifs \| \| \| \| \| \| \|; Node \| Consensus \| Logo \| Logo (Reverse) \| Collections \| Matrix (Transfac format) \| IC \| Nb sites; Node 1 \| **TaYrCACACAhw** \| \| \| coexpression\_cluster2\_11::2\_motifs \| PSSM \| 10.18 \| 103 **cluster\_3** Branch Motifs \| \| \| \| \| \| \|; Node \| Consensus \| Logo \| Logo (Reverse) \| Collections \| Matrix (Transfac format) \| IC \| Nb sites; Node 1 \| **rTRTATryATATATR** \| \| \| coexpression\_cluster2\_11::2\_motifs \| PSSM \| 12.86 \| 404 **cluster\_4** Branch Motifs \| \| \| \| \| \| \|; Node \| Consensus \| Logo \| Logo (Reverse) \| Collections \| Matrix (Transfac format) \| IC \| Nb sites; Singleton \| **wrCGTGCAtw** \| \| \| coexpression\_cluster2\_11 : 1 motif \| PSSM \| 9.78 \| 26 **cluster\_5** Branch Motifs \| \| \| \| \| \| \|; Node \| Consensus \| Logo \| Logo (Reverse) \| Collections \| Matrix (Transfac format) \| IC \| Nb sites; Singleton \| **wwATAATGww** \| \| \| coexpression\_cluster2\_11 : 1 motif \| PSSM \| 9.10 \| 296 **cluster\_6** Branch Motifs \| \| \| \| \| \| \|; Node \| Consensus \| Logo \| Logo (Reverse) \| Collections \| Matrix (Transfac format) \| IC \| Nb sites; Singleton \| **sSsvSMrSSaGCataCGmSkVmS** \| \| \| coexpression\_cluster2\_11 : 1 motif \| PSSM \| 10.03 \| 16 **cluster\_7** Branch Motifs \| \| \| \| \| \| \|; Node \| Consensus \| Logo \| Logo (Reverse) \| Collections \| Matrix (Transfac format) \| IC \| Nb sites; Singleton \| **ksbssbSCRCaCaCaCSsSsvb** \| \| \| coexpression\_cluster2\_11 : 1 motif \| PSSM \| 12.41 \| 28 **cluster\_8** Branch Motifs \| \| \| \| \| \| \|; Node \| Consensus \| Logo \| Logo (Reverse) \| Collections \| Matrix (Transfac format) \| IC \| Nb sites; Singleton \| **CSCCSSCa** \| \| \| coexpression\_cluster2\_11 : 1 motif \| PSSM \| 8.11 \| 14 - **Each cluster has a different color.** - **Click on the upper buttons to display separately the information of each cluster.** - **Click on the circles at each tree to display its corresponding merged motifs.** Definitions - **Logo tree.** This results shows the hierarchical tree with its logo alignment of one cluster. The logos are shown in Forward (left logo) and Reverse (right logo) orientation. - **Branch-motifs.** On the trees are displayed the branch number that is the number in which the motifs were incorporated in the tree. The Branch-Motifs table shows the logo in both orientations and the link to the file in TRANSFAC format with the branch-motif. |
|  | **Individual Motif View** Individual Motif View Motif id \| Motif name \| Cluster \| Collection \| Width \| IC \| Number of sites \| Consensus \| Consensus (Rev) \| Logo \| Logo (Rev); coexpression\_cluster2\_11\_m6\_cluster2\_11\_motif6\_m \| cluster2\_11\_motif6\_m \| cluster\_1 \| coexpression\_cluster2\_11 \| 16 \| 11.60 \| 288 \| **+ARAAAwrrrAAAAaA** \| **TTTTTTYYYWTTTYT+** \| \|; coexpression\_cluster2\_11\_m3\_cluster2\_11\_motif3\_r \| cluster2\_11\_motif3\_r \| cluster\_5 \| coexpression\_cluster2\_11 \| 10 \| 9.10 \| 296 \| **wwATAATGww** \| **WWCATTATWW** \| \|; coexpression\_cluster2\_11\_m4\_cluster2\_11\_motif4\_r \| cluster2\_11\_motif4\_r \| cluster\_4 \| coexpression\_cluster2\_11 \| 10 \| 9.78 \| 26 \| **wrCGTGCATw** \| **WATGCACGYW** \| \|; coexpression\_cluster2\_11\_m13\_cluster2\_11\_motif13\_m \| cluster2\_11\_motif13\_m \| cluster\_7 \| coexpression\_cluster2\_11 \| 22 \| 12.41 \| 28 \| **TvtddtsCRCACACACrhrkmw** \| **WKMYDYGTGTGTGYGSAHHABA** \| \|; coexpression\_cluster2\_11\_m11\_cluster2\_11\_motif11\_m \| cluster2\_11\_motif11\_m \| cluster\_2 \| coexpression\_cluster2\_11 \| 12 \| 10.31 \| 35 \| **+DYACACACAYW** \| **wrTGTGTGTRh+** \| \|; coexpression\_cluster2\_11\_m7\_cluster2\_11\_motif7\_m \| cluster2\_11\_motif7\_m \| cluster\_3 \| coexpression\_cluster2\_11 \| 15 \| 12.62 \| 282 \| **rTryATryATATATR** \| **YATATATRYATRYAY** \| \|; coexpression\_cluster2\_11\_m1\_cluster2\_11\_motif1\_r \| cluster2\_11\_motif1\_r \| cluster\_1 \| coexpression\_cluster2\_11 \| 16 \| 7.38 \| 618 \| **++wwAAAgTGAww+++** \| **+++WWTCACTTTWW++** \| \|; coexpression\_cluster2\_11\_m9\_cluster2\_11\_motif9\_m \| cluster2\_11\_motif9\_m \| cluster\_3 \| coexpression\_cluster2\_11 \| 15 \| 12.00 \| 122 \| **+TATATGCATAT+++** \| **+++ATATGCATATA+** \| \|; coexpression\_cluster2\_11\_m12\_cluster2\_11\_motif12\_m \| cluster2\_11\_motif12\_m \| cluster\_6 \| coexpression\_cluster2\_11 \| 23 \| 10.03 \| 16 \| **wmbmsmahkAGMATACrwkTrhr** \| **YDYAMWYGTATKCTMDTKSKVKW** \| \|; coexpression\_cluster2\_11\_m10\_cluster2\_11\_motif10\_m \| cluster2\_11\_motif10\_m \| cluster\_1 \| coexpression\_cluster2\_11 \| 16 \| 10.98 \| 54 \| **YTTTTTTTCTC+++++** \| **+++++GAGAAAaAAAr** \| \|; coexpression\_cluster2\_11\_m5\_cluster2\_11\_motif5\_m \| cluster2\_11\_motif5\_m \| cluster\_1 \| coexpression\_cluster2\_11 \| 16 \| 15.84 \| 112 \| **+AAAAAAAAATAATAw** \| **WTATTATTTTTTTTT+** \| \|; coexpression\_cluster2\_11\_m2\_cluster2\_11\_motif2\_r \| cluster2\_11\_motif2\_r \| cluster\_1 \| coexpression\_cluster2\_11 \| 16 \| 8.27 \| 574 \| **++wwAAAgTGAAAww+** \| **+WWTTTCACTTTWW++** \| \|; coexpression\_cluster2\_11\_m14\_cluster2\_11\_motif14\_m \| cluster2\_11\_motif14\_m \| cluster\_8 \| coexpression\_cluster2\_11 \| 8 \| 8.11 \| 14 \| **CCCCyCCA** \| **TGGRGGGG** \| \|; coexpression\_cluster2\_11\_m8\_cluster2\_11\_motif8\_m \| cluster2\_11\_motif8\_m \| cluster\_2 \| coexpression\_cluster2\_11 \| 12 \| 11.24 \| 68 \| **TATgCACACAww** \| **WWTGTGTGCATA** \| \| - Click on the column names to change the order of the data. - Write the name of one cluster, collection or a pattern in the *Search* window. **Heatmap View** Heatmap View Distance table \| PDF |

**Additional Files**

#### Exported files

| File | Description |
| --- | --- |
| **Pairwise comparison** | This table shows the pairwise comparison between all the input motifs using different metrics. This is the *compare-matrices* result. |
| **Matrix description** | This table shows information of each input motif. |
| **Cluster-wise Motif Files [zip]** | One TRANSFAC file is generated for each cluster including all the input motifs grouped in the corresponding cluster. |
| **Root motifs** | A file with the root matrix of each cluster. |
| **Motif collections per cluster** | A tab separated file containing the counts of motif of each collection on each cluster. |
| **Clusters** | A tab separated file containing the clusters and their correspondng motifs. |
| **Central motifs IDs** | A tab separated file containing the central motif, that one who is the most similar to all the others motif in the same cluster. |
| **Central motifs (in TRANSFAC format)** | A transfac file containing the central motif, that one who is the most similar to all the others motif within the same cluster. |
| **Input motifs** | The input motifs analyzed with matrix-clustering. |

- **References**
  1. Castro-Mondragon JA et al.  RSAT matrix-clustering: dynamic exploration and redundancy reduction of transcription factor binding motif collections. doi: 10.1093/nar/gkx314; Nucleic Acids Research (2017)
  2. **Last RSAT release:** Nguyen NTT et al. RSAT 2018: regulatory sequence analysis tools 20th anniversary. Nucleic Acids Research (2018).
- **Additional Information**
  - Jacques van Helden course
- **Contact**
  - Jaime Castro-Mondragon - ****
  - Morgane Thomas-Chollier - ****
  - Jacques van Helden - ****
