## Additional file 10 for "Transcriptome analysis reveals a novel DNA element that may interact with chromatin-associated proteins in *Plasmodium berghei* during erythrocytic development": matrix-clustering_SUMMARY_cluster3_5.html

matrix-clustering coexpression\_cluster3\_5

### RSAT - matrix-clustering result

##### Analysis: coexpression\_cluster3\_5 (08/09/2023 15:54)

##### Command

```
matrix-clustering  -v 1 -max_matrices 300 -matrix coexpression_cluster3_5 $RSAT/public_html/tmp/www-data/2023/09/08/matrix-clustering_2023-09-08.155430_2GIcCG/matrix-clustering_query_matrices.transfac transfac -hclust_method average -calc sum -title coexpression_cluster3_5 -metric_build_tree Ncor -lth w 5 -lth cor 0.6 -lth Ncor 0.4 -quick -label_in_tree name -return json,heatmap -o $RSAT/public_html/tmp/www-data/2023/09/08/matrix-clustering_2023-09-08.155430_2GIcCG/matrix-clustering
```

#### Results Summary

|  |  |  |  |  |  |
| --- | --- | --- | --- | --- | --- |
| Nb Input motifs | Nb Input collections | Nb Clusters Found | Download   Root motifs | Complete results [zip] | Parameters used Linkage method \| Similarity metric \| Thresholds to partition the tree |
| 34 | 1 | 9 | **Download** | Download | average \| Ncor \| **Ncor** = 0.4 **cor** = 0.6 |

|  |  |  |  |  |  |  |
| --- | --- | --- | --- | --- | --- | --- |
|  | **Logo Forest (dynamic browsing)**  **Logo Forest (rapid overview - low image quality)**  cluster\_1  cluster\_2  cluster\_3  cluster\_4  cluster\_5  cluster\_6  cluster\_7  cluster\_8  cluster\_9 |  |  |  |  |  |
|  | **Clusters Summary** Clusters Summary Table (Root Motif \| Root Motif (Reverse) \| Cluster ID \| # Motifs \| Motif\_Name::Collection \| Number of Motifs by Collection \| Root Motif (transfac Format)) \| | cluster\_1 | 15 | cluster3\_5\_motif32\_m::coexpression\_cluster3\_5:: cluster3\_5\_motif11\_r::coexpression\_cluster3\_5:: cluster3\_5\_motif12\_r::coexpression\_cluster3\_5:: cluster3\_5\_motif2\_r::coexpression\_cluster3\_5:: cluster3\_5\_motif6\_r::coexpression\_cluster3\_5:: cluster3\_5\_motif7\_r::coexpression\_cluster3\_5:: cluster3\_5\_motif3\_r::coexpression\_cluster3\_5:: cluster3\_5\_motif5\_r::coexpression\_cluster3\_5:: cluster3\_5\_motif10\_r::coexpression\_cluster3\_5:: cluster3\_5\_motif1\_r::coexpression\_cluster3\_5:: cluster3\_5\_motif16\_r::coexpression\_cluster3\_5:: cluster3\_5\_motif8\_r::coexpression\_cluster3\_5:: cluster3\_5\_motif23\_m::coexpression\_cluster3\_5:: cluster3\_5\_motif26\_m::coexpression\_cluster3\_5:: cluster3\_5\_motif28\_m::coexpression\_cluster3\_5:: | coexpression\_cluster3\_5::15\_motifs | Show matrix |
| cluster\_2 \| 10 \| cluster3\_5\_motif31\_m::coexpression\_cluster3\_5:: cluster3\_5\_motif9\_r::coexpression\_cluster3\_5:: cluster3\_5\_motif24\_m::coexpression\_cluster3\_5:: cluster3\_5\_motif25\_m::coexpression\_cluster3\_5:: cluster3\_5\_motif29\_m::coexpression\_cluster3\_5:: cluster3\_5\_motif13\_r::coexpression\_cluster3\_5:: cluster3\_5\_motif19\_r::coexpression\_cluster3\_5:: cluster3\_5\_motif4\_r::coexpression\_cluster3\_5:: cluster3\_5\_motif14\_r::coexpression\_cluster3\_5:: cluster3\_5\_motif34\_m::coexpression\_cluster3\_5:: \| coexpression\_cluster3\_5::10\_motifs \| Show matrix \| \| |  |  |  |  |  |  |
| cluster\_3 \| 2 \| cluster3\_5\_motif15\_r::coexpression\_cluster3\_5:: cluster3\_5\_motif21\_r::coexpression\_cluster3\_5:: \| coexpression\_cluster3\_5::2\_motifs \| Show matrix \| \| |  |  |  |  |  |  |
| cluster\_4 | 2 | cluster3\_5\_motif20\_r::coexpression\_cluster3\_5:: cluster3\_5\_motif22\_r::coexpression\_cluster3\_5:: | coexpression\_cluster3\_5::2\_motifs | Show matrix | | |
| cluster\_5 | 1 | cluster3\_5\_motif30\_m::coexpression\_cluster3\_5 | coexpression\_cluster3\_5 : 1 motif | Show matrix | | |
| cluster\_6 | 1 | cluster3\_5\_motif18\_r::coexpression\_cluster3\_5 | coexpression\_cluster3\_5 : 1 motif | Show matrix | | |
| cluster\_7 | 1 | cluster3\_5\_motif33\_m::coexpression\_cluster3\_5 | coexpression\_cluster3\_5 : 1 motif | Show matrix | | |
| cluster\_8 | 1 | cluster3\_5\_motif27\_m::coexpression\_cluster3\_5 | coexpression\_cluster3\_5 : 1 motif | Show matrix | | |
| cluster\_9 | 1 | cluster3\_5\_motif17\_r::coexpression\_cluster3\_5 | coexpression\_cluster3\_5 : 1 motif | Show matrix | | |

- Note: by default the table displays the first 50 clusters. Click on the *Show X entries* button to display more (or less) clusters.
- Click on the column names to change the order of the data.
- Use the *Search* window to find your motif (s) of interest.

| **Individual Cluster View** Individual Cluster View **Show All** **Hide All** **cluster\_1** **cluster\_2** **cluster\_3** **cluster\_4** **cluster\_5** **cluster\_6** **cluster\_7** **cluster\_8** **cluster\_9** **Display Logo Trees** **cluster\_1** **cluster\_2** **cluster\_3** **cluster\_4** **cluster\_5** **cluster\_6** **cluster\_7** **cluster\_8** **cluster\_9** **Display Branch-Motifs** **cluster\_1** Branch Motifs \| \| \| \| \| \| \|; Node \| Consensus \| Logo \| Logo (Reverse) \| Collections \| Matrix (Transfac format) \| IC \| Nb sites; Node 1 \| **-waAAAAgACAawww** \| \| \| coexpression\_cluster3\_5::2\_motifs \| PSSM \| 9.16 \| 1941; Node 2 \| **-waAAAArAmAAAww** \| \| \| coexpression\_cluster3\_5::3\_motifs \| PSSM \| 9.26 \| 5455; Node 3 \| **-wwTATAkAcAAAww** \| \| \| coexpression\_cluster3\_5::2\_motifs \| PSSM \| 8.70 \| 3205; Node 4 \| **-waAAAArAmAAwww** \| \| \| coexpression\_cluster3\_5::4\_motifs \| PSSM \| 9.07 \| 7669; Node 5 \| **-wwTATAgAcAAAww** \| \| \| coexpression\_cluster3\_5::3\_motifs \| PSSM \| 8.72 \| 4017; Node 6 \| **-waAAAArAmAAwww** \| \| \| coexpression\_cluster3\_5::5\_motifs \| PSSM \| 8.97 \| 8906; Node 7 \| **-wwwATAgAcAAAww** \| \| \| coexpression\_cluster3\_5::4\_motifs \| PSSM \| 8.58 \| 6360; Node 8 \| **-waAAAArAmAwAww** \| \| \| coexpression\_cluster3\_5::6\_motifs \| PSSM \| 8.82 \| 11727; Node 9 \| **-waAAAArAmAwAww** \| \| \| coexpression\_cluster3\_5::7\_motifs \| PSSM \| 8.73 \| 12788; Node 10 \| **AAAAAAAaraAAwAA** \| \| \| coexpression\_cluster3\_5::2\_motifs \| PSSM \| 11.83 \| 550; Node 11 \| **-wwAAwArAmAAAww** \| \| \| coexpression\_cluster3\_5::11\_motifs \| PSSM \| 8.11 \| 19148; Node 12 \| **AAAwAAAAwaAAwAA** \| \| \| coexpression\_cluster3\_5::3\_motifs \| PSSM \| 11.18 \| 856; Node 13 \| **AwwAAwArAmAAAww** \| \| \| coexpression\_cluster3\_5::14\_motifs \| PSSM \| 8.92 \| 20004; Node 14 \| **AwwAAwArAmAAAww** \| \| \| coexpression\_cluster3\_5::15\_motifs \| PSSM \| 8.90 \| 20117 **cluster\_2** Branch Motifs \| \| \| \| \| \| \|; Node \| Consensus \| Logo \| Logo (Reverse) \| Collections \| Matrix (Transfac format) \| IC \| Nb sites; Node 1 \| **--wwwtgTGTGTGCATww----** \| \| \| coexpression\_cluster3\_5::2\_motifs \| PSSM \| 9.46 \| 867; Node 2 \| **---TATaTrYaTrTaTRYA---** \| \| \| coexpression\_cluster3\_5::2\_motifs \| PSSM \| 12.39 \| 1013; Node 3 \| **---taTATrTaTATAtaYA---** \| \| \| coexpression\_cluster3\_5::3\_motifs \| PSSM \| 11.69 \| 4256; Node 4 \| **---wwTGTGTGTAya-------** \| \| \| coexpression\_cluster3\_5::2\_motifs \| PSSM \| 8.37 \| 1037; Node 5 \| **--wwwTGTGTGTryATww----** \| \| \| coexpression\_cluster3\_5::4\_motifs \| PSSM \| 9.00 \| 1904; Node 6 \| **--wwwTrTrTGTryATww----** \| \| \| coexpression\_cluster3\_5::5\_motifs \| PSSM \| 8.59 \| 2847; Node 7 \| **--wwwTrTrTGTryATww----** \| \| \| coexpression\_cluster3\_5::6\_motifs \| PSSM \| 8.57 \| 3055; Node 8 \| **--wtaTaTrTrTryATatA---** \| \| \| coexpression\_cluster3\_5::9\_motifs \| PSSM \| 9.97 \| 7311; Node 9 \| **cywtaTrTrTrTryATatasaw** \| \| \| coexpression\_cluster3\_5::10\_motifs \| PSSM \| 9.68 \| 7442 **cluster\_3** Branch Motifs \| \| \| \| \| \| \|; Node \| Consensus \| Logo \| Logo (Reverse) \| Collections \| Matrix (Transfac format) \| IC \| Nb sites; Node 1 \| **wwAGAAwTww** \| \| \| coexpression\_cluster3\_5::2\_motifs \| PSSM \| 8.48 \| 749 **cluster\_4** Branch Motifs \| \| \| \| \| \| \|; Node \| Consensus \| Logo \| Logo (Reverse) \| Collections \| Matrix (Transfac format) \| IC \| Nb sites; Node 1 \| **wwwTCTATAww** \| \| \| coexpression\_cluster3\_5::2\_motifs \| PSSM \| 8.70 \| 625 **cluster\_5** Branch Motifs \| \| \| \| \| \| \|; Node \| Consensus \| Logo \| Logo (Reverse) \| Collections \| Matrix (Transfac format) \| IC \| Nb sites; Singleton \| **aaataataataat** \| \| \| coexpression\_cluster3\_5 : 1 motif \| PSSM \| 13.83 \| 147 **cluster\_6** Branch Motifs \| \| \| \| \| \| \|; Node \| Consensus \| Logo \| Logo (Reverse) \| Collections \| Matrix (Transfac format) \| IC \| Nb sites; Singleton \| **wwAAGTGTdw** \| \| \| coexpression\_cluster3\_5 : 1 motif \| PSSM \| 9.15 \| 186 **cluster\_7** Branch Motifs \| \| \| \| \| \| \|; Node \| Consensus \| Logo \| Logo (Reverse) \| Collections \| Matrix (Transfac format) \| IC \| Nb sites; Singleton \| **aaaCCGwGGraaa** \| \| \| coexpression\_cluster3\_5 : 1 motif \| PSSM \| 10.63 \| 92 **cluster\_8** Branch Motifs \| \| \| \| \| \| \|; Node \| Consensus \| Logo \| Logo (Reverse) \| Collections \| Matrix (Transfac format) \| IC \| Nb sites; Singleton \| **yaaGaCa** \| \| \| coexpression\_cluster3\_5 : 1 motif \| PSSM \| 7.71 \| 360 **cluster\_9** Branch Motifs \| \| \| \| \| \| \|; Node \| Consensus \| Logo \| Logo (Reverse) \| Collections \| Matrix (Transfac format) \| IC \| Nb sites; Singleton \| **raAACGCCww** \| \| \| coexpression\_cluster3\_5 : 1 motif \| PSSM \| 9.08 \| 36 - **Each cluster has a different color.** - **Click on the upper buttons to display separately the information of each cluster.** - **Click on the circles at each tree to display its corresponding merged motifs.** Definitions - **Logo tree.** This results shows the hierarchical tree with its logo alignment of one cluster. The logos are shown in Forward (left logo) and Reverse (right logo) orientation. - **Branch-motifs.** On the trees are displayed the branch number that is the number in which the motifs were incorporated in the tree. The Branch-Motifs table shows the logo in both orientations and the link to the file in TRANSFAC format with the branch-motif. |
|  | **Individual Motif View** Individual Motif View Motif id \| Motif name \| Cluster \| Collection \| Width \| IC \| Number of sites \| Consensus \| Consensus (Rev) \| Logo \| Logo (Rev); coexpression\_cluster3\_5\_m27\_cluster3\_5\_motif27\_m \| cluster3\_5\_motif27\_m \| cluster\_8 \| coexpression\_cluster3\_5 \| 7 \| 7.71 \| 360 \| **TAAGACA** \| **TGTCTTA** \| \|; coexpression\_cluster3\_5\_m7\_cluster3\_5\_motif7\_r \| cluster3\_5\_motif7\_r \| cluster\_1 \| coexpression\_cluster3\_5 \| 15 \| 9.10 \| 1061 \| **++WWTTTGTCTTWWW** \| **wwwAAgACAAAww++** \| \|; coexpression\_cluster3\_5\_m29\_cluster3\_5\_motif29\_m \| cluster3\_5\_motif29\_m \| cluster\_2 \| coexpression\_cluster3\_5 \| 22 \| 7.91 \| 208 \| **+++++++TGTGTGYR+++++++** \| **+++++++YrCACACa+++++++** \| \|; coexpression\_cluster3\_5\_m13\_cluster3\_5\_motif13\_r \| cluster3\_5\_motif13\_r \| cluster\_2 \| coexpression\_cluster3\_5 \| 22 \| 8.70 \| 943 \| **++++++wwATGTGCATww++++** \| **++++WWATGCACATWW++++++** \| \|; coexpression\_cluster3\_5\_m23\_cluster3\_5\_motif23\_m \| cluster3\_5\_motif23\_m \| cluster\_1 \| coexpression\_cluster3\_5 \| 15 \| 12.47 \| 307 \| **+TTATTTATTTTATT** \| **AATAAAATAAATAA+** \| \|; coexpression\_cluster3\_5\_m19\_cluster3\_5\_motif19\_r \| cluster3\_5\_motif19\_r \| cluster\_2 \| coexpression\_cluster3\_5 \| 22 \| 9.73 \| 291 \| **++wwwtrTGTGTGCAwww++++** \| **++++WWWTGCACACAYAWWW++** \| \|; coexpression\_cluster3\_5\_m10\_cluster3\_5\_motif10\_r \| cluster3\_5\_motif10\_r \| cluster\_1 \| coexpression\_cluster3\_5 \| 15 \| 9.46 \| 2214 \| **+WWAAAARAMAATWW** \| **wwATTkTyTTTTww+** \| \|; coexpression\_cluster3\_5\_m12\_cluster3\_5\_motif12\_r \| cluster3\_5\_motif12\_r \| cluster\_1 \| coexpression\_cluster3\_5 \| 15 \| 9.08 \| 812 \| **++wwwTAgACAAAww** \| **WWTTTGTCTAWWW++** \| \|; coexpression\_cluster3\_5\_m8\_cluster3\_5\_motif8\_r \| cluster3\_5\_motif8\_r \| cluster\_1 \| coexpression\_cluster3\_5 \| 15 \| 9.23 \| 1154 \| **+waAAAAgACAwwww** \| **WWWWTGTCTTTTTW+** \| \|; coexpression\_cluster3\_5\_m33\_cluster3\_5\_motif33\_m \| cluster3\_5\_motif33\_m \| cluster\_7 \| coexpression\_cluster3\_5 \| 13 \| 10.63 \| 92 \| **AAAyCGwkrRAAA** \| **TTTYYMWCGRTTT** \| \|; coexpression\_cluster3\_5\_m22\_cluster3\_5\_motif22\_r \| cluster3\_5\_motif22\_r \| cluster\_4 \| coexpression\_cluster3\_5 \| 11 \| 9.44 \| 512 \| **WWWTCTATAWW** \| **wwTATAGAwww** \| \|; coexpression\_cluster3\_5\_m16\_cluster3\_5\_motif16\_r \| cluster3\_5\_motif16\_r \| cluster\_1 \| coexpression\_cluster3\_5 \| 15 \| 9.35 \| 787 \| **+WAAAAAGACAARWW** \| **wwyTTGTcTTTTtw+** \| \|; coexpression\_cluster3\_5\_m25\_cluster3\_5\_motif25\_m \| cluster3\_5\_motif25\_m \| cluster\_2 \| coexpression\_cluster3\_5 \| 22 \| 10.53 \| 863 \| **+++TATWTRYATRTATRY++++** \| **++++ryAtAyAtRyAwATA+++** \| \|; coexpression\_cluster3\_5\_m18\_cluster3\_5\_motif18\_r \| cluster3\_5\_motif18\_r \| cluster\_6 \| coexpression\_cluster3\_5 \| 10 \| 9.15 \| 186 \| **wwAAGTGTdw** \| **WHACACTTWW** \| \|; coexpression\_cluster3\_5\_m20\_cluster3\_5\_motif20\_r \| cluster3\_5\_motif20\_r \| cluster\_4 \| coexpression\_cluster3\_5 \| 11 \| 9.34 \| 113 \| **wtATCTACAw+** \| **+WTGTAGATAW** \| \|; coexpression\_cluster3\_5\_m2\_cluster3\_5\_motif2\_r \| cluster3\_5\_motif2\_r \| cluster\_1 \| coexpression\_cluster3\_5 \| 15 \| 9.09 \| 2162 \| **+wwTATAkAcAAAww** \| **WWTTTGTMTATAWW+** \| \|; coexpression\_cluster3\_5\_m15\_cluster3\_5\_motif15\_r \| cluster3\_5\_motif15\_r \| cluster\_3 \| coexpression\_cluster3\_5 \| 10 \| 9.20 \| 490 \| **wwAGAAATaw** \| **WTATTTCTWW** \| \|; coexpression\_cluster3\_5\_m28\_cluster3\_5\_motif28\_m \| cluster3\_5\_motif28\_m \| cluster\_1 \| coexpression\_cluster3\_5 \| 15 \| 12.50 \| 412 \| **AAAAAAAmrrAAAAA** \| **TTTTTYYKTTTTTTT** \| \|; coexpression\_cluster3\_5\_m26\_cluster3\_5\_motif26\_m \| cluster3\_5\_motif26\_m \| cluster\_1 \| coexpression\_cluster3\_5 \| 15 \| 15.26 \| 138 \| **AAAAAAAAATAATAA** \| **TTATTATTTTTTTTT** \| \|; coexpression\_cluster3\_5\_m5\_cluster3\_5\_motif5\_r \| cluster3\_5\_motif5\_r \| cluster\_1 \| coexpression\_cluster3\_5 \| 15 \| 8.72 \| 1237 \| **+WWWTGTCTTTTTW+** \| **+waAAAAgACAwww+** \| \|; coexpression\_cluster3\_5\_m11\_cluster3\_5\_motif11\_r \| cluster3\_5\_motif11\_r \| cluster\_1 \| coexpression\_cluster3\_5 \| 15 \| 9.22 \| 2343 \| **+wwAATArAmAAAww** \| **WWTTTKTYTATTWW+** \| \|; coexpression\_cluster3\_5\_m24\_cluster3\_5\_motif24\_m \| cluster3\_5\_motif24\_m \| cluster\_2 \| coexpression\_cluster3\_5 \| 22 \| 18.43 \| 150 \| **+++TATATATATATATATA+++** \| **+++TATATATATATATATA+++** \| \|; coexpression\_cluster3\_5\_m32\_cluster3\_5\_motif32\_m \| cluster3\_5\_motif32\_m \| cluster\_1 \| coexpression\_cluster3\_5 \| 15 \| 11.29 \| 114 \| **+TTTYCCTTWTCT++** \| **++AGAwAAGGRAAA+** \| \|; coexpression\_cluster3\_5\_m31\_cluster3\_5\_motif31\_m \| cluster3\_5\_motif31\_m \| cluster\_2 \| coexpression\_cluster3\_5 \| 22 \| 10.83 \| 132 \| **CYBKDYGTGTGTGCAAAWWSAW** \| **wtswwttTGCACACACrhmvrg** \| \|; coexpression\_cluster3\_5\_m9\_cluster3\_5\_motif9\_r \| cluster3\_5\_motif9\_r \| cluster\_2 \| coexpression\_cluster3\_5 \| 22 \| 10.35 \| 3243 \| **+++waTATrTaTATAtw+++++** \| **+++++WATATATAYATATW+++** \| \|; coexpression\_cluster3\_5\_m6\_cluster3\_5\_motif6\_r \| cluster3\_5\_motif6\_r \| cluster\_1 \| coexpression\_cluster3\_5 \| 15 \| 8.74 \| 1043 \| **+WWWWTAGACAAAWW** \| **wwTTTGTcTAwwww+** \| \|; coexpression\_cluster3\_5\_m30\_cluster3\_5\_motif30\_m \| cluster3\_5\_motif30\_m \| cluster\_5 \| coexpression\_cluster3\_5 \| 13 \| 13.83 \| 147 \| **AAATAATAATAAT** \| **ATTATTATTATTT** \| \|; coexpression\_cluster3\_5\_m17\_cluster3\_5\_motif17\_r \| cluster3\_5\_motif17\_r \| cluster\_9 \| coexpression\_cluster3\_5 \| 10 \| 9.08 \| 36 \| **raAACGCCww** \| **WWGGCGTTTY** \| \|; coexpression\_cluster3\_5\_m34\_cluster3\_5\_motif34\_m \| cluster3\_5\_motif34\_m \| cluster\_2 \| coexpression\_cluster3\_5 \| 22 \| 9.42 \| 183 \| **++++ATGTGTGTGCA+++++++** \| **+++++++TGCACACACAT++++** \| \|; coexpression\_cluster3\_5\_m4\_cluster3\_5\_motif4\_r \| cluster3\_5\_motif4\_r \| cluster\_2 \| coexpression\_cluster3\_5 \| 22 \| 9.64 \| 576 \| **++WWWTGTGTGTGCATWW++++** \| **++++wwATGCACACAcawww++** \| \|; coexpression\_cluster3\_5\_m14\_cluster3\_5\_motif14\_r \| cluster3\_5\_motif14\_r \| cluster\_2 \| coexpression\_cluster3\_5 \| 22 \| 8.65 \| 854 \| **+++WWTGTGTGTATW+++++++** \| **+++++++waTACACACAww+++** \| \|; coexpression\_cluster3\_5\_m3\_cluster3\_5\_motif3\_r \| cluster3\_5\_motif3\_r \| cluster\_1 \| coexpression\_cluster3\_5 \| 15 \| 9.60 \| 2821 \| **+WWTATDTYTTTTWW** \| **wwAAAArAhATAww+** \| \|; coexpression\_cluster3\_5\_m1\_cluster3\_5\_motif1\_r \| cluster3\_5\_motif1\_r \| cluster\_1 \| coexpression\_cluster3\_5 \| 15 \| 9.75 \| 3514 \| **+waAAAArAmAAAww** \| **WWTTTKTYTTTTTW+** \| \|; coexpression\_cluster3\_5\_m21\_cluster3\_5\_motif21\_r \| cluster3\_5\_motif21\_r \| cluster\_3 \| coexpression\_cluster3\_5 \| 10 \| 9.07 \| 259 \| **AWAGAATTWW** \| **wwAATTCTwt** \| \| - Click on the column names to change the order of the data. - Write the name of one cluster, collection or a pattern in the *Search* window. **Heatmap View** Heatmap View Distance table \| PDF |

**Additional Files**

#### Exported files

| File | Description |
| --- | --- |
| **Pairwise comparison** | This table shows the pairwise comparison between all the input motifs using different metrics. This is the *compare-matrices* result. |
| **Matrix description** | This table shows information of each input motif. |
| **Cluster-wise Motif Files [zip]** | One TRANSFAC file is generated for each cluster including all the input motifs grouped in the corresponding cluster. |
| **Root motifs** | A file with the root matrix of each cluster. |
| **Motif collections per cluster** | A tab separated file containing the counts of motif of each collection on each cluster. |
| **Clusters** | A tab separated file containing the clusters and their correspondng motifs. |
| **Central motifs IDs** | A tab separated file containing the central motif, that one who is the most similar to all the others motif in the same cluster. |
| **Central motifs (in TRANSFAC format)** | A transfac file containing the central motif, that one who is the most similar to all the others motif within the same cluster. |
| **Input motifs** | The input motifs analyzed with matrix-clustering. |

- **References**
  1. Castro-Mondragon JA et al.  RSAT matrix-clustering: dynamic exploration and redundancy reduction of transcription factor binding motif collections. doi: 10.1093/nar/gkx314; Nucleic Acids Research (2017)
  2. **Last RSAT release:** Nguyen NTT et al. RSAT 2018: regulatory sequence analysis tools 20th anniversary. Nucleic Acids Research (2018).
- **Additional Information**
  - Jacques van Helden course
- **Contact**
  - Jaime Castro-Mondragon - ****
  - Morgane Thomas-Chollier - ****
  - Jacques van Helden - ****
