## Additional file 10 for "Transcriptome analysis reveals a novel DNA element that may interact with chromatin-associated proteins in *Plasmodium berghei* during erythrocytic development": matrix-clustering_SUMMARY_cluster3_7.html

matrix-clustering coexpression\_cluster3\_7

### RSAT - matrix-clustering result

##### Analysis: coexpression\_cluster3\_7 (08/09/2023 16:02)

##### Command

```
matrix-clustering  -v 1 -max_matrices 300 -matrix coexpression_cluster3_7 $RSAT/public_html/tmp/www-data/2023/09/08/matrix-clustering_2023-09-08.160225_1025zP/matrix-clustering_query_matrices.transfac transfac -hclust_method average -calc sum -title coexpression_cluster3_7 -metric_build_tree Ncor -lth w 5 -lth cor 0.6 -lth Ncor 0.4 -quick -label_in_tree name -return json,heatmap -o $RSAT/public_html/tmp/www-data/2023/09/08/matrix-clustering_2023-09-08.160225_1025zP/matrix-clustering
```

#### Results Summary

|  |  |  |  |  |  |
| --- | --- | --- | --- | --- | --- |
| Nb Input motifs | Nb Input collections | Nb Clusters Found | Download   Root motifs | Complete results [zip] | Parameters used Linkage method \| Similarity metric \| Thresholds to partition the tree |
| 30 | 1 | 9 | **Download** | Download | average \| Ncor \| **Ncor** = 0.4 **cor** = 0.6 |

|  |  |  |  |  |  |  |
| --- | --- | --- | --- | --- | --- | --- |
|  | **Logo Forest (dynamic browsing)**  **Logo Forest (rapid overview - low image quality)**  cluster\_1  cluster\_2  cluster\_3  cluster\_4  cluster\_5  cluster\_6  cluster\_7  cluster\_8  cluster\_9 |  |  |  |  |  |
|  | **Clusters Summary** Clusters Summary Table (Root Motif \| Root Motif (Reverse) \| Cluster ID \| # Motifs \| Motif\_Name::Collection \| Number of Motifs by Collection \| Root Motif (transfac Format)) \| | cluster\_1 | 9 | cluster3\_7\_motif19\_m::coexpression\_cluster3\_7:: cluster3\_7\_motif20\_m::coexpression\_cluster3\_7:: cluster3\_7\_motif5\_r::coexpression\_cluster3\_7:: cluster3\_7\_motif6\_r::coexpression\_cluster3\_7:: cluster3\_7\_motif1\_r::coexpression\_cluster3\_7:: cluster3\_7\_motif2\_r::coexpression\_cluster3\_7:: cluster3\_7\_motif7\_r::coexpression\_cluster3\_7:: cluster3\_7\_motif12\_r::coexpression\_cluster3\_7:: cluster3\_7\_motif3\_r::coexpression\_cluster3\_7:: | coexpression\_cluster3\_7::9\_motifs | Show matrix |
| cluster\_2 \| 3 \| cluster3\_7\_motif11\_r::coexpression\_cluster3\_7:: cluster3\_7\_motif23\_m::coexpression\_cluster3\_7:: cluster3\_7\_motif27\_m::coexpression\_cluster3\_7:: \| coexpression\_cluster3\_7::3\_motifs \| Show matrix \| \| |  |  |  |  |  |  |
| cluster\_3 \| 4 \| cluster3\_7\_motif18\_m::coexpression\_cluster3\_7:: cluster3\_7\_motif21\_m::coexpression\_cluster3\_7:: cluster3\_7\_motif4\_r::coexpression\_cluster3\_7:: cluster3\_7\_motif9\_r::coexpression\_cluster3\_7:: \| coexpression\_cluster3\_7::4\_motifs \| Show matrix \| \| |  |  |  |  |  |  |
| cluster\_4 | 5 | cluster3\_7\_motif26\_m::coexpression\_cluster3\_7:: cluster3\_7\_motif17\_m::coexpression\_cluster3\_7:: cluster3\_7\_motif25\_m::coexpression\_cluster3\_7:: cluster3\_7\_motif15\_r::coexpression\_cluster3\_7:: cluster3\_7\_motif8\_r::coexpression\_cluster3\_7:: | coexpression\_cluster3\_7::5\_motifs | Show matrix | | |
| cluster\_5 | 5 | cluster3\_7\_motif28\_m::coexpression\_cluster3\_7:: cluster3\_7\_motif29\_m::coexpression\_cluster3\_7:: cluster3\_7\_motif24\_m::coexpression\_cluster3\_7:: cluster3\_7\_motif10\_r::coexpression\_cluster3\_7:: cluster3\_7\_motif22\_m::coexpression\_cluster3\_7:: | coexpression\_cluster3\_7::5\_motifs | Show matrix | | |
| cluster\_6 | 1 | cluster3\_7\_motif14\_r::coexpression\_cluster3\_7 | coexpression\_cluster3\_7 : 1 motif | Show matrix | | |
| cluster\_7 | 1 | cluster3\_7\_motif16\_r::coexpression\_cluster3\_7 | coexpression\_cluster3\_7 : 1 motif | Show matrix | | |
| cluster\_8 | 1 | cluster3\_7\_motif13\_r::coexpression\_cluster3\_7 | coexpression\_cluster3\_7 : 1 motif | Show matrix | | |
| cluster\_9 | 1 | cluster3\_7\_motif30\_m::coexpression\_cluster3\_7 | coexpression\_cluster3\_7 : 1 motif | Show matrix | | |

- Note: by default the table displays the first 50 clusters. Click on the *Show X entries* button to display more (or less) clusters.
- Click on the column names to change the order of the data.
- Use the *Search* window to find your motif (s) of interest.

| **Individual Cluster View** Individual Cluster View **Show All** **Hide All** **cluster\_1** **cluster\_2** **cluster\_3** **cluster\_4** **cluster\_5** **cluster\_6** **cluster\_7** **cluster\_8** **cluster\_9** **Display Logo Trees** **cluster\_1** **cluster\_2** **cluster\_3** **cluster\_4** **cluster\_5** **cluster\_6** **cluster\_7** **cluster\_8** **cluster\_9** **Display Branch-Motifs** **cluster\_1** Branch Motifs \| \| \| \| \| \| \|; Node \| Consensus \| Logo \| Logo (Reverse) \| Collections \| Matrix (Transfac format) \| IC \| Nb sites; Node 1 \| **wwAamTAGCTAtTTww-** \| \| \| coexpression\_cluster3\_7::2\_motifs \| PSSM \| 9.27 \| 579; Node 2 \| **-wwttTAGCTAktTww-** \| \| \| coexpression\_cluster3\_7::2\_motifs \| PSSM \| 9.22 \| 503; Node 3 \| **wwAamTAGCTAktTwww** \| \| \| coexpression\_cluster3\_7::3\_motifs \| PSSM \| 9.41 \| 772; Node 4 \| **-wwwtTAGCTAktTww-** \| \| \| coexpression\_cluster3\_7::3\_motifs \| PSSM \| 9.26 \| 732; Node 5 \| **----wtArCTAGCyAw-** \| \| \| coexpression\_cluster3\_7::2\_motifs \| PSSM \| 8.74 \| 177; Node 6 \| **wwwwhTAGCTAktTwww** \| \| \| coexpression\_cluster3\_7::6\_motifs \| PSSM \| 9.26 \| 1504; Node 7 \| **wwwwhTAGCTAktTwww** \| \| \| coexpression\_cluster3\_7::7\_motifs \| PSSM \| 9.13 \| 1628; Node 8 \| **wwwwhTAGCTAkyTwww** \| \| \| coexpression\_cluster3\_7::9\_motifs \| PSSM \| 8.86 \| 1805 **cluster\_2** Branch Motifs \| \| \| \| \| \| \|; Node \| Consensus \| Logo \| Logo (Reverse) \| Collections \| Matrix (Transfac format) \| IC \| Nb sites; Node 1 \| **--TTATkAACAaktmAT** \| \| \| coexpression\_cluster3\_7::2\_motifs \| PSSM \| 14.41 \| 91; Node 2 \| **wwwTATGAACAwwtmAT** \| \| \| coexpression\_cluster3\_7::3\_motifs \| PSSM \| 12.09 \| 584 **cluster\_3** Branch Motifs \| \| \| \| \| \| \|; Node \| Consensus \| Logo \| Logo (Reverse) \| Collections \| Matrix (Transfac format) \| IC \| Nb sites; Node 1 \| **TAYAYAYAyATATATA** \| \| \| coexpression\_cluster3\_7::2\_motifs \| PSSM \| 13.58 \| 392; Node 2 \| **-wwATAgACAwwww--** \| \| \| coexpression\_cluster3\_7::2\_motifs \| PSSM \| 8.50 \| 846; Node 3 \| **TawATAkACAwawaTA** \| \| \| coexpression\_cluster3\_7::4\_motifs \| PSSM \| 11.03 \| 1238 **cluster\_4** Branch Motifs \| \| \| \| \| \| \|; Node \| Consensus \| Logo \| Logo (Reverse) \| Collections \| Matrix (Transfac format) \| IC \| Nb sites; Node 1 \| **--AAAAAAAArAwAAAA** \| \| \| coexpression\_cluster3\_7::2\_motifs \| PSSM \| 12.73 \| 332; Node 2 \| **-wwaAAAAAArww----** \| \| \| coexpression\_cluster3\_7::2\_motifs \| PSSM \| 8.14 \| 5987; Node 3 \| **-wwaAAAAAArwwAAAA** \| \| \| coexpression\_cluster3\_7::4\_motifs \| PSSM \| 11.87 \| 6318; Node 4 \| **AawaAAAAAArwwAAAA** \| \| \| coexpression\_cluster3\_7::5\_motifs \| PSSM \| 12.47 \| 6402 **cluster\_5** Branch Motifs \| \| \| \| \| \| \|; Node \| Consensus \| Logo \| Logo (Reverse) \| Collections \| Matrix (Transfac format) \| IC \| Nb sites; Node 1 \| **----AtdTGTGTGyat------** \| \| \| coexpression\_cluster3\_7::2\_motifs \| PSSM \| 8.69 \| 501; Node 2 \| **cyykdyGTGTGTGYraawwgaw** \| \| \| coexpression\_cluster3\_7::2\_motifs \| PSSM \| 10.47 \| 108; Node 3 \| **----AtdTGTGTGyat------** \| \| \| coexpression\_cluster3\_7::3\_motifs \| PSSM \| 8.72 \| 591; Node 4 \| **cyykatdTGTGTGyawawwgaw** \| \| \| coexpression\_cluster3\_7::5\_motifs \| PSSM \| 8.94 \| 699 **cluster\_6** Branch Motifs \| \| \| \| \| \| \|; Node \| Consensus \| Logo \| Logo (Reverse) \| Collections \| Matrix (Transfac format) \| IC \| Nb sites; Singleton \| **wwTACAAcTww** \| \| \| coexpression\_cluster3\_7 : 1 motif \| PSSM \| 7.40 \| 626 **cluster\_7** Branch Motifs \| \| \| \| \| \| \|; Node \| Consensus \| Logo \| Logo (Reverse) \| Collections \| Matrix (Transfac format) \| IC \| Nb sites; Singleton \| **wwACAAGAww** \| \| \| coexpression\_cluster3\_7 : 1 motif \| PSSM \| 9.37 \| 59 **cluster\_8** Branch Motifs \| \| \| \| \| \| \|; Node \| Consensus \| Logo \| Logo (Reverse) \| Collections \| Matrix (Transfac format) \| IC \| Nb sites; Singleton \| **wwACATTGhw** \| \| \| coexpression\_cluster3\_7 : 1 motif \| PSSM \| 8.95 \| 103 **cluster\_9** Branch Motifs \| \| \| \| \| \| \|; Node \| Consensus \| Logo \| Logo (Reverse) \| Collections \| Matrix (Transfac format) \| IC \| Nb sites; Singleton \| **CCCCSC** \| \| \| coexpression\_cluster3\_7 : 1 motif \| PSSM \| 7.05 \| 19 - **Each cluster has a different color.** - **Click on the upper buttons to display separately the information of each cluster.** - **Click on the circles at each tree to display its corresponding merged motifs.** Definitions - **Logo tree.** This results shows the hierarchical tree with its logo alignment of one cluster. The logos are shown in Forward (left logo) and Reverse (right logo) orientation. - **Branch-motifs.** On the trees are displayed the branch number that is the number in which the motifs were incorporated in the tree. The Branch-Motifs table shows the logo in both orientations and the link to the file in TRANSFAC format with the branch-motif. |
|  | **Individual Motif View** Individual Motif View Motif id \| Motif name \| Cluster \| Collection \| Width \| IC \| Number of sites \| Consensus \| Consensus (Rev) \| Logo \| Logo (Rev); coexpression\_cluster3\_7\_m3\_cluster3\_7\_motif3\_r \| cluster3\_7\_motif3\_r \| cluster\_1 \| coexpression\_cluster3\_7 \| 17 \| 9.21 \| 299 \| **+WWTTTAGCTAKTTWW+** \| **+wwAamTAGCTAAAww+** \| \|; coexpression\_cluster3\_7\_m12\_cluster3\_7\_motif12\_r \| cluster3\_7\_motif12\_r \| cluster\_1 \| coexpression\_cluster3\_7 \| 17 \| 9.85 \| 204 \| **+WWWMTAGCTAKTTWW+** \| **+wwAamTAGCTAkwww+** \| \|; coexpression\_cluster3\_7\_m27\_cluster3\_7\_motif27\_m \| cluster3\_7\_motif27\_m \| cluster\_2 \| coexpression\_cluster3\_7 \| 17 \| 13.59 \| 47 \| **++WTATKAACAaktmA+** \| **+TKAMTTGTTMATAW++** \| \|; coexpression\_cluster3\_7\_m30\_cluster3\_7\_motif30\_m \| cluster3\_7\_motif30\_m \| cluster\_9 \| coexpression\_cluster3\_7 \| 6 \| 7.05 \| 19 \| **MCCCCC** \| **GGGGGK** \| \|; coexpression\_cluster3\_7\_m28\_cluster3\_7\_motif28\_m \| cluster3\_7\_motif28\_m \| cluster\_5 \| coexpression\_cluster3\_7 \| 22 \| 10.97 \| 88 \| **CYBKDYGTGTGTGCAAAWWSAW** \| **wtswwttTGCACACACrhmvrg** \| \|; coexpression\_cluster3\_7\_m9\_cluster3\_7\_motif9\_r \| cluster3\_7\_motif9\_r \| cluster\_3 \| coexpression\_cluster3\_7 \| 16 \| 8.92 \| 417 \| **+wwATAkACATww+++** \| **+++WWATGTMTATWW+** \| \|; coexpression\_cluster3\_7\_m21\_cluster3\_7\_motif21\_m \| cluster3\_7\_motif21\_m \| cluster\_3 \| coexpression\_cluster3\_7 \| 16 \| 12.02 \| 317 \| **+RYAYAyAyATATATA** \| **TATATATRTRTRTRY+** \| \|; coexpression\_cluster3\_7\_m10\_cluster3\_7\_motif10\_r \| cluster3\_7\_motif10\_r \| cluster\_5 \| coexpression\_cluster3\_7 \| 22 \| 7.77 \| 403 \| **+++++TWTGTGTGYAT++++++** \| **++++++atRCACACAwa+++++** \| \|; coexpression\_cluster3\_7\_m17\_cluster3\_7\_motif17\_m \| cluster3\_7\_motif17\_m \| cluster\_4 \| coexpression\_cluster3\_7 \| 17 \| 15.84 \| 132 \| **++AAAAAAAWAATAAAA** \| **TTTTATTWTTTTTTT++** \| \|; coexpression\_cluster3\_7\_m2\_cluster3\_7\_motif2\_r \| cluster3\_7\_motif2\_r \| cluster\_1 \| coexpression\_cluster3\_7 \| 17 \| 9.39 \| 331 \| **WWAWMTAGCTATTTWW+** \| **+wwAAATAGCTAkwTww** \| \|; coexpression\_cluster3\_7\_m14\_cluster3\_7\_motif14\_r \| cluster3\_7\_motif14\_r \| cluster\_6 \| coexpression\_cluster3\_7 \| 11 \| 7.40 \| 626 \| **wwTACAAcTww** \| **WWAGTTGTAWW** \| \|; coexpression\_cluster3\_7\_m19\_cluster3\_7\_motif19\_m \| cluster3\_7\_motif19\_m \| cluster\_1 \| coexpression\_cluster3\_7 \| 17 \| 8.46 \| 91 \| **++++wTAGCTAGCTAw+** \| **+WTAGCTAGCTAW++++** \| \|; coexpression\_cluster3\_7\_m18\_cluster3\_7\_motif18\_m \| cluster3\_7\_motif18\_m \| cluster\_3 \| coexpression\_cluster3\_7 \| 16 \| 18.23 \| 75 \| **TATATATATATATATA** \| **TATATATATATATATA** \| \|; coexpression\_cluster3\_7\_m20\_cluster3\_7\_motif20\_m \| cluster3\_7\_motif20\_m \| cluster\_1 \| coexpression\_cluster3\_7 \| 17 \| 9.57 \| 86 \| **+++++yArCTAGCyAw+** \| **+WTRGCTAGYTR+++++** \| \|; coexpression\_cluster3\_7\_m11\_cluster3\_7\_motif11\_r \| cluster3\_7\_motif11\_r \| cluster\_2 \| coexpression\_cluster3\_7 \| 17 \| 9.03 \| 493 \| **WWTGTTCATAWWW++++** \| **++++wwwTATGAACAww** \| \|; coexpression\_cluster3\_7\_m26\_cluster3\_7\_motif26\_m \| cluster3\_7\_motif26\_m \| cluster\_4 \| coexpression\_cluster3\_7 \| 17 \| 11.17 \| 84 \| **CTTTTTTTTCATT++++** \| **++++AATGAAAAAAAAG** \| \|; coexpression\_cluster3\_7\_m25\_cluster3\_7\_motif25\_m \| cluster3\_7\_motif25\_m \| cluster\_4 \| coexpression\_cluster3\_7 \| 17 \| 13.15 \| 200 \| **++AAAAAAAArRRAAAA** \| **TTTTYYYTTTTTTTT++** \| \|; coexpression\_cluster3\_7\_m4\_cluster3\_7\_motif4\_r \| cluster3\_7\_motif4\_r \| cluster\_3 \| coexpression\_cluster3\_7 \| 16 \| 8.99 \| 429 \| **+wwATAgACAAwww++** \| **++WWWTTGTCTATWW+** \| \|; coexpression\_cluster3\_7\_m16\_cluster3\_7\_motif16\_r \| cluster3\_7\_motif16\_r \| cluster\_7 \| coexpression\_cluster3\_7 \| 10 \| 9.37 \| 59 \| **wwACAAGAww** \| **WWTCTTGTWW** \| \|; coexpression\_cluster3\_7\_m7\_cluster3\_7\_motif7\_r \| cluster3\_7\_motif7\_r \| cluster\_1 \| coexpression\_cluster3\_7 \| 17 \| 9.45 \| 229 \| **++wwdTAGCTAkTTww+** \| **+WWAAMTAGCTAHWW++** \| \|; coexpression\_cluster3\_7\_m29\_cluster3\_7\_motif29\_m \| cluster3\_7\_motif29\_m \| cluster\_5 \| coexpression\_cluster3\_7 \| 22 \| 12.48 \| 21 \| **WKMYDYGTGTGTGYGSAWWABA** \| **TvtwwtsCRCACACACrhrkmw** \| \|; coexpression\_cluster3\_7\_m8\_cluster3\_7\_motif8\_r \| cluster3\_7\_motif8\_r \| cluster\_4 \| coexpression\_cluster3\_7 \| 17 \| 8.95 \| 535 \| **+wwAAgACAAAww++++** \| **++++WWTTTGTCTTWW+** \| \|; coexpression\_cluster3\_7\_m13\_cluster3\_7\_motif13\_r \| cluster3\_7\_motif13\_r \| cluster\_8 \| coexpression\_cluster3\_7 \| 10 \| 8.95 \| 103 \| **wwACATTGhw** \| **WDCAATGTWW** \| \|; coexpression\_cluster3\_7\_m6\_cluster3\_7\_motif6\_r \| cluster3\_7\_motif6\_r \| cluster\_1 \| coexpression\_cluster3\_7 \| 17 \| 10.00 \| 193 \| **wwAamTAGCTAgyTwww** \| **WWWARCTAGCTAKTTWW** \| \|; coexpression\_cluster3\_7\_m1\_cluster3\_7\_motif1\_r \| cluster3\_7\_motif1\_r \| cluster\_1 \| coexpression\_cluster3\_7 \| 17 \| 9.70 \| 248 \| **WWAAMTAGCTAKTTWW+** \| **+wwAamTAGCTAktTww** \| \|; coexpression\_cluster3\_7\_m15\_cluster3\_7\_motif15\_r \| cluster3\_7\_motif15\_r \| cluster\_4 \| coexpression\_cluster3\_7 \| 17 \| 8.24 \| 5452 \| **++waAAAAAArww++++** \| **++++WWYTTTTTTTW++** \| \|; coexpression\_cluster3\_7\_m5\_cluster3\_7\_motif5\_r \| cluster3\_7\_motif5\_r \| cluster\_1 \| coexpression\_cluster3\_7 \| 17 \| 10.55 \| 124 \| **+WWTTTGGCTAKTTWW+** \| **+wwAAmTAGCcAAAww+** \| \|; coexpression\_cluster3\_7\_m23\_cluster3\_7\_motif23\_m \| cluster3\_7\_motif23\_m \| cluster\_2 \| coexpression\_cluster3\_7 \| 17 \| 14.71 \| 44 \| **++TTATkAACAwGwmAT** \| **ATKWCWTGTTMATAA++** \| \|; coexpression\_cluster3\_7\_m24\_cluster3\_7\_motif24\_m \| cluster3\_7\_motif24\_m \| cluster\_5 \| coexpression\_cluster3\_7 \| 22 \| 8.57 \| 90 \| **+++++++TGTGTGYA+++++++** \| **+++++++TrCACACA+++++++** \| \|; coexpression\_cluster3\_7\_m22\_cluster3\_7\_motif22\_m \| cluster3\_7\_motif22\_m \| cluster\_5 \| coexpression\_cluster3\_7 \| 22 \| 10.11 \| 98 \| **++++ATGTGTGTGya+++++++** \| **+++++++TRCACACACAT++++** \| \| - Click on the column names to change the order of the data. - Write the name of one cluster, collection or a pattern in the *Search* window. **Heatmap View** Heatmap View Distance table \| PDF |

**Additional Files**

#### Exported files

| File | Description |
| --- | --- |
| **Pairwise comparison** | This table shows the pairwise comparison between all the input motifs using different metrics. This is the *compare-matrices* result. |
| **Matrix description** | This table shows information of each input motif. |
| **Cluster-wise Motif Files [zip]** | One TRANSFAC file is generated for each cluster including all the input motifs grouped in the corresponding cluster. |
| **Root motifs** | A file with the root matrix of each cluster. |
| **Motif collections per cluster** | A tab separated file containing the counts of motif of each collection on each cluster. |
| **Clusters** | A tab separated file containing the clusters and their correspondng motifs. |
| **Central motifs IDs** | A tab separated file containing the central motif, that one who is the most similar to all the others motif in the same cluster. |
| **Central motifs (in TRANSFAC format)** | A transfac file containing the central motif, that one who is the most similar to all the others motif within the same cluster. |
| **Input motifs** | The input motifs analyzed with matrix-clustering. |

- **References**
  1. Castro-Mondragon JA et al.  RSAT matrix-clustering: dynamic exploration and redundancy reduction of transcription factor binding motif collections. doi: 10.1093/nar/gkx314; Nucleic Acids Research (2017)
  2. **Last RSAT release:** Nguyen NTT et al. RSAT 2018: regulatory sequence analysis tools 20th anniversary. Nucleic Acids Research (2018).
- **Additional Information**
  - Jacques van Helden course
- **Contact**
  - Jaime Castro-Mondragon - ****
  - Morgane Thomas-Chollier - ****
  - Jacques van Helden - ****
