## Additional file 10 for "Transcriptome analysis reveals a novel DNA element that may interact with chromatin-associated proteins in *Plasmodium berghei* during erythrocytic development": matrix-clustering_SUMMARY_cluster3_8.html

matrix-clustering coexpression\_cluster3\_8

### RSAT - matrix-clustering result

##### Analysis: coexpression\_cluster3\_8 (08/09/2023 16:09)

##### Command

```
matrix-clustering  -v 1 -max_matrices 300 -matrix coexpression_cluster3_8 $RSAT/public_html/tmp/www-data/2023/09/08/matrix-clustering_2023-09-08.160939_KQzGMX/matrix-clustering_query_matrices.transfac transfac -hclust_method average -calc sum -title coexpression_cluster3_8 -metric_build_tree Ncor -lth w 5 -lth cor 0.6 -lth Ncor 0.4 -quick -label_in_tree name -return json,heatmap -o $RSAT/public_html/tmp/www-data/2023/09/08/matrix-clustering_2023-09-08.160939_KQzGMX/matrix-clustering
```

#### Results Summary

|  |  |  |  |  |  |
| --- | --- | --- | --- | --- | --- |
| Nb Input motifs | Nb Input collections | Nb Clusters Found | Download   Root motifs | Complete results [zip] | Parameters used Linkage method \| Similarity metric \| Thresholds to partition the tree |
| 30 | 1 | 8 | **Download** | Download | average \| Ncor \| **Ncor** = 0.4 **cor** = 0.6 |

|  |  |  |  |  |  |  |
| --- | --- | --- | --- | --- | --- | --- |
|  | **Logo Forest (dynamic browsing)**  **Logo Forest (rapid overview - low image quality)**  cluster\_1  cluster\_2  cluster\_3  cluster\_4  cluster\_5  cluster\_6  cluster\_7  cluster\_8 |  |  |  |  |  |
|  | **Clusters Summary** Clusters Summary Table (Root Motif \| Root Motif (Reverse) \| Cluster ID \| # Motifs \| Motif\_Name::Collection \| Number of Motifs by Collection \| Root Motif (transfac Format)) \| | cluster\_1 | 13 | cluster3\_8\_motif7\_r::coexpression\_cluster3\_8:: cluster3\_8\_motif12\_r::coexpression\_cluster3\_8:: cluster3\_8\_motif13\_r::coexpression\_cluster3\_8:: cluster3\_8\_motif11\_r::coexpression\_cluster3\_8:: cluster3\_8\_motif1\_r::coexpression\_cluster3\_8:: cluster3\_8\_motif10\_r::coexpression\_cluster3\_8:: cluster3\_8\_motif19\_m::coexpression\_cluster3\_8:: cluster3\_8\_motif5\_r::coexpression\_cluster3\_8:: cluster3\_8\_motif20\_m::coexpression\_cluster3\_8:: cluster3\_8\_motif16\_r::coexpression\_cluster3\_8:: cluster3\_8\_motif4\_r::coexpression\_cluster3\_8:: cluster3\_8\_motif3\_r::coexpression\_cluster3\_8:: cluster3\_8\_motif9\_r::coexpression\_cluster3\_8:: | coexpression\_cluster3\_8::13\_motifs | Show matrix |
| cluster\_2 \| 7 \| cluster3\_8\_motif22\_m::coexpression\_cluster3\_8:: cluster3\_8\_motif25\_m::coexpression\_cluster3\_8:: cluster3\_8\_motif2\_r::coexpression\_cluster3\_8:: cluster3\_8\_motif6\_r::coexpression\_cluster3\_8:: cluster3\_8\_motif24\_m::coexpression\_cluster3\_8:: cluster3\_8\_motif14\_r::coexpression\_cluster3\_8:: cluster3\_8\_motif8\_r::coexpression\_cluster3\_8:: \| coexpression\_cluster3\_8::7\_motifs \| Show matrix \| \| |  |  |  |  |  |  |
| cluster\_3 \| 4 \| cluster3\_8\_motif18\_r::coexpression\_cluster3\_8:: cluster3\_8\_motif21\_m::coexpression\_cluster3\_8:: cluster3\_8\_motif27\_m::coexpression\_cluster3\_8:: cluster3\_8\_motif28\_m::coexpression\_cluster3\_8:: \| coexpression\_cluster3\_8::4\_motifs \| Show matrix \| \| |  |  |  |  |  |  |
| cluster\_4 | 2 | cluster3\_8\_motif26\_m::coexpression\_cluster3\_8:: cluster3\_8\_motif29\_m::coexpression\_cluster3\_8:: | coexpression\_cluster3\_8::2\_motifs | Show matrix | | |
| cluster\_5 | 1 | cluster3\_8\_motif23\_m::coexpression\_cluster3\_8 | coexpression\_cluster3\_8 : 1 motif | Show matrix | | |
| cluster\_6 | 1 | cluster3\_8\_motif15\_r::coexpression\_cluster3\_8 | coexpression\_cluster3\_8 : 1 motif | Show matrix | | |
| cluster\_7 | 1 | cluster3\_8\_motif17\_r::coexpression\_cluster3\_8 | coexpression\_cluster3\_8 : 1 motif | Show matrix | | |
| cluster\_8 | 1 | cluster3\_8\_motif30\_m::coexpression\_cluster3\_8 | coexpression\_cluster3\_8 : 1 motif | Show matrix | | |

- Note: by default the table displays the first 50 clusters. Click on the *Show X entries* button to display more (or less) clusters.
- Click on the column names to change the order of the data.
- Use the *Search* window to find your motif (s) of interest.

| **Individual Cluster View** Individual Cluster View **Show All** **Hide All** **cluster\_1** **cluster\_2** **cluster\_3** **cluster\_4** **cluster\_5** **cluster\_6** **cluster\_7** **cluster\_8** **Display Logo Trees** **cluster\_1** **cluster\_2** **cluster\_3** **cluster\_4** **cluster\_5** **cluster\_6** **cluster\_7** **cluster\_8** **Display Branch-Motifs** **cluster\_1** Branch Motifs \| \| \| \| \| \| \|; Node \| Consensus \| Logo \| Logo (Reverse) \| Collections \| Matrix (Transfac format) \| IC \| Nb sites; Node 1 \| **----wwTATAgAcAAAwww** \| \| \| coexpression\_cluster3\_8::2\_motifs \| PSSM \| 9.40 \| 1719; Node 2 \| **---awATACACACAtaw--** \| \| \| coexpression\_cluster3\_8::2\_motifs \| PSSM \| 8.82 \| 2139; Node 3 \| **----wwTATAgAcAAAwww** \| \| \| coexpression\_cluster3\_8::3\_motifs \| PSSM \| 9.19 \| 2293; Node 4 \| **----wwTATAgACAAAwww** \| \| \| coexpression\_cluster3\_8::4\_motifs \| PSSM \| 8.90 \| 2875; Node 5 \| **---AtaTATAtAyATAta-** \| \| \| coexpression\_cluster3\_8::2\_motifs \| PSSM \| 10.62 \| 2458; Node 6 \| **--wawATACACAcAtaw--** \| \| \| coexpression\_cluster3\_8::3\_motifs \| PSSM \| 8.97 \| 2496; Node 7 \| **---AwaTATAkAyATAta-** \| \| \| coexpression\_cluster3\_8::3\_motifs \| PSSM \| 10.23 \| 3204; Node 8 \| **waTatAtACACAcATaw--** \| \| \| coexpression\_cluster3\_8::4\_motifs \| PSSM \| 9.91 \| 3063; Node 9 \| **----wwwwTAgAcAAAwww** \| \| \| coexpression\_cluster3\_8::5\_motifs \| PSSM \| 8.60 \| 4053; Node 10 \| **waTatAtACACAcATaw--** \| \| \| coexpression\_cluster3\_8::5\_motifs \| PSSM \| 9.94 \| 3176; Node 11 \| **waTatATAyAyAcATAwa-** \| \| \| coexpression\_cluster3\_8::8\_motifs \| PSSM \| 9.93 \| 6380; Node 12 \| **waTawaTAtAbAcAwAwww** \| \| \| coexpression\_cluster3\_8::13\_motifs \| PSSM \| 9.30 \| 10433 **cluster\_2** Branch Motifs \| \| \| \| \| \| \|; Node \| Consensus \| Logo \| Logo (Reverse) \| Collections \| Matrix (Transfac format) \| IC \| Nb sites; Node 1 \| **---wwAAAmAAAAww-** \| \| \| coexpression\_cluster3\_8::2\_motifs \| PSSM \| 9.18 \| 4121; Node 2 \| **wwaAAArAmAAAAww-** \| \| \| coexpression\_cluster3\_8::2\_motifs \| PSSM \| 9.71 \| 3612; Node 3 \| **waaAAArAmAAAAww-** \| \| \| coexpression\_cluster3\_8::3\_motifs \| PSSM \| 9.52 \| 3954; Node 4 \| **---wwAAAmAAAAwwA** \| \| \| coexpression\_cluster3\_8::3\_motifs \| PSSM \| 10.59 \| 4246; Node 5 \| **waawAAAAmAAAAwwA** \| \| \| coexpression\_cluster3\_8::6\_motifs \| PSSM \| 10.86 \| 8200; Node 6 \| **waawAAAAmAAAAwwA** \| \| \| coexpression\_cluster3\_8::7\_motifs \| PSSM \| 10.77 \| 8367 **cluster\_3** Branch Motifs \| \| \| \| \| \| \|; Node \| Consensus \| Logo \| Logo (Reverse) \| Collections \| Matrix (Transfac format) \| IC \| Nb sites; Node 1 \| **GtaTGTACAta** \| \| \| coexpression\_cluster3\_8::2\_motifs \| PSSM \| 10.33 \| 327; Node 2 \| **-TATGCAyATA** \| \| \| coexpression\_cluster3\_8::2\_motifs \| PSSM \| 10.75 \| 179; Node 3 \| **GTATGyAYAta** \| \| \| coexpression\_cluster3\_8::4\_motifs \| PSSM \| 9.40 \| 506 **cluster\_4** Branch Motifs \| \| \| \| \| \| \|; Node \| Consensus \| Logo \| Logo (Reverse) \| Collections \| Matrix (Transfac format) \| IC \| Nb sites; Node 1 \| **wtywwttyrCACACACrhvrrk** \| \| \| coexpression\_cluster3\_8::2\_motifs \| PSSM \| 10.26 \| 180 **cluster\_5** Branch Motifs \| \| \| \| \| \| \|; Node \| Consensus \| Logo \| Logo (Reverse) \| Collections \| Matrix (Transfac format) \| IC \| Nb sites; Singleton \| **aStaGmCarR** \| \| \| coexpression\_cluster3\_8 : 1 motif \| PSSM \| 7.06 \| 190 **cluster\_6** Branch Motifs \| \| \| \| \| \| \|; Node \| Consensus \| Logo \| Logo (Reverse) \| Collections \| Matrix (Transfac format) \| IC \| Nb sites; Singleton \| **awACATGTwt** \| \| \| coexpression\_cluster3\_8 : 1 motif \| PSSM \| 9.24 \| 222 **cluster\_7** Branch Motifs \| \| \| \| \| \| \|; Node \| Consensus \| Logo \| Logo (Reverse) \| Collections \| Matrix (Transfac format) \| IC \| Nb sites; Singleton \| **wwAGAATGtw** \| \| \| coexpression\_cluster3\_8 : 1 motif \| PSSM \| 9.15 \| 104 **cluster\_8** Branch Motifs \| \| \| \| \| \| \|; Node \| Consensus \| Logo \| Logo (Reverse) \| Collections \| Matrix (Transfac format) \| IC \| Nb sites; Singleton \| **KasSmttkBCCSSCC** \| \| \| coexpression\_cluster3\_8 : 1 motif \| PSSM \| 10.82 \| 18 - **Each cluster has a different color.** - **Click on the upper buttons to display separately the information of each cluster.** - **Click on the circles at each tree to display its corresponding merged motifs.** Definitions - **Logo tree.** This results shows the hierarchical tree with its logo alignment of one cluster. The logos are shown in Forward (left logo) and Reverse (right logo) orientation. - **Branch-motifs.** On the trees are displayed the branch number that is the number in which the motifs were incorporated in the tree. The Branch-Motifs table shows the logo in both orientations and the link to the file in TRANSFAC format with the branch-motif. |
|  | **Individual Motif View** Individual Motif View Motif id \| Motif name \| Cluster \| Collection \| Width \| IC \| Number of sites \| Consensus \| Consensus (Rev) \| Logo \| Logo (Rev); coexpression\_cluster3\_8\_m20\_cluster3\_8\_motif20\_m \| cluster3\_8\_motif20\_m \| cluster\_1 \| coexpression\_cluster3\_8 \| 19 \| 11.04 \| 113 \| **+++++RTATGTGTGTGT++** \| **++ACACACACATAy+++++** \| \|; coexpression\_cluster3\_8\_m16\_cluster3\_8\_motif16\_r \| cluster3\_8\_motif16\_r \| cluster\_1 \| coexpression\_cluster3\_8 \| 19 \| 11.27 \| 567 \| **AWATRTGTGTATATATW++** \| **++waTATAtACAcAyATwt** \| \|; coexpression\_cluster3\_8\_m24\_cluster3\_8\_motif24\_m \| cluster3\_8\_motif24\_m \| cluster\_2 \| coexpression\_cluster3\_8 \| 16 \| 15.21 \| 125 \| **+++TATTTTTTTTTTT** \| **AAAAAAAAAAATA+++** \| \|; coexpression\_cluster3\_8\_m17\_cluster3\_8\_motif17\_r \| cluster3\_8\_motif17\_r \| cluster\_7 \| coexpression\_cluster3\_8 \| 10 \| 9.15 \| 104 \| **wwAGAATGtw** \| **WACATTCTWW** \| \|; coexpression\_cluster3\_8\_m23\_cluster3\_8\_motif23\_m \| cluster3\_8\_motif23\_m \| cluster\_5 \| coexpression\_cluster3\_8 \| 10 \| 7.06 \| 190 \| **AgTAGACAAr** \| **YTTGTCTACT** \| \|; coexpression\_cluster3\_8\_m27\_cluster3\_8\_motif27\_m \| cluster3\_8\_motif27\_m \| cluster\_3 \| coexpression\_cluster3\_8 \| 11 \| 10.75 \| 118 \| **+TATGCATAT+** \| **+ATATGCATA+** \| \|; coexpression\_cluster3\_8\_m21\_cluster3\_8\_motif21\_m \| cluster3\_8\_motif21\_m \| cluster\_3 \| coexpression\_cluster3\_8 \| 11 \| 10.59 \| 101 \| **GTATGTACAT+** \| **+ATGTACATAC** \| \|; coexpression\_cluster3\_8\_m29\_cluster3\_8\_motif29\_m \| cluster3\_8\_motif29\_m \| cluster\_4 \| coexpression\_cluster3\_8 \| 22 \| 10.90 \| 120 \| **wtswwttTGCACACACrhmvrg** \| **CYBKDYGTGTGTGCAAAWWSAW** \| \|; coexpression\_cluster3\_8\_m18\_cluster3\_8\_motif18\_r \| cluster3\_8\_motif18\_r \| cluster\_3 \| coexpression\_cluster3\_8 \| 11 \| 9.43 \| 226 \| **+taTGTACAta** \| **TATGTACATA+** \| \|; coexpression\_cluster3\_8\_m5\_cluster3\_8\_motif5\_r \| cluster3\_8\_motif5\_r \| cluster\_1 \| coexpression\_cluster3\_8 \| 19 \| 10.03 \| 2042 \| **++++waTATAtAyATAwa+** \| **+TWTATRTATATATW++++** \| \|; coexpression\_cluster3\_8\_m25\_cluster3\_8\_motif25\_m \| cluster3\_8\_motif25\_m \| cluster\_2 \| coexpression\_cluster3\_8 \| 16 \| 12.17 \| 342 \| **TTTTTTYTTTTTTTT+** \| **+AAAAaAaarAAaAAA** \| \|; coexpression\_cluster3\_8\_m14\_cluster3\_8\_motif14\_r \| cluster3\_8\_motif14\_r \| cluster\_2 \| coexpression\_cluster3\_8 \| 16 \| 9.11 \| 837 \| **+++wwcAAmAAAAww+** \| **+WWTTTTKTTGWW+++** \| \|; coexpression\_cluster3\_8\_m9\_cluster3\_8\_motif9\_r \| cluster3\_8\_motif9\_r \| cluster\_1 \| coexpression\_cluster3\_8 \| 19 \| 8.90 \| 821 \| **+++wwATACACACAwww++** \| **++WWWTGTGTGTATWW+++** \| \|; coexpression\_cluster3\_8\_m1\_cluster3\_8\_motif1\_r \| cluster3\_8\_motif1\_r \| cluster\_1 \| coexpression\_cluster3\_8 \| 19 \| 9.63 \| 851 \| **++++wwTATAgAcAAAAww** \| **WWTTTTGTCTATAWW++++** \| \|; coexpression\_cluster3\_8\_m3\_cluster3\_8\_motif3\_r \| cluster3\_8\_motif3\_r \| cluster\_1 \| coexpression\_cluster3\_8 \| 19 \| 9.10 \| 1318 \| **+++awATAyACAyATaw++** \| **++WTATRTGTRTATWT+++** \| \|; coexpression\_cluster3\_8\_m8\_cluster3\_8\_motif8\_r \| cluster3\_8\_motif8\_r \| cluster\_2 \| coexpression\_cluster3\_8 \| 16 \| 9.56 \| 3284 \| **+++wwAAAmAAAAww+** \| **+WWTTTTKTTTWW+++** \| \|; coexpression\_cluster3\_8\_m28\_cluster3\_8\_motif28\_m \| cluster3\_8\_motif28\_m \| cluster\_3 \| coexpression\_cluster3\_8 \| 11 \| 11.33 \| 61 \| **+TATGCACAyA** \| **TRTGTGCATA+** \| \|; coexpression\_cluster3\_8\_m12\_cluster3\_8\_motif12\_r \| cluster3\_8\_motif12\_r \| cluster\_1 \| coexpression\_cluster3\_8 \| 19 \| 8.95 \| 582 \| **++++WWTATAGACAATWWW** \| **wwwaTTGTcTATAww++++** \| \|; coexpression\_cluster3\_8\_m6\_cluster3\_8\_motif6\_r \| cluster3\_8\_motif6\_r \| cluster\_2 \| coexpression\_cluster3\_8 \| 16 \| 9.77 \| 1127 \| **WWAWAARAMAAAAWW+** \| **+wwTTTTkTyTTwTww** \| \|; coexpression\_cluster3\_8\_m4\_cluster3\_8\_motif4\_r \| cluster3\_8\_motif4\_r \| cluster\_1 \| coexpression\_cluster3\_8 \| 19 \| 10.17 \| 357 \| **++ATATGTGTGTGYATW++** \| **++watrCACACAcATat++** \| \|; coexpression\_cluster3\_8\_m11\_cluster3\_8\_motif11\_r \| cluster3\_8\_motif11\_r \| cluster\_1 \| coexpression\_cluster3\_8 \| 19 \| 9.47 \| 868 \| **++++wwTATAkACAAAwww** \| **WWWTTTGTMTATAWW++++** \| \|; coexpression\_cluster3\_8\_m30\_cluster3\_8\_motif30\_m \| cluster3\_8\_motif30\_m \| cluster\_8 \| coexpression\_cluster3\_8 \| 15 \| 10.82 \| 18 \| **kwtCATTTyCCCCyy** \| **RRGGGGRAAATGAWM** \| \|; coexpression\_cluster3\_8\_m26\_cluster3\_8\_motif26\_m \| cluster3\_8\_motif26\_m \| cluster\_4 \| coexpression\_cluster3\_8 \| 22 \| 12.34 \| 61 \| **TvtwdtgCRCACACACrwrkmw** \| **WKMYWYGTGTGTGYGCAHWABA** \| \|; coexpression\_cluster3\_8\_m2\_cluster3\_8\_motif2\_r \| cluster3\_8\_motif2\_r \| cluster\_2 \| coexpression\_cluster3\_8 \| 16 \| 9.91 \| 2485 \| **+wwAAArAmAAAAww+** \| **+WWTTTTKTYTTTWW+** \| \|; coexpression\_cluster3\_8\_m10\_cluster3\_8\_motif10\_r \| cluster3\_8\_motif10\_r \| cluster\_1 \| coexpression\_cluster3\_8 \| 19 \| 9.07 \| 746 \| **++++WWWTGTMTATATW++** \| **++waTATAkACAwww++++** \| \|; coexpression\_cluster3\_8\_m13\_cluster3\_8\_motif13\_r \| cluster3\_8\_motif13\_r \| cluster\_1 \| coexpression\_cluster3\_8 \| 19 \| 9.29 \| 574 \| **++++wwwrTAgACAAAAww** \| **WWTTTTGTCTAYWWW++++** \| \|; coexpression\_cluster3\_8\_m22\_cluster3\_8\_motif22\_m \| cluster3\_8\_motif22\_m \| cluster\_2 \| coexpression\_cluster3\_8 \| 16 \| 11.10 \| 168 \| **+++TTATTATTWTT++** \| **++AAwAATAATAA+++** \| \|; coexpression\_cluster3\_8\_m19\_cluster3\_8\_motif19\_m \| cluster3\_8\_motif19\_m \| cluster\_1 \| coexpression\_cluster3\_8 \| 19 \| 11.71 \| 416 \| **+++ATRYAYATrCAYAya+** \| **+TRTRTGYATRTRYAT+++** \| \|; coexpression\_cluster3\_8\_m7\_cluster3\_8\_motif7\_r \| cluster3\_8\_motif7\_r \| cluster\_1 \| coexpression\_cluster3\_8 \| 19 \| 9.24 \| 1178 \| **+++++WWTTARAMAAAAWW** \| **wwTTTTkTyTAAww+++++** \| \|; coexpression\_cluster3\_8\_m15\_cluster3\_8\_motif15\_r \| cluster3\_8\_motif15\_r \| cluster\_6 \| coexpression\_cluster3\_8 \| 10 \| 9.24 \| 222 \| **awACATGTwt** \| **AWACATGTWT** \| \| - Click on the column names to change the order of the data. - Write the name of one cluster, collection or a pattern in the *Search* window. **Heatmap View** Heatmap View Distance table \| PDF |

**Additional Files**

#### Exported files

| File | Description |
| --- | --- |
| **Pairwise comparison** | This table shows the pairwise comparison between all the input motifs using different metrics. This is the *compare-matrices* result. |
| **Matrix description** | This table shows information of each input motif. |
| **Cluster-wise Motif Files [zip]** | One TRANSFAC file is generated for each cluster including all the input motifs grouped in the corresponding cluster. |
| **Root motifs** | A file with the root matrix of each cluster. |
| **Motif collections per cluster** | A tab separated file containing the counts of motif of each collection on each cluster. |
| **Clusters** | A tab separated file containing the clusters and their correspondng motifs. |
| **Central motifs IDs** | A tab separated file containing the central motif, that one who is the most similar to all the others motif in the same cluster. |
| **Central motifs (in TRANSFAC format)** | A transfac file containing the central motif, that one who is the most similar to all the others motif within the same cluster. |
| **Input motifs** | The input motifs analyzed with matrix-clustering. |

- **References**
  1. Castro-Mondragon JA et al.  RSAT matrix-clustering: dynamic exploration and redundancy reduction of transcription factor binding motif collections. doi: 10.1093/nar/gkx314; Nucleic Acids Research (2017)
  2. **Last RSAT release:** Nguyen NTT et al. RSAT 2018: regulatory sequence analysis tools 20th anniversary. Nucleic Acids Research (2018).
- **Additional Information**
  - Jacques van Helden course
- **Contact**
  - Jaime Castro-Mondragon - ****
  - Morgane Thomas-Chollier - ****
  - Jacques van Helden - ****
