## Additional file 10 for "Transcriptome analysis reveals a novel DNA element that may interact with chromatin-associated proteins in *Plasmodium berghei* during erythrocytic development": matrix-clustering_SUMMARY_cluster3_9.html

matrix-clustering coexpression\_cluster3\_9

### RSAT - matrix-clustering result

##### Analysis: coexpression\_cluster3\_9 (08/09/2023 16:16)

##### Command

```
matrix-clustering  -v 1 -max_matrices 300 -matrix coexpression_cluster3_9 $RSAT/public_html/tmp/www-data/2023/09/08/matrix-clustering_2023-09-08.161614_08Wx62/matrix-clustering_query_matrices.transfac transfac -hclust_method average -calc sum -title coexpression_cluster3_9 -metric_build_tree Ncor -lth w 5 -lth cor 0.6 -lth Ncor 0.4 -quick -label_in_tree name -return json,heatmap -o $RSAT/public_html/tmp/www-data/2023/09/08/matrix-clustering_2023-09-08.161614_08Wx62/matrix-clustering
```

#### Results Summary

|  |  |  |  |  |  |
| --- | --- | --- | --- | --- | --- |
| Nb Input motifs | Nb Input collections | Nb Clusters Found | Download   Root motifs | Complete results [zip] | Parameters used Linkage method \| Similarity metric \| Thresholds to partition the tree |
| 25 | 1 | 9 | **Download** | Download | average \| Ncor \| **Ncor** = 0.4 **cor** = 0.6 |

|  |  |  |  |  |  |  |
| --- | --- | --- | --- | --- | --- | --- |
|  | **Logo Forest (dynamic browsing)**  **Logo Forest (rapid overview - low image quality)**  cluster\_1  cluster\_2  cluster\_3  cluster\_4  cluster\_5  cluster\_6  cluster\_7  cluster\_8  cluster\_9 |  |  |  |  |  |
|  | **Clusters Summary** Clusters Summary Table (Root Motif \| Root Motif (Reverse) \| Cluster ID \| # Motifs \| Motif\_Name::Collection \| Number of Motifs by Collection \| Root Motif (transfac Format)) \| | cluster\_1 | 8 | cluster3\_9\_motif16\_m::coexpression\_cluster3\_9:: cluster3\_9\_motif1\_r::coexpression\_cluster3\_9:: cluster3\_9\_motif2\_r::coexpression\_cluster3\_9:: cluster3\_9\_motif12\_r::coexpression\_cluster3\_9:: cluster3\_9\_motif5\_r::coexpression\_cluster3\_9:: cluster3\_9\_motif3\_r::coexpression\_cluster3\_9:: cluster3\_9\_motif6\_r::coexpression\_cluster3\_9:: cluster3\_9\_motif9\_r::coexpression\_cluster3\_9:: | coexpression\_cluster3\_9::8\_motifs | Show matrix |
| cluster\_2 \| 3 \| cluster3\_9\_motif24\_m::coexpression\_cluster3\_9:: cluster3\_9\_motif23\_m::coexpression\_cluster3\_9:: cluster3\_9\_motif4\_r::coexpression\_cluster3\_9:: \| coexpression\_cluster3\_9::3\_motifs \| Show matrix \| \| |  |  |  |  |  |  |
| cluster\_3 \| 6 \| cluster3\_9\_motif11\_r::coexpression\_cluster3\_9:: cluster3\_9\_motif7\_r::coexpression\_cluster3\_9:: cluster3\_9\_motif19\_m::coexpression\_cluster3\_9:: cluster3\_9\_motif20\_m::coexpression\_cluster3\_9:: cluster3\_9\_motif17\_m::coexpression\_cluster3\_9:: cluster3\_9\_motif21\_m::coexpression\_cluster3\_9:: \| coexpression\_cluster3\_9::6\_motifs \| Show matrix \| \| |  |  |  |  |  |  |
| cluster\_4 | 3 | cluster3\_9\_motif14\_m::coexpression\_cluster3\_9:: cluster3\_9\_motif15\_m::coexpression\_cluster3\_9:: cluster3\_9\_motif18\_m::coexpression\_cluster3\_9:: | coexpression\_cluster3\_9::3\_motifs | Show matrix | | |
| cluster\_5 | 1 | cluster3\_9\_motif13\_r::coexpression\_cluster3\_9 | coexpression\_cluster3\_9 : 1 motif | Show matrix | | |
| cluster\_6 | 1 | cluster3\_9\_motif8\_r::coexpression\_cluster3\_9 | coexpression\_cluster3\_9 : 1 motif | Show matrix | | |
| cluster\_7 | 1 | cluster3\_9\_motif10\_r::coexpression\_cluster3\_9 | coexpression\_cluster3\_9 : 1 motif | Show matrix | | |
| cluster\_8 | 1 | cluster3\_9\_motif22\_m::coexpression\_cluster3\_9 | coexpression\_cluster3\_9 : 1 motif | Show matrix | | |
| cluster\_9 | 1 | cluster3\_9\_motif25\_m::coexpression\_cluster3\_9 | coexpression\_cluster3\_9 : 1 motif | Show matrix | | |

- Note: by default the table displays the first 50 clusters. Click on the *Show X entries* button to display more (or less) clusters.
- Click on the column names to change the order of the data.
- Use the *Search* window to find your motif (s) of interest.

| **Individual Cluster View** Individual Cluster View **Show All** **Hide All** **cluster\_1** **cluster\_2** **cluster\_3** **cluster\_4** **cluster\_5** **cluster\_6** **cluster\_7** **cluster\_8** **cluster\_9** **Display Logo Trees** **cluster\_1** **cluster\_2** **cluster\_3** **cluster\_4** **cluster\_5** **cluster\_6** **cluster\_7** **cluster\_8** **cluster\_9** **Display Branch-Motifs** **cluster\_1** Branch Motifs \| \| \| \| \| \| \|; Node \| Consensus \| Logo \| Logo (Reverse) \| Collections \| Matrix (Transfac format) \| IC \| Nb sites; Node 1 \| **waAAAtAgCTAGCTAktwwww--** \| \| \| coexpression\_cluster3\_9::2\_motifs \| PSSM \| 11.92 \| 144; Node 2 \| **--wwwtAgCTAGCTATtTtw---** \| \| \| coexpression\_cluster3\_9::2\_motifs \| PSSM \| 10.66 \| 210; Node 3 \| **wwAAAtAgCTAGCTAtttwww--** \| \| \| coexpression\_cluster3\_9::3\_motifs \| PSSM \| 11.42 \| 272; Node 4 \| **-wwwwTAgCTAGCTATtTtw---** \| \| \| coexpression\_cluster3\_9::3\_motifs \| PSSM \| 10.71 \| 337; Node 5 \| **wwwwwTAgCTAGCTATtTwww--** \| \| \| coexpression\_cluster3\_9::6\_motifs \| PSSM \| 10.84 \| 609; Node 6 \| **wwwwwTAgCTAGCTAttTwwwww** \| \| \| coexpression\_cluster3\_9::7\_motifs \| PSSM \| 11.19 \| 635; Node 7 \| **wwwwwtAgCTAGCTAttTwwwww** \| \| \| coexpression\_cluster3\_9::8\_motifs \| PSSM \| 11.23 \| 683 **cluster\_2** Branch Motifs \| \| \| \| \| \| \|; Node \| Consensus \| Logo \| Logo (Reverse) \| Collections \| Matrix (Transfac format) \| IC \| Nb sites; Node 1 \| **-----wwtGTTCATAAww** \| \| \| coexpression\_cluster3\_9::2\_motifs \| PSSM \| 9.21 \| 289; Node 2 \| **kymTGwwtGTTCATAAww** \| \| \| coexpression\_cluster3\_9::3\_motifs \| PSSM \| 13.60 \| 306 **cluster\_3** Branch Motifs \| \| \| \| \| \| \|; Node \| Consensus \| Logo \| Logo (Reverse) \| Collections \| Matrix (Transfac format) \| IC \| Nb sites; Node 1 \| **-wwAAAAAAAAww--** \| \| \| coexpression\_cluster3\_9::2\_motifs \| PSSM \| 8.66 \| 6119; Node 2 \| **rAAAAAAAArAAAAA** \| \| \| coexpression\_cluster3\_9::2\_motifs \| PSSM \| 12.97 \| 375; Node 3 \| **-AAwAAAAATAAWAw** \| \| \| coexpression\_cluster3\_9::2\_motifs \| PSSM \| 13.36 \| 100; Node 4 \| **rAAAAAAAArAAAAA** \| \| \| coexpression\_cluster3\_9::4\_motifs \| PSSM \| 12.12 \| 475; Node 5 \| **rwaAAAAAAAAwwAA** \| \| \| coexpression\_cluster3\_9::6\_motifs \| PSSM \| 10.75 \| 6594 **cluster\_4** Branch Motifs \| \| \| \| \| \| \|; Node \| Consensus \| Logo \| Logo (Reverse) \| Collections \| Matrix (Transfac format) \| IC \| Nb sites; Node 1 \| **ATATATRTATryATRT** \| \| \| coexpression\_cluster3\_9::2\_motifs \| PSSM \| 12.77 \| 271; Node 2 \| **ATATATaTATryATRT** \| \| \| coexpression\_cluster3\_9::3\_motifs \| PSSM \| 12.58 \| 375 **cluster\_5** Branch Motifs \| \| \| \| \| \| \|; Node \| Consensus \| Logo \| Logo (Reverse) \| Collections \| Matrix (Transfac format) \| IC \| Nb sites; Singleton \| **wayAGCCAAAww** \| \| \| coexpression\_cluster3\_9 : 1 motif \| PSSM \| 9.30 \| 58 **cluster\_6** Branch Motifs \| \| \| \| \| \| \|; Node \| Consensus \| Logo \| Logo (Reverse) \| Collections \| Matrix (Transfac format) \| IC \| Nb sites; Singleton \| **wwAAGCTACww** \| \| \| coexpression\_cluster3\_9 : 1 motif \| PSSM \| 7.24 \| 156 **cluster\_7** Branch Motifs \| \| \| \| \| \| \|; Node \| Consensus \| Logo \| Logo (Reverse) \| Collections \| Matrix (Transfac format) \| IC \| Nb sites; Singleton \| **wkGATATCmw** \| \| \| coexpression\_cluster3\_9 : 1 motif \| PSSM \| 8.96 \| 50 **cluster\_8** Branch Motifs \| \| \| \| \| \| \|; Node \| Consensus \| Logo \| Logo (Reverse) \| Collections \| Matrix (Transfac format) \| IC \| Nb sites; Singleton \| **aGttGC** \| \| \| coexpression\_cluster3\_9 : 1 motif \| PSSM \| 7.64 \| 51 **cluster\_9** Branch Motifs \| \| \| \| \| \| \|; Node \| Consensus \| Logo \| Logo (Reverse) \| Collections \| Matrix (Transfac format) \| IC \| Nb sites; Singleton \| **SSCSCCCC** \| \| \| coexpression\_cluster3\_9 : 1 motif \| PSSM \| 6.83 \| 17 - **Each cluster has a different color.** - **Click on the upper buttons to display separately the information of each cluster.** - **Click on the circles at each tree to display its corresponding merged motifs.** Definitions - **Logo tree.** This results shows the hierarchical tree with its logo alignment of one cluster. The logos are shown in Forward (left logo) and Reverse (right logo) orientation. - **Branch-motifs.** On the trees are displayed the branch number that is the number in which the motifs were incorporated in the tree. The Branch-Motifs table shows the logo in both orientations and the link to the file in TRANSFAC format with the branch-motif. |
|  | **Individual Motif View** Individual Motif View Motif id \| Motif name \| Cluster \| Collection \| Width \| IC \| Number of sites \| Consensus \| Consensus (Rev) \| Logo \| Logo (Rev); coexpression\_cluster3\_9\_m17\_cluster3\_9\_motif17\_m \| cluster3\_9\_motif17\_m \| cluster\_3 \| coexpression\_cluster3\_9 \| 15 \| 15.20 \| 65 \| **+AAAAAAAATAATAA** \| **TTATTATTTTTTTT+** \| \|; coexpression\_cluster3\_9\_m15\_cluster3\_9\_motif15\_m \| cluster3\_9\_motif15\_m \| cluster\_4 \| coexpression\_cluster3\_9 \| 16 \| 13.74 \| 63 \| **+TATRTATATGCATAT** \| **ATATGCATATAyATA+** \| \|; coexpression\_cluster3\_9\_m6\_cluster3\_9\_motif6\_r \| cluster3\_9\_motif6\_r \| cluster\_1 \| coexpression\_cluster3\_9 \| 23 \| 11.03 \| 100 \| **++WWWTAGCTAGCTATTTTW+++** \| **+++waAaAtAGCTAGCTAwww++** \| \|; coexpression\_cluster3\_9\_m22\_cluster3\_9\_motif22\_m \| cluster3\_9\_motif22\_m \| cluster\_8 \| coexpression\_cluster3\_9 \| 6 \| 7.64 \| 51 \| **AGTTGC** \| **GCAACT** \| \|; coexpression\_cluster3\_9\_m4\_cluster3\_9\_motif4\_r \| cluster3\_9\_motif4\_r \| cluster\_2 \| coexpression\_cluster3\_9 \| 18 \| 9.07 \| 245 \| **+++++wwtGTTCATAAww** \| **WWTTATGAACAWW+++++** \| \|; coexpression\_cluster3\_9\_m10\_cluster3\_9\_motif10\_r \| cluster3\_9\_motif10\_r \| cluster\_7 \| coexpression\_cluster3\_9 \| 10 \| 8.96 \| 50 \| **wkGATATCmw** \| **WKGATATCMW** \| \|; coexpression\_cluster3\_9\_m12\_cluster3\_9\_motif12\_r \| cluster3\_9\_motif12\_r \| cluster\_1 \| coexpression\_cluster3\_9 \| 23 \| 11.78 \| 78 \| **WAAAATAGCTAGCTAKTWWWW++** \| **++wwwwamTAGCTAGcTaTtTtw** \| \|; coexpression\_cluster3\_9\_m21\_cluster3\_9\_motif21\_m \| cluster3\_9\_motif21\_m \| cluster\_3 \| coexpression\_cluster3\_9 \| 15 \| 14.71 \| 35 \| **+AATAAAAATAAAAT** \| **ATTTTATTTTTATT+** \| \|; coexpression\_cluster3\_9\_m11\_cluster3\_9\_motif11\_r \| cluster3\_9\_motif11\_r \| cluster\_3 \| coexpression\_cluster3\_9 \| 15 \| 9.06 \| 579 \| **+wwGArAAAAAww++** \| **++WWTTTTTYTCWW+** \| \|; coexpression\_cluster3\_9\_m23\_cluster3\_9\_motif23\_m \| cluster3\_9\_motif23\_m \| cluster\_2 \| coexpression\_cluster3\_9 \| 18 \| 11.16 \| 44 \| **+++++HTTGTTCATAAT+** \| **+ATTATGAACAad+++++** \| \|; coexpression\_cluster3\_9\_m5\_cluster3\_9\_motif5\_r \| cluster3\_9\_motif5\_r \| cluster\_1 \| coexpression\_cluster3\_9 \| 23 \| 12.24 \| 66 \| **AAAAATARCTAGCTAKTTWWW++** \| **++wwwAamTAGCTAGyTaTTTtt** \| \|; coexpression\_cluster3\_9\_m24\_cluster3\_9\_motif24\_m \| cluster3\_9\_motif24\_m \| cluster\_2 \| coexpression\_cluster3\_9 \| 18 \| 14.67 \| 17 \| **KYMTGAAYGTTCATA+++** \| **+++TATGaACrTTCAkrm** \| \|; coexpression\_cluster3\_9\_m8\_cluster3\_9\_motif8\_r \| cluster3\_9\_motif8\_r \| cluster\_6 \| coexpression\_cluster3\_9 \| 11 \| 7.24 \| 156 \| **wwAAGCTACww** \| **WWGTAGCTTWW** \| \|; coexpression\_cluster3\_9\_m1\_cluster3\_9\_motif1\_r \| cluster3\_9\_motif1\_r \| cluster\_1 \| coexpression\_cluster3\_9 \| 23 \| 12.63 \| 26 \| **+WWTTTRGMTAGCTAKCYAAAWW** \| **wwTTTrgmTAGCTAkcyAAAww+** \| \|; coexpression\_cluster3\_9\_m16\_cluster3\_9\_motif16\_m \| cluster3\_9\_motif16\_m \| cluster\_1 \| coexpression\_cluster3\_9 \| 23 \| 9.79 \| 48 \| **+++++TRGCTAGYTR++++++++** \| **++++++++yArCTAGCyA+++++** \| \|; coexpression\_cluster3\_9\_m7\_cluster3\_9\_motif7\_r \| cluster3\_9\_motif7\_r \| cluster\_3 \| coexpression\_cluster3\_9 \| 15 \| 8.95 \| 5540 \| **+wwAAAAAAAAww++** \| **++WWTTTTTTTTWW+** \| \|; coexpression\_cluster3\_9\_m9\_cluster3\_9\_motif9\_r \| cluster3\_9\_motif9\_r \| cluster\_1 \| coexpression\_cluster3\_9 \| 23 \| 10.81 \| 110 \| **++wwawAgCTAGCTATwTtw+++** \| **+++WAAWATAGCTAGCTWTWW++** \| \|; coexpression\_cluster3\_9\_m3\_cluster3\_9\_motif3\_r \| cluster3\_9\_motif3\_r \| cluster\_1 \| coexpression\_cluster3\_9 \| 23 \| 11.04 \| 127 \| **+WWTTTAGCTAGCTATWTWW+++** \| **+++wwAwAtAGcTAGCTAAAww+** \| \|; coexpression\_cluster3\_9\_m25\_cluster3\_9\_motif25\_m \| cluster3\_9\_motif25\_m \| cluster\_9 \| coexpression\_cluster3\_9 \| 8 \| 6.83 \| 17 \| **scCsyCCC** \| **GGGRSGGS** \| \|; coexpression\_cluster3\_9\_m14\_cluster3\_9\_motif14\_m \| cluster3\_9\_motif14\_m \| cluster\_4 \| coexpression\_cluster3\_9 \| 16 \| 11.82 \| 105 \| **+ATATAKATATA++++** \| **++++TATATmTATAT+** \| \|; coexpression\_cluster3\_9\_m19\_cluster3\_9\_motif19\_m \| cluster3\_9\_motif19\_m \| cluster\_3 \| coexpression\_cluster3\_9 \| 15 \| 13.04 \| 236 \| **rAAAAAAaarRAAAA** \| **TTTTYYTTTTTTTTY** \| \|; coexpression\_cluster3\_9\_m13\_cluster3\_9\_motif13\_r \| cluster3\_9\_motif13\_r \| cluster\_5 \| coexpression\_cluster3\_9 \| 12 \| 9.30 \| 58 \| **wayAGCCAAAww** \| **WWTTTGGCTRTW** \| \|; coexpression\_cluster3\_9\_m2\_cluster3\_9\_motif2\_r \| cluster3\_9\_motif2\_r \| cluster\_1 \| coexpression\_cluster3\_9 \| 23 \| 11.26 \| 128 \| **WWAAATAGCTAGCTATTTWW+++** \| **+++wwAaATAgcTAgcTATtTww** \| \|; coexpression\_cluster3\_9\_m18\_cluster3\_9\_motif18\_m \| cluster3\_9\_motif18\_m \| cluster\_4 \| coexpression\_cluster3\_9 \| 16 \| 12.34 \| 208 \| **ATATATRTATRYATR+** \| **+yATryATAyATATAT** \| \|; coexpression\_cluster3\_9\_m20\_cluster3\_9\_motif20\_m \| cluster3\_9\_motif20\_m \| cluster\_3 \| coexpression\_cluster3\_9 \| 15 \| 12.93 \| 139 \| **+AAAAAAAAGAAAA+** \| **+TTTTCTTTTTTTT+** \| \| - Click on the column names to change the order of the data. - Write the name of one cluster, collection or a pattern in the *Search* window. **Heatmap View** Heatmap View Distance table \| PDF |

**Additional Files**

#### Exported files

| File | Description |
| --- | --- |
| **Pairwise comparison** | This table shows the pairwise comparison between all the input motifs using different metrics. This is the *compare-matrices* result. |
| **Matrix description** | This table shows information of each input motif. |
| **Cluster-wise Motif Files [zip]** | One TRANSFAC file is generated for each cluster including all the input motifs grouped in the corresponding cluster. |
| **Root motifs** | A file with the root matrix of each cluster. |
| **Motif collections per cluster** | A tab separated file containing the counts of motif of each collection on each cluster. |
| **Clusters** | A tab separated file containing the clusters and their correspondng motifs. |
| **Central motifs IDs** | A tab separated file containing the central motif, that one who is the most similar to all the others motif in the same cluster. |
| **Central motifs (in TRANSFAC format)** | A transfac file containing the central motif, that one who is the most similar to all the others motif within the same cluster. |
| **Input motifs** | The input motifs analyzed with matrix-clustering. |

- **References**
  1. Castro-Mondragon JA et al.  RSAT matrix-clustering: dynamic exploration and redundancy reduction of transcription factor binding motif collections. doi: 10.1093/nar/gkx314; Nucleic Acids Research (2017)
  2. **Last RSAT release:** Nguyen NTT et al. RSAT 2018: regulatory sequence analysis tools 20th anniversary. Nucleic Acids Research (2018).
- **Additional Information**
  - Jacques van Helden course
- **Contact**
  - Jaime Castro-Mondragon - ****
  - Morgane Thomas-Chollier - ****
  - Jacques van Helden - ****
